# Metabolic pathway rerouting in *Paraburkholderia rhizoxinica* evolved long-overlooked derivatives of coenzyme F_420_

**DOI:** 10.1101/670455

**Authors:** Daniel Braga, Daniel Last, Mahmudul Hasan, Huijuan Guo, Daniel Leichnitz, Zerrin Uzum, Ingrid Richter, Felix Schalk, Christine Beemelmanns, Christian Hertweck, Gerald Lackner

**Affiliations:** Junior Research Group Synthetic Microbiology, Leibniz Institute for Natural Product Research and Infection Biology – Hans Knöll Institute, Beutenbergstr 11a, 07745 Jena, Germany; Junior Research Group Chemical Biology of Microbe-Host Interactions, Leibniz Institute for Natural Product Research and Infection Biology – Hans Knöll Institute, Beutenbergstr 11a, 07745 Jena, Germany; Department of Biomolecular Chemistry, Leibniz Institute for Natural Product Research and Infection Biology – Hans Knöll Institute, Beutenbergstr 11a, 07745 Jena, Germany; Friedrich Schiller University, Jena

**Author notes:** Corresponding author: Dr. Gerald Lackner, E-Mail. **Author contributions and conflict of interest** Daniel Braga performed research, analyzed data (molecular biology, malachite green assay, mass spectrometry) and contributed to writing the manuscript, Daniel Last performed research and analyzed data (structure elucidation, Fno assay, biogas plant studies), Mahmudul Hasan performed research (CofC/D enzyme assays), Huijuan Guo performed research and analyzed data (structure elucidation), Daniel Leichnitz performed research (chemical synthesis), Zerrin Uzum performed research (microscopy), Ingrid Richter performed research (microscopy), Felix Schalk performed research (*cofE* constructs), Christine Beemelmanns designed research, acquired funding, analyzed data (structure elucidation, synthesis) and edited the manuscript, Christian Hertweck designed research, acquired funding and edited the manuscript, Gerald Lackner designed the study, acquired funding and wrote the original manuscript. The authors declare no conflict of interest.

**Keywords:** Cofactors, natural products, biocatalysis, symbiosis, deazaflavin

## Abstract

Coenzyme F_420_ is a specialized redox cofactor with a highly negative redox potential. It supports biochemical processes like methanogenesis, degradation of xenobiotics or the biosynthesis of antibiotics. Although well-studied in methanogenic archaea and actinobacteria, not much is known about F_420_ in Gram-negative bacteria. Genome sequencing revealed F_420_ biosynthetic genes in the Gram-negative, endofungal bacterium *Paraburkholderia rhizoxinica*, a symbiont of phytopathogenic fungi. Fluorescence microscopy, high-resolution LC-MS, and structure elucidation by NMR demonstrated that the encoded pathway is active and yields unexpected derivatives of F_420_ (3PG-F_420_). Further analyses of a biogas-producing microbial community showed that these derivatives are more widespread in nature. Genetic and biochemical studies of their biosynthesis established that a specificity switch in the guanylyltransferase CofC re-programmed the pathway to start from 3-phospho-D-glycerate, suggesting a rerouting event during the evolution of F_420_ biosynthesis. Furthermore, the cofactor activity of 3PG-F_420_ was validated, thus opening up perspectives for its use in biocatalysis. The 3PG-F_420_ biosynthetic gene cluster is fully functional in *Escherichia coli*, enabling convenient production of the cofactor by fermentation.

## Introduction

Cofactors are essential for the catalytic power of many enzymes and thus play a key role in virtually all metabolic pathways. Knowledge of their catalytic functions and biosynthesis is highly important for the understanding of biochemical reactions as well as their application in biocatalysis and biotechnology. An important subclass comprises redox-cofactors that mediate electron transfer between molecules. The deazaflavin coenzyme F_420_ (Figure 1) is a specialized redox cofactor with a lower redox potential (−350 mV) than NAD (1). This feature makes F_420_ an ideal electron carrier between H_2_ and NAD(P) in methanogenesis and renders it a strong reducing agent for challenging reactions in biocatalysis (2). For instance, enzymatic processes involving F_420_ facilitate the degradation of pollutants like aromatic nitro compounds (3) or the carcinogen aflatoxin (4). Furthermore, F_420_-dependent enzymes are important for asymmetric ene reductions (5, 6). In actinomycetes, coenzyme F_420_ is involved in the biosynthesis of antibiotics like oxytetracycline (7), pyrrolobenzodiazepines (8), or thiopeptins (9). Additionally, F_420_ has attracted considerable interest as a fitness factor of the human pathogen *Mycobacterium tuberculosis*, being involved in nitrosative stress response (10) or prodrug-activation (11).

**Figure 1.**
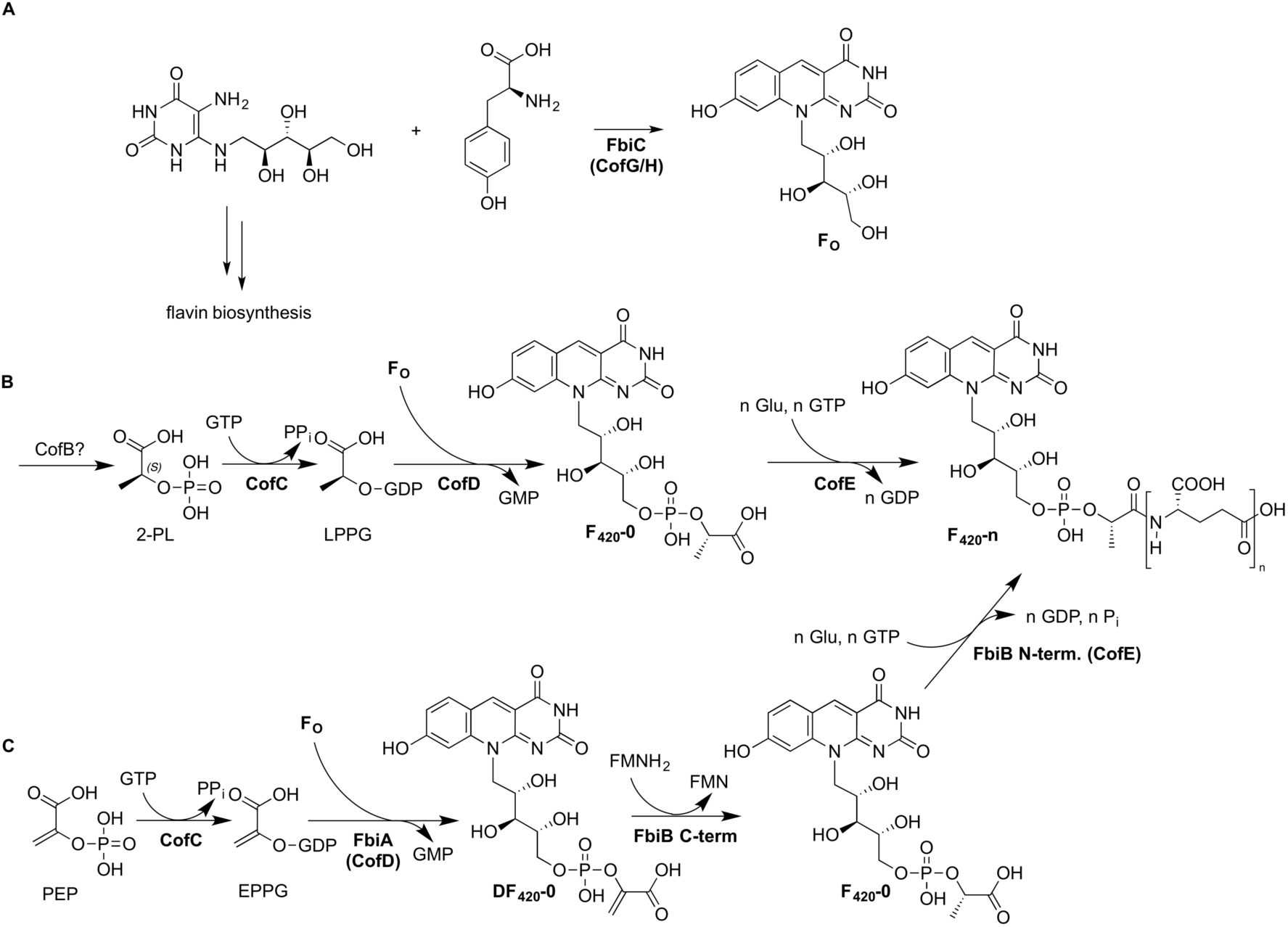
Biosynthesis of coenzyme F_420_ A) F_O_ synthase FbiC (in archaea: CofG/H) catalyzes formation of the deazaflavin ring from tyrosine and 5-amino-6-(ribitylamino)-uracil, an intermediate of riboflavin biosynthesis. B) Biosynthetic scheme of F_420_-n proposed for archaea. CofC and CofD catalyze the activation of 2-PL and transfer of the 2-PL moiety, respectively. CofE performs (oligo-)γ-glutamylation. The number of glutamate residues (n) varies depending on the organism. The origin of 2-PL is unknown. C) Biosynthesis of F_420_-n proposed for mycobacteria: CofC and CofD activate PEP resulting in DF_420_ formation. The C-terminal domain of FbiB reduces DF_420_ to F_420_. EPPG: enolpyruvyl-diphosphoguanosine, LPPG: lactyl-diphosphoguanosine, 2-PL: 2-phospho-L-lactate.

A key step during the biosynthesis of F_420_ is the formation of the deazaflavin fluorophore F_O_ (Figure 1A), a stable metabolic precursor of F_420_ originating from tyrosine and an intermediate of riboflavin biosynthesis. This chemically challenging step is catalyzed by the radical SAM enzyme complex CofG/H in archaea or the homologous dual-domain protein FbiC in actinobacteria (12). F_O_ is then further processed by CofC (EC 2.7.7.68) and CofD (EC 2.7.8.28). This pair of enzymes is responsible for the biosynthesis of the 2-phospho-L-lactate (2-PL) moiety (Figure 1B). Previous studies have shown that CofC from *Methanocaldococcus jannaschii* directly activates 2-phospho-L-lactate (2-PL) by guanylation (13) resulting in the formation of the short-lived metabolite lactyl-diphosphoguanosine (LPPG). CofD then forms F_420_-0 by transfer of the 2-PL moiety from LPPG to F_O_ (14). The enzymes producing 2-PL, however, have remained elusive in all F_420_ producers so far. Recently, Bashiri *et al.* proposed a revised biosynthetic pathway by demonstrating that phosphoenolpyruvate (PEP) instead of 2-PL can serve as a substrate of CofC in *Mycobacteria* (15). The resulting dehydro-F_420_ (DF_420_) is reduced to F_420_ by a flavin-dependent reductase domain present in the FbiB protein (Figure 1C). In mycobacteria, FbiB is a dual-domain protein consisting of a γ-glutamyl ligase domain (CofE-like, EC 6.3.2.31) and the C-terminal DF_420_ reductase domain (16). The γ-glutamyl ligase CofE finally decorates F_420_-0 with a varying number of (oligo-)γ-glutamate residues (17).

F_420_ is not ubiquitous in prokaryotes, but it is associated with certain phyla (18, 19). First discovered in methanogenic archaea (20, 21), it was extensively studied as a potential drug target of pathogenic mycobacteria (10) or as a cofactor enabling antibiotics biosynthesis in streptomycetes (2). Genome sequencing revealed that some Gram-negative bacteria have acquired F_420_ genes by horizontal transfer (19, 22). However, virtually nothing is known about the biosynthesis and role of F_420_ in these organisms. By genome mining, we found a biosynthetic gene cluster (BGC) homologous to those previously implicated in the biosynthesis of F_420_ in the endofungal bacterium *Paraburkholderia rhizoxinica* HKI 454 (Figure 2A and *Supporting Information* Table S3). This organism is an intracellular endosymbiont of the phytopathogenic fungus *Rhizopus microsporus* supplying its host with antimitotic toxins that act as virulence factors during infection of rice plants (23-25). We hypothesized that genes from Gram-negative bacteria related to F_420_ biosynthesis could facilitate F_420_ production in *E. coli* or could reveal novel biosynthetic routes towards this valuable molecule. Therefore, we set out to investigate if the BGC is active and if it can be refactored to produce F_420_ in *E. coli*.

**Figure 2.**
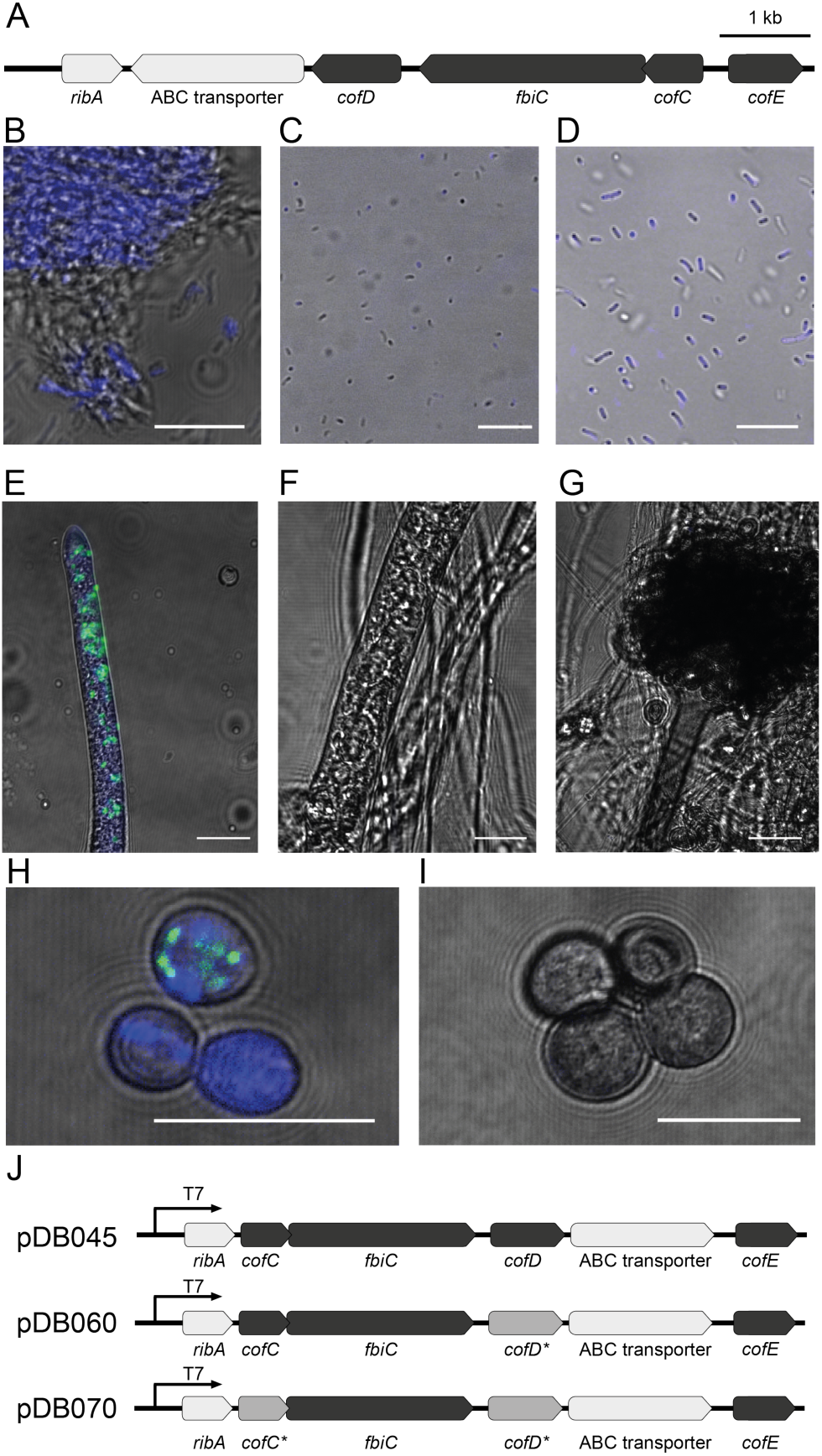
Deazaflavin biosynthesis in *P. rhizoxinica.* A) BGC of 3PG-F_420_. Core genes are shown in dark grey. B-I) Microscopy photographs depict fluorescence characteristic of deazaflavins in blue. B-D) axenic *M. smegmatis* (B), *P. rhizoxinica* (C), and *E. coli* / pDB045 (D). In *R. microsporus* ATCC 62417, deazaflavins are correlated to the presence of *P. rhizoxinica* symbionts (green, Syto9 staining) (E). No fluorescence was detected in cured ATCC 62417 mycelium (F) or in the naturally symbiont-free strain, CBS 344.29 (G). The same pattern was observed in spores of either wild-type ATCC 62417 (H) or CBS 344.29 (I). Scale bars represent 10 µm. J) Refactored versions of the BGC and corresponding plasmids for heterologous expression in *E. coli.* Asterisks mark genes from *M. jannaschii.*

Here, we show that *P. rhizoxinica* produces unexpected F_420_-derivatives (3PG-F_420_) both in symbiosis as well as in axenic culture. Heterologous expression and large-scale production in *E. coli* allowed for elucidation of their chemical structure. By comparative analyses, we discovered related metabolites in a biogas-producing microbial community, thus indicating their broader abundance and relevance. Enzyme assays showed that a switch in substrate specificity of CofC is responsible for the biosynthesis of 3PG-F_420_ and proved that it can serve as a substitute for F_420_ in biochemical reactions.

## Results and Discussion

To test whether *P. rhizoxinica* is capable of producing deazaflavins, we investigated axenic cultures of symbiont (*P. rhizoxinica*) and host (*R. microsporus*) as well as symbiotic cultures by fluorescence microscopy (Figures 2B-I and *Supporting Information* Figures S1-4). Indeed, symbiotic *P. rhizoxinica* and the cytosol of colonized mycelia emitted strong fluorescence characteristic of deazaflavins. Notably, even fluorescence of bacteria present inside of fungal spores was observed. Axenic fungi, however, showed no fluorescence, whereas only low signals were measured from axenic bacteria under the same conditions. To corroborate the results obtained by microscopy, we extracted metabolites from axenic *P. rhizoxinica*, axenic *R. microsporus* as well as fungal host containing endosymbionts and analyzed the extracts by LC-MS/MS. To our surprise, only F_O_ could be detected in axenic *P. rhizoxinica* and in the fungal host containing endosymbionts, but none of the expected F_420_-n species.

To further test the biosynthetic capacity of the full BGC, we refactored it to obtain a single operon under the control of a T7 promoter for heterologous expression yielding *E. coli/*pDB045 (Figure 2J). Examination of transformed bacteria by fluorescence microscopy revealed strong fluorescence as in the endofungal bacteria (Figure 2D), but LC-MS analyses again yielded only mass traces of F_O_ (mainly found in the culture supernatant), but not of F_420_-n. Since the impeded production could be attributed to nonfunctional proteins, we analyzed proteins by SDS-PAGE and found that CofD was poorly soluble in *E. coli*. Replacement of the *cofD* gene with the corresponding *M. jannaschii* homolog (14) provided soluble protein (*E. coli/*pDB060), however, again no trace of F_420_ could be detected. Therefore, we reexamined the metabolome of *E. coli/*pDB045 for characteristic MS/MS fragments derived from the F_O_ moiety (*m/z* 230.06, 346.10, 364.11). Surprisingly, the analysis revealed spectra with a similar fragmentation pattern, yet derived from precursor ions that had a mass shift of 15.995 compared to F_420_, indicating the presence of an additional oxygen atom (Figures 3A-B and Supporting Information Figures 5-8). Further analyses revealed an (oligo-)γ-glutamate series of the oxygenated compound suggesting these species are congeners. According to MS/MS fragmentation, the additional oxygen was present in the “phospholactyl” moiety of F_420_-n thus forming a “phosphoglyceryl” moiety. Extensive 1D- and 2D-NMR experiments (Figure 3C and Supporting Information *Section 2.2*) and comparison to classical F_420_ (20) corroborated that this moiety corresponds to 3-phosphoglycerate (3-PG). Therefore, we named the molecules 3PG-F_420_. This finding was unexpected because 2-phosphoglycerate is structurally more similar to PEP and 2-PL than 3-PG. Chemical degradation followed by chiral UHPLC-MS finally substantiated that the additional stereocenter of 3PG-F_420_ is *R*-configured (Figure 3D and Supporting Information Figure S51). Large-scale cultivation of *E. coli*/pDB045 also revealed traces of dehydro-F_420_ (DF_420_), but the yields were too low for NMR studies. The structure and occurrence of 3PG-F_420_ has not been reported before. To date, the only known derivatives of F_420_-n are factor F_390_-A and F_390_-G, 8-OH-AMP and 8-OH-GMP esters of F_420_, respectively (26). In methanogens, they are formed reversibly, e.g., during oxygen exposure, acting as a reporter compound for hydrogen starvation (27). In contrast, the modifications seen in 3PG-F_420_ are not temporary. Rather, 3PG-F_420_ seems to replace F_420_ as a natural deazaflavin-cofactor in *P. rhizoxinica*. At least in this organism, it does not coexist with classical F_420_. This situation is reminiscent of mycothiol, a specialized thiol cofactor that replaced glutathione in actinobacteria (28).

**Figure 3.**
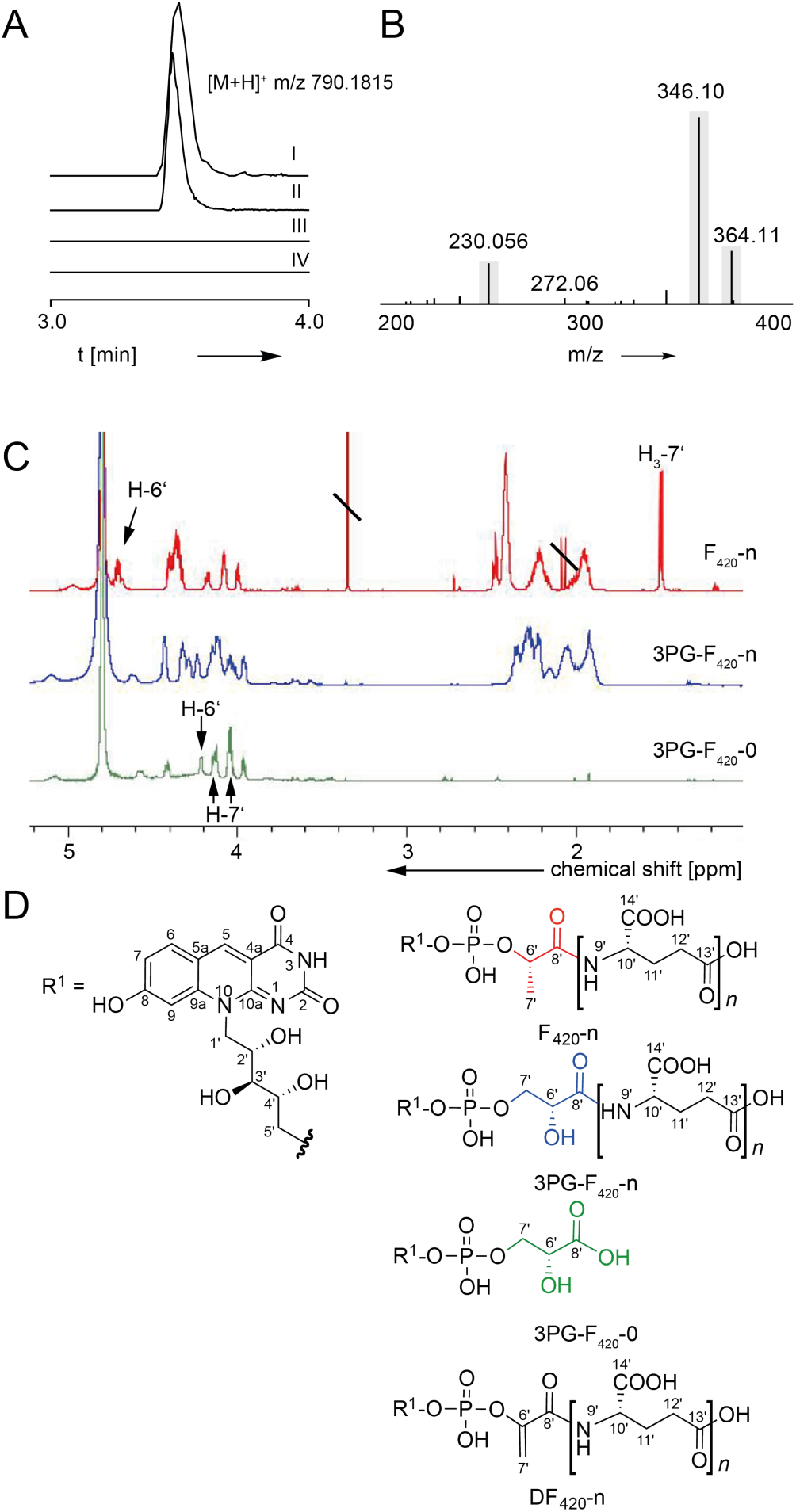
Chemical analysis of 3PG-F_420_. A) Extracted ion chromatograms of 3PG-F_420_-2 produced in *E. coli*. I: *E. coli* / pDB045, II: *cofD* exchanged by *M. jannaschii* homolog (pDB060), III: *cofD* and *cofC* exchanged by *M. jannaschii* homologs (pDB070) IV: empty vector (pETDuet). B) Excerpt of the MS/MS spectrum of 3PG-F_420_-2. Grey bars highlight m/z used for fragment ion search of F_420_ derivatives. C) ^1^H NMR comparison of F_420_-n (D_2_O), 3PG-F_420_-n (0.1% ND_3_ in D_2_O) and 3PG-F_420_-0 (0.1% ND_3_ in D_2_O) indicated the replacement of the lactyl moiety in F_420_ with a glyceryl moiety in 3PG-F_420_. D) Proposed structures of 3PG-F_420_-0, 3PG-F_420_-n, and DF_420_-n.

**Figure 4.**
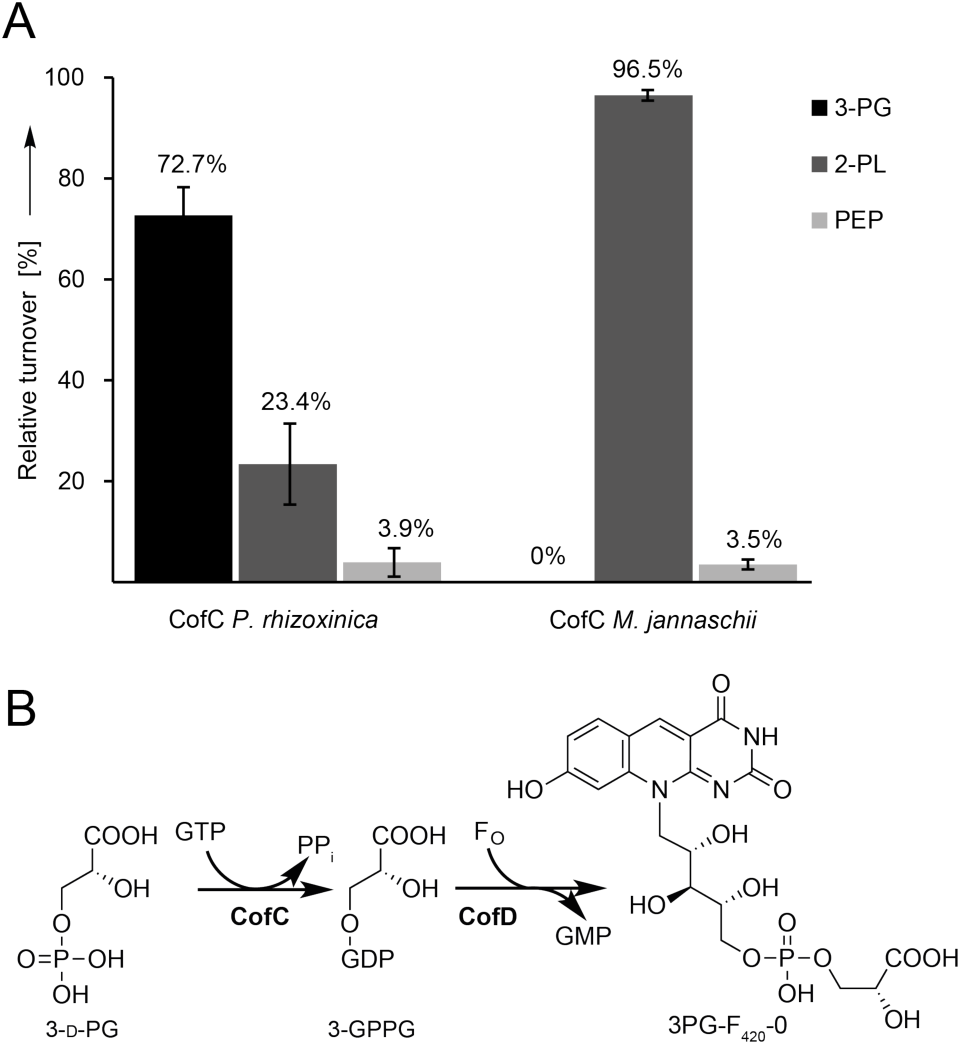
Combined CofC/D in-vitro assay. A) Relative turnover of substrates estimated from a substrate competition assays (D-3-PG, 2-PL, and PEP). CofC from *P. rhizoxinica* accepted 3-PG (72.7%), 2-PL (23.4%), and PEP (3.9%). CofC from *M. jannaschii* preferred 2-PL (96.5%), and PEP (3.5%). 3-PG was not turned over. CofD from *M. jannaschii* was used in all assays. Error bars represent the standard deviation (SD) of three independent biological replicates (N=3). B) Proposed model of 3PG-F_420_ biosynthesis. 3-GPPG: 3-(guanosine-5’-disphospho)-D-glycerate.

**Figure 5.**
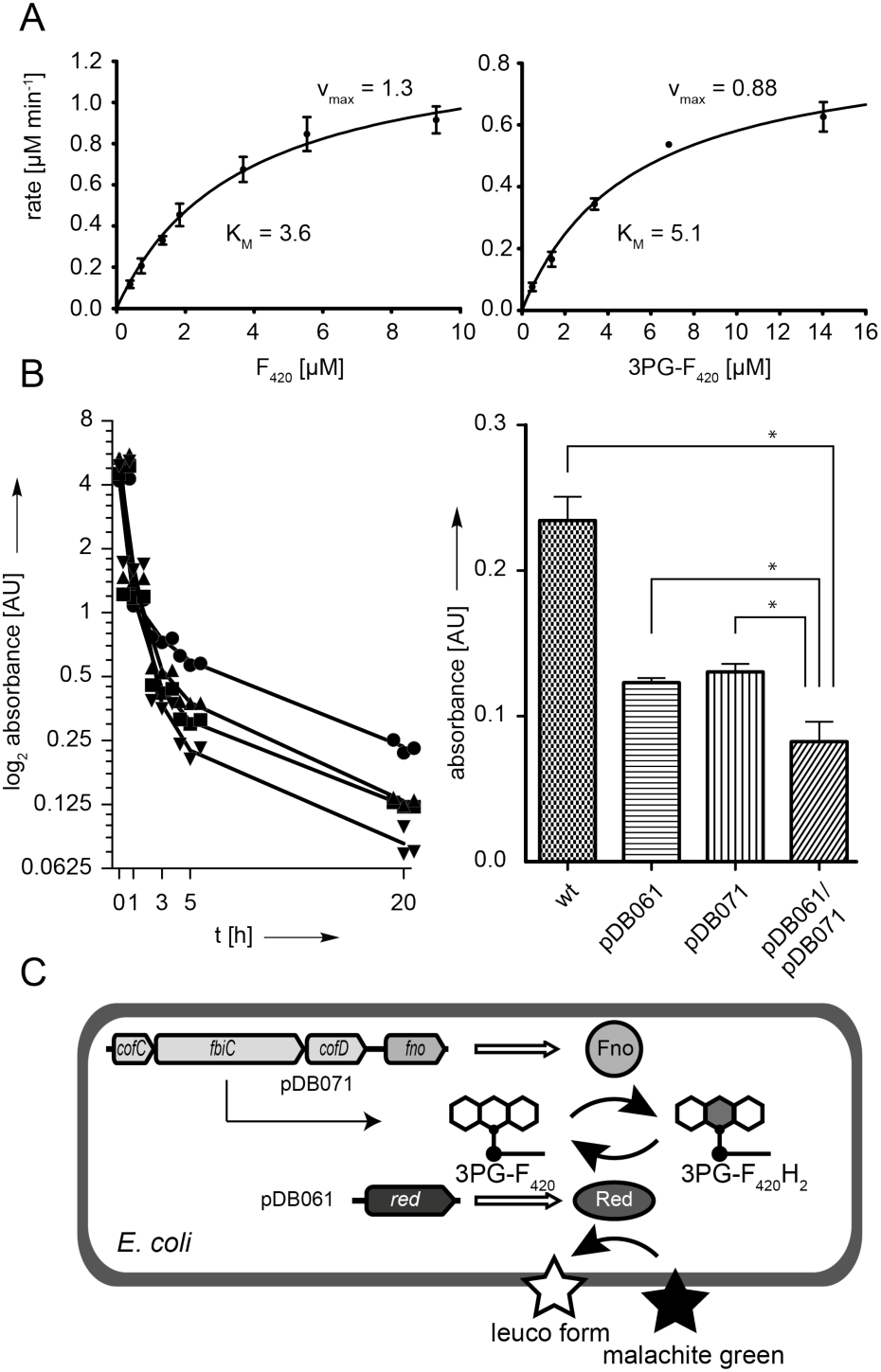
Cofactor function of 3PG-F_420_. A) Michaelis-Menten kinetics of Fno for F_420_ (left) and 3PG-F_420_ as substrates (right). Three biological replicates were used to determine parameters. K_M_ for F_420_ was 3.6 ± 0.7 µM (standard error). K_M_ for 3PG-F_420_ was 5.1 ± 1.0 µM. Error bars indicate standard deviation of replicates (N=3). B) In-vivo reduction of malachite green (absorbance: 618 nm) by the F_420_-dependent reductase MSMEG_5998. Fno was used to regenerate 3PG-F_420_H_2_. Left panel: Time course of the malachite green depletion assay. Right panel: Bar chart of residual malachite green after 20 h: wt: *E. coli* BL21(DE3), pDB061: *E. coli* producing MSMEG_5998, pDB071: *E. coli* producing 3PG-F_420_-0 + Fno. Exact means ± SD of biological triplicates were 0.234 ± 0.017 (wt), 0.124 ± 0.003 (pDB061), 0.169 ± 0.011 (pDB071), and 0.082 ± 0.0139 (pDB061/pDB071). An asterisk indicates statistical significance (one-way ANOVA, p<0.05, N=3). C) Engineered *E. coli* combining 3PG-F_420_, Fno and reductase MSMEG_5998 (*red*) for reduction of malachite green.

To investigate if 3PG-F_420_ is produced by wild-type *P. rhizoxinica*, we reanalyzed LC-MS data for the presence of corresponding mass signals. Indeed, 3PG-F_420_ species were found in samples containing bacteria (axenic culture, symbiosis), but not in symbiont-free host mycelia (Supporting Information Figures S9-14). In extracts of *P. rhizoxinica*, no traces of F_420_ and DF_420_ were detected. The presence of 3PG-F_420_ was restricted to the cell pellet, whereas F_O_ was abundant in culture supernatants. We thus conclude that the fluorescence observed in bacterial cells of *P. rhizoxinica* is derived from 3PG-F_420_ and F_O_. So far, there is only preliminary evidence for the occurrence of deazaflavin cofactors in a few Gram-negative bacteria, e.g., *Oligotropha carboxidivorans* and *Paracoccus dentrificans* (19) as well as in the uncultured, but biosynthetically highly prolific ‘*Candidatus* Entotheonella factor’ (22). The exact structure and function of F_420_ in most Gram-negative bacteria that harbor corresponding biosynthetic genes, however, is unknown. The fact that *P. rhizoxinica* produced 3PG-F_420_ under symbiotic cultivation conditions allows for the conclusion that it provides a fitness benefit in its natural habitat. Notably, none of the well-characterized F_420_-dependent enzyme families (4, 18) are encoded in the *P. rhizoxinica* genome according to BLAST and conserved domains searches, not even any of the widespread regeneration systems like Fno or F_420_-dependent glucose-6-phosphate dehydrogenase. Therefore, future investigations of 3PG-F_420_-producing organisms are likely to reveal novel enzymes, regeneration systems, and cellular pathways depending on this cofactor.

To assess if 3PG-F_420_ is restricted to fungal endosymbionts or if it might be more widespread in the environment, we examined *M. jannaschii* and *M. smegmatis* for the presence of any F_420_ congeners. Only F_420_-n was found from extracts of *M. jannaschii*, while F_420_-n and DF_420_-n were detected in extracts of *M. smegmatis*. (Supporting Information Figures S15-16). None of the 3PG-F_420_-derivatives were found in the reference organisms. Since methanogens are a common source of F_420_ in nature, we analyzed (two independent) sludge samples from a local biogas production plant. To our surprise, extraction of the microbial community present in the biogas-producing sludge followed by LC-MS/MS eluted, besides classical F_420_, a compound with identical retention time, exact mass and MS/MS fragmentation pattern as 3PG-F_420_ (Supporting Information Figure S17). As fluorescence and UV-based detection does usually not resolve classical F_420_ and 3PG-F_420,_ these derivatives might have been misidentified as F_420_ in the past. Since neither *P. rhizoxinica* nor its host *R. microsporu*s are able to grow under anaerobic and thermophilic (temperatures >42 °C) conditions, they can be excluded as the source of these cofactors. The high complexity of biogas-producing microbiomes (29) do not allow for an educated guess of the producer, although methanogens would be reasonable candidates.

In order to rationalize how the biosynthetic pathway was redirected to form 3PG-F_420_ instead of F_420_, we examined key steps of the biosynthesis more closely. We observed that production of 3PG-F_420_ was not abolished by the exchange of c*ofD* from *P. rhizoxinica* by *cofD* from *M. jannaschii* (plasmid pDB060). Hence, the phospholactyl transferase CofD could not be held accountable for the switch towards 3PG-F_420_. According to the existing biosynthetic model, the most plausible scenario was that CofC incorporated 3-phospho-D-glycerate (3-PG), an intermediate of glycolysis, instead of 2-PL to form 3PG-F_420_-0 and, to a minor extent, PEP to form DF_420_-0. To test this hypothesis, we exchanged *cofC* and *cofD* by the corresponding *M. jannaschii* homologs. The resulting strain *E. coli*/pDB070 (Figure 2J and Supporting Information Figure S18) produced neither 3PG-F_420_ nor F_420_ but traces of DF_420_. To further investigate the substrate specificity of CofC, we performed an in-vitro assay using CofC and CofD (13).

Genes *cofC* of *P. rhizoxinica* as well as *cofC* and *cofD* from *M. jannaschii* were cloned, corresponding proteins produced as hexahistidine fusions in *E. coli* and purified by metal affinity chromatography for in-vitro assays. The physiologically relevant isomers 2-phospho-D-glycerate (2-PG) and 3-phospho-D-glycerate (3-PG), as well as PEP and 2-PL, served as substrates. Reaction products were monitored by LC-MS. Indeed, when CofC from *P. rhizoxinica* was tested, the mass of 3PG-F_420_-0 appeared after reaction with D-3-PG eluting at the same retention time as the in-vivo product (Supporting Information Section 2.4). In addition, the formation of DF_420_-0 and F_420_-0 was detected, when the enzymes were incubated with PEP and 2-PL, respectively. In contrast, reaction with 2-PG yielded mass signals close to the noise level. Controls lacking CofC did not generate any of these products. In a direct substrate competition assay (Figure 4A), 3-PG was found to be the preferred substrate with a relative turnover of ca. 73% (2-PL: 23% PEP: 4%). This finding is in agreement with the structure of 3PG-F_420_ and the occurrence of DF_420_ as a minor biosynthetic product in *E. coli*. Note that CofC from *M. jannaschii* displayed a strong turnover of 2-PL (96.5%), weak turnover of PEP (3.5%) and no turnover of 3-PG. This finding supports the notion that the CofC of *P. rhizoxinica* has undergone a substrate specificity switch during evolution.

Recently, Bashiri *et al.* claimed that PEP is the substrate of CofC in prokaryotes (15). Our results confirm the hypothesis that PEP is the physiological substrate in mycobacteria, since we observed turnover of PEP by all CofC homologs tested. However, in contrast to Bashiri *et al.*, 2-PL was the best substrate of *M. jannaschii* CofC in our assay. Since Graupner and White detected significant amounts of 2-PL in methanogenic archaea and observed the conversion of lactate into 2-PL by isotope labeling (30), we conclude that 2-PL might still be a relevant substrate in archaea. From a phylogenetic perspective, our results suggest that multiple metabolic re-wiring events occurred in the evolution of F_420_ biosynthesis. While actinobacteria evolved the DF_420_ reductase (C-terminal domain of FbiB), archaea accomplished to produce the (unusual) metabolite 2-PL. Other organisms, as exemplified by *P. rhizoxinca*, rerouted the biosynthesis to the ubiquitous metabolite 3-PG.

To address the question if the γ-glutamyl ligase CofE adapted its substrate specificity to 3PG-F_420_, we individually co-expressed *cofE* genes from *P. rhizoxinica, M. jannaschii* and *M. smegmatis* (*fbiB*) together with a minimal BGC consisting of *fbiC, cofC*, and *cofD* in a two-plasmid system. Extraction of metabolites and LC-MS/MS revealed that all three CofE homologs elongated 3PG-F_420_-0 to oligo-glutamate chain lengths up to n=6 (Supporting Information Figures S63-65). Thus, we conclude that CofE does not act as an additional specificity filter during chain elongation of 3PG-F_420_.

The successful isolation of 3PG-F_420_ and reconstitution of its biosynthesis in *E. coli* motivated us to address the question of whether 3PG-F_420_ could substitute F_420_ in biocatalysis. To this end, we cloned a gene encoding Fno (F_420_:NADPH oxidoreductase), an enzyme that serves as a regeneration system for F_420_H_2_ using NADPH/H^+^ as an electron donor (31). We first examined if Fno can accept 3PG-F_420_ as a substrate. Indeed, we observed an efficient reduction of 3PG-F_420_ by recombinant Fno as mirrored by a rapid decrease of characteristic UV absorption. An examination of kinetic parameters (Figure 5A) revealed that the apparent K_M_ of Fno for F_420_ was 3.6 ± 0.7 µM. This value is similar to the reported K_M_ of 10 µM (32). Under identical assay conditions, the K_M_ for 3PG-F_420_ was only slightly higher (5.1 ± 1.0 µM). The v_max_ values were in a similar range as well (F_420_: 1.3 ± 0.2 µM min^-1^; 3PG-F_420_: 0.88 ± 0.07 µM min^-1^) pointing towards only a minor reduction of maximal turnover. Encouraged by the finding that 3PG-F_420_ can substitute F_420_, we aimed at an in-vivo application of the cofactor for malachite green reduction as a proof of principle. To this end, we combined the *fno* gene with a minimal BGC producing 3PG-F_420_-0 (Figure 5C and Supporting Information Figure S62) on a single vector (pDB071). Additionally, the F_420_-dependent malachite green reductase gene MSMEG_5998 (33) from *M. smegmatis* was cloned and expressed from a compatible vector backbone (pDB061). Finally, co-expression of all components in *E. coli* yielded a strain (pDB061/pDB071) that was able to decolorize malachite green significantly faster than control strains expressing the reductase or the cofactor alone (Figure 5B). Thus, we conclude that 3PG-F_420_ can substitute F_420_ as a redox cofactor in this case. The production of classical F_420_ and its use for biotransformations in *E. coli* has just recently been achieved in moderate yields using *Mycobacterium* genes including the DF_420_ reductase domain (15).

In summary, we discovered a derivative of the redox cofactor F_420_ that is produced by the Gram-negative endofungal bacterium *P. rhizoxinica.* We fully elucidated its chemical structure and show its potential cofactor function. Thus, our work is a solid basis to unveil unknown enzyme families and bioprocesses depending on 3PG-F_420_. Intriguingly, its presence in a biogas-producing digester suggests that the cofactor is more widespread in nature than expected. Furthermore, we could demonstrate that the guanylyltransferase CofC is responsible for the biosynthetic switch leading to the production of 3PG-F_420_. Our results thus significantly refine and extend the biosynthetic pathway models to deazaflavin cofactors in several phyla. Notably, the pathway discovered here, offered an alternative route to heterologous production and reconstitution of F_420_-dependent bioprocesses in *E. coli*. In recent years, there has been increasing interest in F_420_-dependent enzymes for biocatalysis (5, 6, 34, 35). Future applications will comprise for instance enantioselective biotransformations or the creation of a universal expression host for the production of antibiotics and other high-value compounds.

## Methods

Materials and methods are summarized in Supporting Information (Section 1).

## Acknowledgments

We thank Prof. Dr. Dina Grohmann and Dr. Ghader Bashiri for kindly providing *M. jannaschii* and *M. smegmatis* / *fbiABC*, respectively. We thank Biogas Jena GmbH and Co. KG for the kind donation of biogas plant samples. We thank Heike Heinecke for conducting NMR experiments. G.L. thanks the Deutsche Forschungsgemeinschaft (DFG Grant LA 4424/1-1) and the Carl Zeiss Foundation for funding. Financial support by the DFG (CRC 1127 ChemBioSys) to C.H. and C.B., and, BE-4799/2-1 to C.B., and Leibniz Award to C.H., by the ERC (MSCA-IF-EF-RI Project 794343, to I.R.) and the JSMC to Z.U. is gratefully acknowledged.

