## Supporting Information for "Metabolic pathway rerouting in *Paraburkholderia rhizoxinica* evolved long-overlooked derivatives of coenzyme F_420_"

**This PDF file includes:**

Supplementary text (see table of contents) including:
Figures S1 to S65
Tables S1 to S9
References for SI reference citations

**Table of contents:**

**1. Material and methods** **3**

1.1. Chemicals, microorganisms, and synthetic DNA 3

1.2. Microbial strains and cultivation conditions 3

1.3. Fluorescence microscopy 3

1.4. Plasmid design and cloning 4

1.5. Heterologous expression of BGCs in *Escherichia coli* 5

1.6. Extraction of deazaflavins from bacteria and fungi (small scale) 5

1.7. UHPLC-MS analyses of deazaflavins 6

1.8. HPLC of $F_O$ 6

1.9. Large-scale production and purification of 3PG-$F_{420}$ and $F_{420}$ 7

1.10. Extraction of deazaflavins from a biogas–producing microbial community 8

1.11. Acid hydrolysis of 3PG-$F_{420}$ and $F_{420}$ followed by chiral UPLC-MS analysis 8

1.12. Nuclear magnetic resonance (NMR) spectroscopy 9

1.13. Chemical synthesis of 2-phospho-L-lactate (2-PL) 9

1.14. Heterologous production and purification of CofC and CofD 9

1.15. In-vitro CofC/CofD combined enzyme assay 10

1.16. Heterologous production of Fno and enzymatic reduction of 3PG-$F_{420}$ 10

1.17. In-vivo reduction of malachite green 11

1.18. Fluorescence spectroscopy 11

**2. Results** **16**

2.1. Fluorescence Microscopy 16

2.2. Detection of 3PG-$F_{420}$ by LC-MS/MS 18

2.3. Structure elucidation of deazaflavins 30

2.3.1. Structure elucidation of classical $F_{420}$ (control) 30

2.3.2. Structure elucidation of 3PG-$F_{420}$-0 43

2.3.3. Structure elucidation of 3PG-$F_{420}$-n 51

2.3.4. Determination of the absolute stereochemistry of 3PG-$F_{420}$ 57

2.4. Combined CofC/D in-vitro enzyme assay 59

2.5. Fno and malachite green reduction assays 63

2.6. Plasmids sequences (Fasta format) 67

**3. References** **88**

### 1 Material and methods

#### 1.1 Chemicals, microorganisms, and synthetic DNA

Chemicals and media components were purchased from VWR, Roth, Sigma-Aldrich, Acros Organics and Alfa Aesar. All chemicals were of the highest purity available and the solvents were of mass spectrometry grade. Endonucleases and DNA polymerases were purchased from New England Biolabs. Synthetic genes (ThermoFisher Scientific, BioCat) were optimized for expression in *E. coli*.

#### 1.2 Microbial strains and cultivation conditions

*Rhizopus microsporus* van Tieghem ATCC 62417 harboring *Paraburkholderia rhizoxinica* HKI 454 (former name: *Burkholderia rhizoxinica*) and naturally endosymbiont-free *Rhizopus microsporus* var. *chinensis* CBS 344.29 were used in this study. Endobacteria were eliminated from *R. microsporus* ATCC 62417 by antibiotic treatment as described (1). Bacterial symbionts were isolated from the mycelium of *R. microsporus* ATCC 62417 as previously reported (2). For $F_{420}$ extraction, *R. microsporus* strains were cultured in 100 mL of NB medium (Merck Millipore, Darmstadt, Germany) in 500 mL baffled Erlenmeyer flasks at 30 °C, 110 rpm for 7 days. Isolates of *P. rhizoxinica* were cultured in 50 mL MGY M9 medium (0.12% yeast extract, 1% glycerol, 40 mM $K_2HPO_4$, 14 mM $KH_2PO_4$, 2.2 mM sodium citrate, 7.5 mM $(NH_4)_2SO_4$, and 0.8 mM $Mg_2SO_4$) in 300 mL baffled Erlenmeyer flasks at 30 °C, 110 rpm for 7 days. *E. coli* was routinely grown on lysogeny broth (LB, 10 g $L^{-1}$ tryptone, 5 g $L^{-1}$ yeast extract, 10 g $L^{-1}$ NaCl) at 37 °C and 210 rpm. *E. coli* BL21 (DE3) (New England Biolabs) was used for heterologous gene expression. *Mycolicibacterium* (former name: *Mycobacterium*) *smegmatis* was grown on LB supplemented with 0.05% (v/v) tween 80. Whenever used for the production of coenzyme $F_{420}$, tween was substituted by sterile glass beads.

#### 1.3 Fluorescence microscopy

To visualize the fluorescence of deazaflavins, *R. microsporus* ATCC 62417, cured *R. microsporus* ATCC 62417, symbiont-free *R. microsporus* CBS 344.29, *M. smegmatis* (3 days old), *E. coli* BL21(DE3)/pETDuet, *E. coli* BL21(DE3)/pDB045, and axenic *P. rhizoxinica* (7 days old) were analyzed using a Zeiss LSM 710 Confocal Laser Scanning Microscope (CLSM, Zeiss, Oberkochen, Germany) with excitation/emission of 405/470 nm. To visualize the localization of *P. rhizoxinica* within the fungal hyphae, a piece of fresh *R. microsporus* ATCC 62417 mycelium was transferred to an Eppendorf tube containing 200 µL physiological saline. The bacterial cells were stained with 5 µM Syto 9 green-fluorescent nucleic acid stain (Invitrogen, USA). The samples were incubated in the dark for 5 min and then analyzed using a Zeiss LSM 710 CLSM at 480/500 nm.

##### 1.4 Plasmid design and cloning

Unless otherwise specified, cloning was based on DNA recombination following the Fast Cloning protocol (3). 50 µL-polymerase chain reactions were composed by 0.2 mM each dNTP, 3 mM $MgCl_2$, 0.2 µM each primer (Table S2), as well as Q5 DNA Polymerase (1 U) and its respective buffer. PCRs started with initial denaturation at 98 °C for 1 minute followed by 18 cycles at 98 °C for 10 s, X °C for 30 s, and 72 °C for X' minutes. A final elongation at 72 °C for two minutes ended the reaction. Annealing temperatures varied with the primers and followed recommendations of the primer manufacturer. Elongation times were calculated as 2 kb per minute. All plasmids created during this study are listed in Table S3. Full plasmid sequences are provided in section 2.6.

The biosynthetic gene cluster for coenzyme F420 from *P. rhizoxinica* HKI 454 (Table S1) was amplified from its genomic DNA and was cloned in two steps. First, ribA and cofE were individually amplified (primers oDB01 and oDB02, oDB05 and oDB06, respectively) and were assembled in pETDuet-1 (Merck) previously linearized with XbaI and BamHI. The resulting construct, pDB044 (6,840 bp), was generated from a tripartite assembly of the two genes and vector backbone using the infusion cloning kit (Takara Clonetech). The gene sequences encoding CofC, the ABC transporter, and CofD were amplified as a single fragment (primers oDB07 and oDB08). pDB044 was PCR-linearized with the primers FC_pDB044_FP and FC_pDB044_RP to receive the latter amplicon. The construct bearing the complete gene cluster for 3PG-$F_{420}$ was termed pDB045 (13,135 bp).

Plasmid pDB045 served as the base to have either *P. rhizoxinica cofD* or both *cofC* and *cofD* swapped by *M. jannaschii* homologous gene sequences. The *cofD* gene (MJ_RS06720) from *M. jannaschii* was obtained in the form of a gene string and amplified with the primers oDB023 and oDB024. pDB045 was amplified with the primers oDB021 and oDB022 in a manner the PCR product would lack *P. rhizoxinica cofD*. The resulting construct (pDB060, 13,045 bp), therefore, bore the BGC for $F_{420}$ with *P. rhizoxinica cofD* replaced by that from *M. jannaschi*. The same strategy was used to exchange *cofC*. This gene from *M. jannaschii* (MJ_0887) was also purchased as a gene string and amplified with the primers oDB128 and oDB069. pDB060 served as template for the amplification/linearization of the plasmid lacking *P. rhizoxinica cofC* using the primers oDB070 and oDB071. The construct obtained (pDB070, 13,136 bp) has, therefore, the BGC for $F_{420}$ from *P. rhizoxinica* with the genes *cofC* and *cofD* replaced by the homologs from *M. jannaschii*. To construct plasmids encoding $F_{420}$-dependent enzymes, the gene encoding $F_{420}$-dependent malachite green reductase (MSMEG_5998) was ordered as a gene string. The sequence was amplified (primers DB_FP_MBP_5998 and DB_RP_MBP_5998) and cloned in pMAL-c2x (New England Biolabs) linearized with the primers DB_FP_pMal-c2X and DB_RP_pMal-c2X. The resulting construct was termed pDB061 (7,155 bp). For Fno production ($F_{420}$:NADPH oxidoreductase from *Archaeoglobus fulgidus)*, the codon-optimized *fno* gene string (Fno, AFULGI_RS04660) was amplified by PCR (primers: FC_fno_ins_fw, FC_fno_ins_rv) and ligated to the pET28a backbone (primers: FC_fno_vec_fw, FC_fno_vec_rv) by Fast Cloning yielding plasmid pDL008. Plasmids intended to yield reduced 3PG-$F_{420}$ were built with the pCDFDuet-1 backbone (Merck) and bore the codon-optimized sequence of *fno* in the multiple cloning site (MCS) 1. The optimized *fno* sequence was amplified with the primers oDB084 and oDB122. The vector was linearized with the primers oDB081 and oDB123. FastCloning of both fragments yielded pDB065 (4,385 bp), which served as the vector to receive the gene sequence(s) from *P. rhizoxinica* encoding proteins for the production of deazaflavins in its MCS2. pDB065 was linearized with primers oDB105 and oDB106 to receive the trio *cofC*, *fbiC*, and *cofD*, amplified with primers oDB107 and oDB108. The construct was intended to yield reduced 3PG-$F_{420}$-0 (pDB071, 8,656 bp). For heterologous expression of *cofC* and *cofD*, *cofC* (*P. rhizoxinica*) was amplified by PCR using primers CofC-fw and CofC-rv from genomic DNA. The vector backbone of pACYCDuet was amplified using primers pACYCDuet-fw and pACYCDuet-rv. Both PCR products were joined by FastCloning yielding plasmid pFS03 encoding an N-terminal hexahistidine ($His_6$) fusion protein of CofC. The coding sequence of *cofD* (*M. jannaschii*) was obtained as synthetic gene string and PCR amplified using primers CofD-fw and CofD-rv. The backbone of pET28a was amplified by primers pET28a-fw and pET28a-rv and ligated with the insert by FastCloning yielding plasmid pFS04 encoding an N-terminal hexahistidine (His6) fusion protein $His_6$-CofD. For coexpression with *cofE* homologs, a minimal cluster containing cofC, fbiC and cofD from pDB045 was amplified using primers oDB115 and oDB129B. oDB081 and oDB123 primers were used to amplify vector pCDFDuet-1 resulting in plasmid pMH02 (8,085 bp). *CofE* from *Methanocaldococcus jannaschii* was obtained as synthetic gene cloned into pET28a between BamHI and HindIII *sites.* FbiB was obtained as codon-optimized string and amplified by primers fbiB_fw and fbiB_rv, *cofE* from *Paraburkholderia rhizoxinica* was amplified using primers cofE_fw and cofE_rv. Both PCR products were fast-cloned into pET28a (amplified by primers pET28a-fw and pET28a-rv). All *cofE* constructs were designed in frame with the N-terminal $His_6$-tag.

##### 1.5 Heterologous expression of BGCs in *Escherichia coli*

For the heterologous expression of the biosynthetic gene clusters, *E. coli* BL21 (DE3) transformed, e.g. with plasmid pDB045 was grown to saturation in LB (50 µg $mL^{-1}$ carbenicillin) at 37 °C and 210 rpm. An overnight pre-culture was used (1:100) to inoculate 200 mL of main culture and induced with 200 µM IPTG upon $OD_{600}$ of 0.7. Protein and metabolite production proceeded for additional 20 h in the same culture conditions.

#### 1.6 Extraction of deazaflavins from bacteria and fungi (small scale)

Metabolite extraction for both *E. coli* and *M. smegmatis* followed the same protocol: The biomass was pelleted by centrifugation (4 °C, 8,000 × *g*, 25 min), washed with $dH_2O$, suspended in -20°C MeOH (MeOH) and left in an ultrasonic homogenizer for 10 minutes. Cell debris was removed by centrifugation. The extract was collected and the solvent was removed by rotary evaporation. The dried extract was dissolved in 500 µl MeOH. If the produced deazaflavin(s) were to be used in an enzymatic assay, the dried extract was dissolved in $ddH_2O$ and subsequently loaded onto a C18 Chromabond cartridge (Macherey-Nagel). Samples were washed and eluted with 20% and 30% (v/v) $MeOH:H_2O$, respectively. Except for the steps of cell washing and lysis, solvent systems were acidified with 0.1% formic acid (v/v). Mycelium of *R. microsporus* was strained and resuspended in 10 mL ice-cold HPLC-grade MeOH (VWR Chemicals, Darmstadt, Germany), while *P. rhizoxinica* cultures were snap-frozen in liquid nitrogen and freeze-dried. Dry *P. rhizoxinica* cultures were then resuspended in 10 mL ice-cold MeOH. Samples were sonicated for 20 min (Sonorex RK100 Ultrasonic bath, Bandelin) and then shaken (250 rpm) for 1 h followed by centrifugation (10.000 rpm) for 15 min. The supernatant was filtered through a paper filter and the solvent was evaporated. Dry extracts were dissolved in 2 mL $H_2O$.

#### 1.7 UHPLC-MS analyses of deazaflavins

Ultra-high performance liquid chromatography coupled with mass spectrometry (UHPLC-HRMS) was performed on a Dionex Ultimate3000 system combined with a Q-Exactive Plus mass spectrometer (Thermo Scientific) with a heated electrospray ion source (HESI). Samples were separated by reverse-phase chromatography in a Luna Omega C18 column (100 x 2.1 mm, 1.6 µm, 100 Å, Phenomenex). $H_2O$ (A) or acetonitrile (B), both acidified with formic acid 0.1% (v/v), served as mobile phases. 5 µl of the bacterial extracts were submitted to a gradient as follows: 0 min, 5% B; 1 min, 15% B; 3 min, 25% B; 6 min, 40% B; 7-9 min, 97% B; 10 min, 5% B at a constant flow rate of 300 µL $min^{-1}$ and at 40 °C. The chromatographic eluent was ionized in positive mode. The MS was carried out within a range of m/z 350 – 1800 followed by a data-dependent $MS^2$ analysis. $MS^1$ measurements were set to a resolving power of 70,000, $MS^2$ experiments to 17,500 at m/z 200, isolation windows m/z 1.0, normalized collision energy (NCE) 30.

#### 1.8 HPLC of $F_O$

The purification of $F_O$ for subsequent biochemical assays was carried out by analytical HPLC on a Shimadzu system equipped with a Nucleodur 100-5 C8ec column (150 x 4.6 mm, 5 µm, 300 Å, Macherey-Nagel). Solvents were the same as described for LC-HRMS. 20 µl of the SPE-purified samples were submitted to a gradient as follows: 0-2.5 min, 15% B; 5 min, 22% B; 6-11 min, 99% B;

11.5-16.5 min, 15% B. Flow rate and column temperature were held constant at 1 mL $min^{-1}$ and 25 °C, respectively. Compounds were analyzed by a diode array- and a fluorescence detector arranged in tandem. Excitation and emission at 420 and 480 nm, respectively, were set up for the fluorescence-based detection.

##### 1.9 Large-scale production and purification of 3PG-$F_{420}$ and $F_{420}$

3PG-$F_{420}$ was produced from a total volume of 50 L of *E. coli* BL21 (DE3)/pDB045 grown in LB medium (100 µg/mL carbenicillin) in 2L flasks in several batches. At an $OD_{600nm}$ of 0.6, IPTG was added (0.2 mM) followed by 20 h of further incubation at 37 °C and 160 rpm. *E. coli* cells were harvested in batches by centrifugation (30 min at 3,100 × g), washed with Tris-HCl buffer (25mM, pH 7.4) and stored temporarily at -20°C. Combined *E. coli* pellets (213 g wet weight) were resuspended in Tris-HCl buffer (50 mM, 100 mM NaCl, pH 7.4), to a concentration of about 0.5 g/mL (wet weight). Subsequently, the suspensions were autoclaved for 15 min at 121°C to release 3PG-$F_{420}$. Extracts were centrifuged for 30 min at 3,100 × g, supernatants were further processed. To generate 3PG-$F_{420}$-0, the 3PG-$F_{420}$ species were digested by the addition of carboxypeptidase G from *Pseudomonas* sp. (5 units, Sigma-Aldrich) and zinc sulfate (0.2 mM final concentration), and subsequent incubation for 18 hours at 37 °C and 180 rpm. The reaction product was acidified to pH 3.5 with HCl and applied to Chromabond C18 SPE cartridge (10 g, Macherey-Nagel). Step-wise elution was performed with increasing shares of MeOH in $H_2O$ containing 0.25% formic acid. Fluorescence measurements of 1:50 diluted samples in sodium phosphate buffer (50 mM, pH 7.5) with FLUOstar Omega (BMG Labtech) indicated positive fractions. Solvent was evaporated to dryness under reduced pressure in a rotary evaporator. The residues were dissolved in sodium phosphate buffer (25 mM, pH 7.4). Next, anion-exchange chromatography of 3PG-$F_{420}$ was performed on a FPLC system (NGC Quest, Bio-Rad) equipped with a HiPrep QFF column (GE Healthcare) pre-equilibrated with sodium phosphate buffer (25 mM, pH 7.4). After loading, the column was subsequently washed with ten column volumes (CV) of the same buffer. Elution of 3PG-$F_{420}$ was achieved by a NaCl gradient during the elution phase: 0-2 CV: 0-150 mM NaCl, 2-18 CV: 150-700 mM NaCl, 18-20 CV: 700-1000 mM NaCl and monitored by UV/VIS absorption at 420 nm. Retention time depended on the length of the glutamic acid tail, e. g., 5-7 ≥ 450 mM NaCl. Fractions with absorption at 420 nm were applied to C18 SPE cartridges as described above. Finally, the desalted 3PG-$F_{420}$-0 containing FPLC fractions (A19–A22) were submitted to a semi-preparative HPLC equipped with a Phenomenex Luna C8(2) 250 mm ×10 mm to yield 3PG-$F_{420}$-0 (golden yellow solid, 2.62 mg, $t_R$ = 13.27 min) using the following gradient: 0–5 min, 20% B; 5–25 min, 20%–100% B; 25–30 min, 100% B (A: dd$H_2O$ + 0.1% formic acid; B: MeCN) with a flow rate of 2.0 mL/min. The desalted 3PG-$F_{420}$-n containing FPLC fractions (A28–B9) were submitted to semi-preparative HPLC equipped with a Phenomenex Luna C8(2) 250mm × 10 mm to yield 3PG-$F_{420}$-n (3.96 mg, $t_R$ = 15.03 min) using the following gradient: 0–5 min, 20% B; 5–20 min, 20%–60% B; 20–25 min, 60%–100% B; 25–30 min, 100% B (A: dd $H_2O$ + 0.1% formic acid; B: MeCN) with a flow rate of 2.0 mL/min.

Classical $F_{420}$ was produced in *M. smegmatis* mc$^2$4517 pYUBDuet-fbiABC as described previously (4). At the end of the cultivations, cells were washed with sodium phosphate buffer (25 mM, pH 7.4). *M. smegmatis* cells (135 g wet weight) were resuspended in sodium phosphate buffer (25 mM, pH 7.4), to about 0.5 g/mL. Subsequently, the suspensions were autoclaved for 15 min at 121°C. Extracts were centrifuged for 30 min at 3,100 x g, the supernatants were further processed.

Anion-exchange purification of $F_{420}$ was performed as described above. Fractions with absorption at 420 nm were applied to C18 SPE cartridges as described above. At this step, for instance, the purity of $F_{420}$-5-7 eluting at 30% MeOH was already estimated by NMR to be ≥ 95%.

##### 1.10 Extraction of deazaflavins from a biogas–producing microbial community

Samples were kindly provided by the Biogas Jena GmbH & Co. KG (Jena, Germany) two times and with an interval of three months. The substrates of the fermenters are about 30% goat manure and 70% mixed plant residues. Fermenting conditions are anaerobic at temperatures over 42 °C. For extraction of metabolites, 500 mL of MeOH was added to 65 g of the liquid portion of the sludge. The suspension was kept in an ultrasonic homogenizer for 30 minutes and stirred for another three hours followed by filtration through several paper filters and solvent evaporation to dryness under reduced pressure. The remaining solid (300 mg) was resuspended in 50 mL $H_2O$ containing 0.25% formic acid. Subsequently, deazaflavins were SPE-purified as described above on a 10 g Chromabond C18 cartridge (Macherey-Nagel). As high amounts of unknown polymers interfered with LC-MS analyses, these were almost entirely removed by upstream HPLC purification similar to the description of analytical HPLC in section 1.7.

##### 1.11 Acid hydrolysis of 3PG-$F_{420}$ and $F_{420}$ followed by chiral UPLC-MS analysis

3PG-$F_{420}$-0 (**5**, 1.45 mg yellow solid) was dissolved in 100 µL HCl (1 M) and kept at 100 °C for 12 h. Afterwards, the reaction was neutralized by NaOH (1 M), centrifuged, and submitted to chiral UHPLC-MS on a Shimadzu LCMS-2020 system (single quadrupole) using an Astec CHIROBIOTIC R (10 cm x 4.6 mm, 5 µm, Sigma) column at 40 °C. Scan range of the MS was set to *m*/*z* 70 to 1,000 with a scan speed of 5,000 u/s and event time of 0.25 s under positive and negative mode. DL temperature was set to 250 °C with an interface temperature of 350 °C and a heat block of 400 °C. The nebulizing gas flow was set to 1.5 L/min and dry gas flow to 15 L/min. UPLC-MS analysis was conducted using an isocratic condition: 0–16 min, 15% A/85% B (A: dd $H_2O$ with 33.3 mM $NH_4OAc$; B: MeCN) with a flow rate of 0.7 mL/min under negative selected ion monitor (SIM) mode to detect the *m*/*z* 105.3 $(M–H)^-$.

#### 1.12 Nuclear magnetic resonance (NMR) spectroscopy

All measurements were performed on a Bruker AVANCE II 300 MHz and 600 MHz spectrometer equipped with a Bruker Cryoplatform. The chemical shifts are reported in parts per million (ppm) relative to the solvent residual peak of $D_2O$ ($^1$H: 4.79 ppm, singlet) and DMSO-$d_6$ ($^1$H: 2.50 ppm, quintet; $^{13}$C: 39.52 ppm, heptet).

#### 1.13 Chemical synthesis of 2-phospho-l-lactate (2-PL)

Synthesis was performed as described in the literature (5). To a solution of benzyl (*S*)-2-hydroxypropanoate (750 mg, 4.16 mmol) in pyridine (5.55 mL) was dropped at 0°C diphenyl phosphoryl chloride (905 µL, 4.37 mmol, 1.05 equiv.) and the suspension was stirred at 25 °C for 24 h. The reaction was stopped by addition of $H_2O$ (1 mL) and the volatiles removed *in vacuo*. The residue was dissolved in toluene (7 mL) and washed with $H_2O$, 1 M aq. HCl, sat. $NaHCO_3$-solution and brine successively. The organic phase was dried over $MgSO_4$, filtered and the volatiles removed *in vacuo* to yield 1.54 g of colorless oil, which was directly used in the next step. The oil was dissolved in EtOH (41.7 mL) and $PtO_2$ (82.6 mg, 636 µmol) was added. Hydrogen gas from a balloon was bubbled through the stirred black suspension for 3 h, then the mixture was left stirring under hydrogen atmosphere for additional 22 h with renewal of the atmosphere, when the balloon was depleted. After filtration of the mixture through a pad of Celite the volatiles were removed in vacuo to yield a greenish highly viscous oil. 2-PL was further purified via Chromabond C18 SPE cartridge (Macherey-Nagel). The flow-through was lyophilized and yielded 629 mg of 2-PL as colorless oil (47% yield over two steps). The analytical data was consistent with the literature.

**$^1$H NMR** (300 MHz, $D_2O$) δ 4.71 (dq, *J* = 7.5, 6.9 Hz, 1H, 2-H), 1.44 (t, *J* = 6.9 Hz, 3H, 3-H) ppm.
**$^{13}$C NMR** (75 MHz, $D_2O$) δ 175.6 (d, C-1), 70.6 (dd, C-2), 18.8 (dq, C-3) ppm.

#### 1.14 Heterologous production and purification of CofC and CofD

*E. coli* BL21 (DE3) was individually transformed with pFS03 (encoding His$_6$-CofC from *P. rhizoxinica*) or pFS04 (His$_6$-CofD from *M. jannaschii)*. A culture of 100 mL (50 µg/mL kanamycin or 25 µg/mL chloramphenicol) was grown at 37 °C and 180 rpm until $OD_{600}$ = 0.7 and induced with 1 mM isopropyl β-D-1-thiogalactopyranoside (IPTG). Protein production followed for 20 h at 17 °C and 180 rpm. Harvesting and purification steps of proteins were conducted at 4 °C. Cells were pelleted for 25 min at 8000 × g and suspended in lysis buffer ($NaH_2PO_4$ 50 mM and NaCl 300 mM, 20 mM imidazole). Pulsed sonication was applied for 60 s with 10 s pause to lyse cells (SONOPLUS Ultrasound-

Homogenizer HD 2070, BANDELIN). Cell debris was separated by centrifugation for 30 min at 11000 × g and the hexa-histidine fusion proteins were purified from the supernatant by metal-affinity chromatography (HisPur$^{TM}$ Ni-NTA Resin column, Thermo Scientific) using lysis buffer with increasing concentration of imidazole (elution by 500 mM imidazole). Re-buffering of proteins was performed in a PD10 column (Sephadex$^{TM}$ G-25 M, GE Healthcare).

**1.15 In-vitro CofC/CofD combined enzyme assay**

Enzyme assays combining CofC and CofD to assess their substrate specificity were performed analogously to Grochowski *et al.* (6) and product formation of $F_{420}$-0 derivatives was monitored by LC-MS. Reactions were carried in 50 µl-reaction mixtures of 100 mM HEPES buffer (pH 7.4), 2 mm GTP, 2 mM $MgCl_2$, 50 ng Fo (0.14 nM), 34 µm CofD (WP_010870769), and 0.5 mM of substrates (either 3-phospho-D-glyceric acid, 2-phospho-D-glyceric acid, phosphoenolpyruvic acid or 2-phospho-L-lactate). The reactions were initiated upon addition of 26 µM CofC (WP_013435883). For the competition assay, several substrates were used at the same time. Reactions were stopped after 0, 20, 40 and 60 min to monitor the rate of product formation. All reactions were carried out at 37 °C and 300 rpm. Reactions were quenched by addition of one volume of acetonitrile and 10 µl of formic acid (20 %) and centrifuged at 13000 × g for 30 minutes to pellet precipitates. Supernatants were analyzed by LC-MS. The LC method was the same as stated using a Luna® Omega 1.6 µm C18 100 Å and size of 100 x 2.1 mm (Phenomenex®). ESI-MS analysis was performed with a resolution of 70,000 and restricted to a mass range of 350 to 600 m/z to enhance sensitivity. Individual XICs ($[M+H]^+$) were extracted from raw LC/MS data (3PG-$F_{420}$-0: m/z 532.09364-532.09896, $F_{420}$-0: m/z 516.09881-516.10397, $DF_{420}$-0: m/z 514.08317-514.08831. Quantification was achieved by integration of specific extracted ion chromatograms (XICs). Area under curve (AUC) for each product was then normalized to the area corresponding to $F_0$ at time point zero. The normalized area was plotted over time and slopes of product formation were determined from the near-linear time range of the assay (20 min to 40 min) to mirror substrate turnover. Relative percentages of substrate turnover were calculated, their mean and standard deviation (SD) were derived from three independent biological replicates (N=3). Error bars of bar charts represent the SD.

**16 Heterologous production of Fno and enzymatic reduction of 3PG-$F_{420}$ (kinetics)**

For Fno production, *E. coli* BL21 (DE3) / pDL008 was grown in 150 mL of LB added with kanamycin (50 µg/mL) at 37 °C and 180 rpm to an $OD_{600}$ of 0.45. Induction with 1 mM IPTG was followed by 20 h of further incubation at 20 °C and 180 rpm. The pellet was washed with 20 mL potassium phosphate buffer (1.5 M, pH 8.0), resuspended in 5 mL of the same buffer and lysed on ice by 3 cycles of 1 min sonication (SONOPLUS Ultrasound-Homogenizer HD 2070, BANDELIN) with 30 s between each cycle. After centrifugation (15 min at 3,100 × g), the supernatant was incubated for 30 min at 90 °C to denature most host proteins. This was followed by another centrifugation step (15 min at 3,100 × g) and one-to-four dilution with sodium phosphate buffer (50 mM, pH 7.5, 300 mM NaCl, 20 mM imidazole). Subsequently, metal-affinity chromatography (HisPur$^{TM}$ Ni-NTA Resin column, Thermo Scientific) was performed. After sample loading, the column was washed with two column volumes of the same buffer. Elution of the protein was achieved with buffer containing 500 mM imidazole. Finally, the buffer was exchanged to 1 mL potassium phosphate buffer (50 mM, pH 6) via centrifugal filtration (10 kDa cut-off). Fno-mediated turnover of oxidized 3PG-$F_{420}$ (absorbing at 400 nm, $\varepsilon_{400}$=25.7 $mM^{-1}$ $cm^{-1}$) to reduced 3PG-$F_{420}H_2$ (non-absorbing) was measured by FLUOstar Omega reader (BMG Labtech). The assay contained 0-14 µM 3PG-$F_{420}$ or 0-9.3 µM $F_{420}$, 250 µM NADPH and 0.149 µg/mL Fno in potassium phosphate buffer (50 mM, pH 6). Initial reaction rates were obtained by linear regression and plotted against 3PG-$F_{420}$ or $F_{420}$ concentrations. Three biological replicates (N=3) were measured and combined in one analysis. Michaelis-Menten parameters ($v_{max}$ and $K_M$ and corresponding standard errors) were obtained by nonlinear regression using SigmaPlot 12 with Enzyme Kinetics Wizard. Error bars show the standard deviation of the initial reaction speeds of the three replicates.

##### 1.17 In-vivo reduction of malachite green

*E. coli* BL21 (DE3) transformed with pDB061, pDB071, or a combination of both was cultivated overnight as detailed. At this moment, gene expression was induced with 200 µM IPTG and the cultivation received 50 mg $L^{-1}$ malachite green as described in the literature (7). 1-mL aliquots were centrifuged and UV absorbance at 618 nm of supernatants was measured. The reduction of malachite green to leucomalachite green was indicated by the reduction of absorbance over time. Values were recorded from three independent experiments. All data represent mean ± SD. Statistical comparison was performed in GraphPad Prism version 5.0 for Windows by using one-way analysis of variance (ANOVA) followed by Tukey's multiple comparison test. The value of $p < 0.05$ was considered statistically significant.

##### 1.18 Fluorescence spectroscopy

Purified 3PG-$F_{420}$ and $F_{420}$ were dissolved in sodium phosphate buffer (50 mM, pH 7.5). Fluorescence spectra were recorded in a CLARIOstar reader (BMG Labtech).

**Table S1**: The 3PG-$F_{420}$ biosynthetic gene cluster from *P. rhizoxinica* HKI454 (GenBank accession number: NC_014722).

| Gene | Locus_tag | Proposed function |
|---|---|---|
| ribA | RBRH_RS08950 | GTP cyclohydrolase II |
| | RBRH_RS08955 | ABC transporter |
| cofD | RBRH_RS08960 | 2-phospho-L-lactate transferase |
| fbiC | RBRH_RS08965 | FO synthase |
| cofC | RBRH_RS08970 | 2-phospho-L-lactate guanylyltransferas |
| cofE | RBRH_RS08970 | coenzyme $F_{420}$-0: γ-L-glutamate ligase |

**Table S2**. Primers used in this study (nucleotide sequences).

| **Name** | **5' - 3' sequences** |
|---|---|
| cofC - fw | AGGTCGACAAGCTTGCGATGTCTCCTGTTTCTGTCGC |
| cofC - re | CTTAAGCATTATGCGGCCTAGATTTGGCATAGCACCCT |
| cofD - fw | AAATGGGTCGCGGATCCATGGCAAAGTATGTTGCGTTGTG |
| cofD - re | CGACGGAGCTCGAATTCTTATGCGGTGCTGTGCTGCA |
| cofE_fw | AAATGGGTCGCGGATCCATGACTGTATCCGCCATC |
| cofE_rv | CGACGGAGCTCGAATTCTTAGGCCATTTCCACCG |
| DB_pDB044_FP | AAACCAGCAAGAAGCATGACTGTATCC |
| DB_pDB044_RP | CCTCTACAATGACAACGGGCAC |
| DB_FP_MBP_5998 | GGATCGAGGGAAGGATTTCAGAA |
| DB_RP_MBP_5998 | AAGCTTGCCTGCAGGTCGAC |
| DB_FP_pMal-c2X | GTCGACCTGCAGGCAAGCTT |
| DB_RP_pMal-c2X | TGAAATCCTTCCCTCGATCC |
| fbiB_fw | AAATGGGTCGCGGATCCATGAGCGCAGCAGCAAATGCA |
| fbiB_rv | CGACGGAGCTCGAATTCTTATTTACGAACCAGCAGTTCATCGG |
| oDB08 | CATGCTTCTTGCTGGTTTAAACTTATTCAGGCTGCTTGTGTTGCAATG |
| oDB021 | TTTGTGGTAGCCTGTAAGGTTGATGGCTATACAGAAA |
| oDB022 | CTCAGAACGGTAATCACTATCTTCCTGCGTAAAGATC |
| oDB023 | GTGATTACCGTTCTGAGCG |
| oDB024 | TTACAGGCTACCACAAAATTCAATG |
| oDB069 | ATTCCTCAAGGTCGGCATTAACGACGTTTAACAAC |
| oDB070 | ACCATTGCATCTCATATAGGC |
| oDB071 | TGCCGACCTTGAGGAATTC |

| | |
|---|---|
| oDB081 | GGATCCGAATTCGAGCTC |
| oDB084 | AGCTCGAATTCGGATCCTCACAGAAATTTGATACCCAGTTCG |
| oDB085 | GGTACCCTCGAGTCTGGT |
| oDB088 | CCAGACTCGAGGGTACCTCATTGTTCCGCTTCACTCAAG |
| oDB105 | CTAATATACTAAGATGGGGAATTG |
| oDB106 | CGAGTCTGGTAAAGAAACC |
| oDB107 | CAATTCCCCATCTTAGTATATTAGGAGATGCAATGGTATGTCTCC |
| oDB108 | GGTTTCTTTACCAGACTCGTTATGCGGTGCTGTGCTGC |
| oDB115 | GAGCTCGAATTCGGATCCTTATGCGGTGCTGTGCTG |
| oDB122 | GTTTAACTTTAATAAGGAGATATACCATGCGTGTGGCTCTGCTG |
| oDB123 | GGTATATCTCCTTATTAAAGTTAAACA |
| oDB124 | ATGTATATCTCCTTCTTATACTTAACT |
| oDB125 | GTTAAGTATAAGAAGGAGATATACATATGATCCAAGAGGTCGAACG |
| oDB128 | GCCTATATGAGATGCAATGGTATGAACTGCGGCATCAAAATGA |
| oDB129 | GAGCTCGAATTCGGATCCTTATGCGGTGCTGTGCTG |
| oDB129B | TAATAAGGAGATATACCATGTCTCCTGTTTCTGTCGC |
| pACYCduet-fw | GCCGCATAATGCTTAAGTCG |
| pACYCduet-rv | CGCAAGCTTGTCGACCTG |
| pET28a-fw | GAATTCGAGCTCCGTCGACAAG |
| pET28a-rv | GGATCCGCGACCCATTTGCTGT |

**Table S3**. Plasmids used in the study.

| Plasmid | Backbone | Inserted genes | Size (bp) |
|---|---|---|---|
| pDB045 | pETDuet-1 | *ribA*, *cofC*, *fbiC*, *cofD*, ABC transporter, *cofE* | 13,135 |
| pDB060 | pETDuet-1 | *ribA*, *cofC*, *fbiC*, *cofD**, ABC transporter, *cofE* | 13,045 |
| pDB061 | pMAL-c2x | MSMEG_5998 | 6,067 |
| pDB065 | pCDFDuet-1 | *fno* (MCS1) | 4,385 |
| pDB070 | pETDuet-1 | *ribA*, *cofC**, *fbiC*, *cofD**, ABC transporter, and *cofE* | 13,136 |
| pDB071 | pCDFDuet-1 | *fno* (MCS1) and *cofC*, *fbiC, and cofD* (MCS2) | 8,656 |
| pDL008 | pET28a | *fno (Archaeoglobus fulgidus)* | 5,878 |
| pMH01 | pET28a | His6-cofE (*Methanocaldoccoccus jannaschii*) | 6,100 |
| pMH02 | pCDFDuet | c*ofC*, *fbiC*, *cofD* (*P. rhizoxinica*) | 8,085 |
| pFS01 | pET28a | His6-*cofE (P.rhizoxinica*) | |
| pFS03 | pACYCDuet | His6-*cofC (P. rhizoxinica)* | 4,659 |
| pFS04 | pET28a | His6-*cofD (M. jannaschii)* | 6,296 |
| pFS06 | pER28a | His6-*fbiB (Mycolicibacterium smegmatis)* | 6,734 |

*genes from *Methanocaldococcus jannaschii.*

### 2 Results

#### 2.1 Fluorescence Microscopy

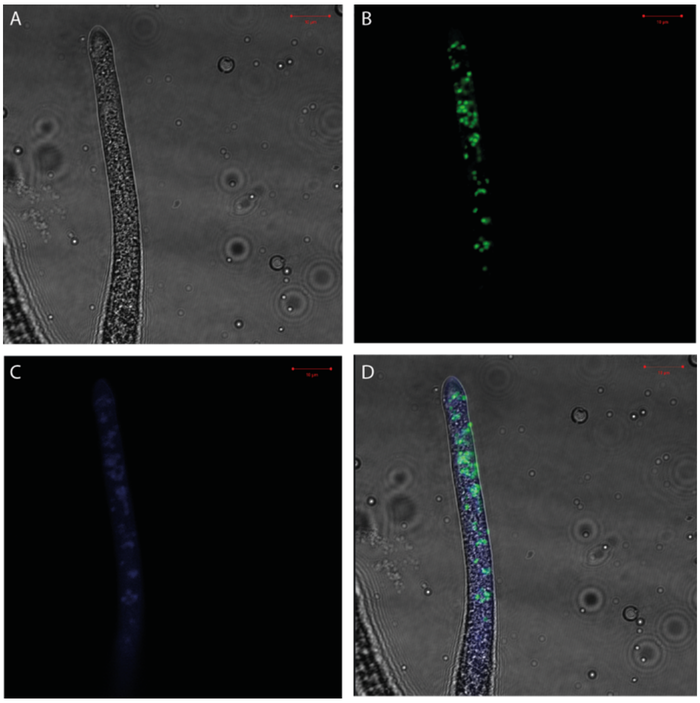

**Figure S1**. Fluorescence microscopy images of a hyphal tip of *Rhizopus microsporus* ATTC 62417 harbouring *Paraburkholderia rhizoxinica* endosymbionts (green). A) Brightfield B) Syto 9 channel in green C) deazaflavin-channel in blue (excitation: 405 nm, emission: approx. 470 nm; D) Overlay. Symbionts and fluorescence of deazaflavins are present within hyphae. Scale bars represent 10 µm.

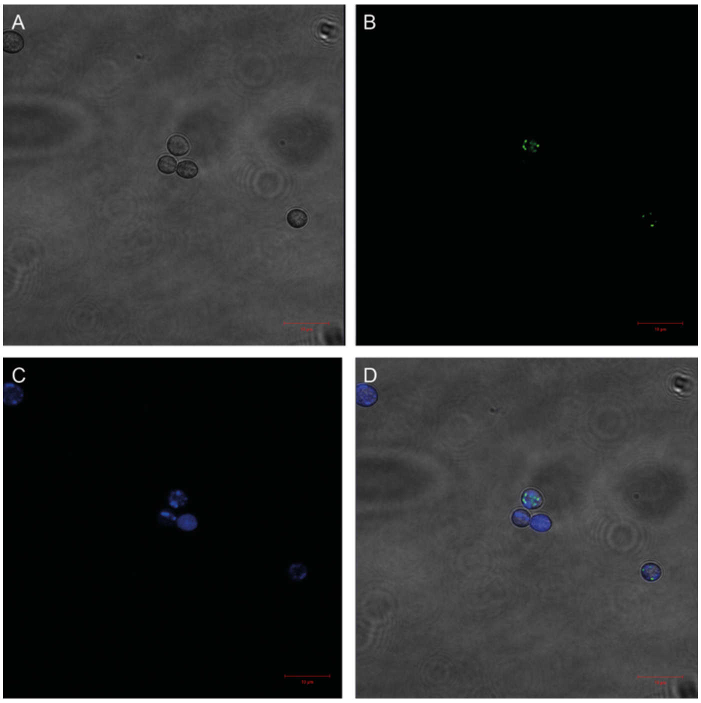

**Figure S2**. Fluorescence microscopy images of spores of *Rhizopus microsporus* ATTC 62417 harboring *Paraburkholderia rhizoxinica* endosymbionts (stained in green). A) Brightfield B) Syto 9 channel in green C) deazaflavin-channel in blue (excitation: 405 nm, emission: approx. 470 nm; D) Overlay. Symbionts and fluorescence of deazaflavins are present within spores. Scale bars represent 10 µm.

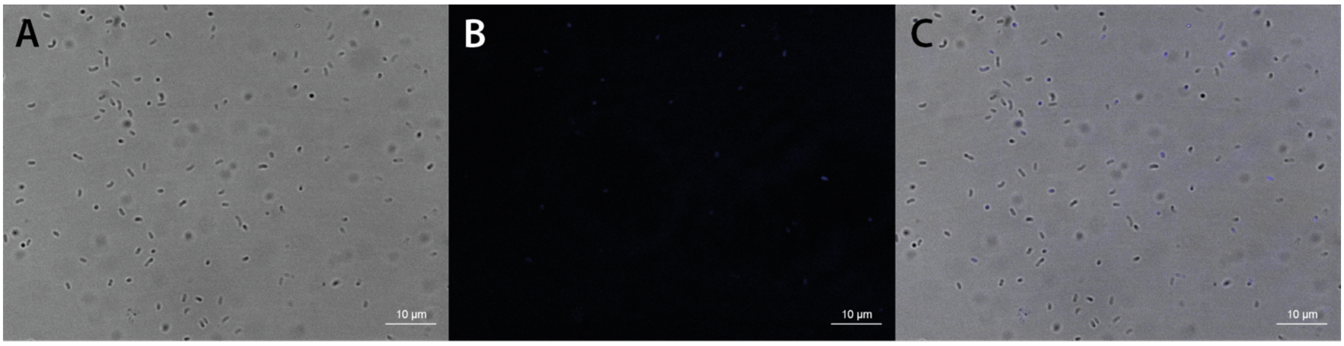

**Figure S3**. Fluorescence microscopy images of *Paraburkholderia rhizoxinica* in axenic culture. A) Brightfield B) Deazaflavin-channel in blue (excitation: 405 nm, emission: approx. 470 nm; C) Overlay. Bacteria display detectable, but weak, deazaflavin-related fluorescence. Scale bars represent 10 µm.

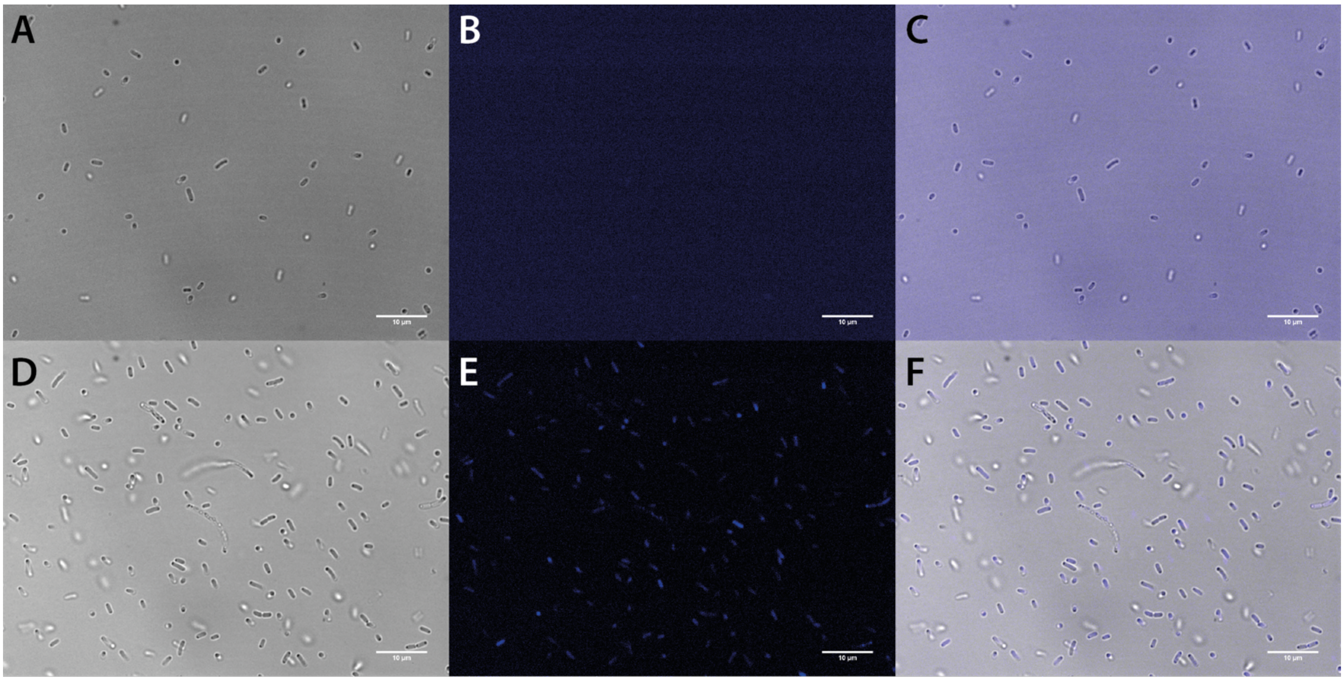

**Figure S4**. Fluorescence microscopy images of *E. coli*. A – C) Empty vector control (*E. coli* / pETDuet). D – F) 3PG-$F_{420}$-producing *E. coli* / pDB045. Images shown correlate to brightfield (A, D), blue deazaflavin-channel (excitation: 405 nm, emission: approx. 470 nm; B, E), and brightfield-deazaflavin overlay (C, F). Engineered bacteria *E. coli* / pDB045 display strong deazaflavin-related fluorescence, fluorescence of vector controls *E. coli* / pETDuet is close to noise level.

#### 2.2 Detection of 3PG-$F_{420}$ by LC-MS/MS

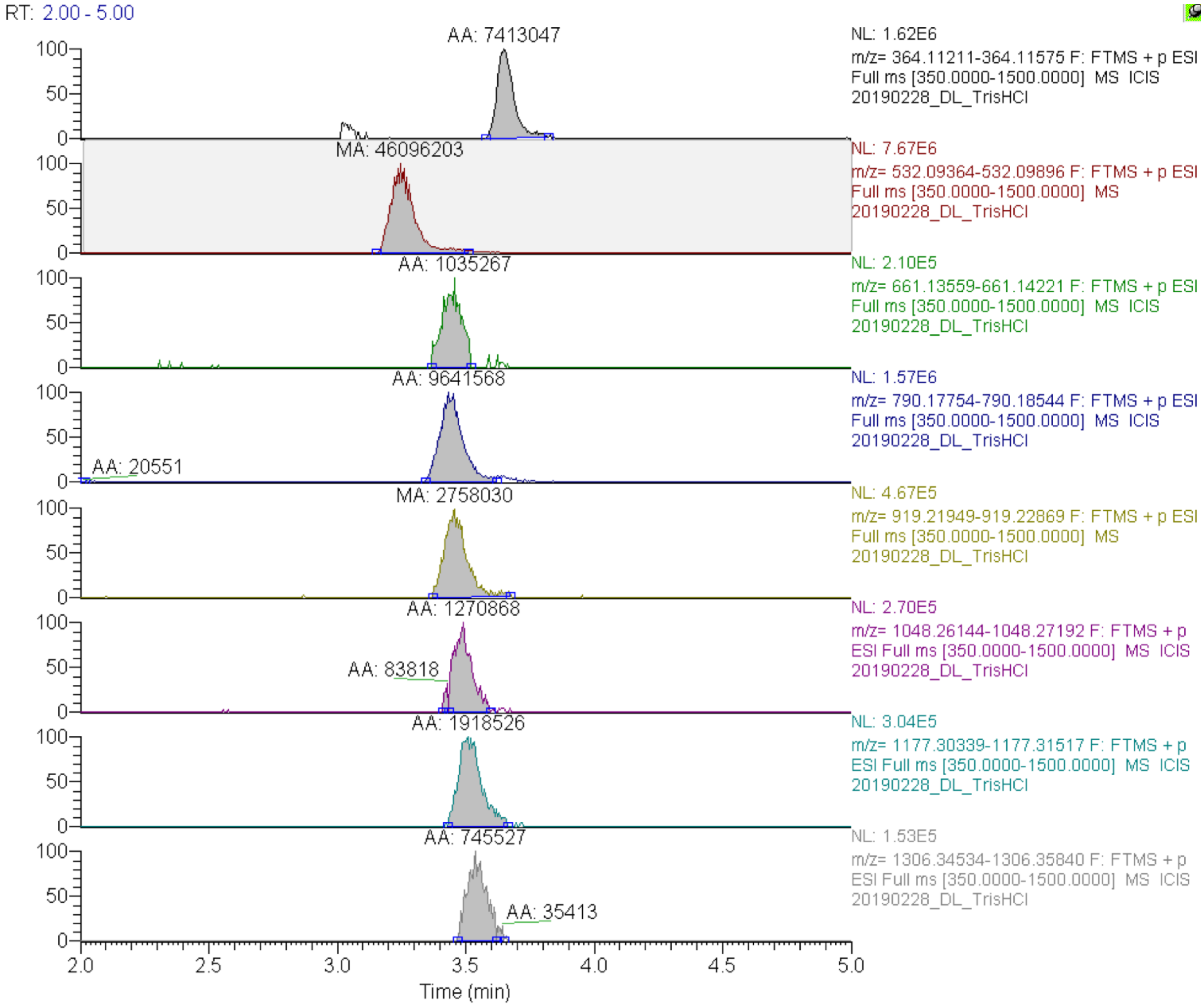

**Figure S5**. LC-MS analysis of extracts of 3PG-$F_{420}$-producing *E. coli* BL21(DE3) /pDB045 showing XICs of 3PG-$F_{420}$-n species with a varying number of (oligo)-γ-glutamate residues. Expected masses ($[M+H]^+$, 10 ppm mass tolerance): $F_O$: 364.11393, 3PG-$F_{420}$-0: 532.09630, 3PG-$F_{420}$-1: 661.13890, 3PG-$F_{420}$-2: 790.18149, 3PG-$F_{420}$-3: 919.22409, 3PG-$F_{420}$-4: 1048.26668, 3PG-$F_{420}$-5: 1177.30928, 3PG-$F_{420}$-6: 1306.35187. Areas under the curve are indicated on top of each peak.

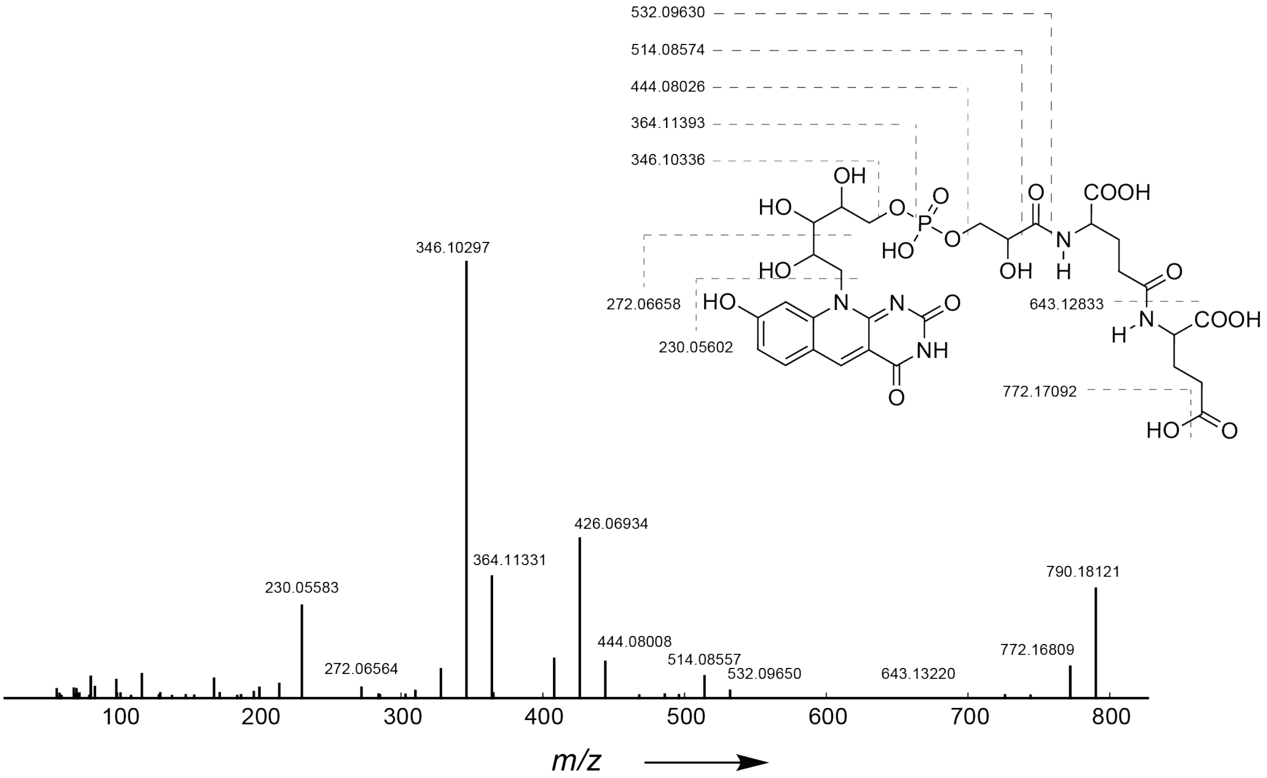

**Figure S6** Tandem-mass spectrum ($MS^2$) of 3PG-$F_{420}$-2 (precursor mass: $[M+H]^+$ m/z 790.18212, calculated: 790.18149) measured on a Thermo Q Exactive mass spectrometer (NCE 30, resolution: 17,500). Measured masses of characteristic peaks are indicated in the spectrum. The proposed fragmentation pattern with calculated fragment masses is shown (top right).

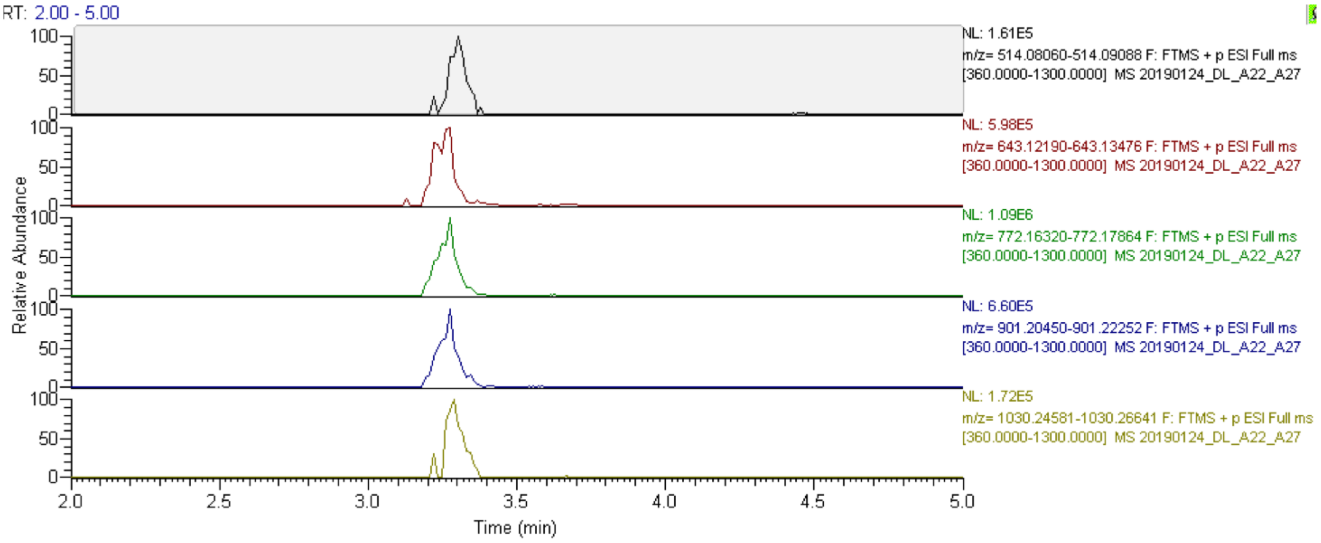

**Figure S7**. LC-MS analysis of extracts (large-scale cultivation) of *E. coli* BL21(DE3) /pDB45 showing XICs of dehydro-$F_{420}$-n ($DF_{420}$-n) species with a varying number of (oligo)-γ-glutamate residues. Expected masses ($[M+H]^+$, 5 ppm mass tolerance): $DF_{420}$-0: 514.08574, $DF_{420}$-1: 643.12833, $DF_{420}$-2: 772.17092, $DF_{420}$-3: 901.21351, $DF_{420}$-4: 1030.25611.

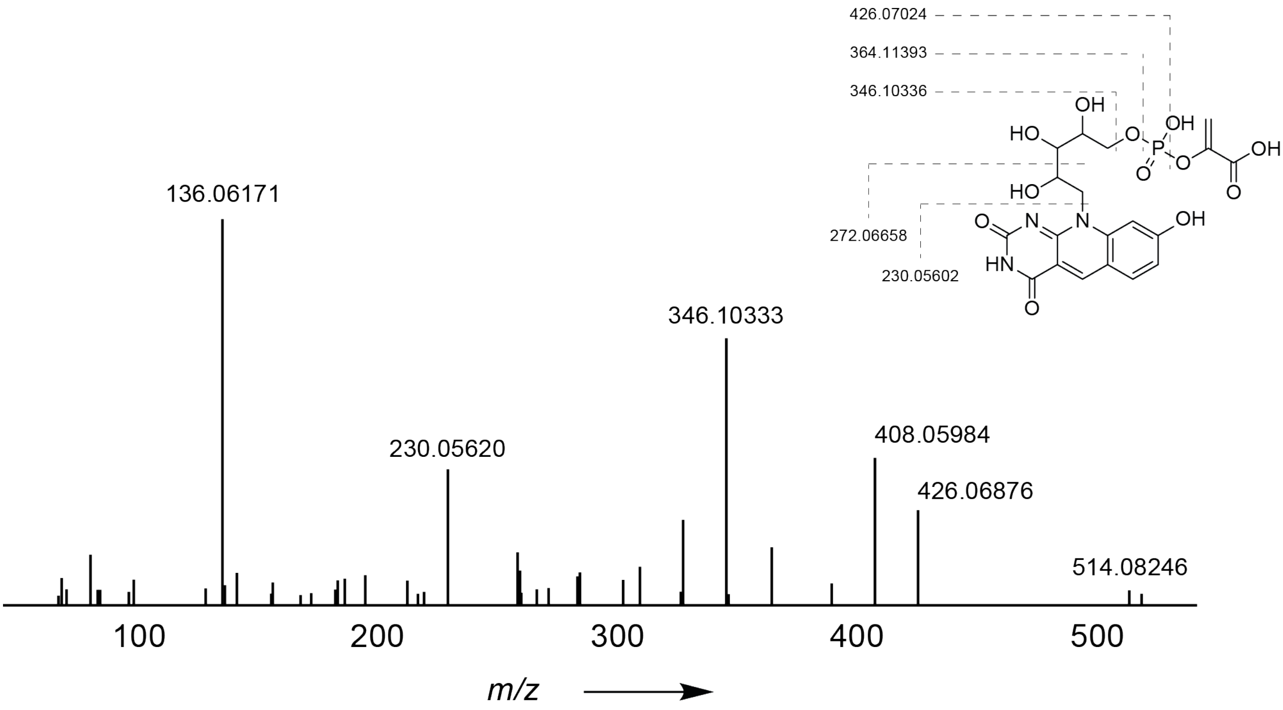

**Figure S8** Tandem-mass spectrum ($MS^2$) of dehydro-$F_{420}$-0 ($DF_{420}$-0, precursor mass: $[M+H]^+$:m/z 514.08246, calculated: 514.08574) measured on a Thermo Q Exactive mass spectrometer (NCE 30, resolution: 17,500). Measured masses of characteristic peaks are indicated in the spectrum. The proposed fragmentation pattern with calculated fragment masses is shown (top right). The peak at m/z 408.06 can be explained as fragment ion m/z 426.07 minus $H_2O$ (-18.01).

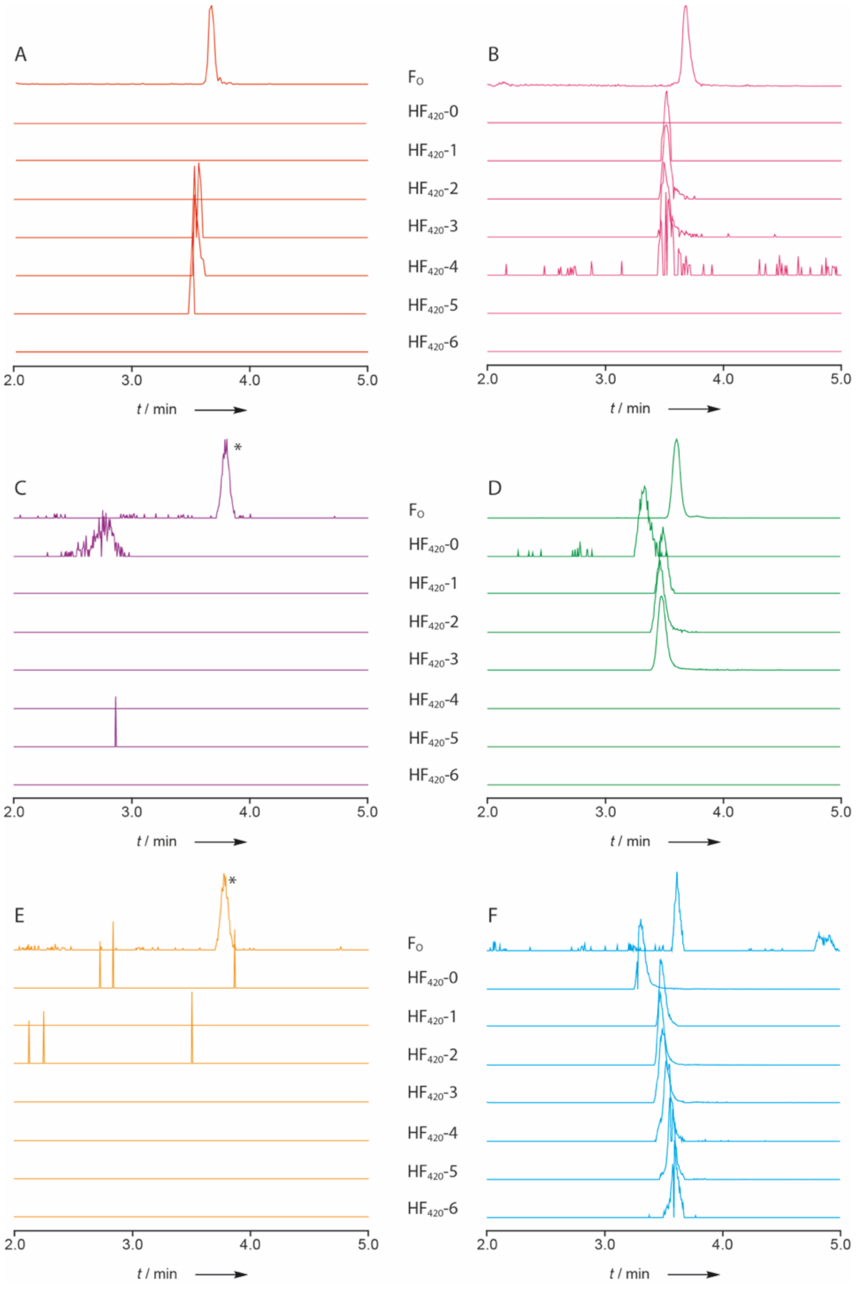

**Figure S9**. Overview of presence and absence of 3PG-$F_{420}$-n species with a varying number of (oligo)-γ-glutamate residues as assessed by LC-MS of microbial extracts (extracted ion chromatograms). Intensities are not drawn to scale, instead, minor peaks are zoomed for better visibility. A) *P. rhizoxinica* axenic culture (red), B) Symbiotic *P. rhizoxinica* isolated from *R. microsporus* ATCC 62417 (magenta), C) cured *R. microsporus* ATCC 62417 without symbionts (purple). D) *R. microsporus* ATCC 62417 with intracellular symbionts (green), E) Naturally symbiont-free *R. microsporus* CBS 344.29 control (yellow), G) 3PG-$F_{420}$ producing *E. coli* BL21(DE3)/pDB045 (blue). Extracted ion chromatograms (10 ppm mass tolerance) were extracted using the following exact masses ($[M+H]^+$): $F_O$: 364.11393, 3PG-$F_{420}$-0: 532.09630, 3PG-$F_{420}$-1: 661.13890, 3PG-$F_{420}$-2: 790.18149, 3PG-$F_{420}$-3: 919.22409, 3PG-$F_{420}$-4: 1048.26668, 3PG-$F_{420}$-5: 1177.30928, 3PG-$F_{420}$-6: 1306.35187. *The asterisks mark a fungal background peak with an m/z similar to the calculated m/z of $F_O$, but consistently shifted retention time and lower intensity (see Figure S12).

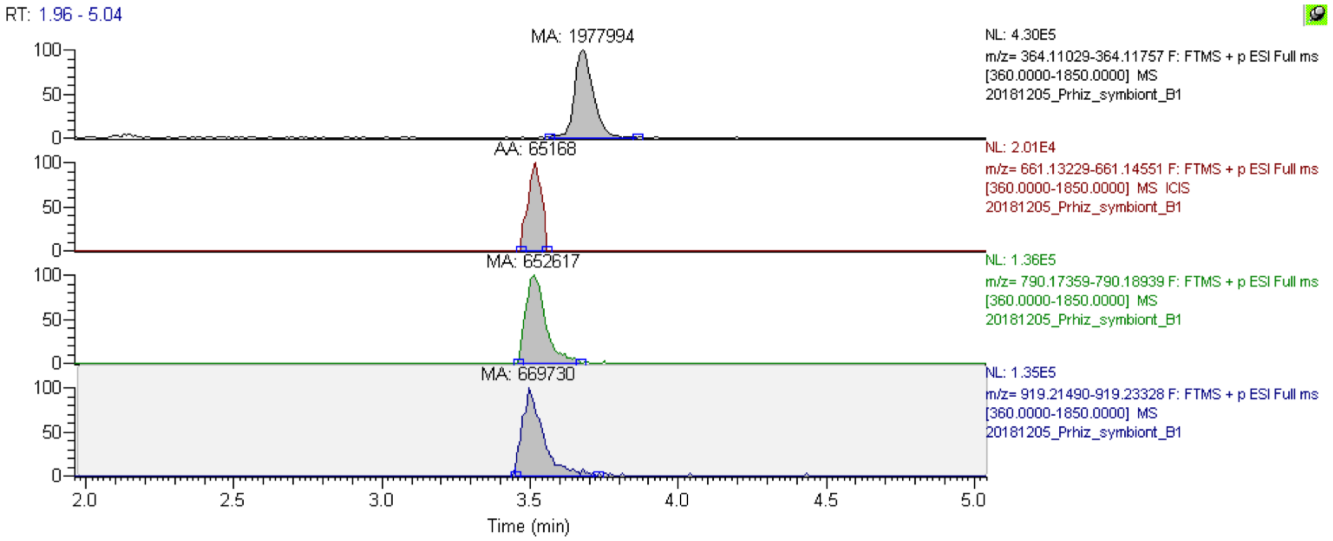

**Figure S10**. LC-MS analysis of extracts of *P. rhizoxinica* pellets isolated from symbiotic growth conditions showing extracted ion chromatograms (XIC) of 3PG-$F_{420}$-n species with a varying number of (oligo)-γ-glutamate residues. Expected masses ($[M+H]^+$, 10 ppm mass tolerance): $F_O$: 364.11393, 3PG-$F_{420}$-1: 661.13890, 3PG-$F_{420}$-2: 790.18149, 3PG-$F_{420}$-3: 919.22409. XICs of missing 3PG-$F_{420}$-n species are not shown. Areas under the curve are indicated on top of each peak.

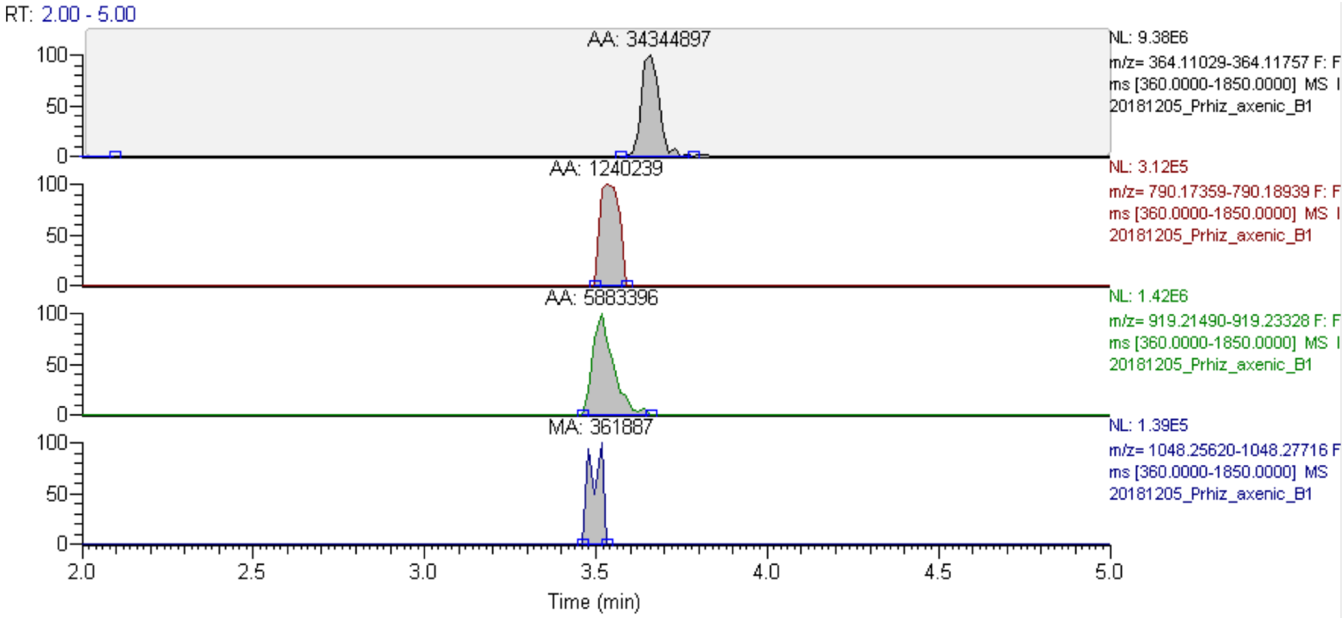

**Figure S11**. LC-MS analysis of extracts of *P. rhizoxinica* grown in axenic culture showing XICs of 3PG-$F_{420}$-n species with a varying number of (oligo)-γ-glutamate residues Expected masses ($[M+H]^+$, 10 ppm mass tolerance): $F_O$: 364.11393, 3PG-$F_{420}$-2: 790.18149, 3PG-$F_{420}$-3: 919.22409, 3PG-$F_{420}$-4: 1048.26668. XICs of missing 3PG-$F_{420}$-n species are not shown. Areas under the curve are indicated on top of each peak.

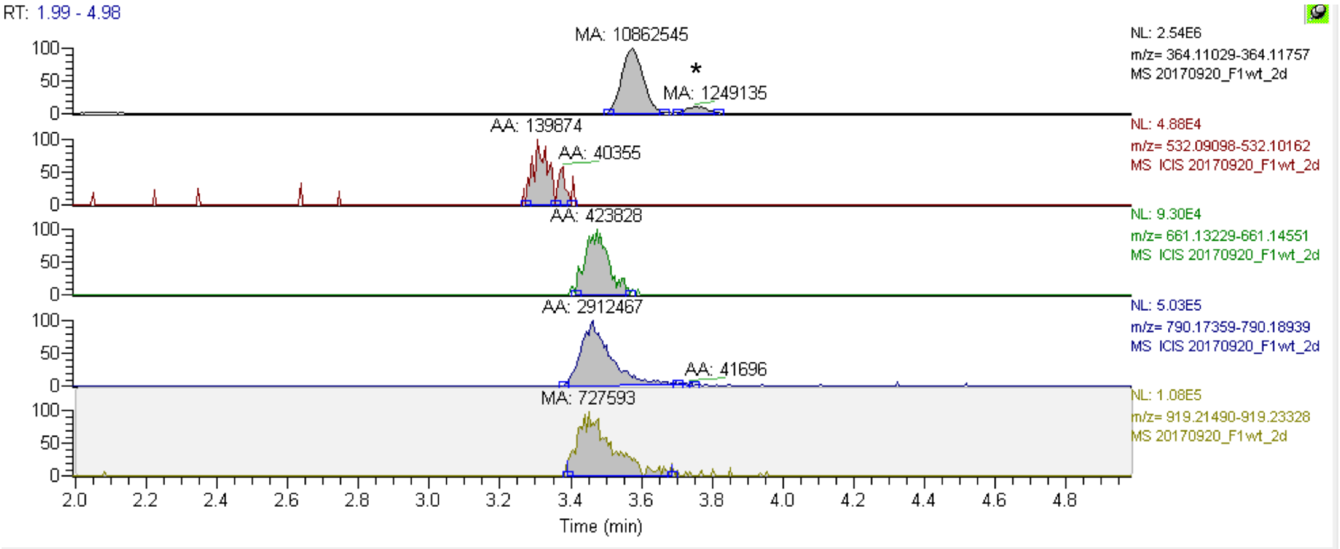

**Figure S12.** LC-MS analysis of extracts of *R. microsporus* ATCC 62417 with intracellular symbionts (*P. rhizoxinica*) showing XICs of 3PG-$F_{420}$-n species with a varying number of (oligo)-γ-glutamate residues. Expected masses ($[M+H]^+$, 10 ppm mass tolerance): $F_O$: 364.11393, 3PG-$F_{420}$-0: 532.09630, 3PG-$F_{420}$-1: 661.13890, 3PG-$F_{420}$-2: 790.18149, 3PG-$F_{420}$-3: 919.22409. Areas under the curve are indicated on top of each peak. XICs of missing 3PG-$F_{420}$-n species are not shown. *A fungal background peak with an m/z close to 364.11393 ($F_O$) elutes shortly after $F_O$ at 3.8 min. This isobaric compound is not related to deazaflavins.

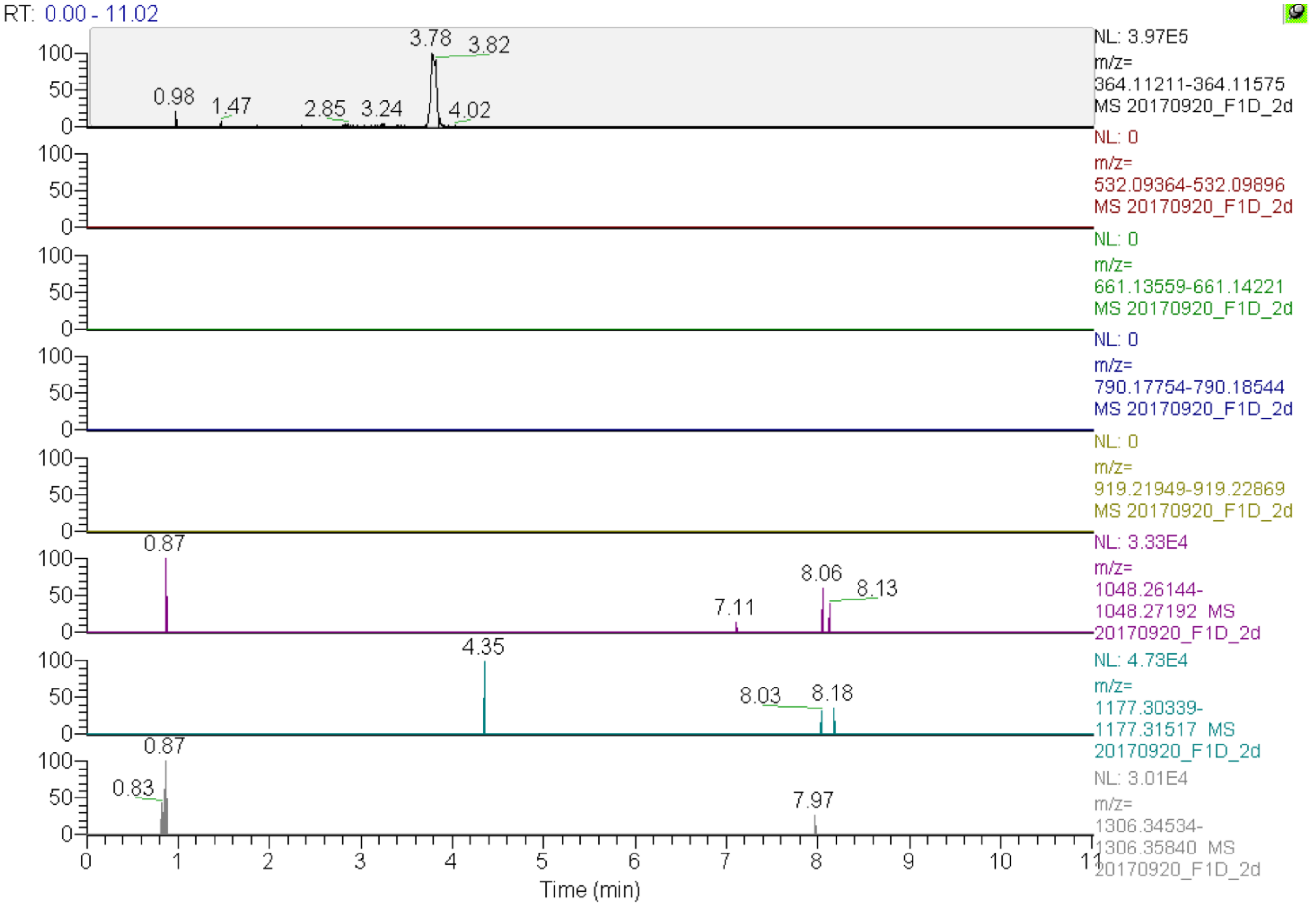

**Figure S13**. LC-MS analysis of extracts of cured *R. microsporus* ATCC 62417 without symbionts (*P. rhizoxinica*) showing XICs of 3PG-$F_{420}$-n species with a varying number of (oligo)-γ-glutamate residues. Expected masses ($[M+H]^+$, 10 ppm mass tolerance): $F_O$: 364.11393, 3PG-$F_{420}$-0: 532.09630, 3PG-$F_{420}$-1: 661.13890, 3PG-$F_{420}$-2: 790.18149, 3PG-$F_{420}$-3: 919.22409, 3PG-$F_{420}$-4: 1048.26668, 3PG-$F_{420}$-5: 1177.30928, 3PG-$F_{420}$-6: 1306.35187. The peak eluting at 3.8 is not $F_O$, but corresponds to the isobaric background compound mentioned above (Figure S12).

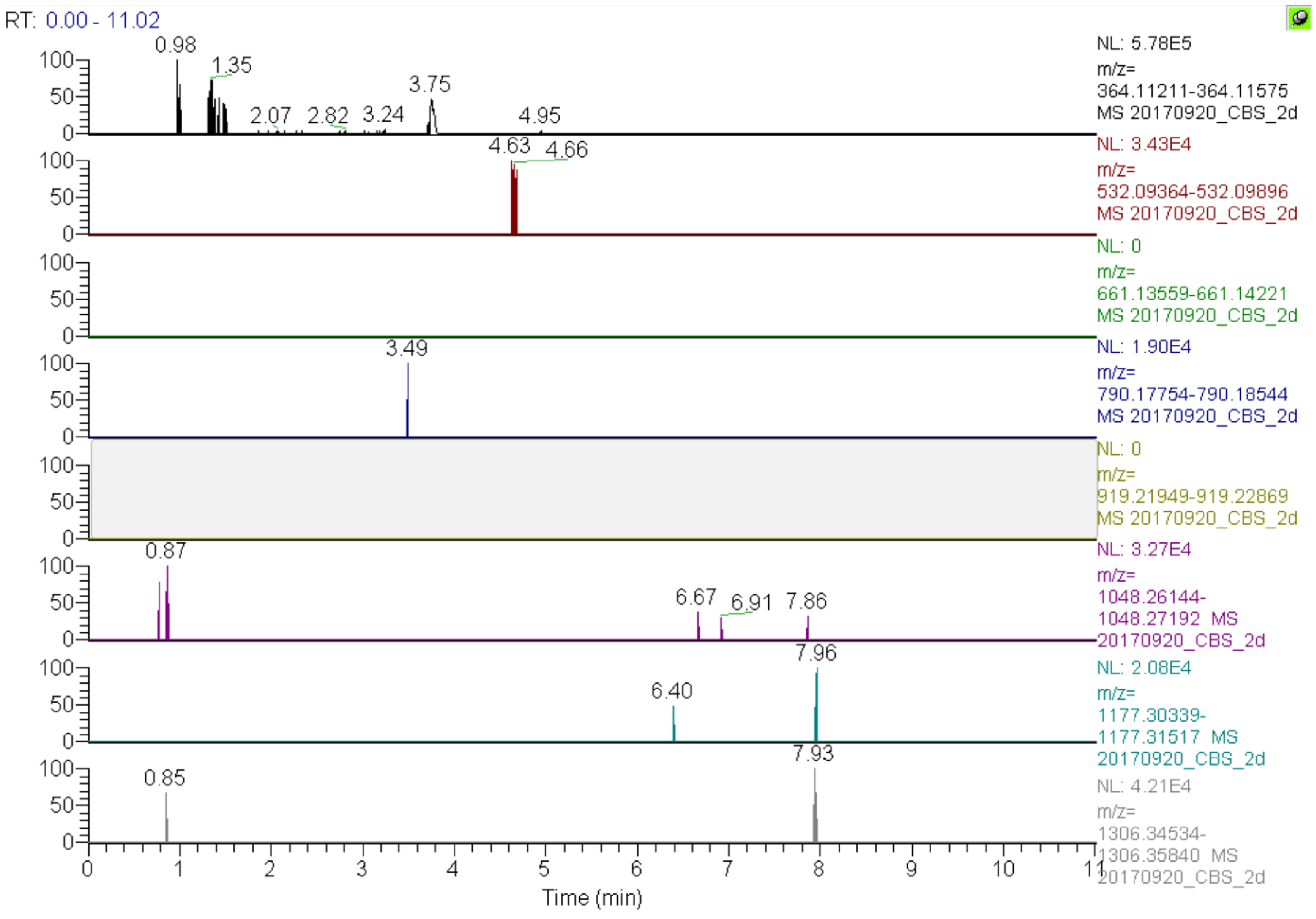

**Figure S14**. LC-MS analysis of extracts of naturally symbiont-free *R. microsporus* CBS 344.29 showing XICs of 3PG-$F_{420}$-n species with a varying number of (oligo)-γ-glutamate residues. Expected masses ($[M+H]^+$, 10 ppm mass tolerance): $F_O$: 364.11393, 3PG-$F_{420}$-0: 532.09630, 3PG-$F_{420}$-1: 661.13890, 3PG-$F_{420}$-2: 790.18149, 3PG-$F_{420}$-3: 919.22409, 3PG-$F_{420}$-4: 1048.26668, 3PG-$F_{420}$-5: 1177.30928, 3PG-$F_{420}$-6: 1306.35187. The peak eluting at 3.8 is not $F_O$, but corresponds to the isobaric background compound mentioned above (Figure S12).

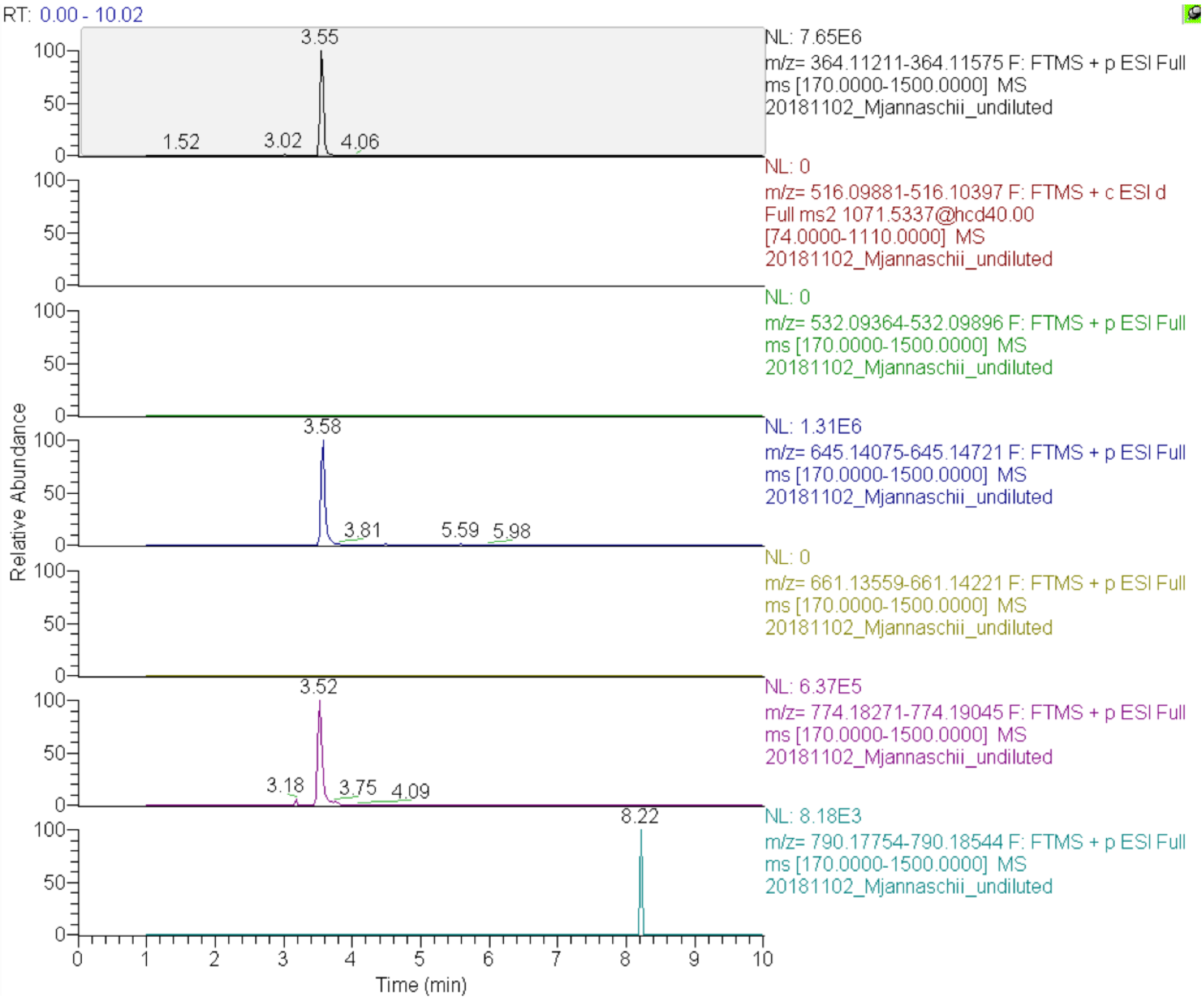

**Figure S15**. LC-MS analysis of extracts of *M. jannaschii* showing XICs of $F_{420}$-n and 3PG-$F_{420}$-n with a varying number of (oligo)-γ-glutamate residues. Expected masses ($[M+H]^+$, 5 ppm mass tolerance): $F_O$: 364.11393, $F_{420}$-0: 516.10139, 3PG-$F_{420}$-0: 532.09630, $F_{420}$-1: 645.14398, 3PG-$F_{420}$-1: 661.13890, $F_{420}$-2: 774.18658, 3PG-$F_{420}$-2: 790.18149. While classical $F_{420}$-n was produced in good yields, 3PG-$F_{420}$-0 and $F_{420}$-0 were not detected.

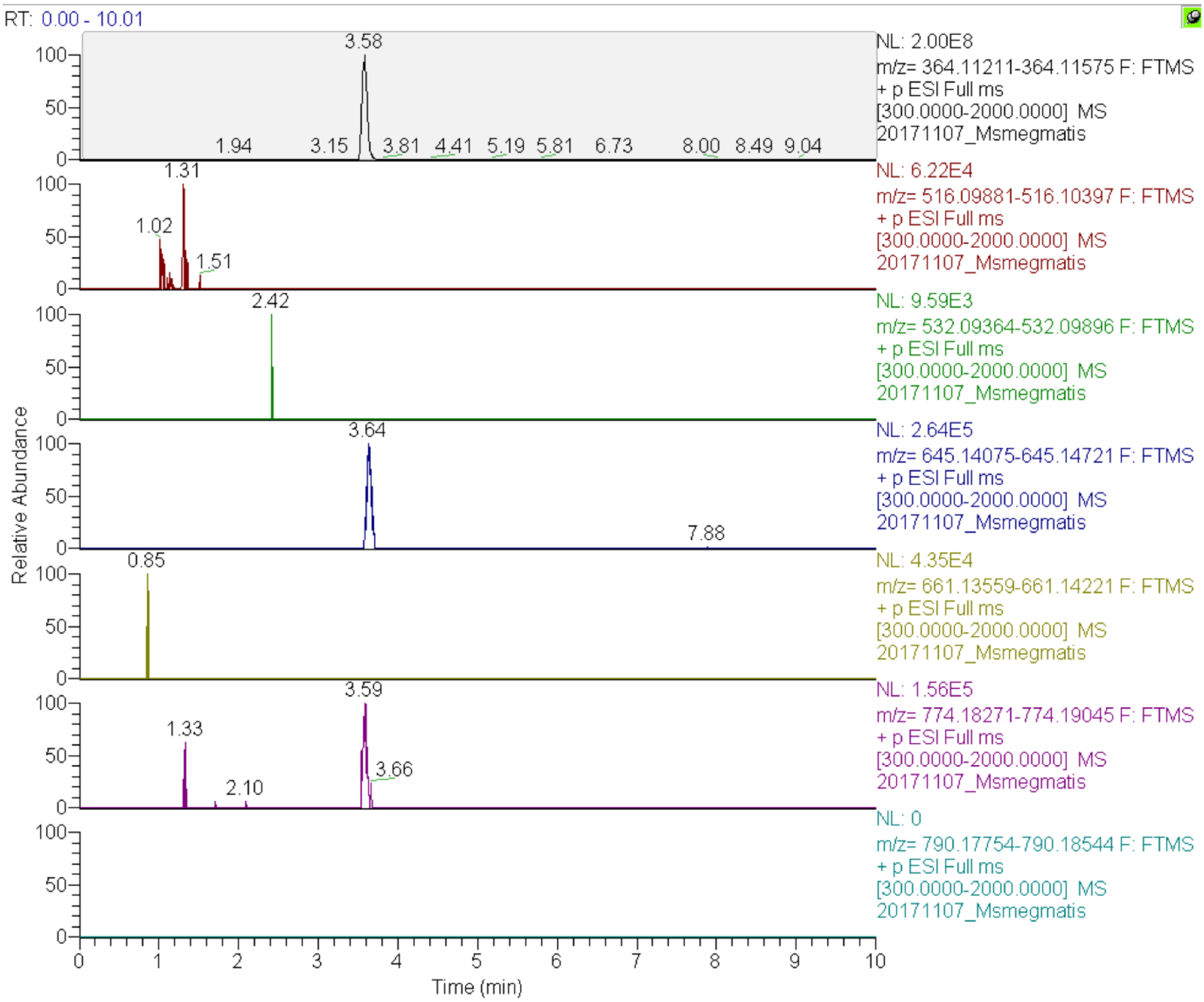

**Figure S16**. LC-MS analysis of extracts of *M. smegmatis* showing XICs of $F_{420}$-n and 3PG-$F_{420}$-n with a varying number of (oligo)-γ-glutamate residues. Expected masses ($[M+H]^+$, 5 ppm mass tolerance): $F_O$: 364.11393, $F_{420}$-0: 516.10139, 3PG-$F_{420}$-0: 532.09630, $F_{420}$-1: 645.14398, 3PG-$F_{420}$-1: 661.13890, $F_{420}$-2: 774.18658, 3PG-$F_{420}$-2: 790.18149. While classical $F_{420}$-n was produced in good yields, 3PG-$F_{420}$ was not detectable. $F_{420}$-0 was not found.

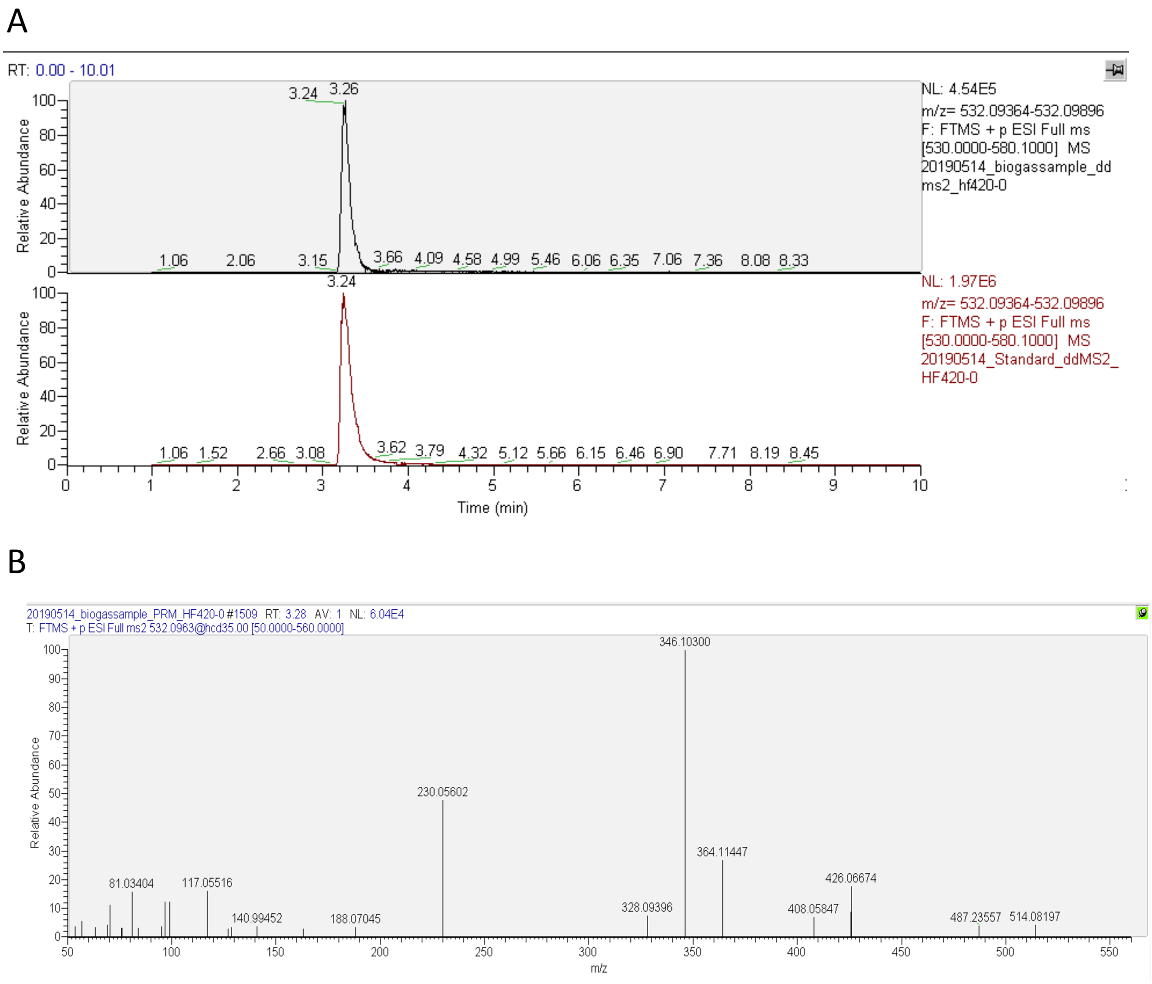

**Figure S17**. LC-MS analysis of extracts of a biogas-producing microbial community after HPLC purification. A) XIC at m/z 532.09630 (corresponds to 3PG-$F_{420}$-0 $[M+H]^+$). Upper chromatogram: sample. Lower chromatogram: 3PG-$F_{420}$-0 standard. B) The MS/MS spectrum of the peak eluting at RT 3.34 min displaying characteristic fragments of $F_{420}$ derivatives (m/z 230.06, 346.10, 364.11, 408.06, 426.07, 514.08).

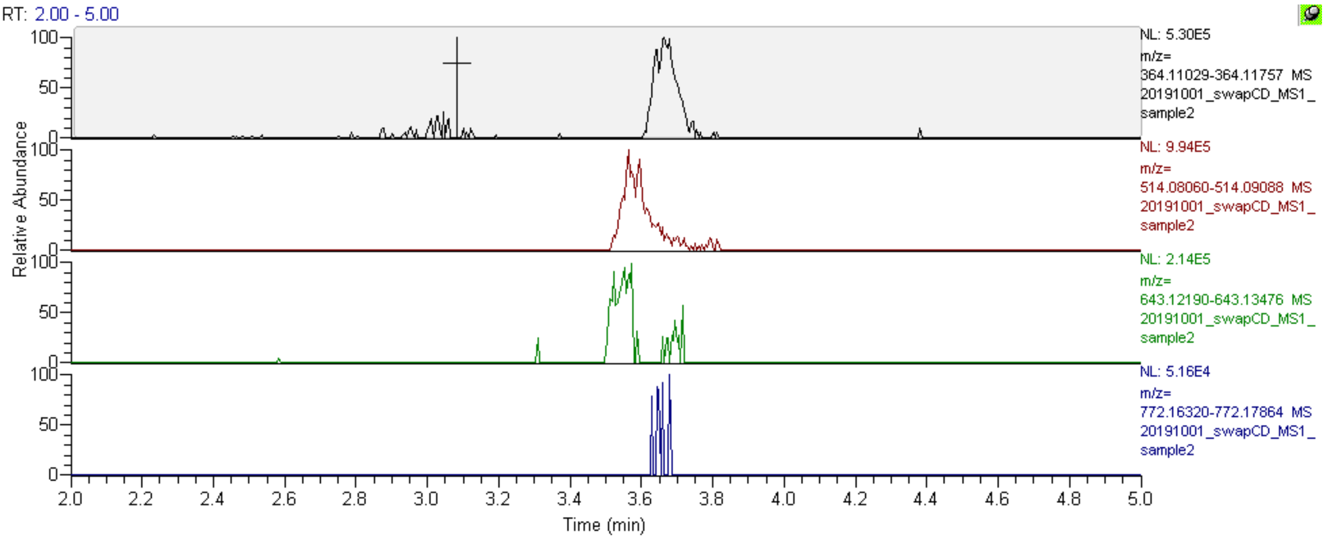

**Figure S18**. LC-MS analysis of extracts of *E. coli* BL21(DE3) /pDB070 showing XICs of dehydro-$F_{420}$-n ($DF_{420}$-n) species with a varying number of (oligo)-γ-glutamate residues. Expected masses ($[M+H]^+$, 10 ppm mass tolerance): $DF_{420}$-0: 514.08574, $DF_{420}$-1: 643.12833, $DF_{420}$-2: 772.17092.

#### 2.3 Structure elucidation of deazaflavins

##### *2.3.1 Structure elucidation of classical $F_{420}$ (control)*

Classical $F_{420}$-n was obtained as described above as yellow solid directly eluted from FPLC and submitted for 1D and 2D NMR analysis. The composition of mixed $F_{420}$-n was determined as $F_{420}$-4 with the molecular formula of $C_{39}H_{50}O_{24}N_7P$ based on the ESI-HRMS analysis (*m/z* 1032.26904 ($[M+H]^+$ calcd. 1032.27176 Δ = – 2.63 ppm); $F_{420}$-5 with the molecular formula of $C_{44}H_{57}O_{27}N_8P$ based on the ESI-HRMS analysis (*m/z* 1161.31250 ($[M+H]^+$ calcd. 1161.31435 Δ = – 1.59 ppm); $F_{420}$-6 with the molecular formula of $C_{49}H_{64}O_{30}N_9P$ based on the ESI-HRMS analysis (*m/z* 1290.35486 ($[M+H]^+$ calcd. 1290.35694 Δ = – 1.62 ppm); $F_{420}$-7 with the molecular formula of $C_{54}H_{73}O_{33}N_{10}P$ based on the ESI-HRMS analysis (*m/z* 710.20178 ($[2M+H]^{2+}$ calcd. 710.20341 Δ = – 2.29 ppm). Thorough interpretation of the $^1H$ NMR and $^{13}C$ NMR spectra ($D_2O$) indicated the typical 5-deazaflavin moiety with the observation of $\delta_{H\text{-}5}$ 8.82 ppm, $\delta_{C\text{-}5}$ 144.8 ppm; $\delta_{H\text{-}6}$ 7.91 ppm, $\delta_{C\text{-}6}$ 135.5 ppm; $\delta_{H\text{-}7}$ 7.16 ppm, $\delta_{C\text{-}7}$ 118.1 ppm; $\delta_{H\text{-}9}$ 7.31 ppm, $\delta_{C\text{-}9}$ 102.3 ppm. The ribityl moiety was deduced from the observation of $\delta_{H2\text{-}1'}$ 4.67/4.97 ppm, $\delta_{C\text{-}1'}$ 49.31 ppm; $\delta_{H\text{-}2'}$ 4.37 ppm, $\delta_{C\text{-}2'}$ 70.2 ppm; $\delta_{H\text{-}3'}$ 3.99 ppm, $\delta_{C\text{-}3'}$ 73.3 ppm; $\delta_{H\text{-}4'}$ 4.07 ppm, $\delta_{C\text{-}4'}$ 71.8 ppm; $\delta_{H2\text{-}5'}$ 4.07/4.17 ppm, $\delta_{C\text{-}5'}$ 67.4 ppm. The lactyl moiety was assigned based on the observation of resonance at $\delta_{H\text{-}6'}$ 4.70 ppm, $\delta_{C\text{-}6'}$ 72.6 ppm; $\delta_{H3\text{-}7'}$ 1.49 ppm, $\delta_{C\text{-}7'}$ 19.9 ppm; $\delta_{C\text{-}8'}$ 175.4 ppm. Due to the overlapped signals from the glutamyl chain, the assignment of one unit can be deduced from the observation of $\delta_{H\text{-}10'}$ 4.35 ppm, $\delta_{C\text{-}10'}$ 52.5 ppm; $\delta_{H2\text{-}11'}$ 1.94/2.21 ppm, $\delta_{C\text{-}11'}$ 27.0 ppm; $\delta_{H2\text{-}12'}$ 2.41 ppm, $\delta_{C\text{-}12'}$ 32.0 ppm; $\delta_{C\text{-}13'}$ 175.6 ppm; $\delta_{C\text{-}14'}$ 175.7 ppm. The doublets of C-4', C-5', C-6', C-7' and C-8' indicated the $^{13}C$–$^{31}P$ coupling ($^3J_{C\text{-}4'\text{—}P}$ = 7.56 Hz, $^2J_{C\text{-}5'\text{—}P}$ = 5.06 Hz, $^2J_{C\text{-}6'\text{—}P}$ = 4.94 Hz, $^3J_{C\text{-}7'\text{—}P}$ = 2.93 Hz, $^3J_{C\text{-}8'\text{—}P}$ = 6.60 Hz) between the carbon atoms of the ribityl and lactyl moieties assigned to these resonances and the phosphorus of the phosphate group. Finally, the classical $F_{420}$-n was confirmed based on the match with literature reported NMR data recorded in $D_2O$.(8) Shift in $^{13}C$ resonances are presumably a result of the presence of ammonium salts and/or pH differences as indicated in the literature. Detailed assignment is present in Table S4-6.

**Table S4.** NMR Data (DMSO-$d_6$, at 300 K) for $F_{420}$-n .[a]

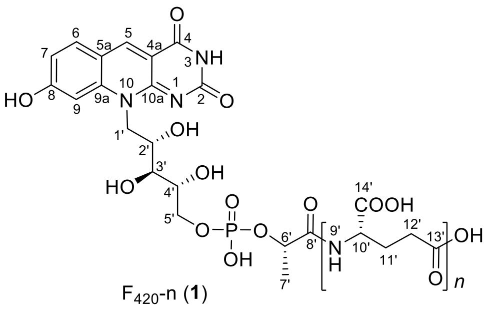

$F_{420}$-n (**1**)

| | $F_{420}$-n | | | | |
|---|---|---|---|---|---|
| position | $\delta_C$, mult.[b] (*J* in Hz)[c] | $\delta_H$, mult. (*J* in Hz) | COSY | HMBC | NOESY |
| *N* (1) | | | | | |
| 2 | n.d. | | | | |
| *N*H (3) | | 11.04, br s | | 4a | |
| 4 | 162.37, qC | | | | |
| 4a | 110.65, qC | | | | |
| 5 | 141.66, CH | 8.90, s | | 4, 5a, 6, 9a, 10a | 6 |
| 5a | 115.64, qC | | | | |
| 6 | 133.83, CH | 8.04, d (*8.87*) | 7 | 5, 5a, 8, 9a | 5 |
| 7 | 115.51, CH | 7.04, dd (*8.87,* [illegible] | 6, 9 | 5a, 9 | |
| 8 | 164.53, qC | | | | |
| 9 | 102.16, CH | 7.40, s | 7 | 5a, 8, 9a | 1'a, 1'b |
| 9a | 144.01, qC | | | | |
| 10 | | | | | |
| 10a | 157.80, qC | | | | |
| 1'a | 48.14, $CH_2$ | 4.80, m | | | 9 |
| 1'b | | 4.65, m | | | 9 |
| 2' | 69.28, CH | 4.25, m | 3' | | |
| 3' | 73.67, CH | 3.62, t (*5.37*) | 2', 4' | 1', 2', 4' | |
| 4' | 70.98, $CH_2$ (*6.97*) | 3.89, m | 2', 3' | 3', 5' | |
| 5'a | 68.25, $CH_2$ | 4.13, m | 4', 5'b | 3' | |
| 5'b | | 3.94, m | 5'a | 4' | |
| 6' | 71.72, CH (*4.96*) | 4.67, q (*7.12*) | 8' | 7', 8' | *N*H (9') |
| 7' | 19.62, $CH_3$ (*3.19*) | 1.39, d (*7.12*) | 6' | 6', 8' | *N*H (9') |
| 8' | 170.77, qC | | | | |
| *N*H (9') [d] | | 8.10, d (*8.45*) | 10' | | 6', 7' |
| 10' [d] | 51.50, CH | 4.15, m | 9', 11'b | 11', 12', 13', 14' | |
| 11'a [d] | 26.97, $CH_2$ | 1.96, m | 10', 11'a, 12' | 10', 12', 13', 14' | |
| 11'b [d] | | 1.74, m | 10', 11'a. 12' | 10', 12', 13', 14' | |
| 12' [d] | 31.50, $CH_2$ | 2.20, m | 11'a, 11'b | 10', 11', 13' | |
| 13' [d] | 171.45, qC | | | | |
| 14' [d] | 173.37, qC | | | | |

[a] 600 MHz for $^1$H NMR and 150 MHz for $^{13}$C NMR
[b] numbers of attached protons were determined by analysis of 2D spectra.
[c] coupling constant indicated $^{13}$C–$^{31}$P coupling
[d] NMR resonance are overlapped by glutamate chains and only one unit was presented.

**Table S5.** NMR Data ($D_2O$, at 300 K) for $F_{420}$-n .[a]

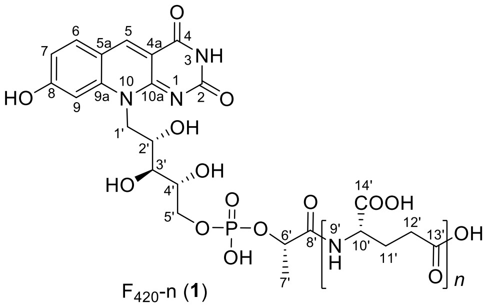

| | $F_{420}$-n | | | | |
|---|---|---|---|---|---|
| position | $\delta_C$, mult.[b] (*J* in Hz)[c] | $\delta_H$, mult. (*J* in Hz) | COSY | HMBC | NOESY |
| *N*H (1) | | | | | |
| 2 | 156.58, qC | | | | |
| *N*H (3) | | | | | |
| 4 | 163.56, qC | | | | |
| 4a | 109.67, qC | | | | |
| 5 | 144.76, CH | 8.82, s | | 4, 5a, 6, 9a, 10a | 6 |
| 5a | 117.91, qC | | | | |
| 6 | 135.44, CH | 7.92, d (*8.80*) | 7 | 5a, 8, 9a | 5, 7 |
| 7 | 118.69, CH | 7.16, dd (*8.80, 1.70*) | 6 | 5a, 9 | 5 |
| 8 | 166.33, qC | | | | |
| 9 | 102.33, CH | 7.31, s | | 5a, 8, 9a | |
| 9a | 143.96, qC | | | | |
| 10 | | | | | |
| 10a | 155.79, qC | | | | |
| 1'a | 49.30, $CH_2$ | 4.96, m | 2' | | 4' |
| 1'b | | 4.69, m | 2' | | |
| 2' | 70.19, CH | 4.37, m | 1'a, 1'b, 3' | | 4' |
| 3' | 73.24, CH | 3.99, t (*3.65*) | 2', 4' | 1', 2', 4', 5' | |
| 4' | 71.76, $CH_2$, d, (*7.56*) | 4.07, m | 3' | | 1', 2' |
| 5'a | 67.42, $CH_2$, d, (*5.06*) | 4.17, m | 5'b | 3' | |
| 5'b | | 4.07, m | 5'a | 3' | |
| 6' | 72.53, CH d, (*4.94*) | 4.70, q (*7.12*) | 7' | 7', 8' | |
| 7' | 19.93, $CH_3$, d, (*2.93*) | 1.49, d (*6.89*) | 6' | 6', 8' | |
| 8' | 175.29, qC, d, (*6.60*) | | | | |
| *N*H (9') [d] | | | | | |
| 10' [d] | 52.31, CH | 4.35, m | 11'a, 11'b | 11', 12', 14' | |
| 11'a [d] | 26.98, $CH_2$ | 2.21, m | 10', 12' | 10', 12', 13', 14' | |
| 11'b [d] | | 1.96, m | 10', 12' | 10', 12', 13', 14' | |
| 12' [d] | 31.98, $CH_2$ | 2.41, m | 11'a, 11'b | 10', 11', 13' | |
| 13' [d] | 175.59, qC | | | | |
| 14' [d] | 175.54, qC | | | | |

[a] 600 MHz for $^1$H NMR and 150 MHz for $^{13}$C NMR
[b] numbers of attached protons were determined by analysis of 2D spectra.
[c] coupling constant indicated $^{13}$C–$^{31}$P coupling
[d] NMR resonance are overlapped by glutamate chains and only one unit was presented.

**Table S6.** $^{13}$C NMR comparison of $F_{420}$-n with reported for $F_{420}$-2.(8)

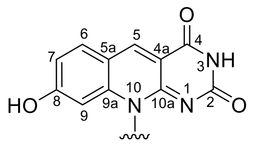

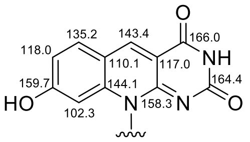

$F_{420}$-2 (Wolfe, 1978)

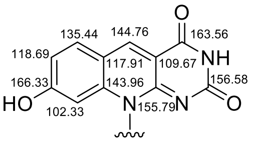

$F_{420}$-n (**1**)

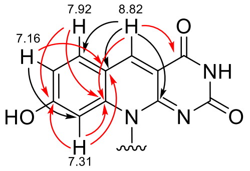

$F_{420}$-n (**1**)

| | $F_{420}$-n in $D_2O$ | $F_{420}$-2 (ammonium salt in $D_2O$) |
|---|---|---|
| position | $\delta_C$, mult.[b] (*J* in Hz)[c] | $\delta_C$, mult. (*J* in Hz) |
| *N* (1) | | |
| 2 | 156.58, qC | 164.4 |
| *N*H (3) | | |
| 4 | 163.56, qC | 166.0 |
| 4a | 109.67, qC | 117.0 |
| 5 | 144.76, CH | 143.4 |
| 5a | 117.91, qC | 110.1 |
| 6 | 135.44, CH | 135.2 |
| 7 | 118.69, CH | 118.0 |
| 8 | 166.33, qC | 159.7 |
| 9 | 102.33, CH | 102.3 |
| 9a | 143.96, qC | 144.1 |
| 10 | | |
| 10a | 155.79, qC | 158.3 |
| 1'a | 49.30, $CH_2$ | 48.3 |
| 1'b | | |
| 2' | 70.19, CH | 70.5 |
| 3' | 73.24, CH | 73.7 |
| 4' | 71.76, $CH_2$, d (*7.56*) | 71.9, d (*7.9*) |
| 5'a | 67.42, $CH_2$, d (*5.06*) | 67.9, d (*5.1*) |
| 5'b | | |
| 6' | 72.53, CH, d (*4.94*) | 73.0, d (*5.2*) |
| 7' | 19.93, $CH_3$, d (*2.93*) | 20.5 |
| 8' | 175.31, qC, d (*6.60*) | 174.9, d (*6.1*) |
| *N*H (9') | | |
| 10' | 52.31, CH | 55.0 |
| 11'a | 26.98, $CH_2$ | 28.5 |
| 11'b | | |
| 12' | 31.98, $CH_2$ | 32.9 |
| 13' | 175.59, qC | 175.8 |
| 14' | 175.54, qC | 178.5 |

[a]chemical shifts in the table and key HMBC correlations are highlighted in red; shift in $^{13}$C resonances are presumably a result of the presence of ammonium salts and/or pH differences as indicated in the literature.

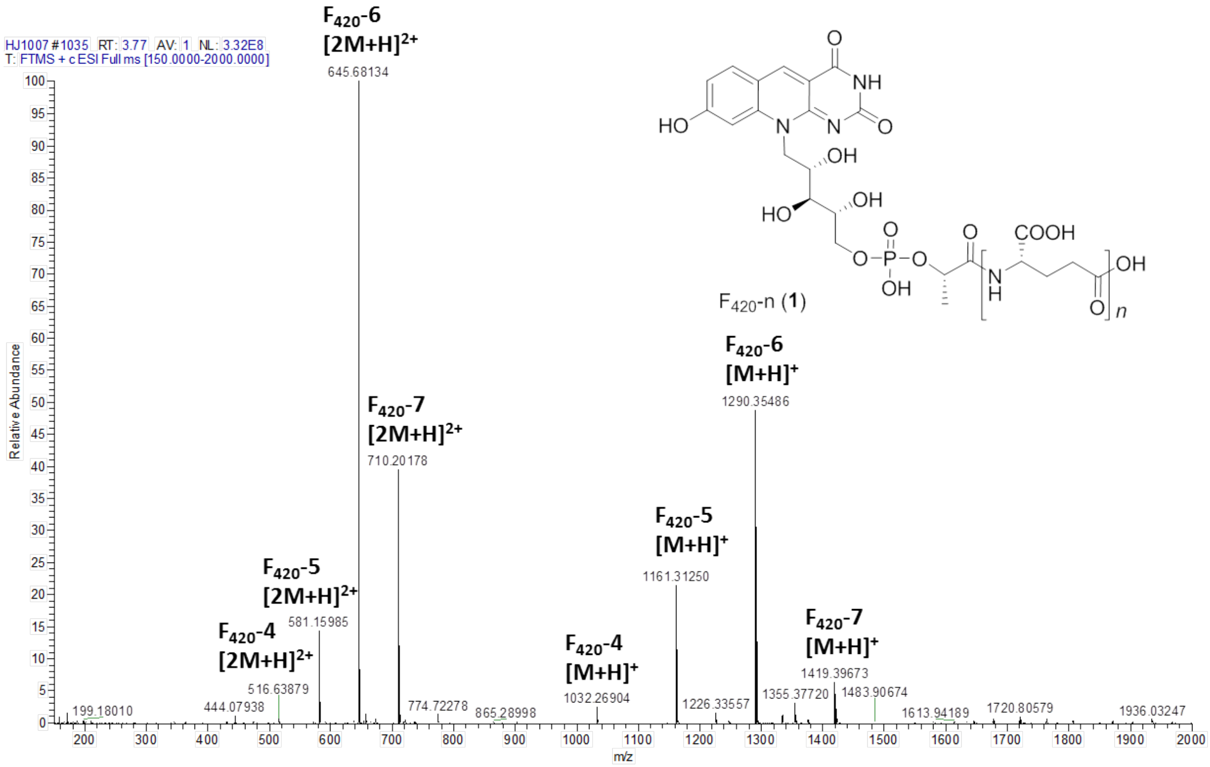

**Figure S19**. ESI-HRMS spectrum of $F_{420}$-n.

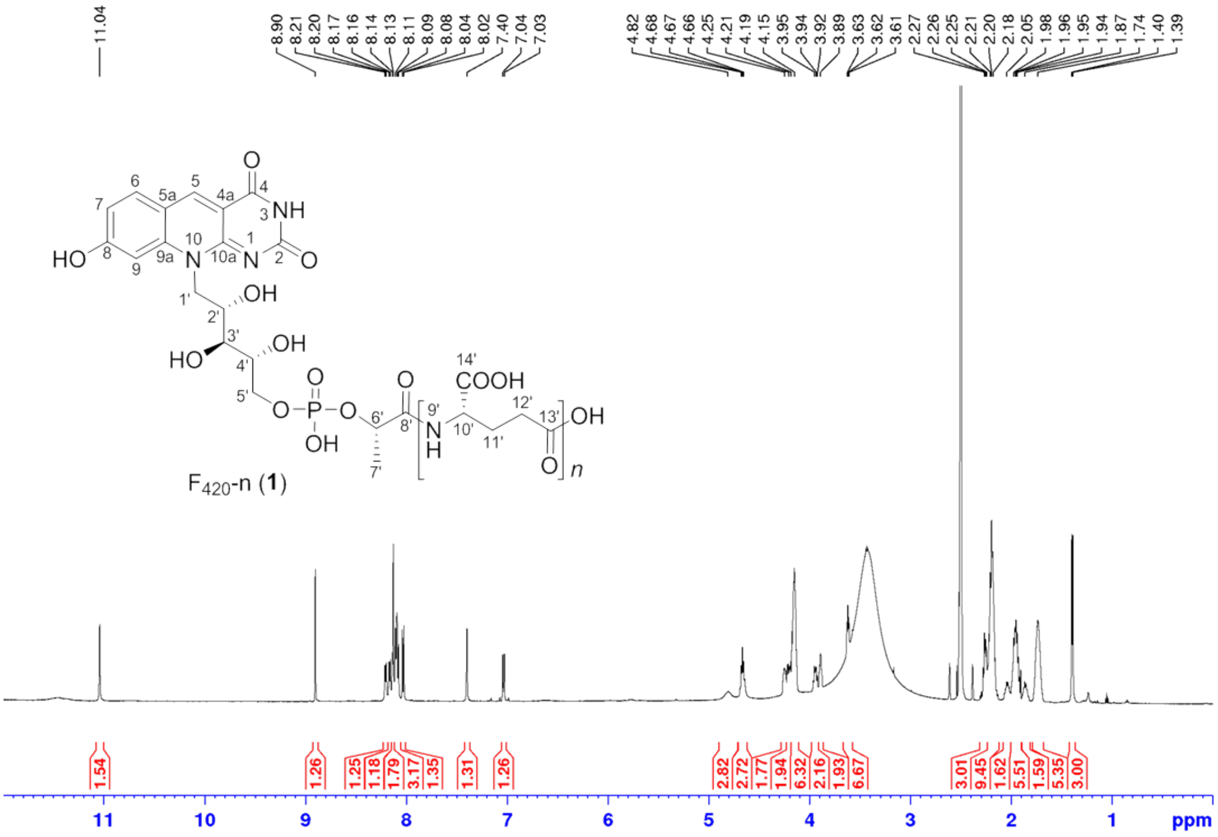

**Figure S20**. $^{1}$H NMR spectrum of $F_{420}$-n (DMSO-$d_6$, 600 MHz, 300 K).

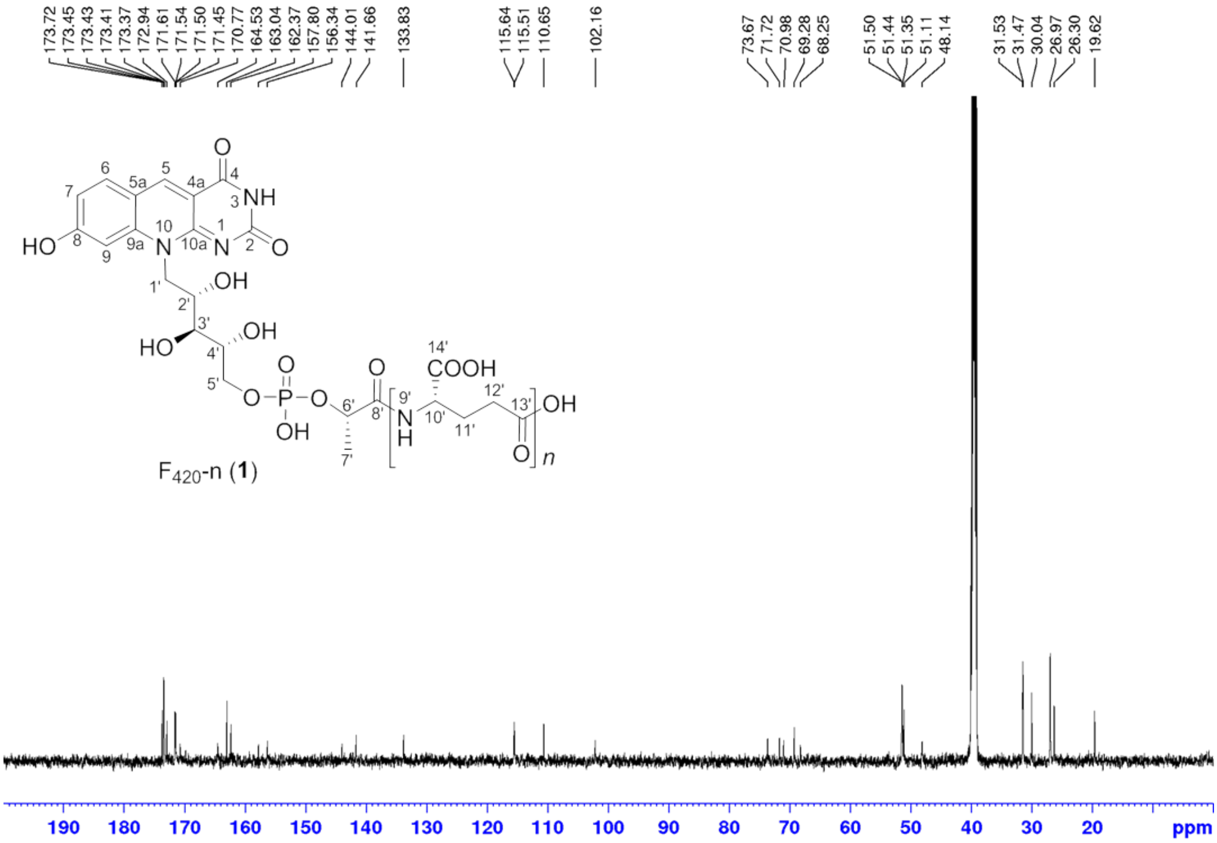

**Figure S21**. $^{13}$C NMR spectrum of $F_{420}$-n (DMSO-$d_6$, 150 MHz, 300 K).

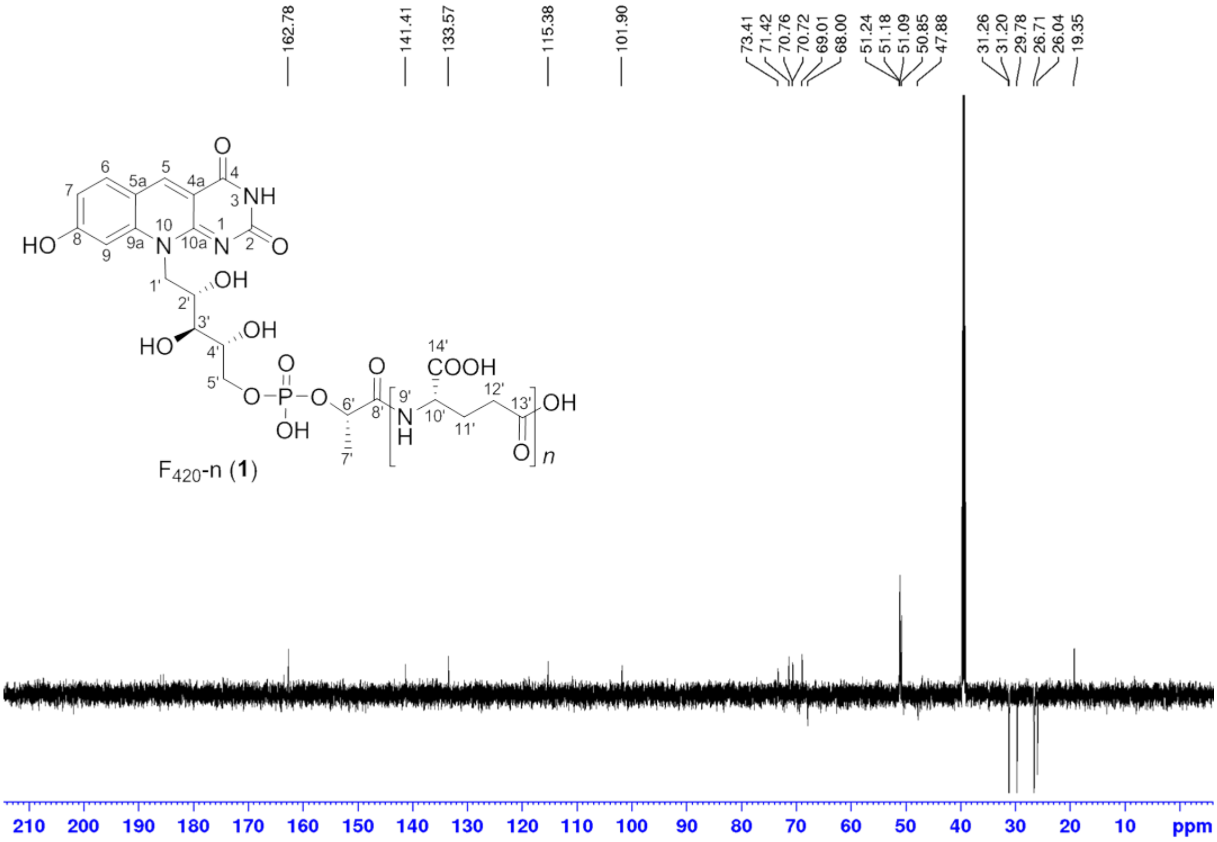

**Figure S22**. DEPT 135 spectrum of $F_{420}$-n (DMSO-$d_6$, 150 MHz, 300 K).

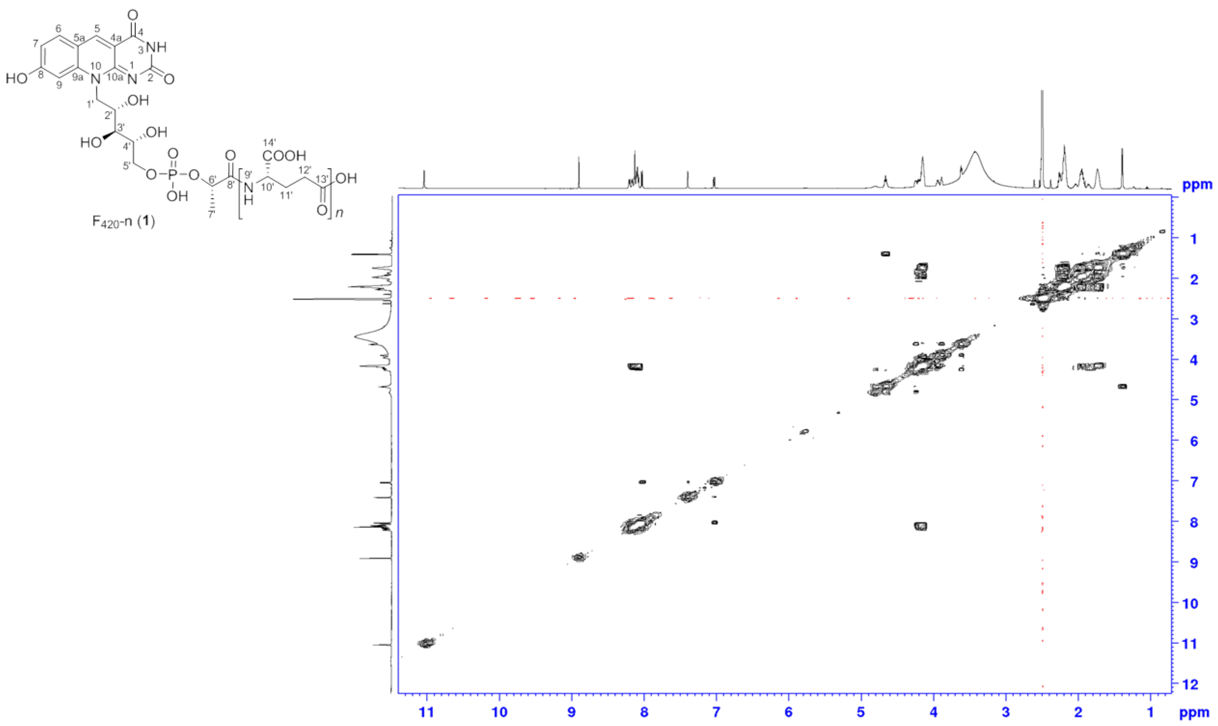

**Figure S23**. $^{1}H$-$^{1}H$ COSY spectrum of $F_{420}$-n (DMSO-$d_6$, 600 MHz, 300 K).

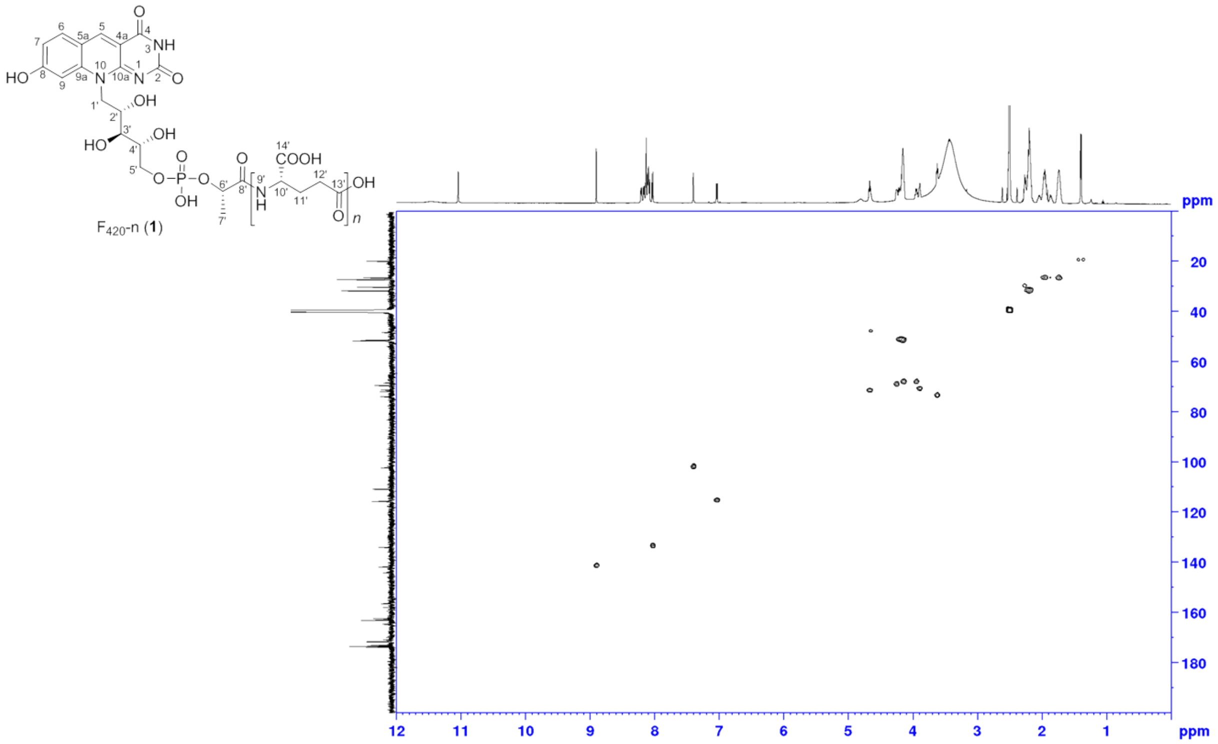

**Figure S24**. $^{1}H$-$^{13}C$ HSQC spectrum of $F_{420}$-n (DMSO-$d_6$, 600 MHz, 300 K).

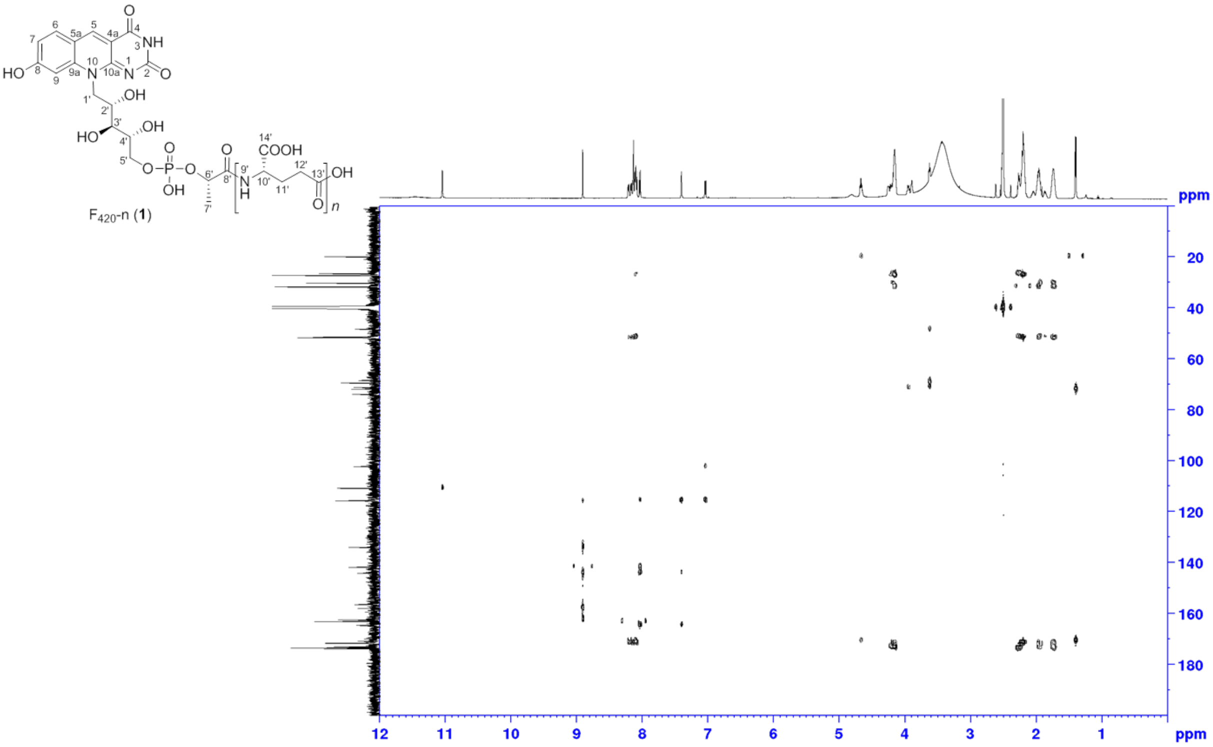

**Figure S25**. $^{1}H$-$^{13}C$ HMBC spectrum of $F_{420}$-n (DMSO-$d_6$, 600 MHz, 300 K).

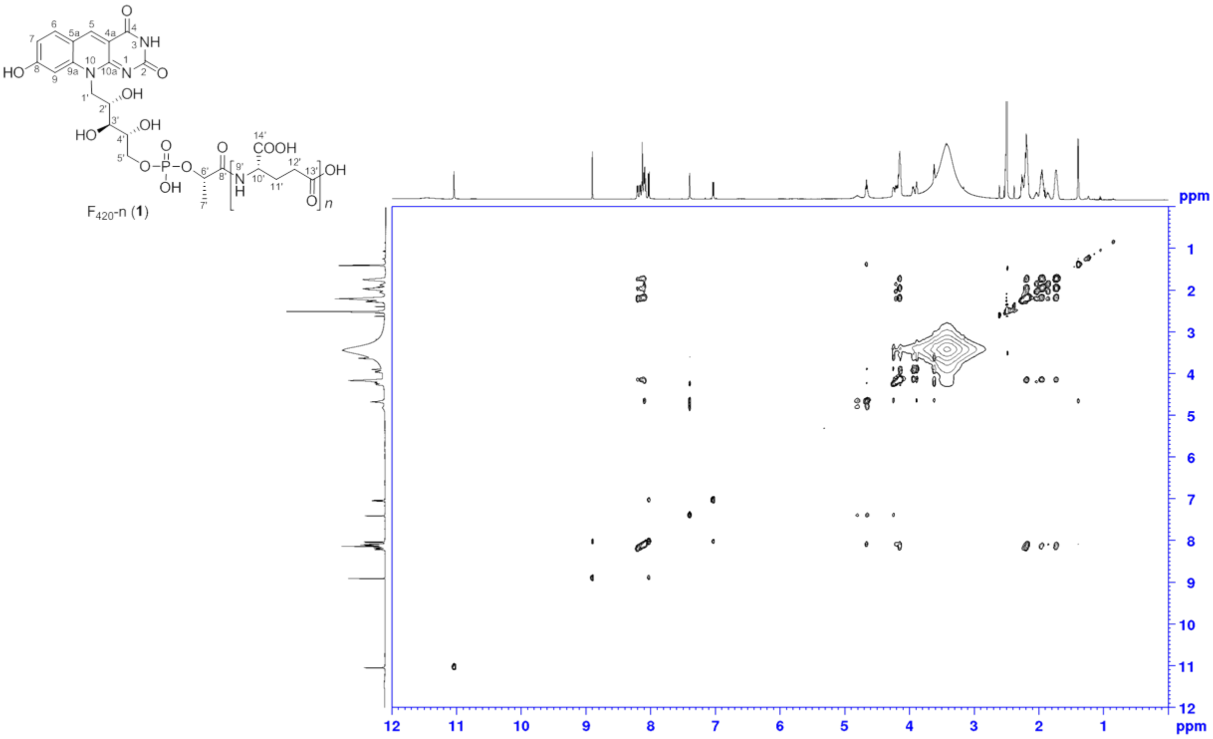

**Figure S26**. NOESY spectrum of $F_{420}$-n (DMSO-$d_6$, 600 MHz, 300 K).

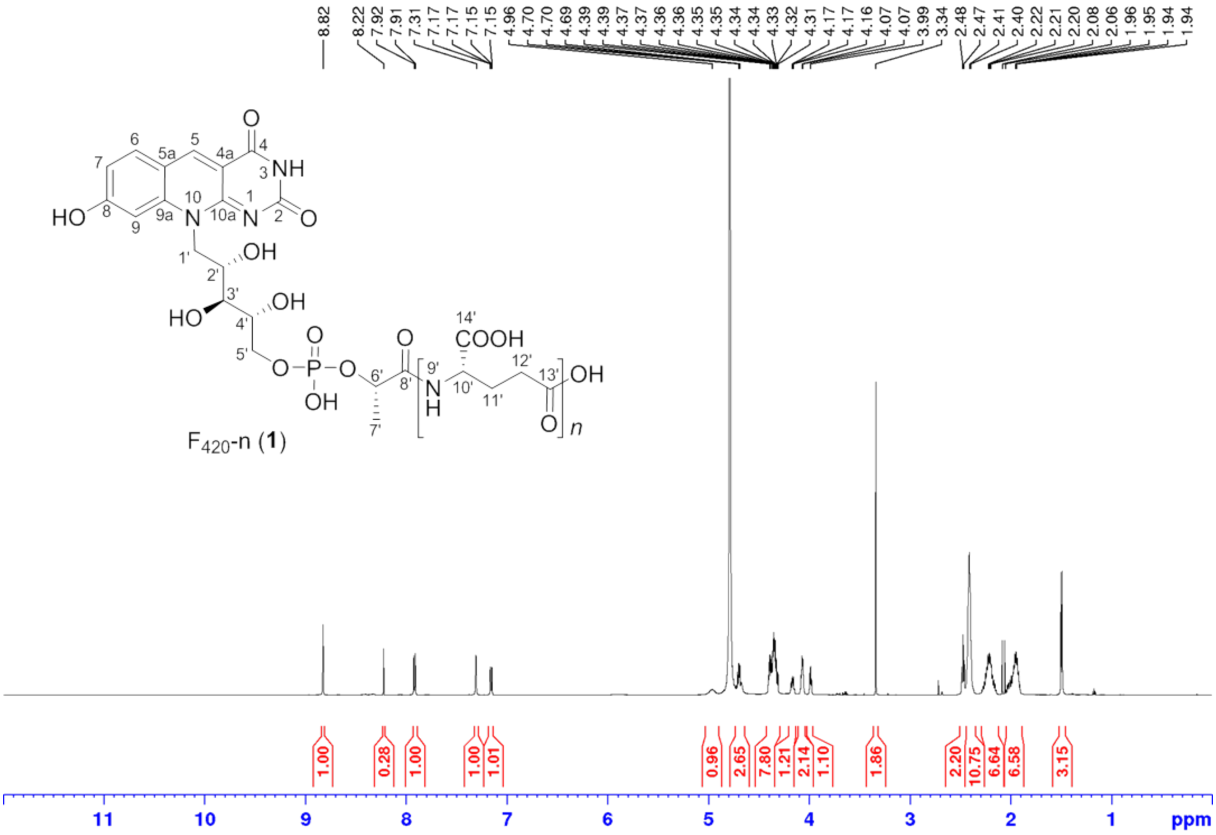

**Figure S27**. $^{1}$H NMR spectrum of $F_{420}$-n ($D_2O$, 600 MHz, 300 K).

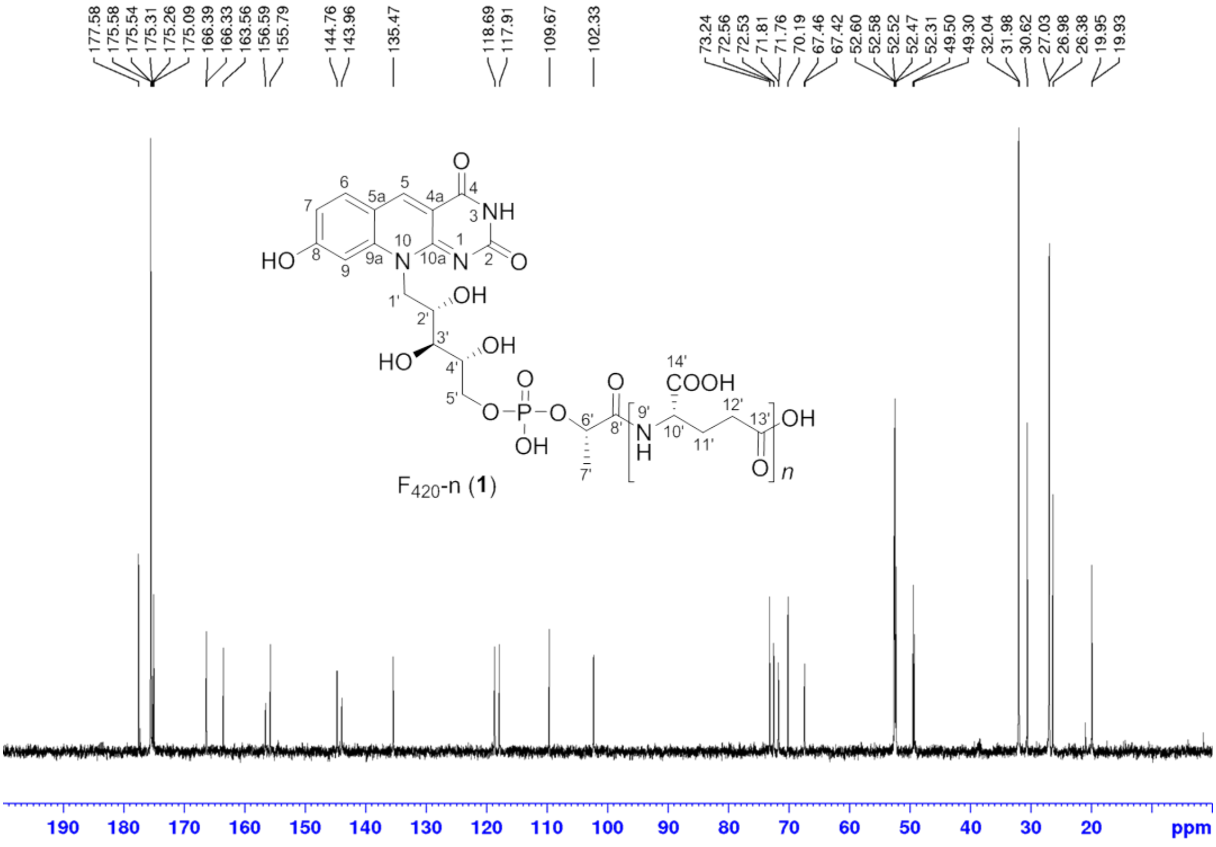

**Figure S28**. $^{13}$C NMR spectrum of $F_{420}$-n ($D_2O$, 150 MHz, 300 K).

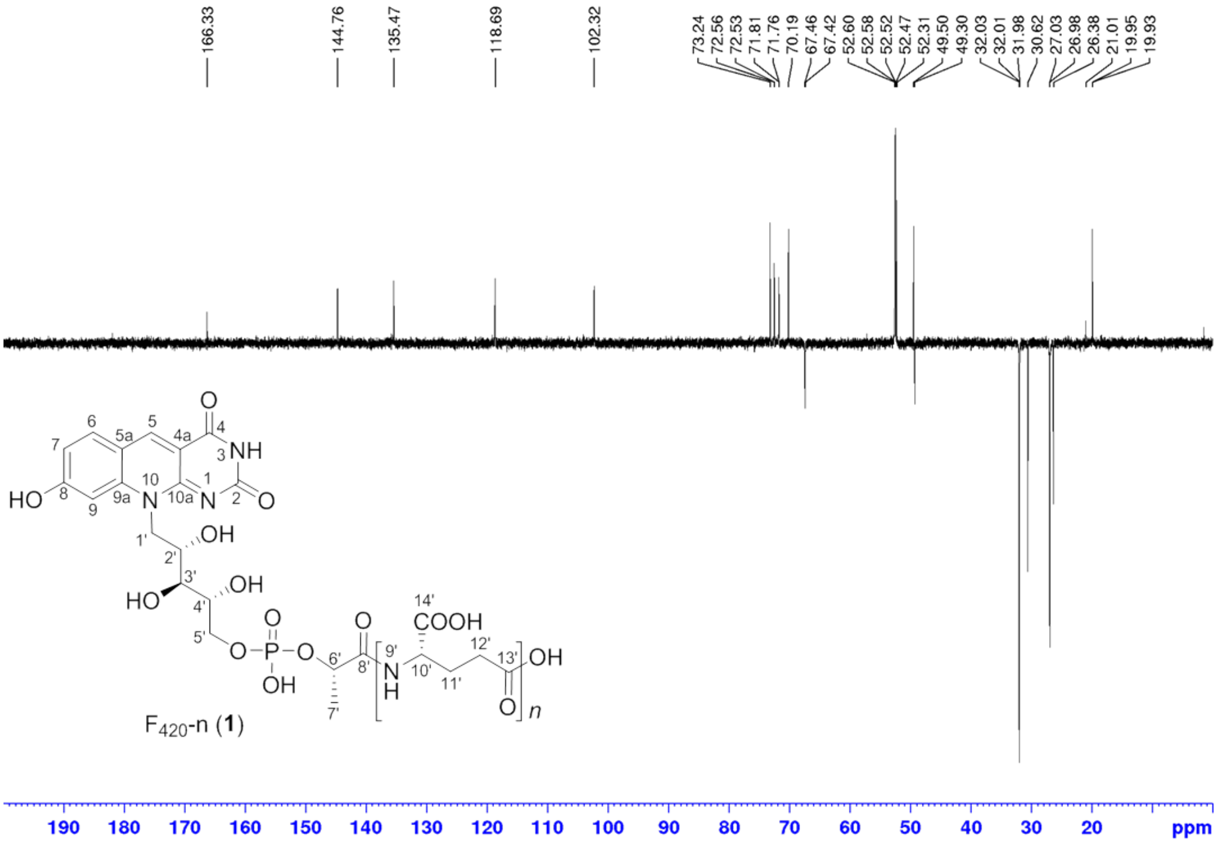

**Figure S29**. DEPT 135 spectrum of $F_{420}$-n ($D_2O$, 150 MHz, 300 K).

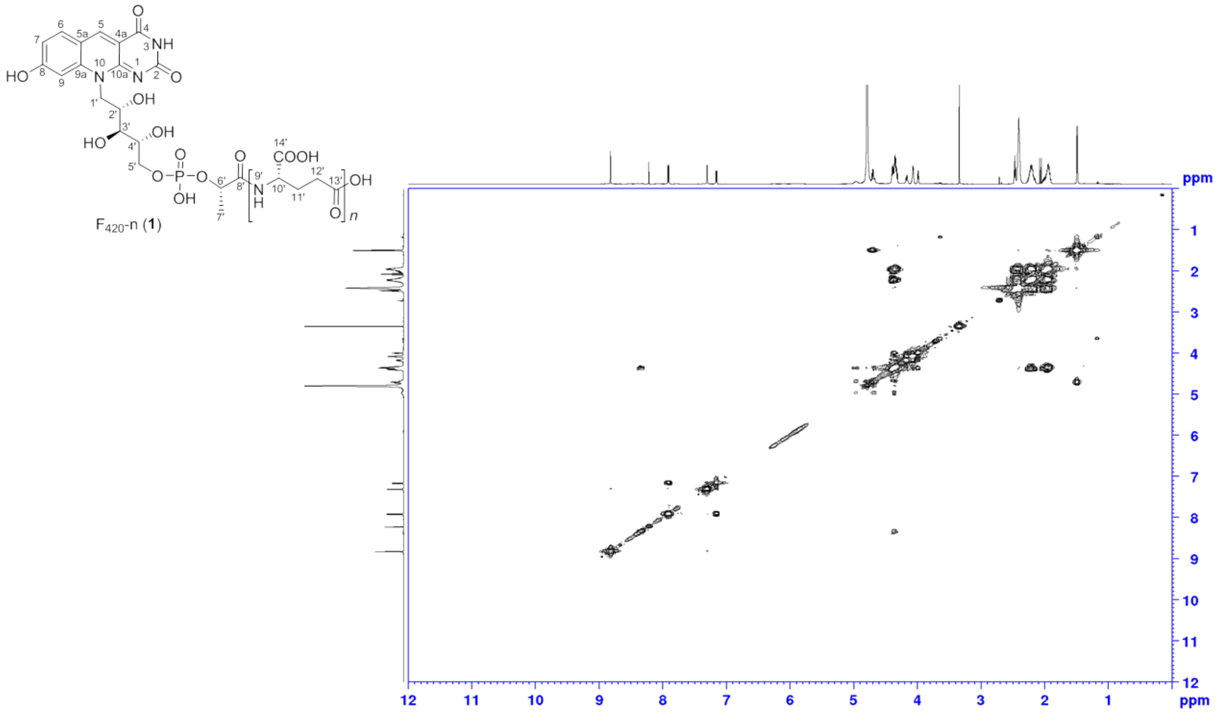

**Figure S30**. $^1$H-$^1$H COSY spectrum of $F_{420}$-n ($D_2O$, 600 MHz, 300 K).

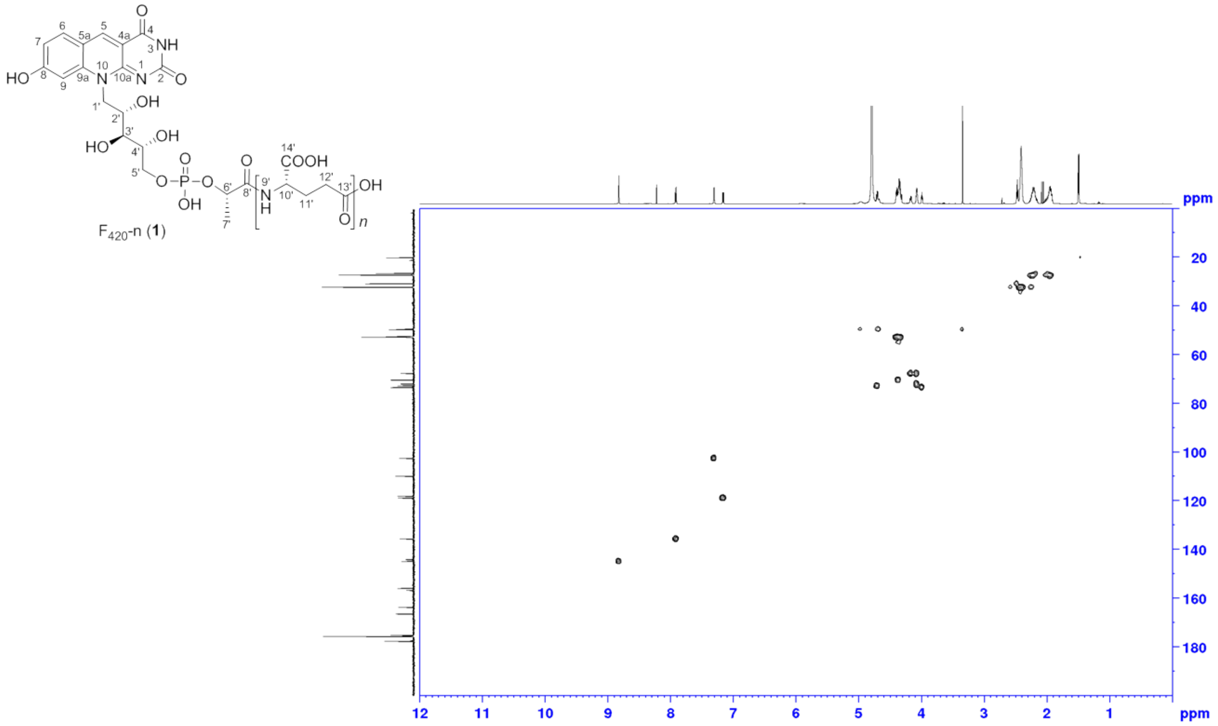

**Figure S31**. $^{1}H$-$^{13}C$ HSQC spectrum of $F_{420}$-n ($D_2O$, 600 MHz, 300 K).

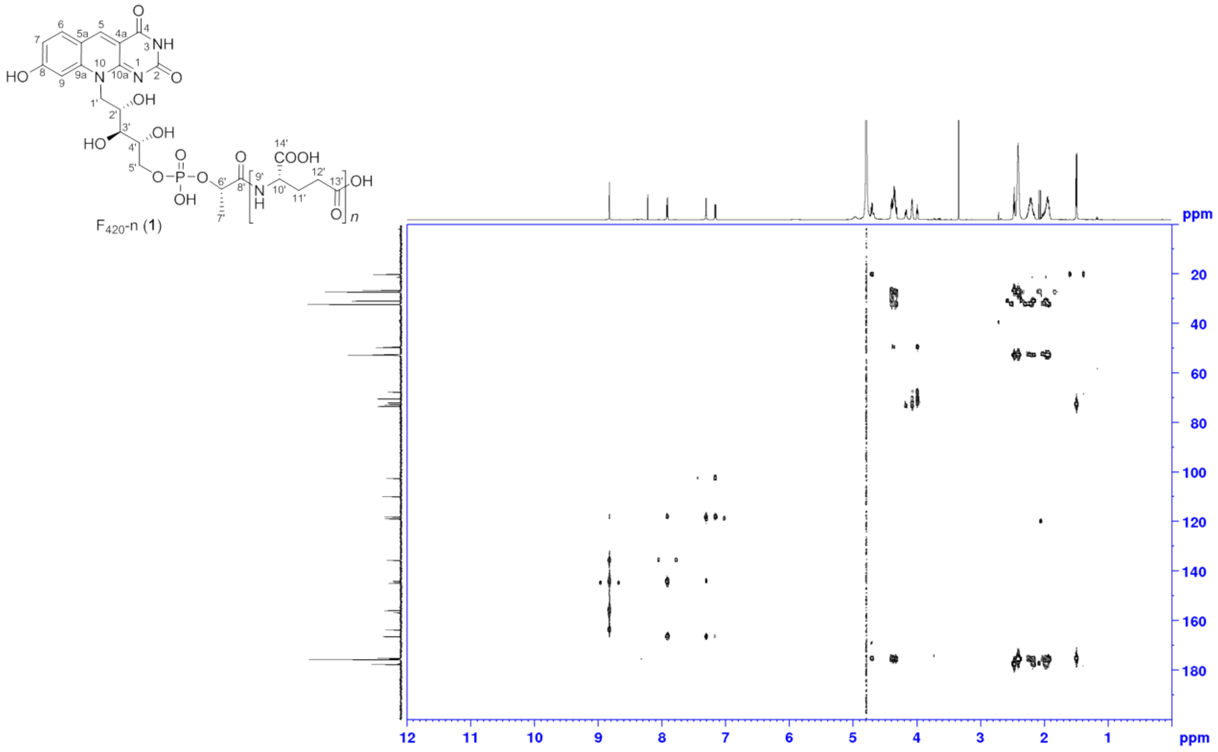

**Figure S32**. $^{1}H$-$^{13}C$ HMBC spectrum of $F_{420}$-n ($D_2O$, 600 MHz, 300 K).

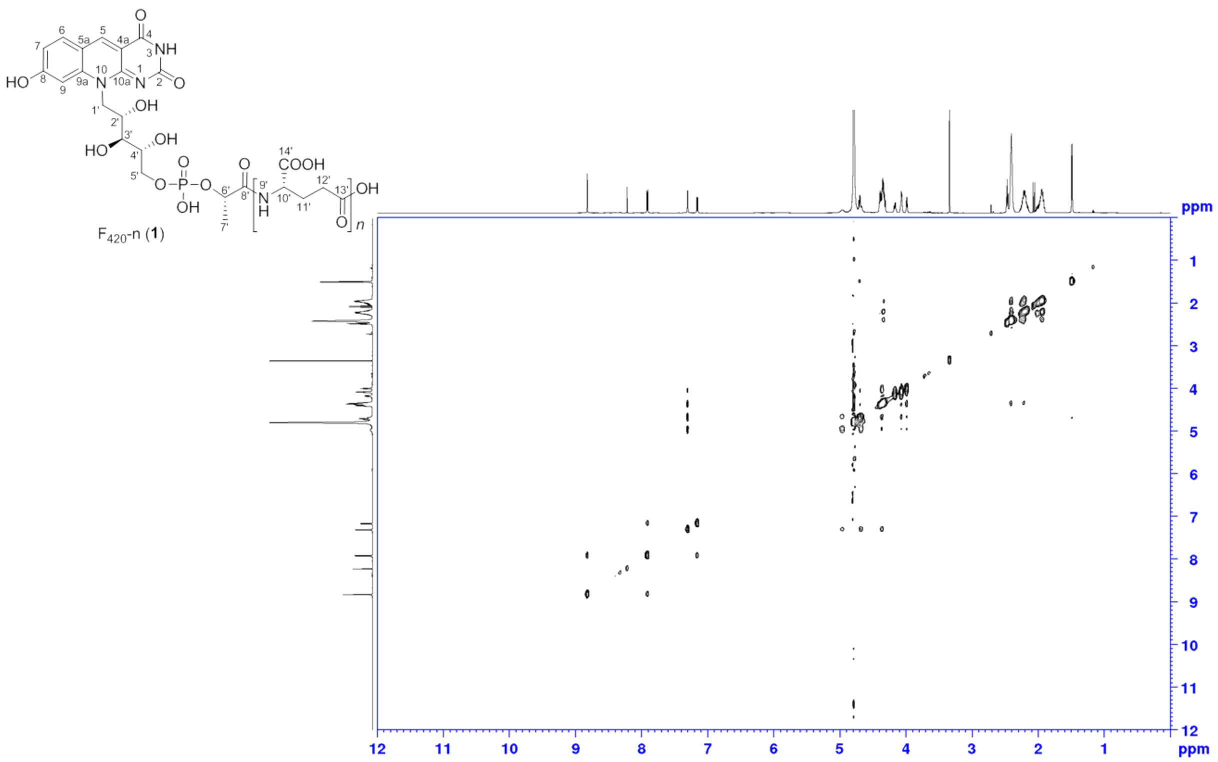

**Figure S33**. NOESY spectrum of $F_{420}$-n ($D_2O$, 600 MHz, 300 K).

*2.3.2 Structure elucidation of 3PG-$F_{420}$-0*

3PG-$F_{420}$ species were isolated from large-scale fermentations of *E. coli* / pDB045 as described in Experimental Procedures (section 1.9). The elution from the anion-exchange column is depicted in Figure S34.

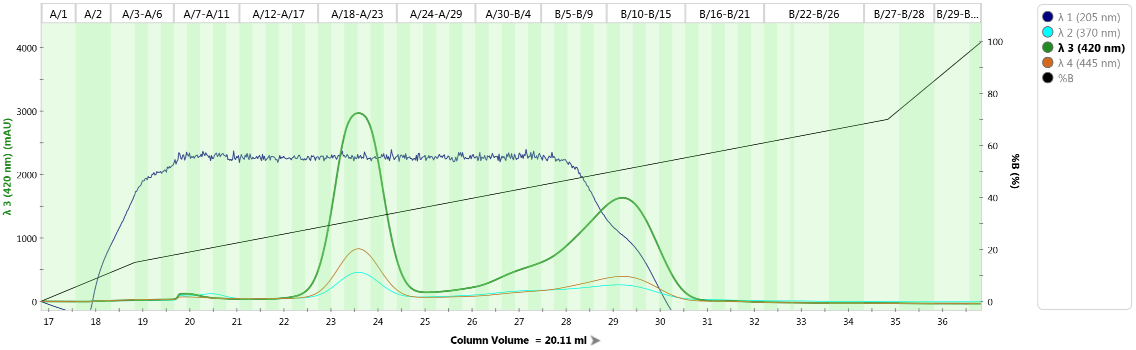

**Figure S34** Preparative ion exchange chromatography of *E. coli* / pDB045 lysate on a NGC Quest FPLC (Bio-Rad) equipped with a HiPrep QFF column (GE Healthcare) pre-equilibrated with sodium phosphate buffer (25mM, pH 7.4). A gradient of NaCl was used to elute different 3PG-$F_{420}$-species as peaks monitored by UV/VIS absorption at 420 nm (3PG-$F_{420}$-0: 22.5 to 25 CV, 3PG-$F_{420}$-n: 25-31 CV).

**3PG-$F_{420}$-0 (5)** was obtained as golden yellow solid. Due to the solubility, 1D and 2D NMR spectra were recorded in 0.1% $ND_3$ in $D_2O$. The molecular formula of 3PG-$F_{420}$-0 (**5**) was determined as $C_{19}H_{22}O_{13}N_3P$ based on the ESI-HRMS analysis (*m/z* 532.09534 ($[M+H]^+$ calcd. 532.09630 Δ = – 1.81 ppm) and $^1$H and $^{13}$C NMR spectra. Detailed comparison of the $^1$H NMR and $^{13}$C NMR spectra with classical **$F_{420}$**-n revealed the similar structure feature of 5-deazaflavin moiety and the ribityl moiety, based on comparable $^{13}$C resonance, with the slight chemical shift on the 5-deazaflavin moiety due to the ammonium salt and basic condition. The observation of a missing methyl group from the lactyl moiety and additional oxygenated methylene group ($\delta_{H\text{-}7'}$ 4.13/4.03 ppm, $\delta_{C\text{-}7'}$ 68.15 ppm) in 3PG-**$F_{420}$**-0 (**5**), and the shift of oxymethine group ($\delta_{H\text{-}6'}$ 4.21 ppm, $\delta_{C\text{-}6'}$ 72.10 ppm) suggested the hydroxy group on the lactyl moiety. HMBC correlations of H-6' and H-7'a to C-8' further suggested the presence of a glyceryl moiety. The doublets of C-4', C-5', C-6' and C-7' indicated the $^{13}$C–$^{31}$P coupling ($^3J_{C\text{-}4'\text{–}P}$ = 8.34 Hz, $^2J_{C\text{-}5'\text{–}P}$ = 5.56 Hz, $^3J_{C\text{-}6'\text{–}P}$ = 8.34 Hz, $^2J_{C\text{-}7'\text{–}P}$ = 5.37 Hz) between the carbon atoms of the ribityl and glyceryl moieties assigned to these resonances and the phosphorus of the phosphate group. $^{13}$C–$^{31}$P coupling also suggested the presence of 3-phosphoglycerate, with the relatively smaller $^2J_{C\text{-}P}$ and larger $^3J_{C\text{-}P}$ value. This hypothesis was confirmed by the measurement of $^2J_{C\text{-}P}$ and $^3J_{C\text{-}P}$ value from 2-phosphoglycerate and 3-phosphoglycerate. The coupling constants of 3PG-**$F_{420}$**-0(**5**) exactly matched to 3-phosphoglycerate, $^1$H and $^{13}$C chemical shift were similar. Therefore, the planar structure of 3PG-

$F_{420}$-0 (**5**) is assigned as glyceric acid phosphodiester of 7,8-dedemethyl-8-hydroxy-5-deazariboflavin 5'-phosphate. (**Table S7** and **S8**)

**Spectral Data:** 3PG-$F_{420}$-0 (**5**): golden yellow solid; UV ($H_2O$, pH 7.5) $\lambda_{max}$ 250, 420 nm; IR (ATR) $\nu_{max}$ 3233, 2954, 2926, 2867, 2358, 2330, 1587, 1506, 1428, 1354, 1152, 1035, 853, 800 $cm^{-1}$; NMR spectral data, see Table S7; ESI-HRMS $[M+H]^+$ *m/z* 532.09534 (calcd. for $C_{19}H_{23}O_{13}N_3P$, 532.09630).

**Table S7.** NMR Data (0.1% $ND_3$ in $D_2O$, at 300 K) for 3PG-$F_{420}$-0 (**5**).[a]

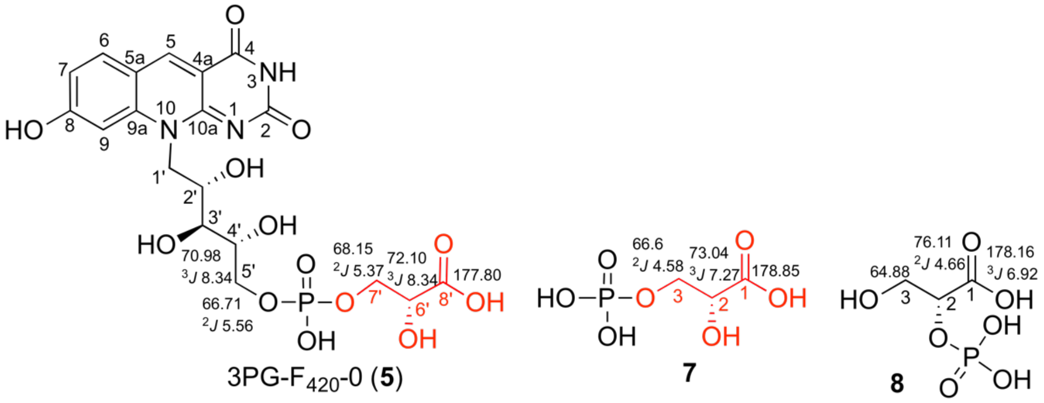

| | 3PG-$F_{420}$-0 (**5**) | | | | |
|---|---|---|---|---|---|
| position | $\delta_C$, mult.[b] (*J* in Hz)[c] | $\delta_H$, mult. (*J* in Hz) | COSY | HMBC | NOESY |
| *N* (1) | | | | | |
| 2 | 159.33, qC | | | | |
| *N*H (3) | | | | | |
| 4 | 165.45, qC | | | | |
| 4a | 104.13, qC | | | | |
| 5 | 140.80, CH | 8.56, s | | 4, 6, 9a, 10a | |
| 5a | 115.41, qC | | | | |
| 6 | 134.51, CH | 7.72, d (*9.06*) | 7 | 5, 5a, 9a, 8 | |
| 7 | 123.52, CH | 6.82, d (*8.98*) | 6 | 5a, 9 | |
| 8 | 179.48, qC | | | | |
| 9 | 102.74, CH | 6.77, s | | 5a, 7, 9a, 8 | |
| 9a | 145.56, qC | | | | |
| 10 | | | | | |
| 10a | 157.75, qC | | | | |
| 1'a | 46.31, $CH_2$ | 5.07, br s | 1'b, 2' | | |
| 1'b | | 4.56, br d (*13.14*) | 1'a | | |
| 2' | 69.31, CH | 4.41, m | 1'b, 3' | | |
| 3' | 72.54, CH | 3.96, m | 2', 4' | 1', 2', 4', 5' | |
| 4' | 70.98, CH, d (*8.34*) | 4.03, m | 3' | 2' | |
| 5'a | 66.71, $CH_2$, d (*5.56*) | 4.13, m | 5'b | 3' | |
| 5'b | | 4.03, m | 5'a | 3', 4' | |
| 6' | 72.10, CH, d (*8.34*) | 4.21, m | 7'b | 8' | |
| 7' | 68.15, $CH_2$, d (*5.37*) | 4.13, m | 7'b | 8' | |
| | | 4.03, m | 6, 7'a | | |
| 8' | 177.80, qC | | | | |

[a] 600 MHz for $^{1}H$ NMR and 150 MHz for $^{13}C$ NMR
[b] numbers of attached protons were determined by analysis of 2D spectra
[c] coupling constant indicated $^{13}C$–$^{31}P$ coupling

**Table S8.** NMR Data (0.1% $ND_3$ in $D_2O$, at 300 K) for 3-phospho-D-glyceric acid (**7**) and 2-phospho-D-glyceric acid (**8**).

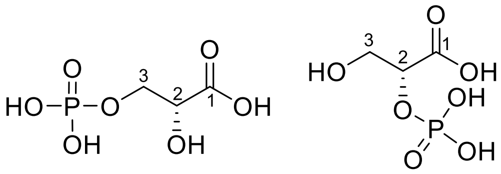

**7** **8**

| | 3-phospho-d-glyceric acid (**7**) | | 2-phospho-D-glyceric acid (**8**) | |
|---|---|---|---|---|
| position | $\delta_C$, mult.[b] (*J* in Hz)[c] | $\delta_H$, mult. (*J* in Hz) | $\delta_C$, mult.[b] (*J* in Hz)[c] | $\delta_H$, mult. (*J* in Hz) |
| 1 | 178.85, qC, s | | 178.16, qC, d (*6.92*) | |
| 2 | 73.04, CH, d (*7.07*) | 4.18, dd (*6.64, 2.70*) | 76.11, CH, d (*4.66*) | 4.47, dd (*5.81, 2.86*) |
| 3a | 66.65, $CH_2$, d (*4.58*) | 4.02, ddd (*11.07 6.14, 2.79*) | 64.88, $CH_2$, s | 3.90, dd (*11.62, 5.81*) |
| 3b | | 3.86, ddd (*11.20, 6.57,* [illegible] | | 3.79, dd (*11.62,3.02*) |

[a] 600 MHz for $^1$H NMR and 150 MHz for $^{13}$C NMR
[b] numbers of attached protons were determined by analysis of 2D spectra.
[c] coupling constant indicated $^{13}$C–$^{31}$P coupling

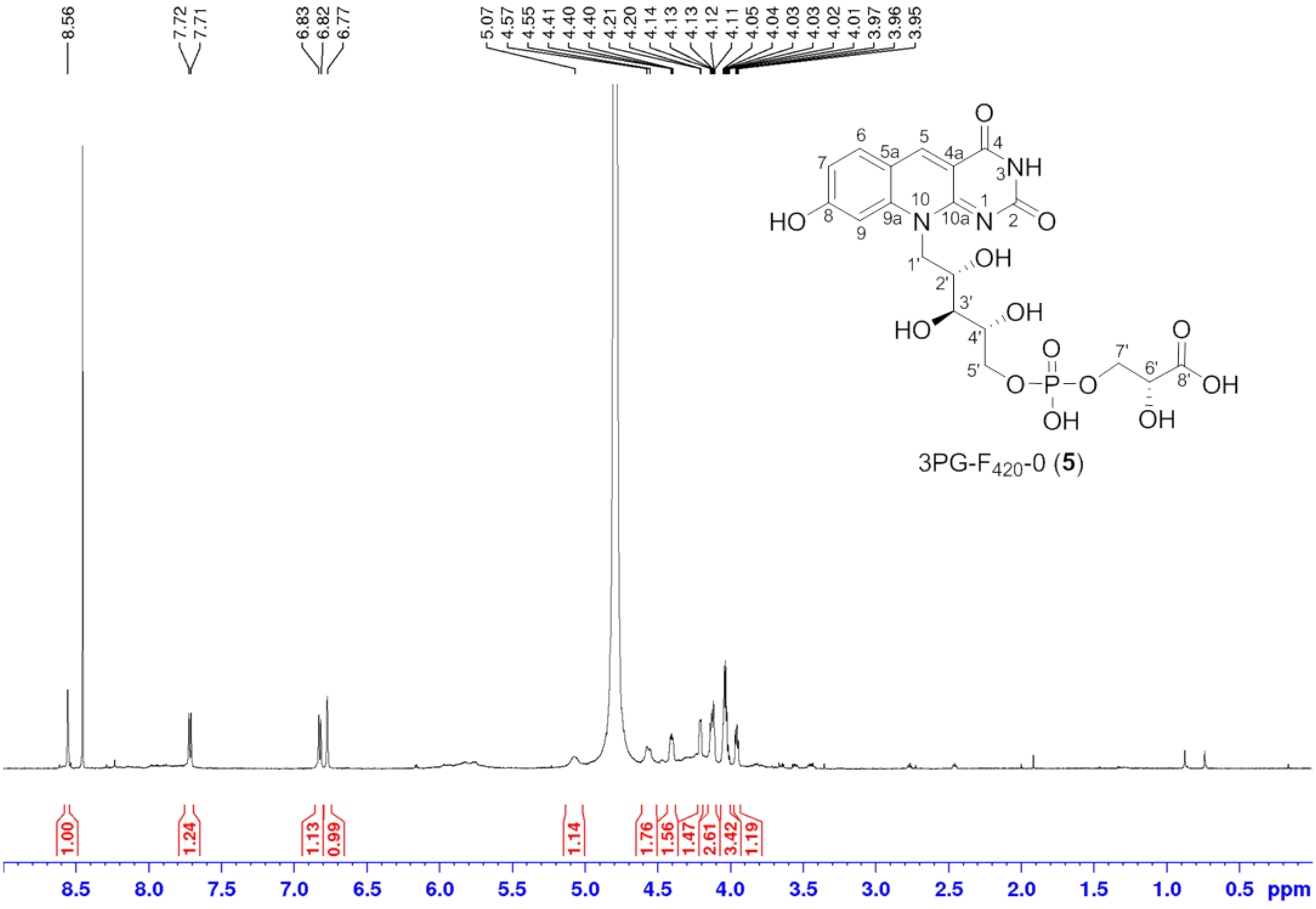

**Figure S35**. $^1$H NMR spectrum of 3PG-$F_{420}$-0 (**5**) (0.1% $ND_3$ in $D_2O$, 600 MHz, 300 K).

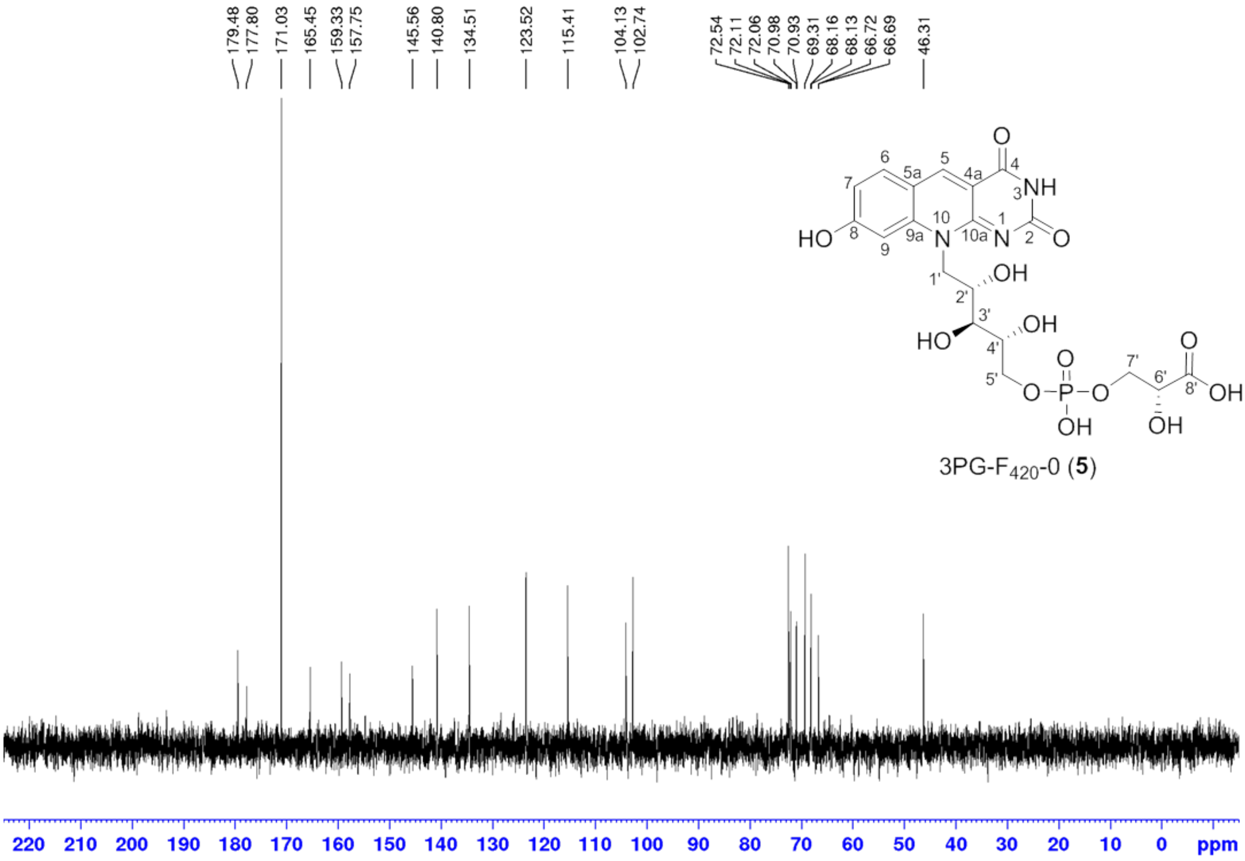

**Figure S36**. $^{13}C$ NMR spectrum of 3PG-$F_{420}$-0 (**5**) (0.1% $ND_3$ in $D_2O$, 150 MHz, 300 K).

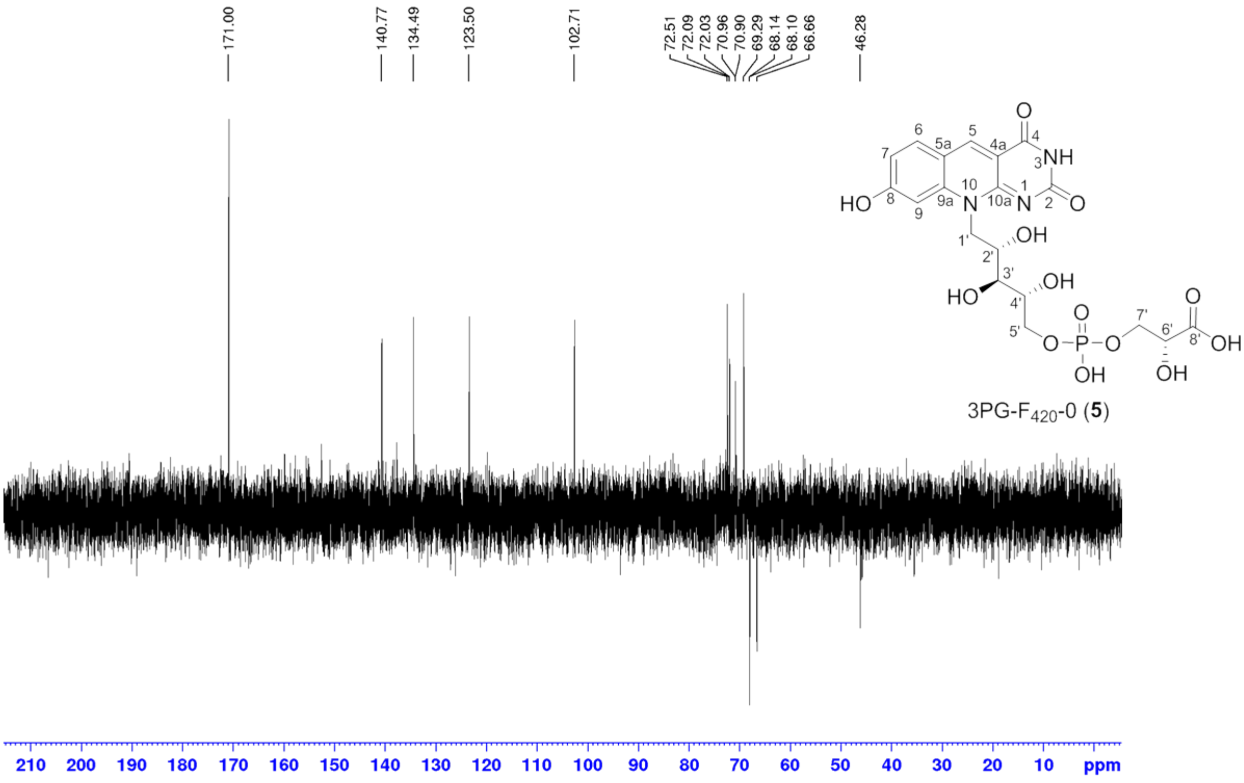

**Figure S37**. DEPT 135 spectrum of 3PG-$F_{420}$-0 (**5**) (0.1% $ND_3$ in $D_2O$, 150 MHz, 300 K).

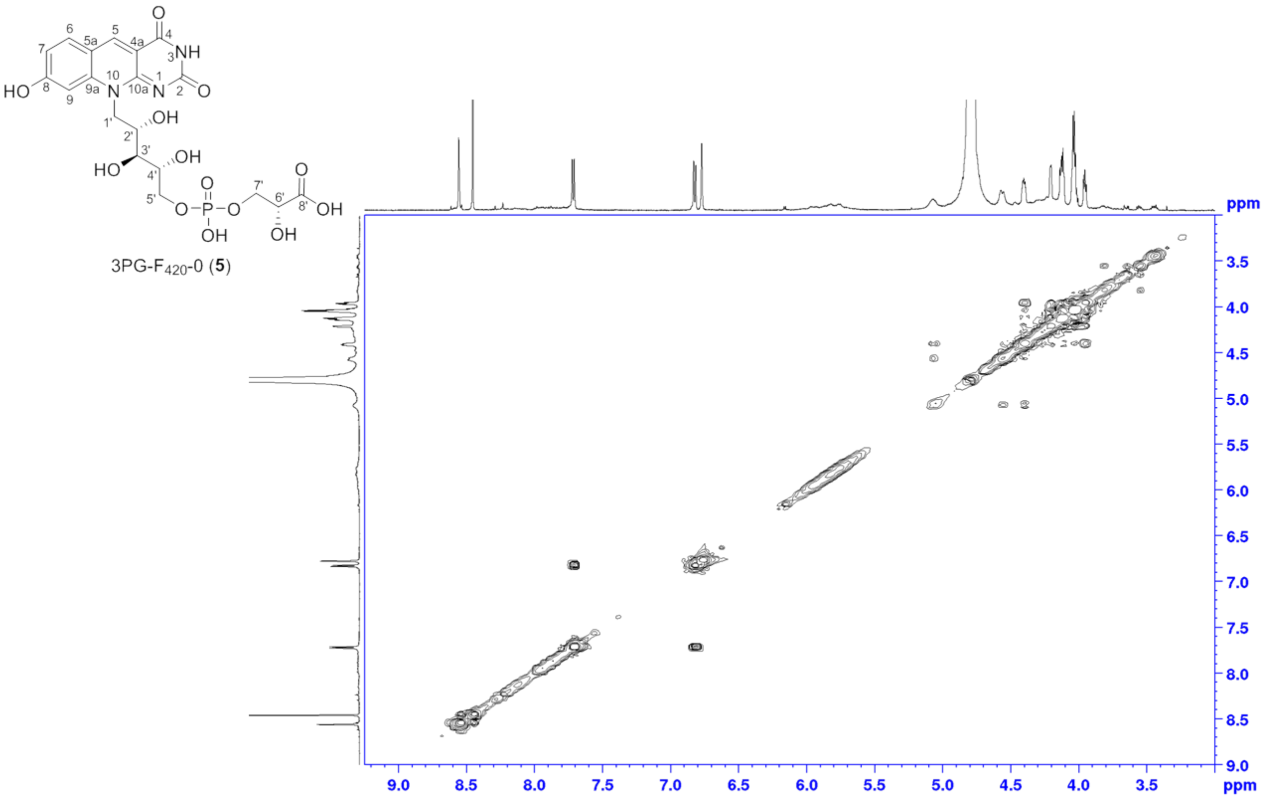

**Figure S38**. [1]H-[1]H COSY spectrum of 3PG-$F_{420}$-0 (**5**) (0.1% $ND_3$ in $D_2O$, 600 MHz, 300 K).

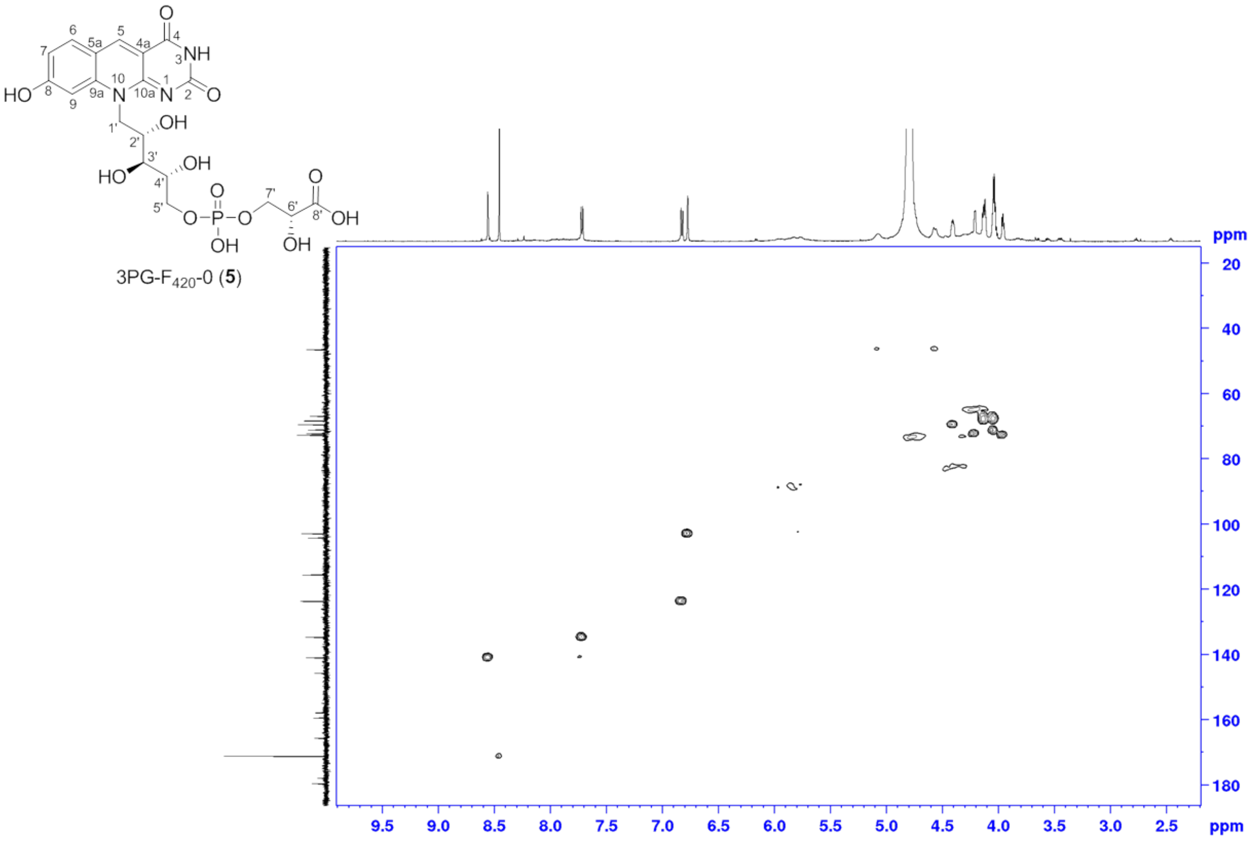

**Figure S39**. [1]H-[13]C HSQC spectrum of 3PG-$F_{420}$-0 (**5**) (0.1% $ND_3$ in $D_2O$, 600 MHz, 300 K).

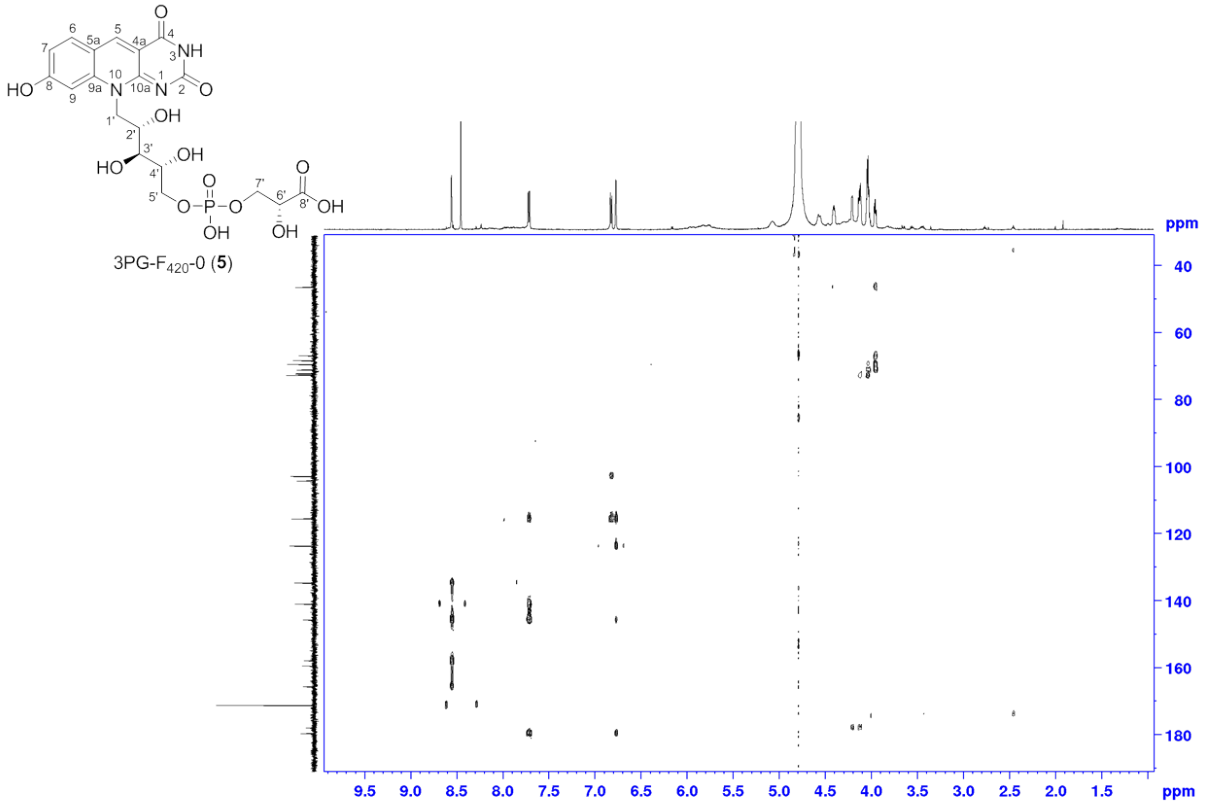

**Figure S40**. $^{1}H$-$^{13}C$ HMBC spectrum of 3PG-$F_{420}$-0 (**5**) (0.1% $ND_3$ in $D_2O$, 600 MHz, 300 K).

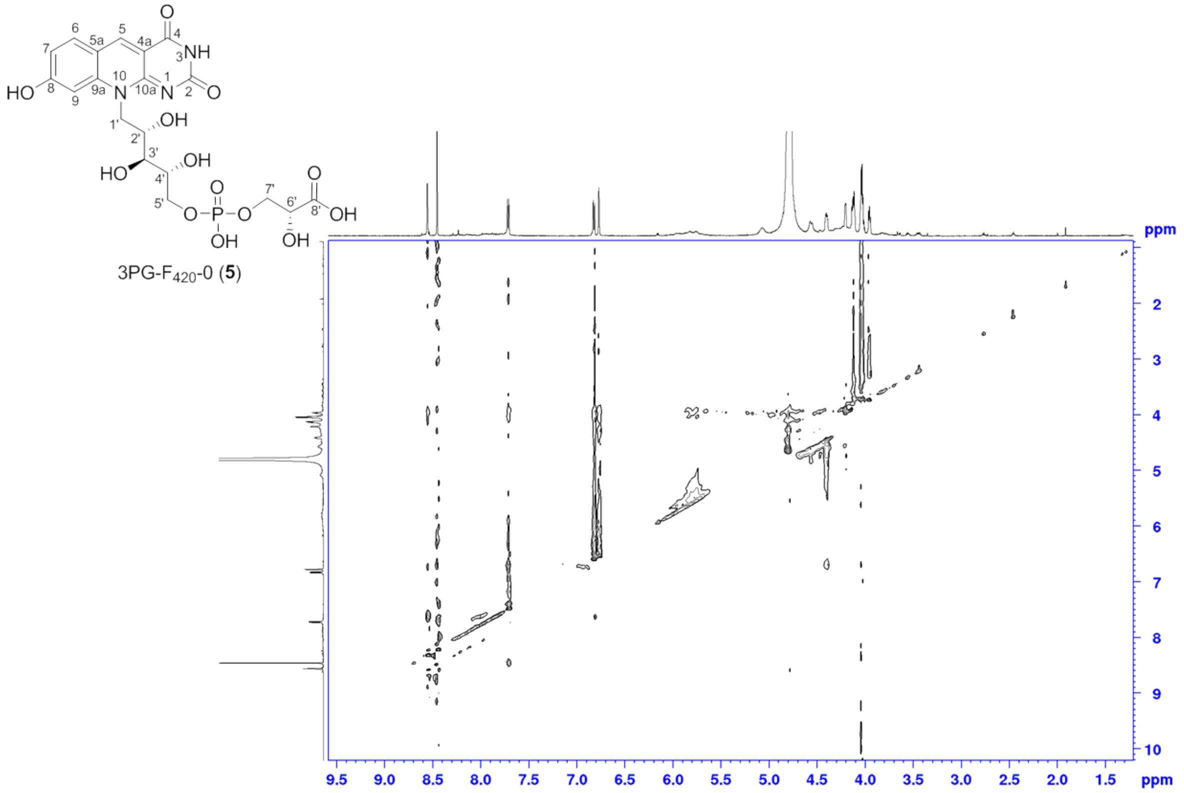

**Figure S41**. NOESY spectrum of 3PG-$F_{420}$-0 (**5**) (0.1% $ND_3$ in $D_2O$, 600 MHz, 300 K).

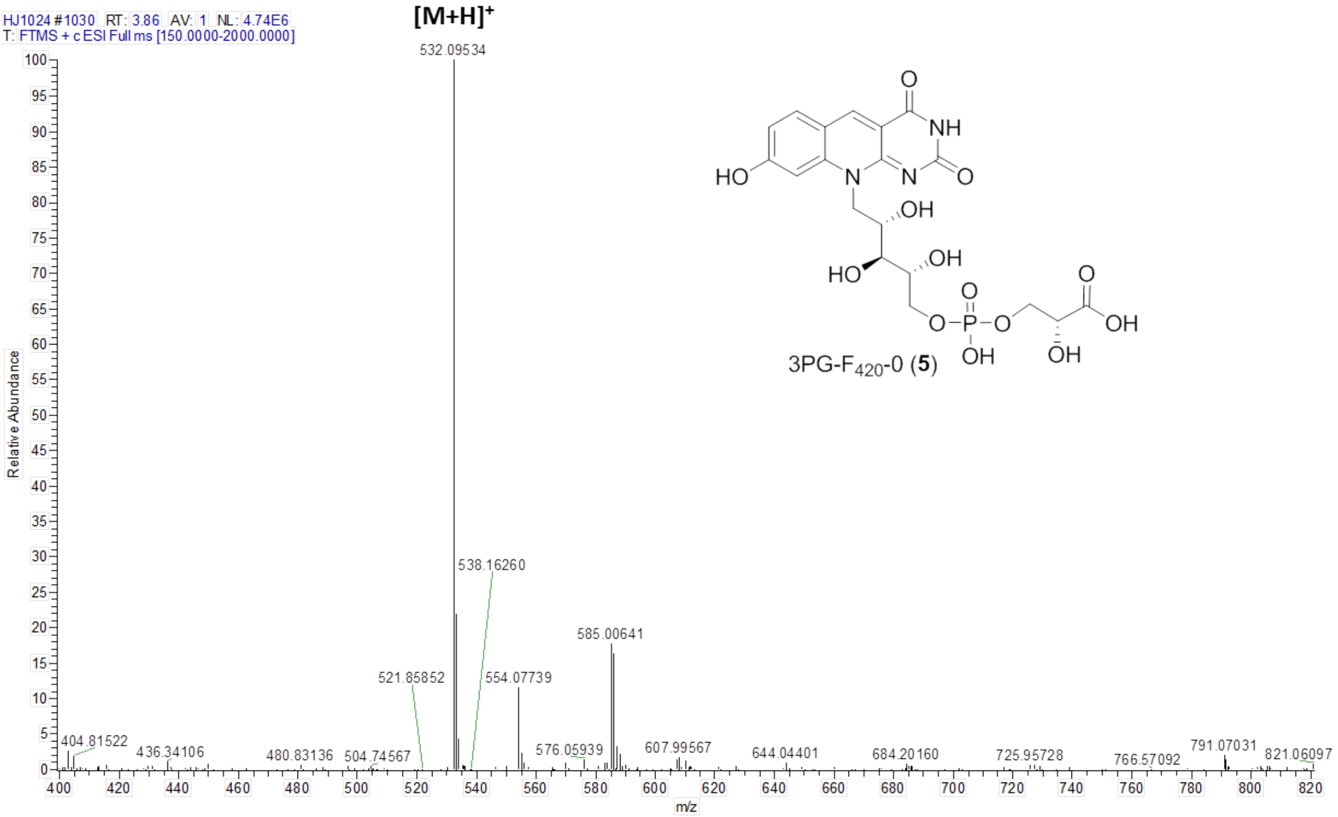

**Figure S42.** ESI-HRMS spectrum of 3PG-$F_{420}$-0 (**5**).

*2.3.3 Structure elucidation of 3PG-$F_{420}$-n*

**3PG-F420-n (4):** 3PG-$F_{420}$-n was obtained as golden yellow solid. 1D and 2D NMR spectra were recorded in 0.1% $ND_3$ in $D_2O$ as well. The composition of mixed 3PG-$F_{420}$-n was determined as 3PG-$F_{420}$-1 with the molecular formula of $C_{24}H_{29}O_{16}N_4P$ based on the ESI-HRMS analysis (*m*/*z* 661.13757 ($[M+H]^+$ calcd. 661.13889 $\Delta$ = – 2.00 ppm); 3PG-$F_{420}$-2 with the molecular formula of $C_{29}H_{36}O_{19}N_5P$ based on the ESI-HRMS analysis (*m*/*z* 790.17981 ($[M+H]^+$ calcd. 790.18149 $\Delta$ = – 2.12 ppm); 3PG-$F_{420}$-3 with the molecular formula of $C_{34}H_{43}O_{22}N_6P$ based on the ESI-HRMS analysis (*m*/*z* 919.22223 ($[M+H]^+$ calcd. 919.22408 $\Delta$ = – 2.01 ppm); 3PG-$F_{420}$-4 with the molecular formula of $C_{39}H_{50}O_{25}N_7P$ based on the ESI-HRMS analysis (*m*/*z* 1048.26489 ($[M+H]^+$ calcd. 1048.26667 $\Delta$ = – 1.70 ppm); 3PG-$F_{420}$-5 with the molecular formula of $C_{44}H_{57}O_{28}N_8P$ based on the ESI-HRMS analysis (*m*/*z* 1177.30750 ($[M+H]^+$ calcd. 1177.30927 $\Delta$ = – 1.50 ppm); 3PG-$F_{420}$-6 with the molecular formula of $C_{49}H_{64}O_{31}N_9P$ based on the ESI-HRMS analysis (*m*/*z* 1306.34961 ($[M+H]^+$ calcd. 1306.35186 $\Delta$ = – 1.72 ppm). Detailed comparison of $^1H$ NMR data of 3PG-$F_{420}$-n with 3PG-$F_{420}$-0 (**5**) and $\mathbf{F_{420}}$-n revealed the signals from glutamyl moiety ($\delta_{H'}$ 1.86–2.35 ppm; $\delta_H$ 4.25–4.25 ppm).

**Spectral Data:** 3PG-$F_{420}$-n**:** golden yellow solid; UV ($H_2O$, pH 7.5) $\lambda_{max}$ 250, 420 nm; IR (ATR) $\nu_{max}$ 3738, 3339, 2926, 2356, 1678, 1581, 1512, 1365, 1206, 1181, 1138, 1014, 841, 805, 769 722, 664 $cm^{-1}$;

**Table S9.** NMR Data (0.1% $ND_3$ in $D_2O$, at 300 K) for 3PG-$F_{420}$-n .[a]

| | 3PG-$F_{420}$-n | | | | |
|---|---|---|---|---|---|
| position | $\delta_C$, mult.[b] (*J* in Hz)[c] | $\delta_H$, mult. (*J* in Hz) | COSY | HMBC | NOESY |
| *N*H (1) | | | | | |
| 2 | 159.37, qC | | | | |
| *N*H (3) | | | | | |
| 4 | 165.59, qC | | | | |
| 4a | 104.29, qC | | | | |
| 5 | 140.93, CH | 8.61, s | | 4, 6, 9a, 10a | 6 |
| 5a | 115.47, qC | | | | |
| 6 | 134.55, CH | 7.75, d (*8.50*) | 7 | 5, 8, 9a | 5, 7 |
| 7 | 123.50, CH | 6.84, d (*8.60*) | 6 | 5a, 9 | 5 |
| 8 | 179.45, qC | | | | |
| 9 | 102.75, CH | 6.80, s | | 5a, 7, 9a | |
| 9a | 145.63, qC | | | | |
| 10 | | | | | |
| 10a | 157.86, qC | | | | |
| 1'a | 46.37, $CH_2$ | 5.09, br s | 1'b, 2' | | |
| 1'b | | 4.61, *d* (13.24) | 1'a | | |
| 2' | 69.40, CH | 4.41, m | 1'a, 3' | | |
| 3' | 72.60, CH | 3.95, t (*5.12*) | 2' | 1', 2', 4', 5' | |
| 4' | 70.94, $CH_2$, d, (*7.81*) | 4.04, m | | | |
| 5'a | 66.95, $CH_2$, d, (*4.82*) | 4.13, m | | | |
| 5'b | | 4.01, m | | | |
| 6' | 71.12, CH d, (*8.60*) | 4.42, br s | 7'a | 8' | |
| 7'a | 66.85, $CH_2$, d, (*3.82*) | 4.13, m | 6' | 8' | |
| 7'b | | 4.01, m | | | |
| 8' | 172.86, qC | | | | |
| *N*H (9') [d] | | | | | |
| 10' [d] | 53.64, CH | 4.32, m | 11'a, 11'b | 8', 11', 12' | |
| 11'a [d] | 27.86, $CH_2$ | 2.04, m | 10', 11b' | 10', 12', 13', 14' | |
| 11'b [d] | | 1.91, m | 10', 11'a | 10', 12', 13', 14' | |
| 12'a [d] | 33.57, $CH_2$ | 2.28, m | | 10', 11', 13' | |
| 12'b | | 2.22, m | | 10', 11', 13' | |
| 13' [d] | 181.56, qC | | | | |
| 14' [d] | 173.63, qC | | | | |

[a] 600 MHz for $^1H$ NMR and 150 MHz for $^{13}C$ NMR

[b] numbers of attached protons were determined by analysis of 2D spectra.

[c] coupling constant indicated $^{13}C$–$^{31}P$ coupling

[d] NMR resonance are overlapped in the five times glutamate chains and only one unit was presented.

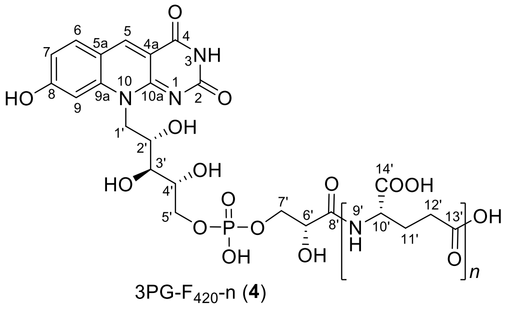

3PG-$F_{420}$-n (**4**)

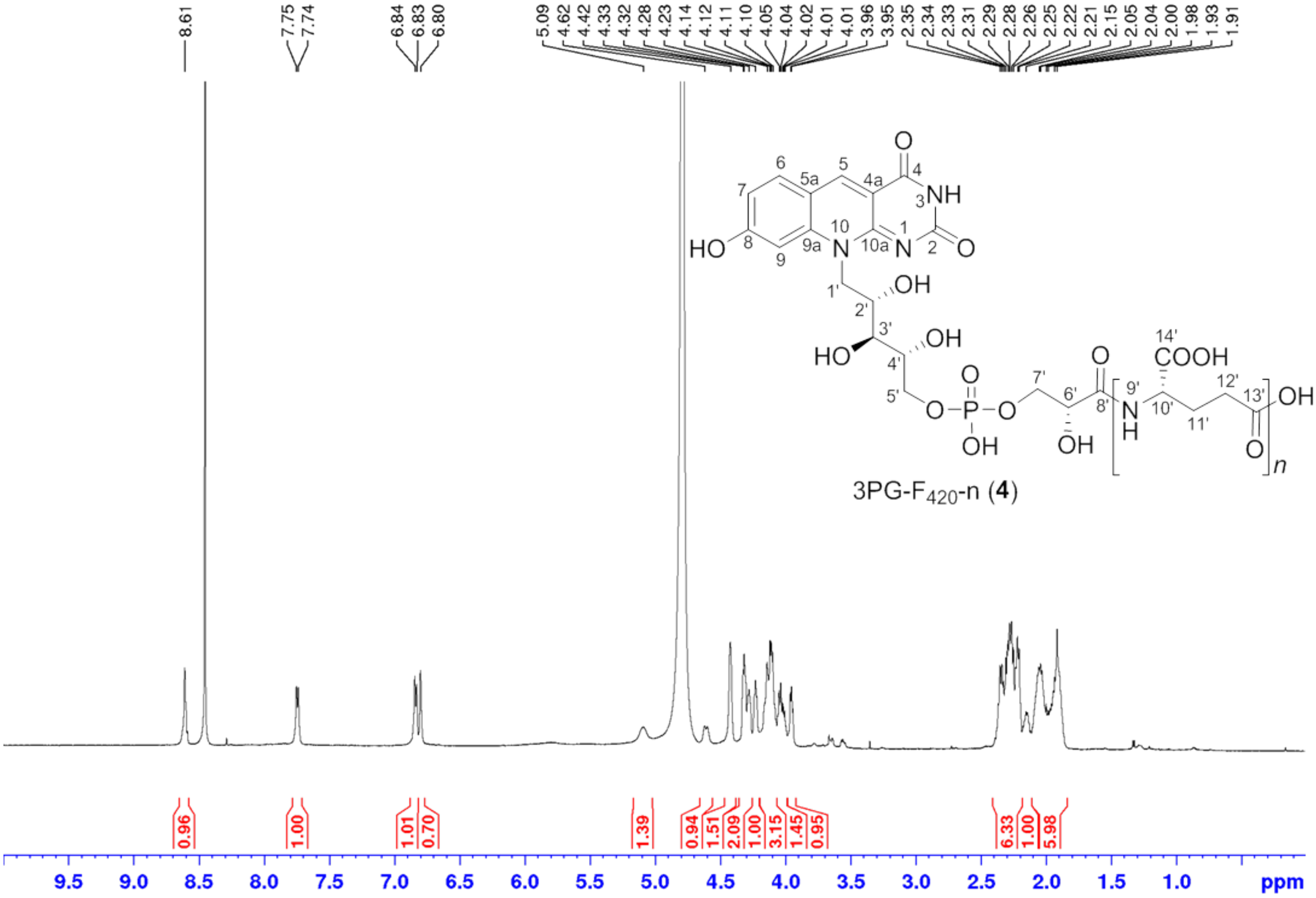

**Figure S43**. [1]H NMR spectrum of 3PG-$F_{420}$-n (**4**, 0.1% $ND_3$ in $D_2O$, 300 MHz, 300 K).

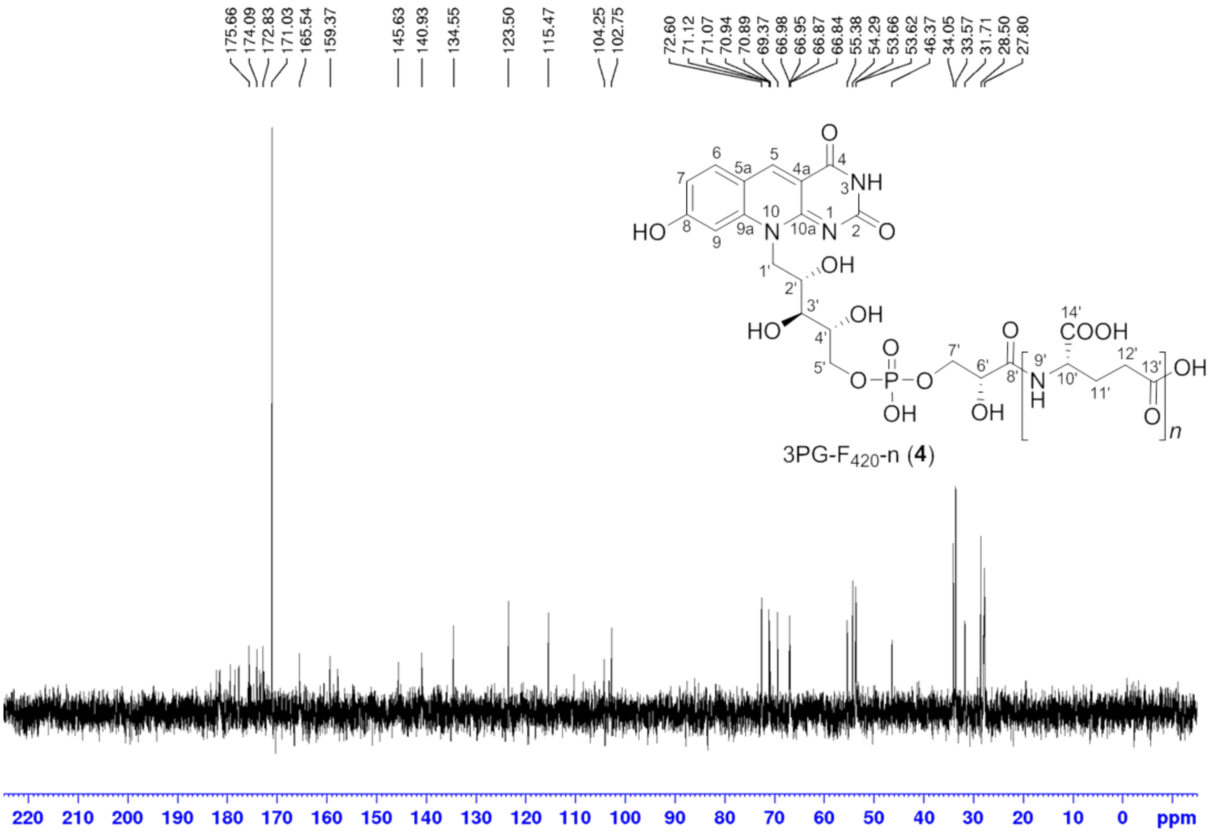

**Figure S44**. [13]C NMR spectrum of 3PG-$F_{420}$-n (**4**, 0.1% $ND_3$ in $D_2O$, 300 MHz, 300 K).

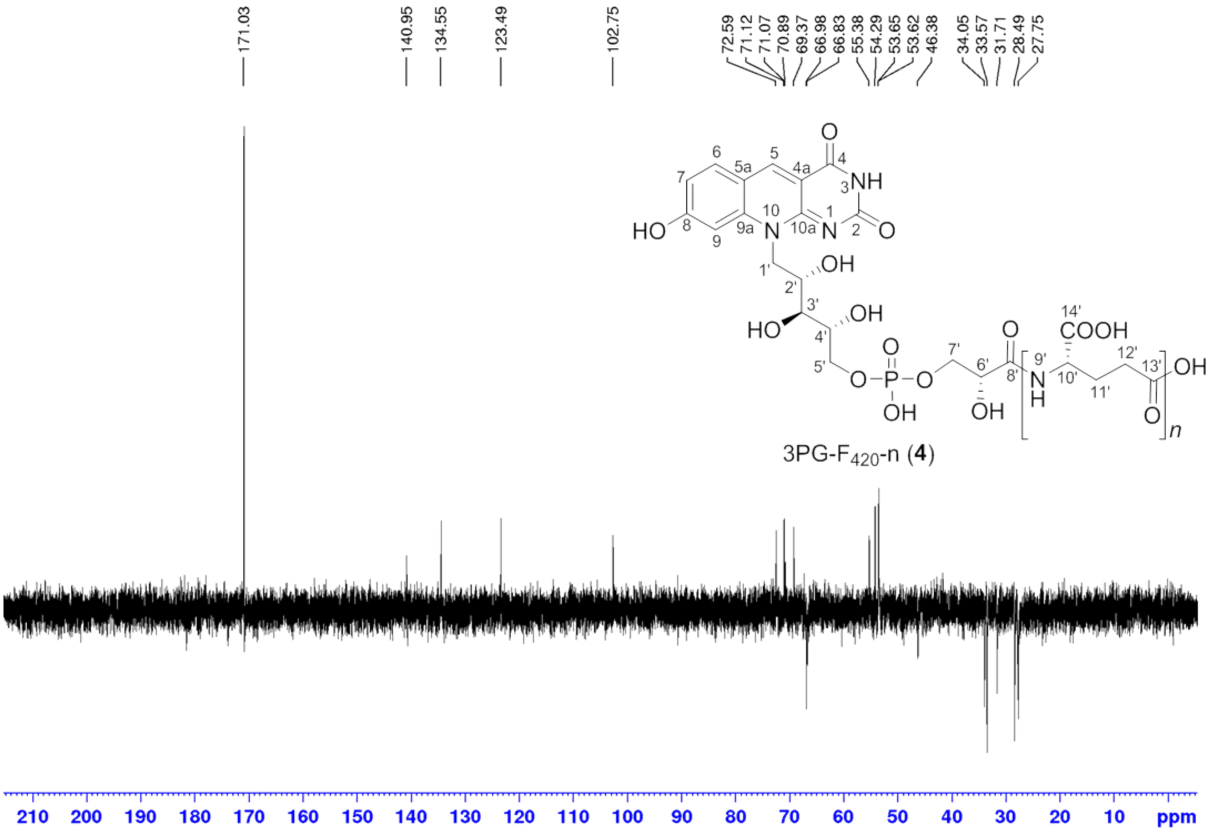

**Figure S45**. DEPT135 NMR spectrum of 3PG-$F_{420}$-n (**4**, 0.1% $ND_3$ in $D_2O$, 300 MHz, 300 K).

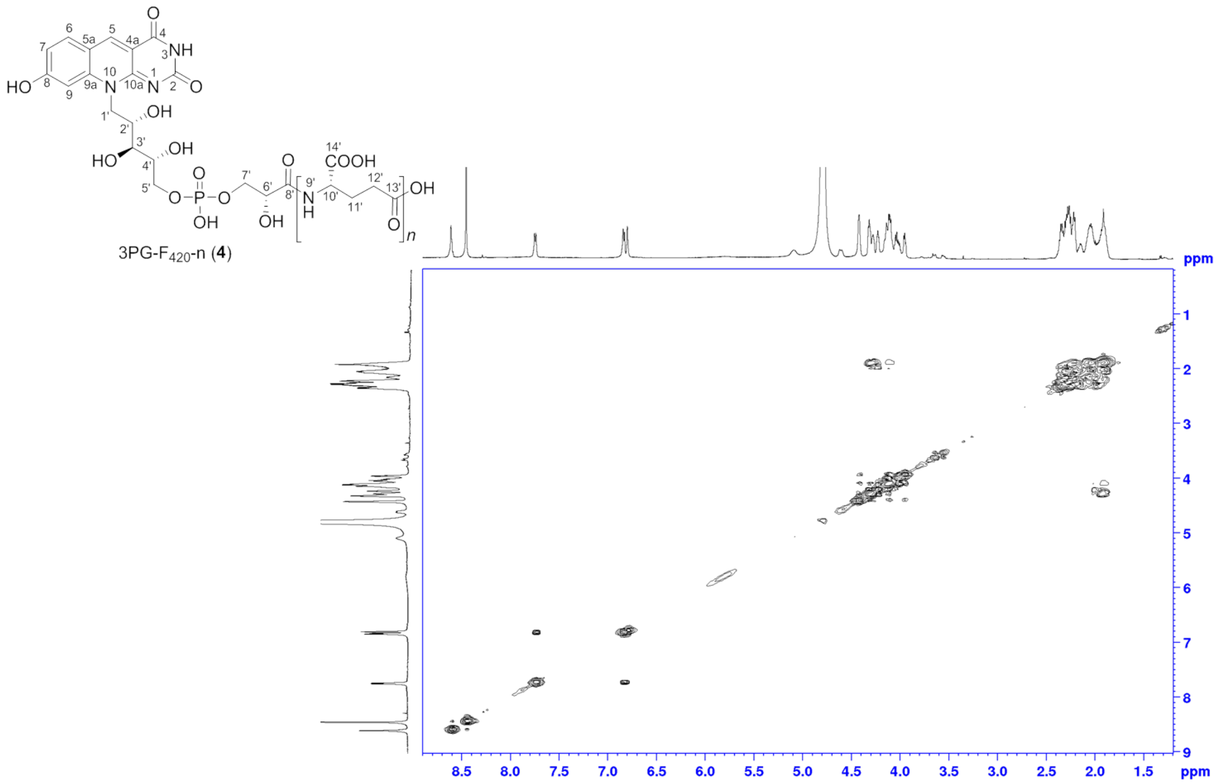

**Figure S46**. $^1H$-$^1H$ COSY NMR spectrum of 3PG-$F_{420}$-n (**4**, 0.1% $ND_3$ in $D_2O$, 300 MHz, 300 K).

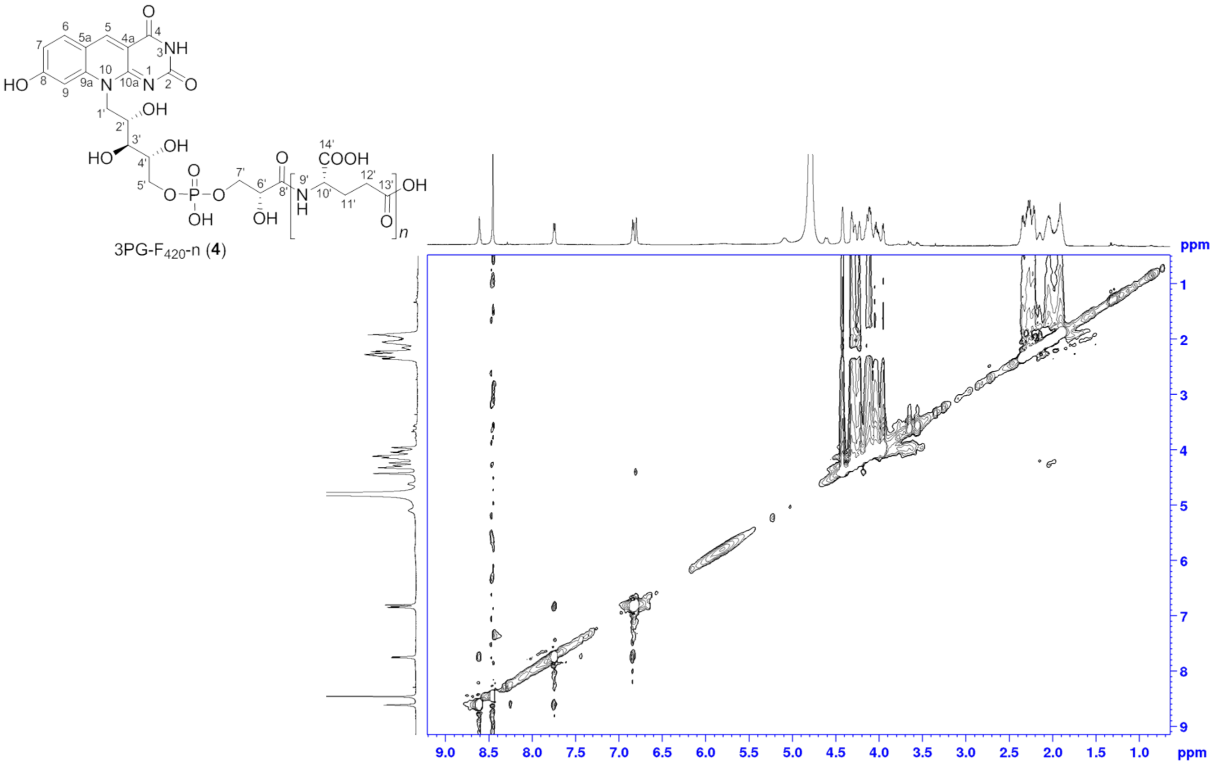

**Figure S47**. $^1$H-$^1$H NOESY NMR spectrum of 3PG-$F_{420}$-n (**4**, 0.1% $ND_3$ in $D_2O$, 300 MHz, 300 K).

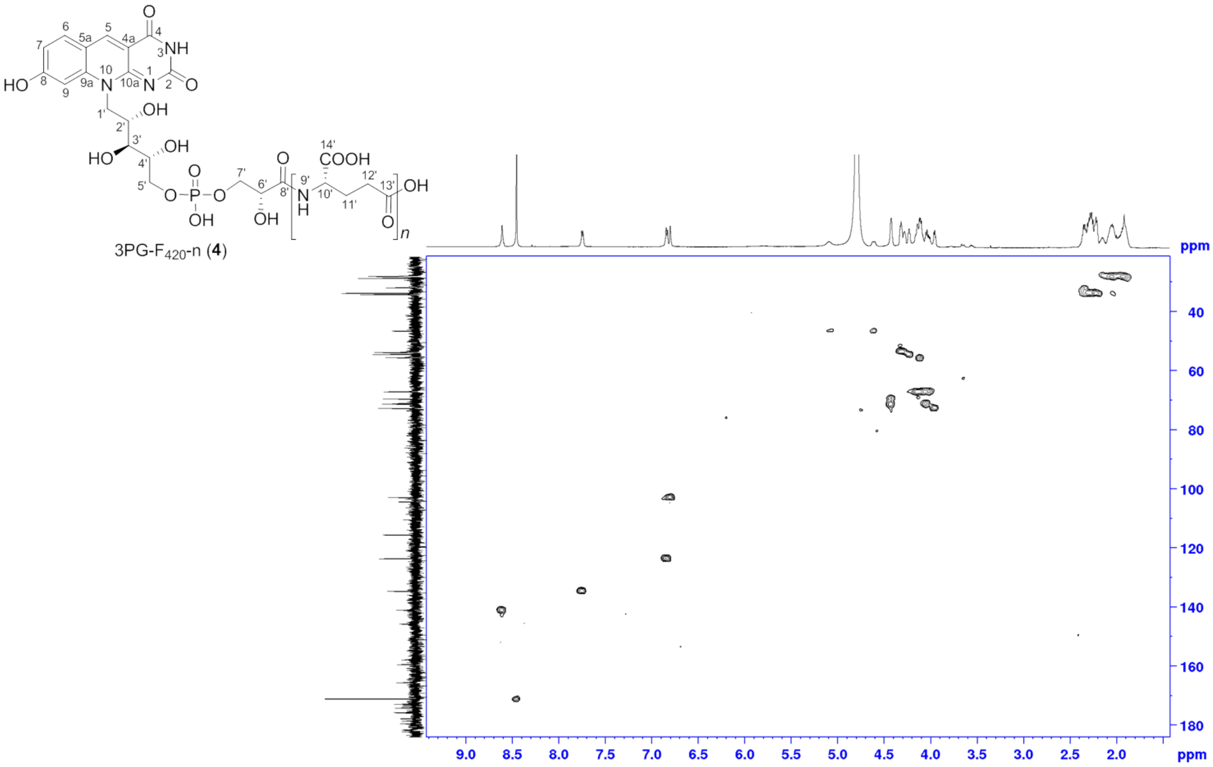

**Figure S48**. $^1$H-$^{13}$C HSQC NMR spectrum of 3PG-$F_{420}$-n (**4**, 0.1% $ND_3$ in $D_2O$, 300 MHz, 300 K).

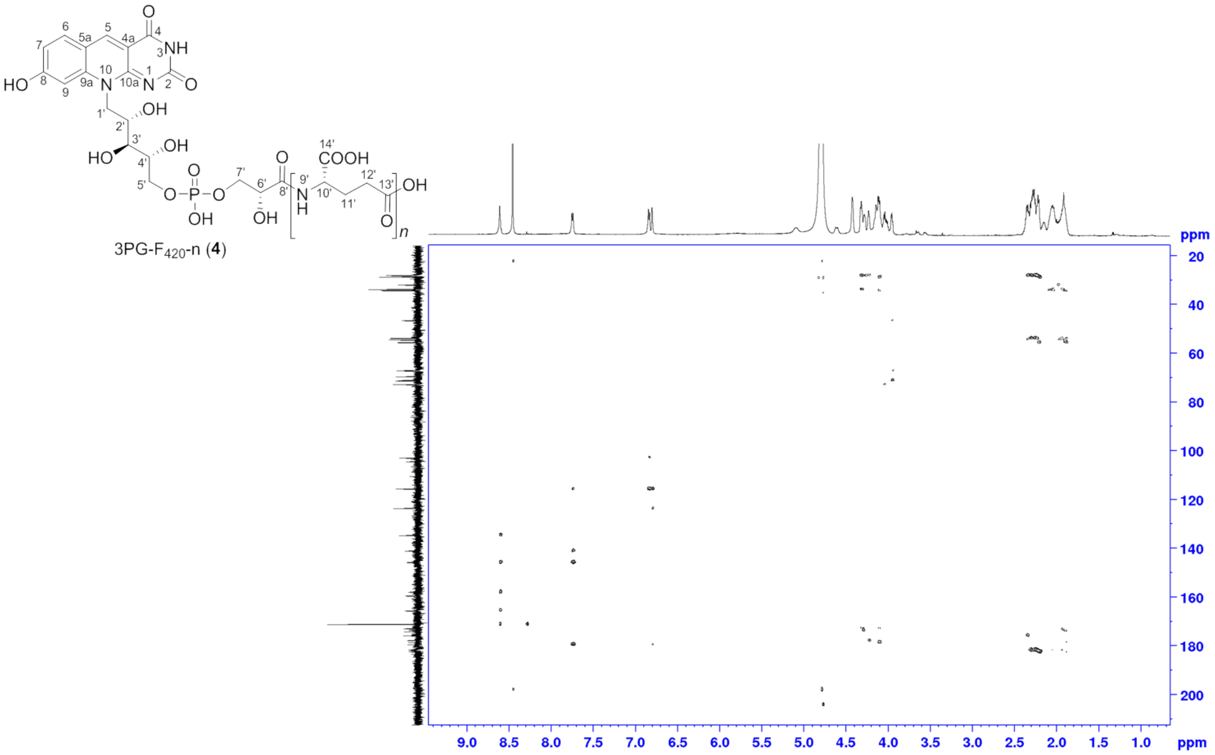

**Figure S49**. $^1H$-$^{13}C$ HMBC NMR spectrum of 3PG-$F_{420}$-n (**4**, 0.1% $ND_3$ in $D_2O$, 300 MHz, 300 K).

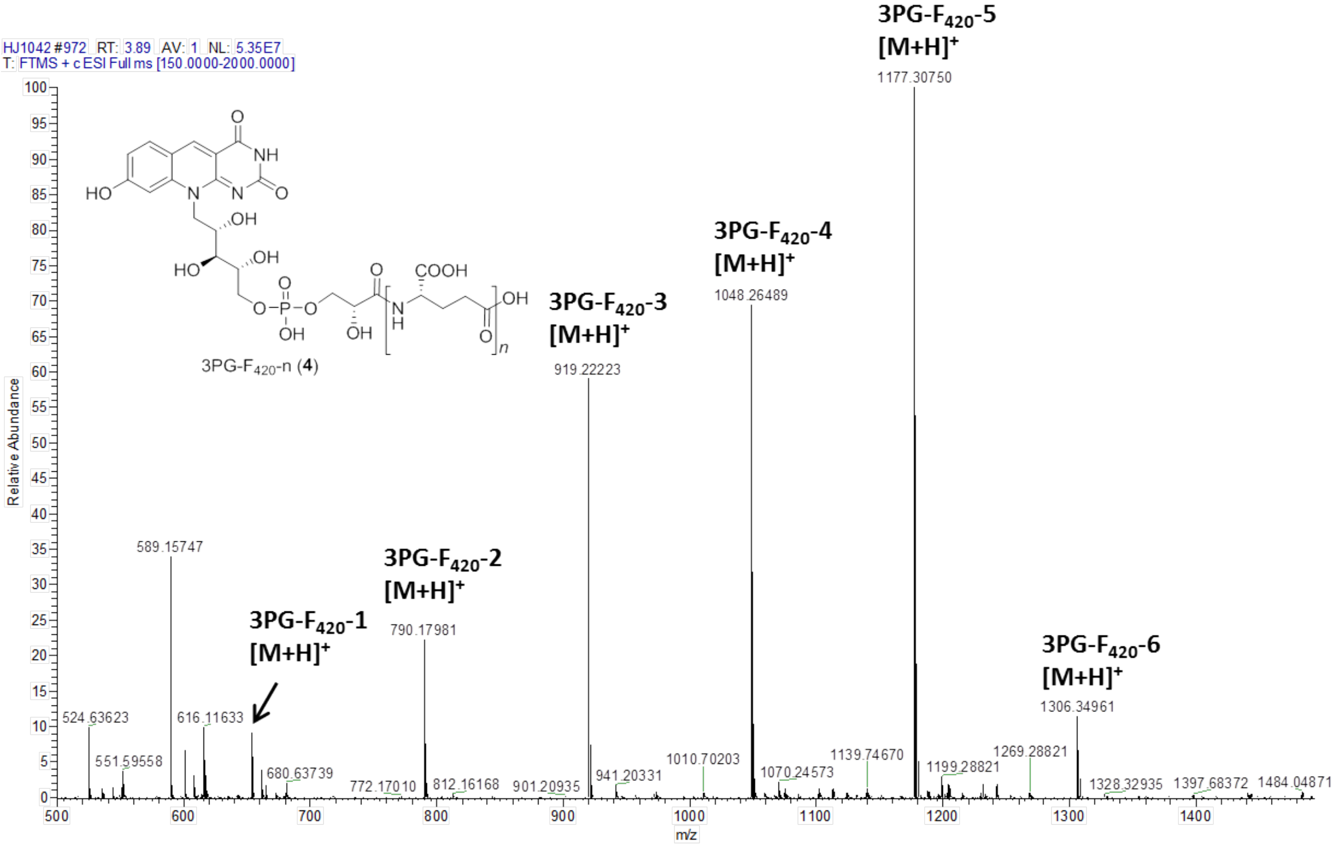

**Figure S50**. ESI-HRMS spectrum of 3PG-$F_{420}$-n (**4**).

##### *2.3.4 Determination of the absolute stereochemistry of 3PG-$F_{420}$*

In order to determine the stereochemistry of the 3-phosphoglyerate moiety, we performed acid hydrolysis of 3PG-$F_{420}$-0 (**5**) and analyzed the reaction products by UHPLC-MS as described in Experimental Procedures. Comparison with synthetic standards demonstrated that the retention time of the hydrolyzed product of 3PG-$F_{420}$-0 (**5**) was identical to the one of D-glyceric acid. Therefore, the stereochemistry of 3-phosphoglycerate moiety in 3PG-$F_{420}$-0 (**5**) side chain is assigned as the D-configuration (**Figure S51a-c**).

As a control, classical $F_{420}$-n was submitted to the same analysis. The hydrolyzed products as well as standards of D-lactic acid and L-lactic acid were analyzed under the same analytical condition. The identical retention time of hydrolysis products of classical $F_{420}$-n with L-lactic acid indicated the stereochemistry of the lactyl moiety as L-configuration as described in the literature(8) (**Figure S51d-f**).

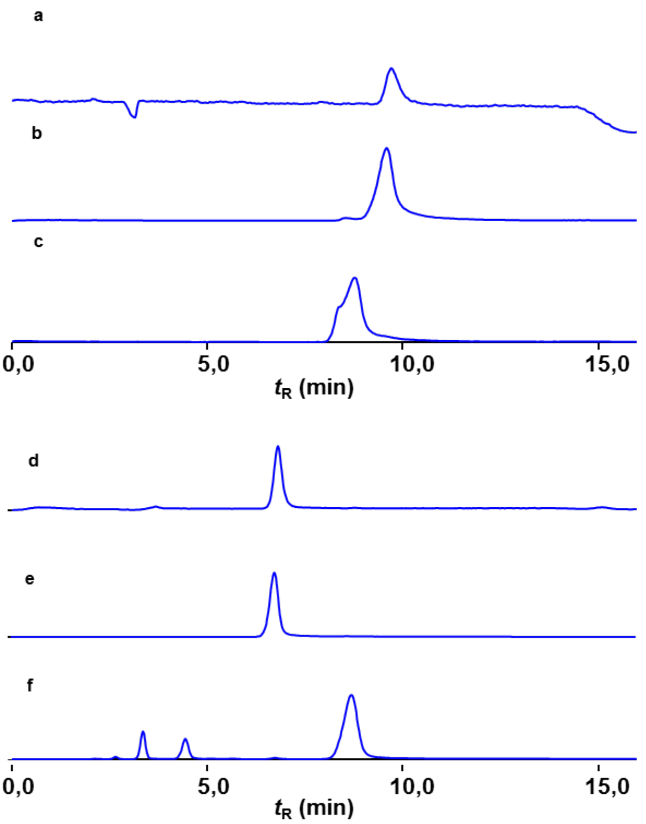

**Figure S51.** UPLC-MS chromatogram of acid-hydrolyzed product of 3PG-$F_{420}$-0 and $F_{420}$-n under SIM mode (–): a) SIM (–) with *m*/*z* 105.3 of acid hydrolyzed product of 3PG-$F_{420}$-0, $t_R$ = 9.707 min; b) SIM (–) with *m*/*z* 105.3 of D-glyceric acid, $t_R$ = 9.607 min; c) SIM (–) with *m*/*z* 105.3 of L-glyceric acid, $t_R$ = 8.747 min; d) SIM (–) with *m*/*z* 89.0 of acid-hydrolyzed product of $F_{420}$-n, $t_R$ = 6.775 min; e) SIM (–) with *m*/*z* 89.0 of L-lactic acid, $t_R$ = 6.733 min; f) SIM (–) with *m*/*z* 89.0 of D-lactic acid, $t_R$ = 8.681 min. AnAstec CHIROBIOTIC R (10 cm x 4.6 mm, 5 µm, Sigma) chiral column was used for chromatographic separation of enantiomers.

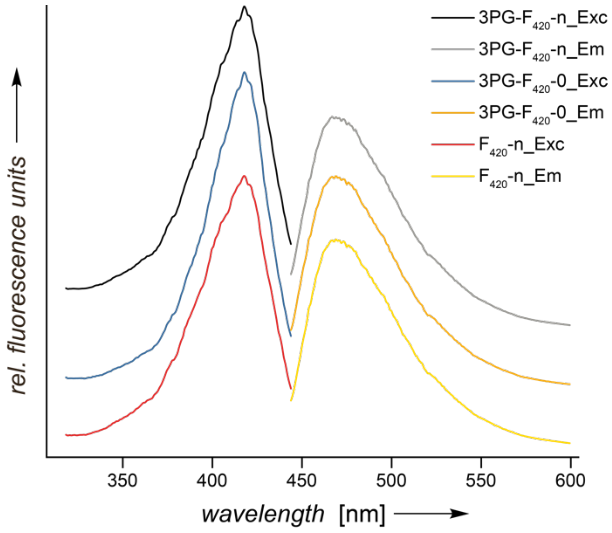

**Figure S52:** Fluorescence emission and excitation spectra of classical $F_{420}$-n, 3PG-$F_{420}$-n and 3PG-$F_{420}$-0 (50 mM sodium phosphate buffer, pH 7.5). The fluorescence spectra were distributed along the y-axis for better visibility. Em: Emission, Exc: Excitation.

#### 2.4 Combined CofC/D in-vitro enzyme assay

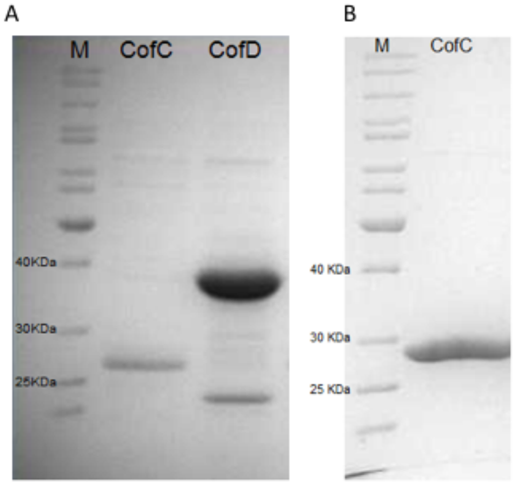

**Figure S53.** SDS-PAGE analysis of of recombinant proteins used for CofC/D assays. A) $His_6$-CofC from *P. rhizoxinica* (25.8 kDa) and $His_6$-CofD from *Methanocaldococcus jannaschii* (37,8 kDa) after metal affinity chromatography (Ni-NTA resin). B) $His_6$-CofC from *M. jannaschii* (27.65 kDa) after metal affinity chromatography (Ni-NTA resin). All proteins were produced in *E. coli* BL21(DE3). Page Ruler™ (Thermo Scientific) 10-200 KDa protein ladder (M) was used as a marker.

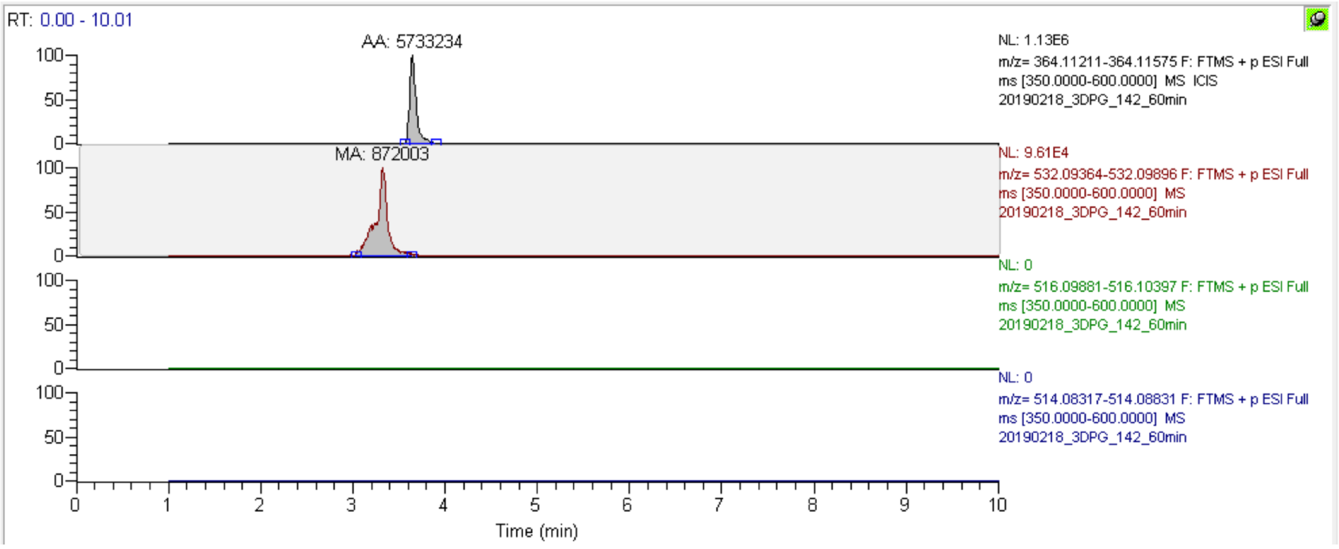

**Figure S54**. LC-MS analysis of combined CofC/D enzyme (CofC from *P. rhizoxinica*) assays with 3-phospho-D-glycerate showing XICs of reaction products after 60 min of incubation. Expected masses ($[M+H]^+$, 5 ppm mass tolerance): $F_O$: 364.11393, 3PG-$F_{420}$-0: 532.09630, $F_{420}$-0: $F_{420}$-0: 516.10139, D$F_{420}$-0: 514.08246. Areas under the curve are indicated on top of each peak. 3PG-$F_{420}$ is the only product formed.

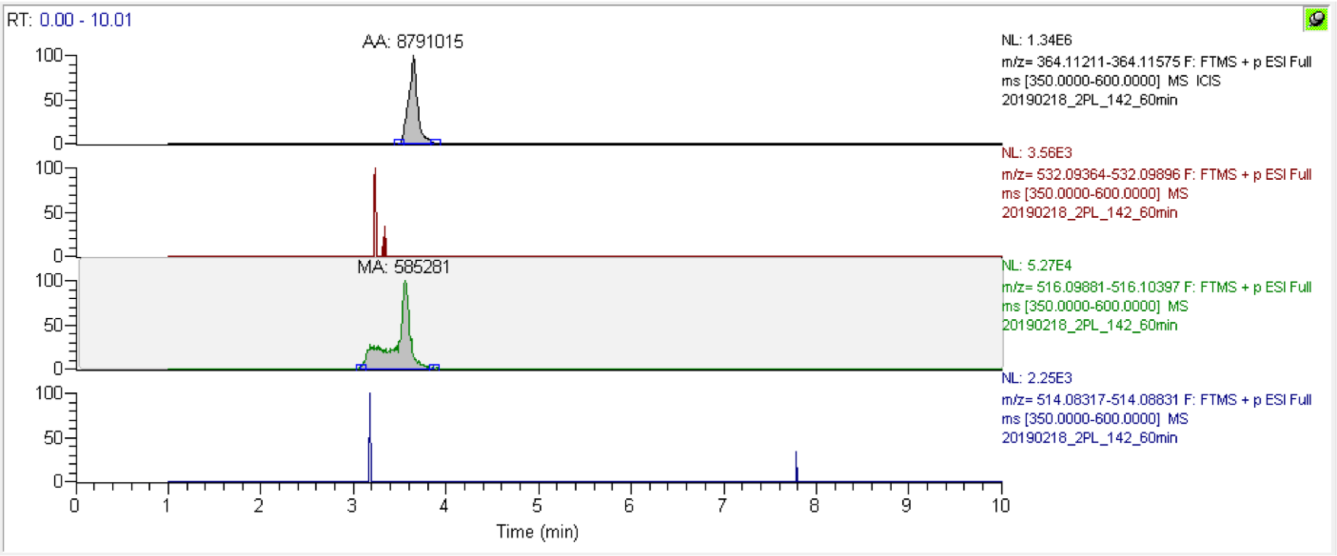

**Figure S55**. LC-MS analysis of combined CofC/D enzyme assays (CofC from *P. rhizoxinica*) with 2-phospho-L-lactate (2-PL) showing XICs of reaction products after 60 min of incubation. Expected masses ($[M+H]^+$, 5 ppm mass tolerance): $F_O$: 364.11393, 3PG-$F_{420}$-0: 532.09630, $F_{420}$-0: $F_{420}$-0: 516.10139, $DF_{420}$-0: 514.08246. Areas under the curve are indicated on top of each peak. $F_{420}$ is the only product formed. Traces corresponding to the mass of other species are below the noise level.

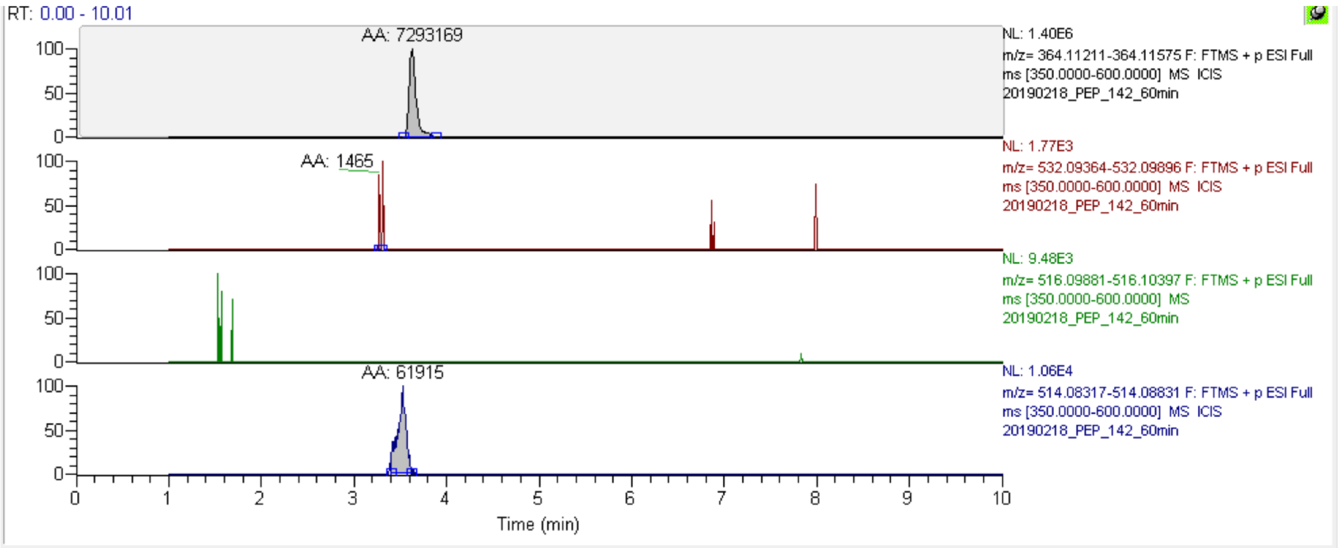

**Figure S56**. LC-MS analysis of combined CofC/D enzyme assays (CofC from *P. rhizoxinica*) with phosphoenolpyruvate (PEP) showing XICs of reaction products after 60 min of incubation. Expected masses ($[M+H]^+$, 5 ppm mass tolerance): $F_O$: 364.11393, 3PG-$F_{420}$-0: 532.09630, $F_{420}$-0: $F_{420}$-0: 516.10139, $DF_{420}$-0: 514.08246. Areas under the curve are indicated on top of each peak. $DF_{420}$ is the only product formed. Traces corresponding to the mass of other species are below the noise level.

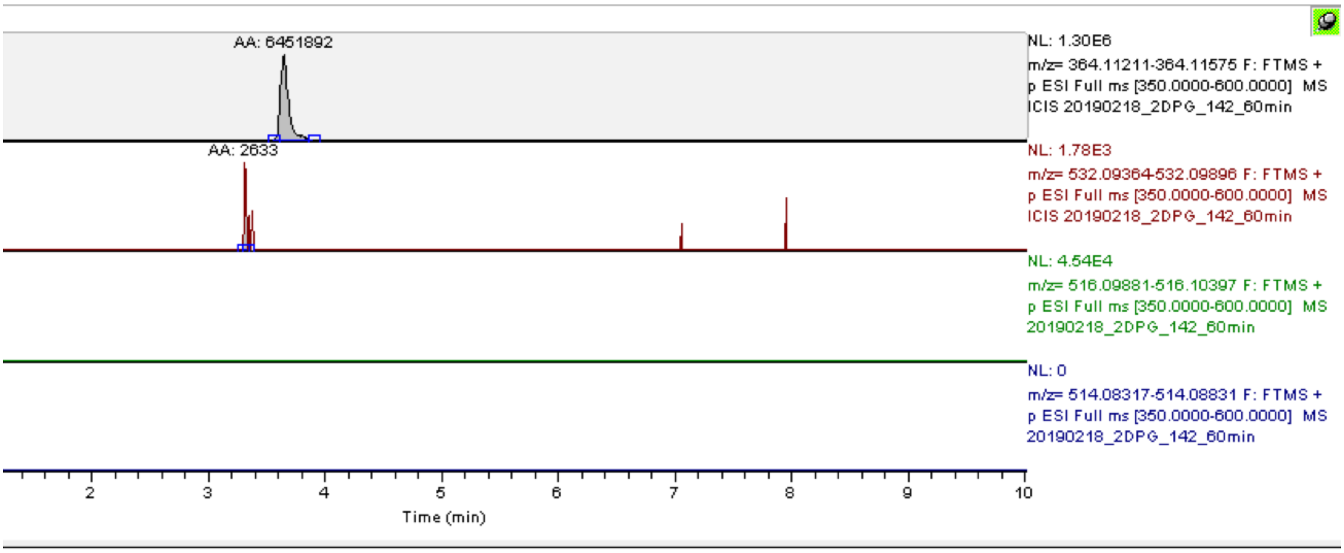

**Figure S57**. LC-MS analysis of combined CofC/D enzyme assays (CofC from *P. rhizoxinica*) with 2-phospho-D-glycerate (2-D-PG) showing XICs of reaction products after 60 min of incubation. Expected masses ($[M+H]^+$, 5 ppm mass tolerance): $F_O$: 364.11393, 3PG-$F_{420}$-0: 532.09630, $F_{420}$-0: $F_{420}$-0: 516.10139, $DF_{420}$-0: 514.08246. Areas under the curve are indicated on top of each peak. Traces corresponding to the mass of 2PG-$F_{420}$ species are below the noise level.

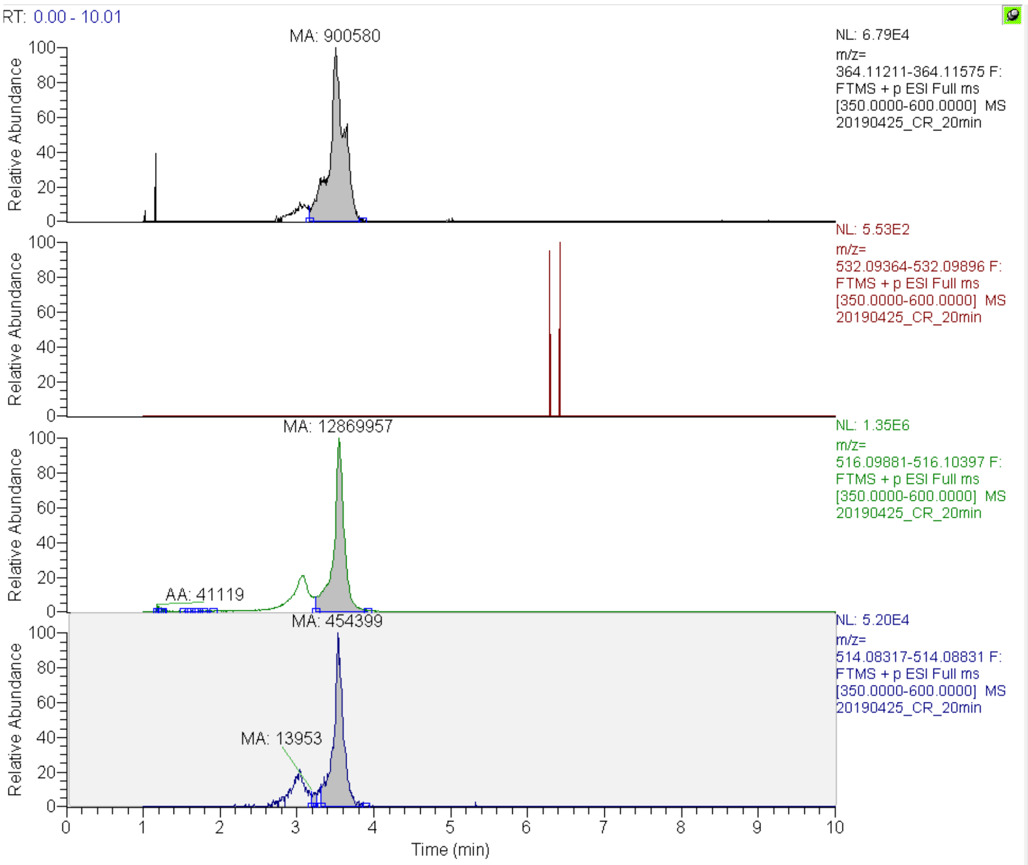

**Figure S58**. LC-MS analysis of combined CofC/D enzyme assays (CofC from *M. jannaschii*) carried out as substrate competition assay (2-PL, 3-PG, PEP) showing XICs of reaction products after 20 min of incubation. Expected masses ($[M+H]^+$, 5 ppm mass tolerance): $F_O$: 364.11393, 3PG-$F_{420}$-0: 532.09630, $F_{420}$-0: $F_{420}$-0: 516.10139, $DF_{420}$-0: 514.08246. Areas under the curve are indicated on top of each peak. Classical $F_{420}$-0 and $DF_{420\text{-}0}$ are formed. Traces corresponding to the mass of other species are below the noise level.

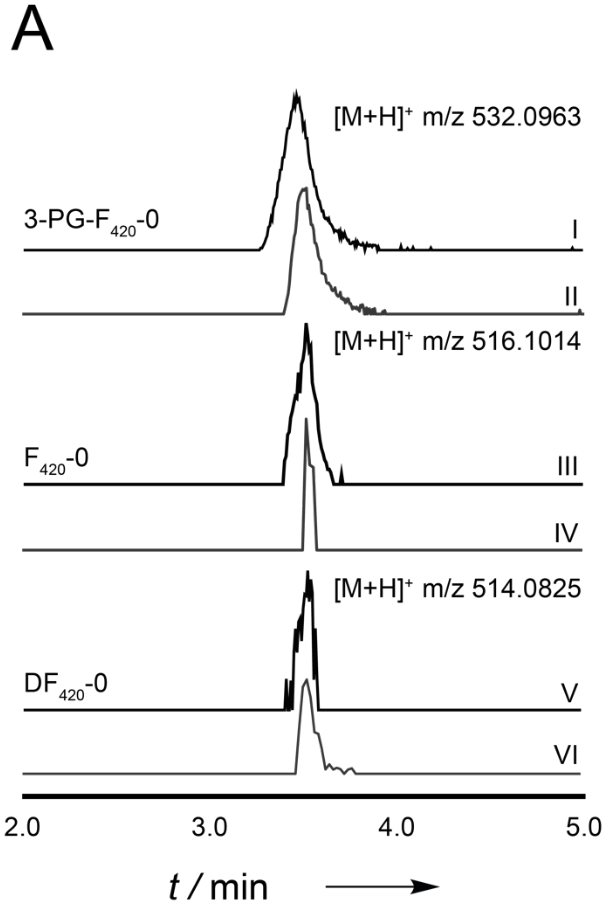

**Figure S59**. Combined CofC/D enzyme assay carried out as substrate competition assay (3-PG, 2-PL, PEP). LC-MS analyses (XICs) shows reaction products in comparison to corresponding in-vivo products: HF420-0 in vitro, III: F420-0 in vitro, and V: DF420-0 in vitro. II, IV, VI show corresponding in-vivo products A) Reaction with CofC of *P.rhizoxinica*. B) Reaction with CofC from *M. jannaschii*. Expected masses ($[M+H]^+$, 5 ppm mass tolerance): $F_O$: 364.11393, 3PG-$F_{420}$-0: 532.09630, $F_{420}$-0: $F_{420}$-0: 516.10139, $DF_{420}$-0: 514.08246. Intensities are not drawn to scale.

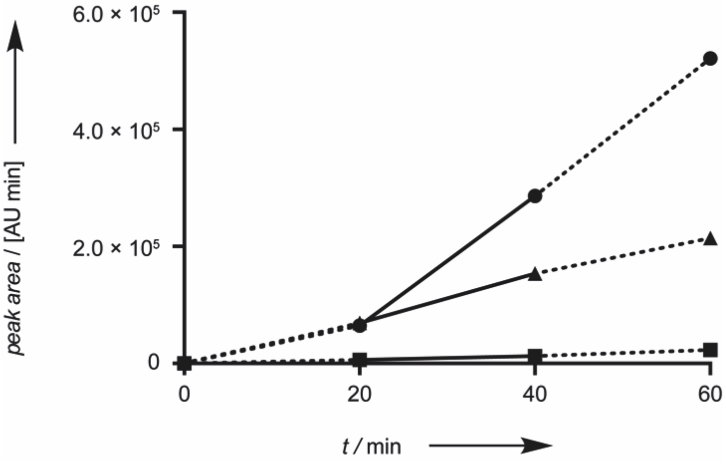

**Figure S60.** Time course of a combined CofC/D enzyme assay (CofC from *P. rhizoxinica*) with competing substrates (3-phospho-D-glycerate, 2-phospho-L-lactate, phosphoenolpyruvate). Corresponding product formation (circles: 3PG-$F_{420}$, triangles: $DF_{420}$ squares: $F_{420}$) was monitored by LC-MS and shown as area under the curve of extracted ion chromatograms (5 ppm mass deviation). Theoretical masses $[M+H]^+$: 3PG-$F_{420}$: m/z 532.09630, $F_{420}$-0: m/z 516.10139, and $DF_{420}$-0: m/z 514.08574. Solid lines indicate the near-linear range of the reaction that was used to determine the rate of product formation.

#### 2.5 Fno and malachite green reduction assays

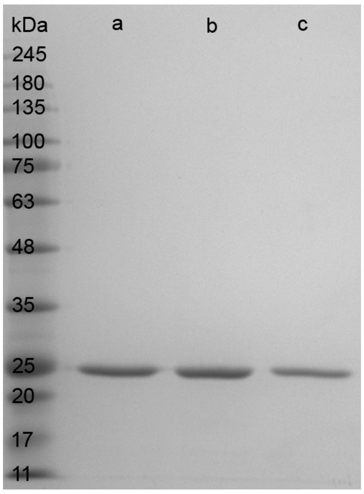

**Figure S61** SDS-PAGE gel of recombinant, purified His$_6$-Fno from *A. fulgidus* (24.1 kDa). Lanes a, b, and c show biological replicates used for kinetic studies. BlueEye Prestained Protein Marker (Jena Bioscience) was used as a marker.

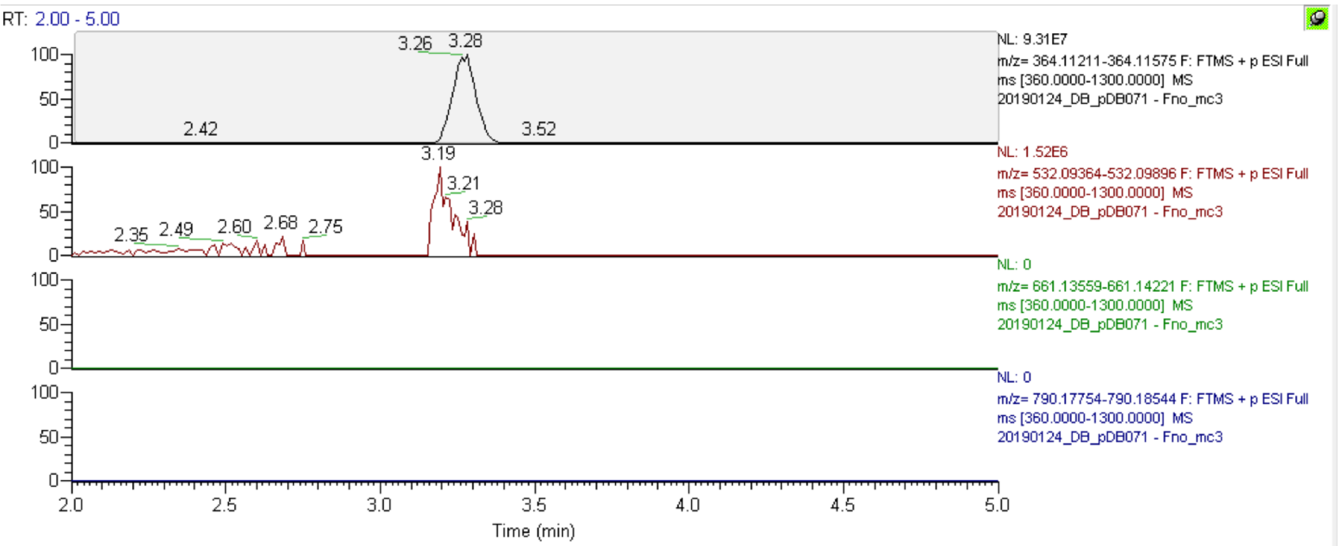

**Figure S62**. LC-MS analysis of extracts of *E. coli* BL21(DE3) / pDB071 (minimal BGC for 3PG-F$_{420}$-0 and *fno*) showing XICs of 3PG-F$_{420}$-n species with a varying number of (oligo)-γ-glutamate residues measured on a Kinetex XB-C18 column (Phenomenex). Expected masses ($[M+H]^+$, 5 ppm mass tolerance): F$_O$: 364.11393, 3PG-F$_{420}$-0: 532.09630, 3PG-F$_{420}$-1: 661.13890, 3PG-F$_{420}$-2: 790.18149. As expected only F$_O$ and 3PG-F$_{420}$-0 were produced.

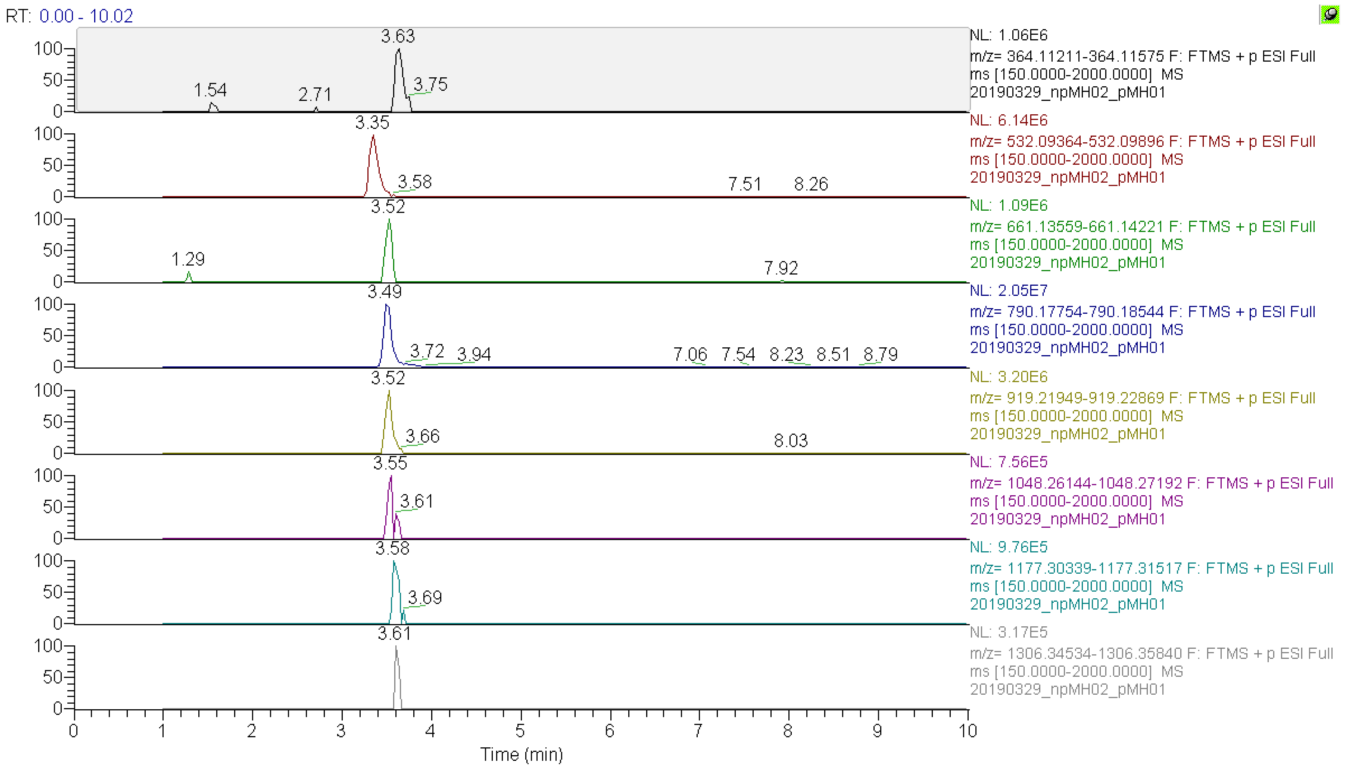

**Figure S63:** LC-MS analysis of extracts from *E. coli* BL21(DE3) coexpressing *cofE* from *M. jannaschii* (pMH01) and a minimal gene cluster (pMH02) producing 3PG-F420-0. Extracted ion chromatograms (5 ppm mass tolerance) were extracted using the following exact masses ($[M+H]^+$): $F_O$: 364.11393, 3PG-$F_{420}$-0: 532.09630, 3PG-$F_{420}$-1: 661.13890, 3PG-$F_{420}$-2: 790.18149, 3PG-$F_{420}$-3: 919.22409, 3PG-$F_{420}$-4: 1048.26668, 3PG-$F_{420}$-5: 1177.30928, 3PG-$F_{420}$-6: 1306.35187

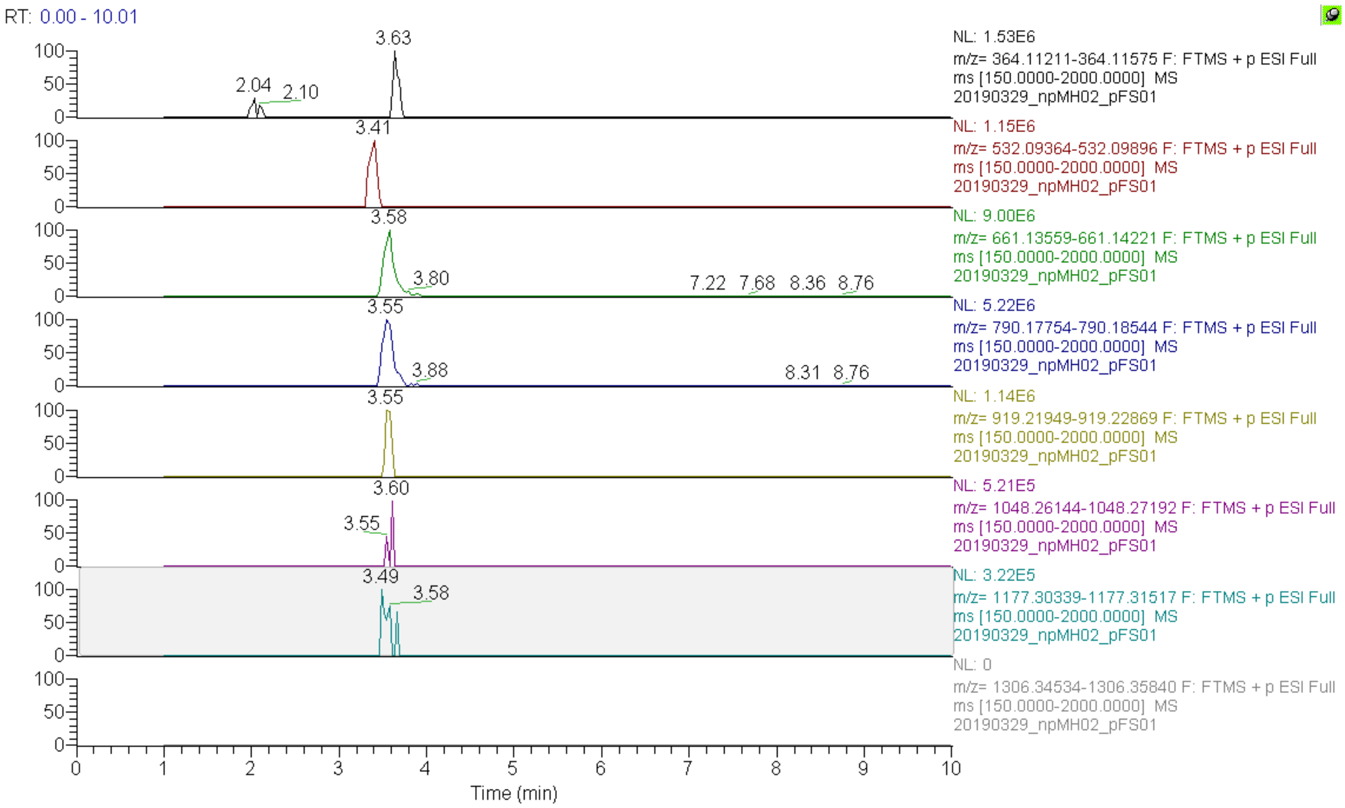

**Figure S64:** LC-MS analysis of extracts from *E. coli* BL21(DE3) coexpressing *cofE* from *P. rhizoxinica* (pFS01) and a minimal gene cluster (pMH02) producing 3PG-F420-0. $F_O$: 364.11393, 3PG-$F_{420}$-0: 532.09630, 3PG-$F_{420}$-1: 661.13890, 3PG-$F_{420}$-2: 790.18149, 3PG-$F_{420}$-3: 919.22409, 3PG-$F_{420}$-4: 1048.26668, 3PG-$F_{420}$-5: 1177.30928, 3PG-$F_{420}$-6: 1306.35187

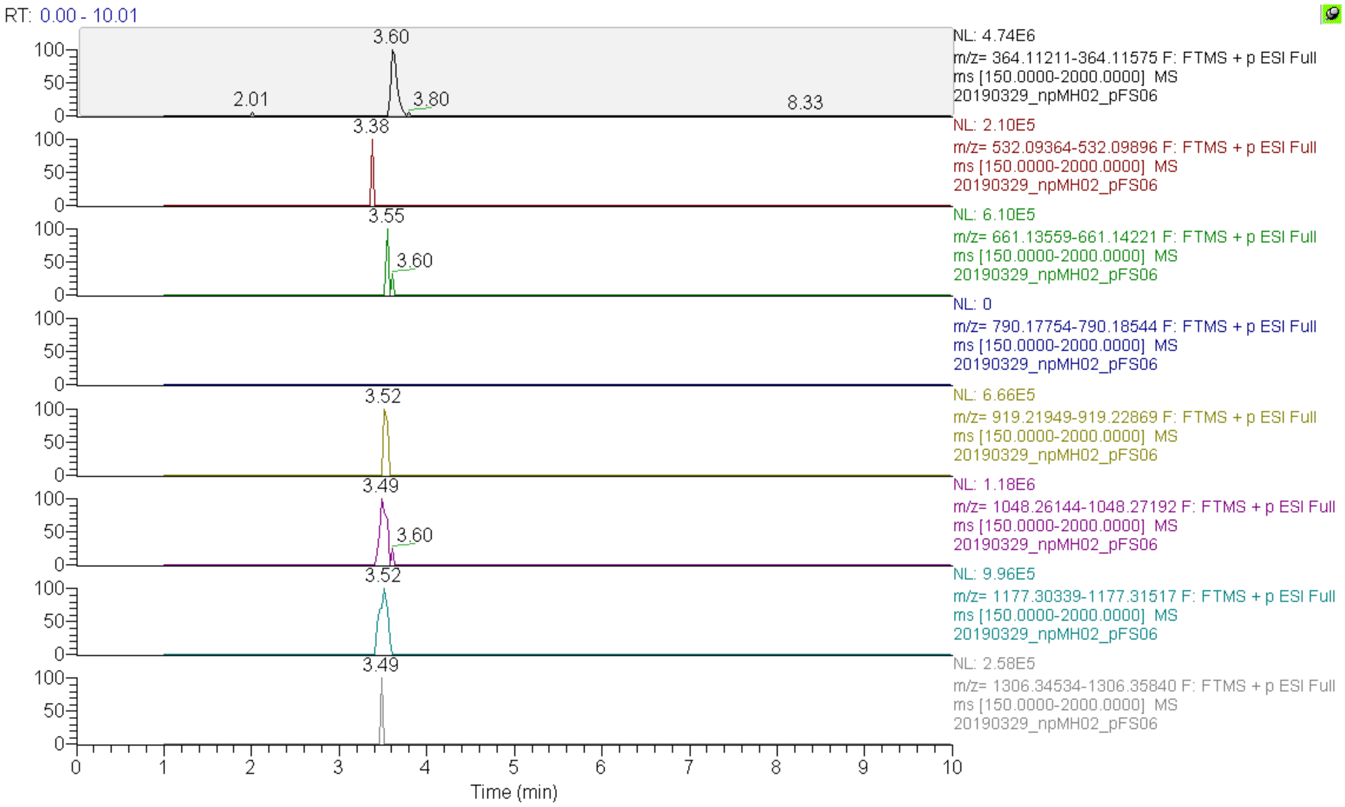

**Figure S65:** LC-MS analysis of extracts from *E. coli* BL21(DE3) coexpressing *fbiB* from *M. smegmatis* and a minimal gene cluster (pMH02) producing 3PG-F420-0. $F_O$: 364.11393, 3PG-$F_{420}$-0: 532.09630, 3PG-$F_{420}$-1: 661.13890, 3PG-$F_{420}$-2: 790.18149, 3PG-$F_{420}$-3: 919.22409, 3PG-$F_{420}$-4: 1048.26668, 3PG-$F_{420}$-5: 1177.30928, 3PG-$F_{420}$-6: 1306.35187

#### 2.6 Plasmids sequences (Fasta format)

>pDB045
GGGGAATTGTGAGCGGATAACAATTCCCCTCTAGACCCGGGGGTGATCCGATGCAACTACAATGCCATGCTGGAAGCAGC
GAGTACGTGACACATGTCGGCAAAGCAGCGATGCCAACTCGTTACGGAAATTTTGTTGCCCATGCATTTCGTTCGACAAG
CGATAACAATGAGTATCTCGCGCTGGTGATGGGTGACGTTTGCCAACGTGAATCAGTACTGACGCGACTGCACTCCGAGT
GCCTCACCGGTGATGTACTAGGATCACTGCGCTGCGATTGTGGTGAACAATTGGACGCCGCATTACGCCATATCGCATCT
GAAGGCGTTGGCGCGCTATTGTATTTGCGAGGCCATGAAGGACGCGGCATTGGCTTGTGTAACAAAATTCTTGCATACGG
ACTCCAAGAACAAGGACTTGACACCGTCGATGCCAATCGCGACCTGGGTTTGCCGGATGACGCGCGCGAGTATGACTGCG
CGGCGAGCATCGTGCGCCAACTCGGCATCCTGTCGGTACGGCTGATGAGCAACAATCCGGATAAGTTTGAAGCGCTGCAG
CGCCATGGCATTCCAGTATGTGAGCGCGTTGACCTCGCAATTGCATTACGCGAGGAAAATGAGCGTTATATCTGGACTAA
GCGCAATCGTTTCGGGCATTATTTTGACGAGAGTGAGCTTCAGTGCCCGTTGTCATTGTAGAGGCCTATATGAGATGCAA
TGGTATGTCTCCTGTTTCTGTCGCCCAGCGCAATACGGGCATCTGGGCGGTCGTGCCGTTGAAAGCGCCTGAGTGTGCCA
AAACCCGGTTGTCTGGCGTTTTAAGCCATGCTGCGCGCCAAGCGTTGTTTTTTTCCATGGCCAGTCATGTCATTGGTACG
TTACGCGCATCGCCTCGGATTGCTTCCCTATTGGTGGTGACGCCGTCGGAAAGTACGGCGGAAATGGCCCGGGCAGCCGG
TGCTGAGATCTTGTGGGGACCACCGGACGAAGGCATGGCGAATGCCTGTTCGCGAGCGATGGCTCATATTGCGGCAGCAG
GCGGAGAGCGTGTGATGTTTGTCCCCGGTGATTTGCCCCTGTTGGATGGGGCGGCCATCGACATGTTGAGCCGTGCACCG
GTCGATGCGATTGGCATGGCGCCAAACCGGGACGGTCATGGCACGAATGGGCTGATTTGCCGACCTGGCGCTATTCCGTT
GTTTTTCAGCGGGCCAAGCTTTTCCGCTCATCAAAACGCCGCTCGGTGCGCCGGAATTGATGTTTGGATTGTCCGTTCAA
GAGAGTGGGCGTTGGATGTAGACTTGCCTGCCGACCTTGAGGAATTCGAGTCTTCCATCAAAGACGCCAAGAGGAGGGTG
CTATGCCAAATCTAGTAAGCAATCCTATTTACAAAATGATCCAAGAGGTCGAACGTTCCGCTAAGGGCAGTCAGCCAAGG
CTAGATGACAAGGCTTTGGCGTATGAGCTTGAAGCGGTGCAAGATTGGCGTGAGTTGGCGGGTCTGGCATCGGCATGGCG
CGATATCGGTTGGGGCAACGTCATCACCTATTCGCGCAAGGTTTTTATTCCGCTCACCCATCTTTGTCGAGATGTTTGCC
ACTACTGCACCTTTGCCAAAGCGCCACGGGTGATTGGCCAAGCATTTCTTACCGTGGACCAAGCGCTGGACATTGCGCGG
GCGGGTGCTGCAGTTGGCTGTCGTGAGGCTTTGTTTACGCTCGGAGATCGGCCCGAAGCGCGTTATGCCGCTGCACGCGA
TGCGCTGCAGCAACTGGGTCATGCTACGACAGCCGACTATGTCGCTGAGGTCGCGCAACGCGTACGAGCTGAAACCGGGC
TGCTGCCCCACTTTAATATGGGCGTGCTAAGCGCGGCAGAATACCAGATGCTGCGGCCCCATGCCCCTTCCTTCGGGTTG
ATGCTAGAGACGGCTTCTGAACGTCTTTCCGAGCGTGGTGGGCCGCATTACGGCTCTCCCGATAAGCACCCGGCAGCGCG
GCTTGAAACGCTTCGCCTTGCCGGCGAAGCGCATATACCGATCACCTCAGGCATATTGATTGGTATCGGGGAGACACGCC
GCGAGCGGCTTGAATCGCTCTTTGCGCTGCGGGATTTGCATGATCGCTATGGTCATGTGCAGGAGGTCATTATTCAGAAC
TTTCGCGCGAAACCCGGCACCAAAATGGTTAACGCGCCTGAGCCGTCAATGGACGAATTGTGTTGGACAACAGCGGTCGC
CCGGTTGGTCCTGGGTTCCGCAATGAGCATCCAGGTGCCGCCTAATCTTTTCGACGGTGACTTGACCGATTTGATTCGCG
CGGGAATCAATGATTGGGGCGGCGTATCGCCGGTCACGCCTGATCACGTTAATCCAGAAGCGCCGTGGCCGCACCTTGAC
CGCTTAAGTGAGGATACAGACAGGGGCGGGAAGACGTTGACCGAGCGGATAACCGTCTATCCTGCCTATATCGAGGAACG
TGATCGTTGGATCGACCCGGGCTTGCATGCTGATGTGCTACGCCATTCAGATTGCAGTGGATTGGCAACAAGCGATATCT
GGAAAGCCGGCTCCACGACGATAGGGGATATCGCTTGCCTGGCAGTCACTCCCACGGCTTCCCAGGTTGATCGTGTTGGT
TCGCAGACGATCGATATCGTCGAGAAATGCGTGGCGGGTGAGCGCCTCGAAGAAGCCGAGCTTGTACATTTATTTAACGC
ACGCGGCCATGATTTTACGCATGTGACGCACACGGCGGACCGTTTGCGCAGGCAGACAAAGGGCGACACGATCACTTACG
TTGTCAATCGGAACATCAACTACACCAACGTCTGCCAATACCATTGCAATTTTTGTGCTTTTTCACGAGGACCGATACAA
GAAGACTTGCGCGAAAAGCCCTATATCGTTGACCTCAACGAAATCCGTCGCCGCGTCAAAGAAGCGTGGGATCGAGGGGC
GACGGAGGTCTGCTTGCAAGGGGGCATTCATCCCGACTATACGGGACGTACTTATCTTGACATCTGCGAAGCGGCTAAAA
CAGAATGTCCAGACATGCACATTCACGCCTTTTCGCCCCTTGAGGTGCTGCATGGTGCCACGACCCTGGGGACGTCAATC
AGCGCTTTTTTGTCTGATTTGAAAAGCGCTGGATTAAGTACGTTGCCTGGGACCGCAGCCGAGATCCTGGACGACGAAGT
GCGGCGACAAATCTGTCCGGATAAGTTGAATACGCAAGAATGGTTGACCGTCGTGCGATCGGCACATGAGTTGGGATTGC
GGACCACTTGCACGATCATGTTCGGTCATGTCGAATCGTATAAGCATTGGGCGCGACATATCCTCAGGCTTGCCGAGCTG
CAACGTGATACAGGCGGGCTGACTGAGTTTGTTCCATTGCCTTTTGTCCATGAGGAAGCGCCATTATTTAAAAAGAAAGG
TGCGCGTCAGGGGCCGACTTTACGAGAAGCCATTTTGATGCATGCCGTCGGGCGTCTGGCGTTTCATGGCTTGATTGACA
ATATTCAAACGTCTTGGGTGAAAATGGGTGAGCACGGCGCACAGCTTTGCTTGCAGGCGGGGGCTAACGATTTGGGCGGT
ACCCTAATGAATGAATCGATTAGCCGCGCAGCCGGTGCAGCCCACGGACAGGAAATGCCGCCTGCCGCCATGGAAGCCCT
GGCCGCCAGACTCGGGCGCAATGCGATGCAGCGTACCCCGCTTTATCGTAACGTCAGTCCTGAGCGGCAGCAAGCTTCGC
ATGCTGCTCCTGCGCTGGTGCCTGTGCGATTCACTAAGGCCGGACGCTTGGTGCGTGGGCACGGCCATCAGGTCTTGAGT
GAAGCGGAACAATGATTGGACCTGCATAGGTGGATCATAGGTGGATTCATGGTGTGGCTTGCATGTTGGCAGCCTCAAAT
CGATGTTGTGATGGCTCTCACGCAGCGCTTCCCATCATGGATTCACCAAGGTGCTTAATTACCCTATTAAATTAGACAGA
CGCTATTCGGTTTGACACGATCGGAAGGGTGATCGGTTTGGGTCAGAGAAATGCTTGATCTTTACGCAGGAAGATAATGG
CAAAGTATGTTGCGTTGTGCGGCGGGGTGGGCGGCGCTAAGCTGGCCTATGGTCTAGCGCAGGTGCTTTCTGCCGACGAG
TTGACGATCGTTGTGAACACGGGCGACGATTTTGAGCATCTTGGCTTGCTCATCTGTCCAGACCTGGACACGGTTGTCTA
TACGCTGGCCGATGTGGCAGATGCAAAAAAAGGATGGGGGCGGGCGAACGAGAGCTGGTCTTCGCAAGAAGCGCTCGCTC
GTTTGGGCGGACCGGTTTGGTTCCAGCTGGGTGACAAGGACCTTGCGCTGCATCTTTATCGGCGCAGCTTGCTTGACGGT
GGTGCCAGCTTGTGTGATGTAACCGAGACCATTGCGAAAGCGTTAGGTGTCAAGCATTCAATCGTCCCAATGTCTGATGA
TCCGGTCAGAACCATCGTTGAGACAGACGAAGGCGATTTGCCTTTCCAGACGTATTTCGTTAAACGGCGATGTGAGCCAC
GCGTGTCGGGGTTCCGCTTTGAAGGGGCTTCGCTAGCACGTTTATCGTCGCCATTTGAAGCAGCGTTGAGTACGCCTGAC

CTGGCTGGCGTAATCTTATGCCCTTCGAATCCTTTTGTCAGCATTGGACCGATTCTGGCATTACCGGGCGTGCGTGACAG
GCTGCATGCGTGTAATGTGCCTGTTTTAGCTGTTGCGCCGCTTGTCGGTGGTGAAGCAGTCAAAGGCCCGTTGACTAAGA
TGATGCATGAACTGGGCATGAGTGTTTCGGTTGGCGAGATTGCTTCGCTTTATGCTGATTTCCTTGATCTGCTGGTGATT
GATCCGTTGGATGAATGCGATCATGATCTTTTGCGTAAGGATCGTGTGGCAGTCAATAAAGTCAAGACGCTCATGACAAC
ACCTGATGAGCGTATCGCGCTTGCGCGACACGTGCTGGCTTGCGTTGAACATCACCGAAACAATCGACAGACAGCTCACG
TTGTGCAGCACAGCACCGCATAAATCACATTGACTATAATTTTTATTAATTAGGTTGATGGCTATACAGAAATAAAAAA
TGATCAATGGCATGCGGTACGGCAAACGCTTGCTGAACCACTCACGAAGAATGCACTATCCACTACGCATGCGCCGCCTT
TGGGCTTTTGTTTCTCCCTTGTCATCTTCGAATTGGCGCGGCGACTGGCAGACAATCGCTGCGCTATTTCCCTATTTGAT
GGTTTATAAAGGGCGCGTCGCCTTCGCACTTGCCTGTCTGGTTGCCGCAAAACTCGCTAATCTAGGGGTTCCGGTGGTGA
TGAAAGCGATTGTCGACCAATTATCAGAAATTGACCGGTTGGCAACGCAGGAATCGTCATTTTCCTTGTTCCACGGCGTG
AGTTTTCTTATTGTCGCTTATGCGGTCGTGCGTTTATCAAGCTCGCTTTTTGCTGAGCTGCGTGAAATTATCTTTGGAAA
AGTGGCTTACAACGCTGCGCGACAAGTGGCACTCGCAGTATTCCGGCATCTGCACACGTTGTCGTTGTGTTTTCACCTTG
AGCGGCAAACAGGTGGCCTATCGCGCGATATTGAGCGAGGCACGCGTGGCGTCAAGACGCTTGTTTCATATTCGCTCTAT
AATATCCTGCCAATATGTGTTGAAGTCATATTGGTACTCATTTTTTTTGCTATTCGTTACGACATCTATTATACGGTCGT
CACGCTATCCGTGCTTGCGGCTTACATCACGTTTACCGTGATGGTGACAGAATGGCGTACGCGTTTAAGGCAGGAGATGA
ATAAGCTTGATTCACGCTCCAATACGCTAATGGTTGATTCGCTTATTAATTACGAGACAGTCAAGAACTTTGGTAACGAG
CAACACGAAGTACAGCGCTATGATGAAAGCATGATGCTTTATCATGATGCGGCAGTTCGATCGCAGAAATCGCTTTCGTT
CATGAATCTTGGCCAGCAGTCGATTATTGCAATCTGCATGATTGCCGTTTTATGGAGGGCGACGCAGCAGGTTGTGGATA
AGCAATTGACATTGGGCGACTTTGTGTTAATCAATACATTTATGTTGCAGATATATATTCCGCTGAGTTTTTTGGGGAAT
ATGTATCGAACTTTGAAGCAAAGTCTGACCGATATGGATCAGATGTTTTCTTTGTTAAGGCTCAGACGTGAAGTTGACGA
TATTCAAGGTTCATCTCCGCTTGCTGTAAGAAGCGCTGAAGTACGCTTTGAGCATGTCAGTTTTTCATATGAACCGCAAC
GGCAAATTTTGCGGGATGTCACCTTTACGATTGCAGCGGGTACGACGACCGCAATTGTCGGACATAGCGGCTCAGGTAAG
TCGACACTCGCACGGCTGTTGCTTCGCTTTTATGATGTAGAGCATGGCGCTGGTCGTATTTTGATTGATGGCCAAGACAT
TCGGGCTGTGACGCAAGACTCGCTACGCGCGGCCGTTGGCATTGTGCCGCAAGATACGGTGCTTTTCCGTGACACGATCT
ATTACAATATTGCGTACGGACGGTTGTCTGCGTCGCCAGAAGAAGTGATAGCGGCTGCGCGCGCCGCTCATATTCATGCT
TTTATTGAAAGCTTGCCCGCTGGCTATTCCACTATAGTGGGTGAGCGTGGTTTGAAATTATCTGGCGGTGAAAAACAGCG
CATCGCAATTGCCCGTACCTTGCTAAAAAAGCCGCCTATTCTCATTTTTGATGAAGCGACTTCAGCACTCGATTCACGTG
CCGAGCGTGCGATTCAGCGCGAGTTGAAACAGCTTGCGCGTCATCGTACAACACTAATCATTGCGCATCGTCTGTCGACC
ATTACGCATGCGCAACAGATTCTTGTTATGGATCAAGGTCGTATCGTTGAGTGTGGCACCCACAGAACATTGCTGAATGC
AGGTGGCCTATATGCGCAAATGTGGGCATTGCAACACAAGCAGCCTGAATAAGTTTAAACCAGCAAGAAGCATGACTGTA
TCCGCCATCGGTGGCATTCCCTTAATACAAACAGGGGATGATCTTGGCCAGATTATTAAGGAGGCGATCCACAAAAACGG
TATCGCTCTAGAAAATGGCGACGTGTTAGTGCTTGCGCAAAAGATCGTTTCAAAGTCCGAAGGACGATGGGCTGCACTAT
CTTCTGTCACGCCCGGCAAACAAGCTATCGATTTGGCGCAGAAAGTGGATAAAGATCCACGGCTCGTTGAACTGATCCTA
TCGGAGTCGGCAGAAATCGTTGCACACAGGCAAGACGGTGTACTGATCACCGCTCACCGTCTCGGATGCGTGATGGCGAA
TGCCGGTATCGATCATTCCAATGTTGGTGATGAAGACAGTGTGCTTCTGCTTCCGAAAGACCCGGATCACAGTGCACGCG
AATTAAAAAAACAGTTTCATCGCTTGTGCGGCGTCGATGTCCACATCATCATCAACGATAGTTTTGGCAGAGTGTGGCGC
CATGGCACGGCAGGATGCGCAATCGGTGTAGCGGGCTTTTCCCCGCTTAAAAATTATATTGGGAAGCCGGATCTGTTTGG
GCAGCTCTTACGAACAACCCAAGTGGCAGTGGCCGATGAACTCGCCGCAGCAGCGTCTTTCTTAATGGGACAAGCCGATG
AAGGATCGCCTGTTGTGCTGATCCGCGGCGCGAATTTGCCCCCCGCTGATGGAACGGTCCAGCAGCTTATTCGCCCCAAA
GAGGAAGATTTATTTCGTGAAACCTCCTCCACACTGGCGGTGGAAATGGCCTAAACTAGTGGATCCGAATTCGAGCTCGG
CGCGCCTGCAGGTCGACAAGCTTGCGGCCGCATAATGCTTAAGTCGAACAGAAAGTAATCGTATTGTACACGGCCGCATA
ATCGAAATTAATACGACTCACTATAGGGGAATTGTGAGCGGATAACAATTCCCCATCTTAGTATATTAGTTAAGTATAAG
AAGGAGATATACATATGGCAGATCTCAATTGGATATCGGCCGGCCACGCGATCGCTGACGTCGGTACCCTCGAGTCTGGT
AAAGAAACCGCTGCTGCGAAATTTGAACGCCAGCACATGGACTCGTCTACTAGCGCAGCTTAATTAACCTAGGCTGCTGC
CACCGCTGAGCAATAACTAGCATAACCCCTTGGGGCCTCTAAACGGGTCTTGAGGGGTTTTTTGCTGAAAGGAGGAACTA
TATCCGGATTGGCGAATGGGACGCGCCCTGTAGCGGCGCATTAAGCGCGGCGGGTGTGGTGGTTACGCGCAGCGTGACCG
CTACACTTGCCAGCGCCCTAGCGCCCGCTCCTTTCGCTTTCTTCCCTTCCTTTCTCGCCACGTTCGCCGGCTTTCCCCGT
CAAGCTCTAAATCGGGGGCTCCCTTTAGGGTTCCGATTTAGTGCTTTACGGCACCTCGACCCCAAAAAACTTGATTAGGG
TGATGGTTCACGTAGTGGGCCATCGCCCTGATAGACGGTTTTTCGCCCTTTGACGTTGGAGTCCACGTTCTTTAATAGTG
GACTCTTGTTCCAAACTGGAACAACACTCAACCCTATCTCGGTCTATTCTTTTGATTTATAAGGGATTTTGCCGATTTCG
GCCTATTGGTTAAAAAATGAGCTGATTTAACAAAAATTTAACGCGAATTTTAACAAAATATTAACGTTTACAATTTCTGG
CGGCACGATGGCATGAGATTATCAAAAAGGATCTTCACCTAGATCCTTTTAAATTAAAAATGAAGTTTTAAATCAATCTA
AAGTATATATGAGTAAACTTGGTCTGACAGTTACCAATGCTTAATCAGTGAGGCACCTATCTCAGCGATCTGTCTATTTC
GTTCATCCATAGTTGCCTGACTCCCCGTCGTGTAGATAACTACGATACGGGAGGGCTTACCATCTGGCCCCAGTGCTGCA
ATGATACCGCGAGACCCACGCTCACCGGCTCCAGATTTATCAGCAATAAACCAGCCAGCCGGAAGGGCCGAGCGCAGAAG
TGGTCCTGCAACTTTATCCGCCTCCATCCAGTCTATTAATTGTTGCCGGGAAGCTAGAGTAAGTAGTTCGCCAGTTAATA
GTTTGCGCAACGTTGTTGCCATTGCTACAGGCATCGTGGTGTCACGCTCGTCGTTTGGTATGGCTTCATTCAGCTCCGGT
TCCCAACGATCAAGGCGAGTTACATGATCCCCCATGTTGTGCAAAAAAGCGGTTAGCTCCTTCGGTCCTCCGATCGTTGT
CAGAAGTAAGTTGGCCGCAGTGTTATCACTCATGGTTATGGCAGCACTGCATAATTCTCTTACTGTCATGCCATCCGTAA
GATGCTTTTCTGTGACTGGTGAGTACTCAACCAAGTCATTCTGAGAATAGTGTATGCGGCGACCGAGTTGCTCTTGCCCG
GCGTCAATACGGGATAATACCGCGCCACATAGCAGAACTTTAAAAGTGCTCATCATTGGAAAACGTTCTTCGGGGCGAAA
ACTCTCAAGGATCTTACCGCTGTTGAGATCCAGTTCGATGTAACCCACTCGTGCACCCAACTGATCTTCAGCATCTTTTA
CTTTCACCAGCGTTTCTGGGTGAGCAAAAACAGGAAGGCAAAATGCCGCAAAAAAGGGAATAAGGGCGACACGGAAATGT
TGAATACTCATACTCTTCCTTTTTCAATCATGATTGAAGCATTTATCAGGGTTATTGTCTCATGAGCGGATACATATTTG

AATGTATTTAGAAAAATAAACAAATAGGTCATGACCAAAATCCCTTAACGTGAGTTTTCGTTCCACTGAGCGTCAGACCC
CGTAGAAAAGATCAAAGGATCTTCTTGAGATCCTTTTTTTCTGCGCGTAATCTGCTGCTTGCAAACAAAAAAACCACCGC
TACCAGCGGTGGTTTGTTTGCCGGATCAAGAGCTACCAACTCTTTTTCCGAAGGTAACTGGCTTCAGCAGAGCGCAGATA
CCAAATACTGTCCTTCTAGTGTAGCCGTAGTTAGGCCACCACTTCAAGAACTCTGTAGCACCGCCTACATACCTCGCTCT
GCTAATCCTGTTACCAGTGGCTGCTGCCAGTGGCGATAAGTCGTGTCTTACCGGGTTGGACTCAAGACGATAGTTACCGG
ATAAGGCGCAGCGGTCGGGCTGAACGGGGGGTTCGTGCACACAGCCCAGCTTGGAGCGAACGACCTACACCGAACTGAGA
TACCTACAGCGTGAGCTATGAGAAAGCGCCACGCTTCCCGAAGGGAGAAAGGCGGACAGGTATCCGGTAAGCGGCAGGGT
CGGAACAGGAGAGCGCACGAGGGAGCTTCCAGGGGGAAACGCCTGGTATCTTTATAGTCCTGTCGGGTTTCGCCACCTCT
GACTTGAGCGTCGATTTTTGTGATGCTCGTCAGGGGGGCGGAGCCTATGGAAAAACGCCAGCAACGCGGCCTTTTTACGG
TTCCTGGCCTTTTGCTGGCCTTTTGCTCACATGTTCTTTCCTGCGTTATCCCCTGATTCTGTGGATAACCGTATTACCGC
CTTTGAGTGAGCTGATACCGCTCGCCGCAGCCGAACGACCGAGCGCAGCGAGTCAGTGAGCGAGGAAGCGGAAGAGCGCC
TGATGCGGTATTTTCTCCTTACGCATCTGTGCGGTATTTCACACCGCATATATGGTGCACTCTCAGTACAATCTGCTCTG
ATGCCGCATAGTTAAGCCAGTATACACTCCGCTATCGCTACGTGACTGGGTCATGGCTGCGCCCCGACACCCGCCAACAC
CCGCTGACGCGCCCTGACGGGCTTGTCTGCTCCCGGCATCCGCTTACAGACAAGCTGTGACCGTCTCCGGGAGCTGCATG
TGTCAGAGGTTTTCACCGTCATCACCGAAACGCGCGAGGCAGCTGCGGTAAAGCTCATCAGCGTGGTCGTGAAGCGATTC
ACAGATGTCTGCCTGTTCATCCGCGTCCAGCTCGTTGAGTTTCTCCAGAAGCGTTAATGTCTGGCTTCTGATAAAGCGGG
CCATGTTAAGGGCGGTTTTTTCCTGTTTGGTCACTGATGCCTCCGTGTAAGGGGGATTTCTGTTCATGGGGGTAATGATA
CCGATGAAACGAGAGAGGATGCTCACGATACGGGTTACTGATGATGAACATGCCCGGTTACTGGAACGTTGTGAGGGTAA
ACAACTGGCGGTATGGATGCGGCGGGACCAGAGAAAAATCACTCAGGGTCAATGCCAGCGCTTCGTTAATACAGATGTAG
GTGTTCCACAGGGTAGCCAGCAGCATCCTGCGATGCAGATCCGGAACATAATGGTGCAGGGCGCTGACTTCCGCGTTTCC
AGACTTTACGAAACACGGAAACCGAAGACCATTCATGTTGTTGCTCAGGTCGCAGACGTTTTGCAGCAGCAGTCGCTTCA
CGTTCGCTCGCGTATCGGTGATTCATTCTGCTAACCAGTAAGGCAACCCCGCCAGCCTAGCCGGGTCCTCAACGACAGGA
GCACGATCATGCTAGTCATGCCCCGCGCCCACCGGAAGGAGCTGACTGGGTTGAAGGCTCTCAAGGGCATCGGTCGAGAT
CCCGGTGCCTAATGAGTGAGCTAACTTACATTAATTGCGTTGCGCTCACTGCCCGCTTTCCAGTCGGGAAACCTGTCGTG
CCAGCTGCATTAATGAATCGGCCAACGCGCGGGGAGAGGCGGTTTGCGTATTGGGCGCCAGGGTGGTTTTTCTTTTCACC
AGTGAGACGGGCAACAGCTGATTGCCCTTCACCGCCTGGCCCTGAGAGAGTTGCAGCAAGCGGTCCACGCTGGTTTGCCC
CAGCAGGCGAAAATCCTGTTTGATGGTGGTTAACGGCGGGATATAACATGAGCTGTCTTCGGTATCGTCGTATCCCACTA
CCGAGATGTCCGCACCAACGCGCAGCCCGGACTCGGTAATGGCGCGCATTGCGCCCAGCGCCATCTGATCGTTGGCAACC
AGCATCGCAGTGGGAACGATGCCCTCATTCAGCATTTGCATGGTTTGTTGAAAACCGGACATGGCACTCCAGTCGCCTTC
CCGTTCCGCTATCGGCTGAATTTGATTGCGAGTGAGATATTTATGCCAGCCAGCCAGACGCAGACGCGCCGAGACAGAAC
TTAATGGGCCCGCTAACAGCGCGATTTGCTGGTGACCCAATGCGACCAGATGCTCCACGCCCAGTCGCGTACCGTCTTCA
TGGGAGAAAATAATACTGTTGATGGGTGTCTGGTCAGAGACATCAAGAAATAACGCCGGAACATTAGTGCAGGCAGCTTC
CACAGCAATGGCATCCTGGTCATCCAGCGGATAGTTAATGATCAGCCCACTGACGCGTTGCGCGAGAAGATTGTGCACCG
CCGCTTTACAGGCTTCGACGCCGCTTCGTTCTACCATCGACACCACCACGCTGGCACCCAGTTGATCGGCGCGAGATTTA
ATCGCCGCGACAATTTGCGACGGCGCGTGCAGGGCCAGACTGGAGGTGGCAACGCCAATCAGCAACGACTGTTTGCCCGC
CAGTTGTTGTGCCACGCGGTTGGGAATGTAATTCAGCTCCGCCATCGCCGCTTCCACTTTTTCCCGCGTTTTCGCAGAAA
CGTGGCTGGCCTGGTTCACCACGCGGGAAACGGTCTGATAAGAGACACCGGCATACTCTGCGACATCGTATAACGTTACT
GGTTTCACATTCACCACCCTGAATTGACTCTCTTCCGGGCGCTATCATGCCATACCGCGAAAGGTTTTGCGCCATTCGAT
GGTGTCCGGGATCTCGACGCTCTCCCTTATGCGACTCCTGCATTAGGAAGCAGCCCAGTAGTAGGTTGAGGCCGTTGAGC
ACCGCCGCCGCAAGGAATGGTGCATGCAAGGAGATGGCGCCCAACAGTCCCCCGGCCACGGGGCCTGCCACCATACCCAC
GCCGAAACAAGCGCTCATGAGCCCGAAGTGGCGAGCCCGATCTTCCCCATCGGTGATGTCGGCGATATAGGCGCCAGCAA
CCGCACCTGTGGCGCCGGTGATGCCGGCCACGATGCGTCCGGCGTAGAGGATCGAGATCGATCTCGATCCCGCGAAATTA
ATACGACTCACTATA
>pDB060
GGGGAATTGTGAGCGGATAACAATTCCCCTCTAGACCCGGGGGTGATCCGATGCAACTACAATGCCATGCTGGAAGCAGC
GAGTACGTGACACATGTCGGCAAAGCAGCGATGCCAACTCGTTACGGAAATTTTGTTGCCCATGCATTTCGTTCGACAAG
CGATAACAATGAGTATCTCGCGCTGGTGATGGGTGACGTTTGCCAACGTGAATCAGTACTGACGCGACTGCACTCCGAGT
GCCTCACCGGTGATGTACTAGGATCACTGCGCTGCGATTGTGGTGAACAATTGGACGCCGCATTACGCCATATCGCATCT
GAAGGCGTTGGCGCGCTATTGTATTTGCGAGGCCATGAAGGACGCGGCATTGGCTTGTGTAACAAAATTCTTGCATACGG
ACTCCAAGAACAAGGACTTGACACCGTCGATGCCAATCGCGACCTGGGTTTGCCGGATGACGCGCGCGAGTATGACTGCG
CGGCGAGCATCGTGCGCCAACTCGGCATCCTGTCGGTACGGCTGATGAGCAACAATCCGGATAAGTTTGAAGCGCTGCAG
CGCCATGGCATTCCAGTATGTGAGCGCGTTGACCTCGCAATTGCATTACGCGAGGAAAATGAGCGTTATATCTGGACTAA
GCGCAATCGTTTCGGGCATTATTTTGACGAGAGTGAGCTTCAGTGCCCGTTGTCATTGTAGAGGCCTATATGAGATGCAA
TGGTATGTCTCCTGTTTCTGTCGCCCAGCGCAATACGGGCATCTGGGCGGTCGTGCCGTTGAAAGCGCCTGAGTGTGCCA
AAACCCGGTTGTCTGGCGTTTTAAGCCATGCTGCGCGCCAAGCGTTGTTTTTTTCCATGGCCAGTCATGTCATTGGTACG
TTACGCGCATCGCCTCGGATTGCTTCCCTATTGGTGGTGACGCCGTCGGAAAGTACGGCGGAAATGGCCCGGGCAGCCGG
TGCTGAGATCTTGTGGGGACCACCGGACGAAGGCATGGCGAATGCCTGTTCGCGAGCGATGGCTCATATTGCGGCAGCAG
GCGGAGAGCGTGTGATGTTTGTCCCCGGTGATTTGCCCCTGTTGGATGGGGCGGCCATCGACATGTTGAGCCGTGCACCG
GTCGATGCGATTGGCATGGCGCCAAACCGGGACGGTCATGGCACGAATGGGCTGATTTGCCGACCTGGCGCTATTCCGTT
GTTTTTCAGCGGGCCAAGCTTTTCCGCTCATCAAAACGCCGCTCGGTGCGCCGGAATTGATGTTTGGATTGTCCGTTCAA
GAGAGTGGGCGTTGGATGTAGACTTGCCTGCCGACCTTGAGGAATTCGAGTCTTCCATCAAAGACGCCAAGAGGAGGGTG
CTATGCCAAATCTAGTAAGCAATCCTATTTACAAAATGATCCAAGAGGTCGAACGTTCCGCTAAGGGCAGTCAGCCAAGG
CTAGATGACAAGGCTTTGGCGTATGAGCTTGAAGCGGTGCAAGATTGGCGTGAGTTGGCGGGTCTGGCATCGGCATGGCG
CGATATCGGTTGGGGCAACGTCATCACCTATTCGCGCAAGGTTTTATTCCGCTCACCCATCTTTGTCGAGATGTTTGCC

ACTACTGCACCTTTGCCAAAGCGCCACGGGTGATTGGCCAAGCATTTCTTACCGTGGACCAAGCGCTGGACATTGCGCGG
GCGGGTGCTGCAGTTGGCTGTCGTGAGGCTTTGTTTACGCTCGGAGATCGGCCCGAAGCGCGTTATGCCGCTGCACGCGA
TGCGCTGCAGCAACTGGGTCATGCTACGACAGCCGACTATGTCGCTGAGGTCGCGCAACGCGTACGAGCTGAAACCGGGC
TGCTGCCCCACTTTAATATGGGCGTGCTAAGCGCGGCAGAATACCAGATGCTGCGGCCCCATGCCCCTTCCTTCGGGTTG
ATGCTAGAGACGGCTTCTGAACGTCTTTCCGAGCGTGGTGGGCCGCATTACGGCTCTCCCGATAAGCACCCGGCAGCGCG
GCTTGAAACGCTTCGCCTTGCCGGCGAAGCGCATATACCGATCACCTCAGGCATATTGATTGGTATCGGGGAGACACGCC
GCGAGCGGCTTGAATCGCTCTTTGCGCTGCGGGATTTGCATGATCGCTATGGTCATGTGCAGGAGGTCATTATTCAGAAC
TTTCGCGCGAAACCCGGCACCAAAATGGTTAACGCGCCTGAGCCGTCAATGGACGAATTGTGTTGGACAACAGCGGTCGC
CCGGTTGGTCCTGGGTTCCGCAATGAGCATCCAGGTGCCGCCTAATCTTTTCGACGGTGACTTGACCGATTTGATTCGCG
CGGGAATCAATGATTGGGGCGGCGTATCGCCGGTCACGCCTGATCACGTTAATCCAGAAGCGCCGTGGCCGCACCTTGAC
CGCTTAAGTGAGGATACAGACAGGGGCGGGAAGACGTTGACCGAGCGGATAACCGTCTATCCTGCCTATATCGAGGAACG
TGATCGTTGGATCGACCCGGGCTTGCATGCTGATGTGCTACGCCATTCAGATTGCAGTGGATTGGCAACAAGCGATATCT
GGAAAGCCGGCTCCACGACGATAGGGGATATCGCTTGCCTGGCAGTCACTCCCACGGCTTCCCAGGTTGATCGTGTTGGT
TCGCAGACGATCGATATCGTCGAGAAATGCGTGGCGGGTGAGCGCCTCGAAGAAGCCGAGCTTGTACATTTATTTAACGC
ACGCGGCCATGATTTTACGCATGTGACGCACACGGCGGACCGTTTGCGCAGGCAGACAAAGGGCGACACGATCACTTACG
TTGTCAATCGGAACATCAACTACACCAACGTCTGCCAATACCATTGCAATTTTTGTGCTTTTTCACGAGGACCGATACAA
GAAGACTTGCGCGAAAAGCCCTATATCGTTGACCTCAACGAAATCCGTCGCCGCGTCAAAGAAGCGTGGGATCGAGGGGC
GACGGAGGTCTGCTTGCAAGGGGGCATTCATCCCGACTATACGGGACGTACTTATCTTGACATCTGCGAAGCGGCTAAAA
CAGAATGTCCAGACATGCACATTCACGCCTTTTCGCCCCTTGAGGTGCTGCATGGTGCCACGACCCTGGGGACGTCAATC
AGCGCTTTTTTGTCTGATTTGAAAAGCGCTGGATTAAGTACGTTGCCTGGGACCGCAGCCGAGATCCTGGACGACGAAGT
GCGGCGACAAATCTGTCCGGATAAGTTGAATACGCAAGAATGGTTGACCGTCGTGCGATCGGCACATGAGTTGGGATTGC
GGACCACTTGCACGATCATGTTCGGTCATGTCGAATCGTATAAGCATTGGGCGCGACATATCCTCAGGCTTGCCGAGCTG
CAACGTGATACAGGCGGGCTGACTGAGTTTGTTCCATTGCCTTTTGTCCATGAGGAAGCGCCATTATTTAAAAAGAAAGG
TGCGCGTCAGGGGCCGACTTTACGAGAAGCCATTTGATGCATGCCGTCGGGCGTCTGGCGTTTCATGGCTTGATTGACA
ATATTCAAACGTCTTGGGTGAAAATGGGTGAGCACGGCGCACAGCTTTGCTTGCAGGCGGGGGCTAACGATTTGGGCGGT
ACCCTAATGAATGAATCGATTAGCCGCGCAGCCGGTGCAGCCCACGGACAGGAAATGCCGCCTGCCGCCATGGAAGCCCT
GGCCGCCAGACTCGGGCGCAATGCGATGCAGCGTACCCCGCTTTATCGTAACGTCAGTCCTGAGCGGCAGCAAGCTTCGC
ATGCTGCTCCTGCGCTGGTGCCTGTGCGATTCACTAAGGCCGGACGCTTGGTGCGTGGGCACGGCCATCAGGTCTTGAGT
GAAGCGGAACAATGATTGGACCTGCATAGGTGGATCATAGGTGGATTCATGGTGTGGCTTGCATGTTGGCAGCCTCAAAT
CGATGTTGTGATGGCTCTCACGCAGCGCTTCCCATCATGGATTCACCAAGGTGCTTAATTACCCTATTAAATTAGACAGA
CGCTATTCGGTTTGACACGATCGGAAGGGTGATCGGTTTGGGTCAGAGAAATGCTTGATCTTTACGCAGGAAGATAGTGA
TTACCGTTCTGAGCGGTGGTACAGGCACCCCGAAACTGCTGCAGGGTCTGAAACGTGTTGTTAATAATGAAGAACTGGCC
GTGATTGTGAATACCGGTGAAGATACCTGGATTGGTGATCTGTATCTGAGTCCGGATGTTGATACCGTGCTGTATACCCT
GGCAGATCTGATTAACGAAGAAACCTGGTATGGTGTGAAAGAGGATACCTTTTATACCCACGAACAGCTGAAAAATCTGG
GCTTTGATGAAGTTCTGCGCATTGGTGATAAAGATCGTGCACTGAAAATGCACAAAACCTATTATCTGAAACGCGGTCAT
AAACTGAGCGAAGTTGTTGATATGGAAAAAGTTGCCCTGGGCATTAAAGCAAAAGTTATTCCGATGACCGATGATCGTGT
GGAAACCAAAATTCTGGCAAAAGTTGATGGTAAAGTGGACCTGCTGAAATTCCATGATTTTTGGGTTAAACGCAAAGGTG
ATGTTGAAGTGCTGGATGTGATTTATGAAAACAGCCTGTATGCAAAACCGTGCGAAAAAGCAGTTGAAGCCATTAAAAAC
AGCGATCTGGTTATTATTGGTCCGAGCAATCCGATTACCAGCATCGGTCCGATTCTGAGCCTGAATGGTATTAAAGAACT
GCTGAAAGACAAAAAAGTTGTTGTGGTTAGCCCGATTGTTGGTAATAGCGCAGTTAGCGGTCCGGCAGGTAAACTGATGA
AAGCCAAAGGTTATGATGTTAGCGTGAAAGGCATCTACGAGTTCTATAAAGATATTGTGGATGTGCTGGTGATCGACAAC
GTGGATAAAGAAATTGCAAAAGAAATTCCGTGCGAAGTGCTGATTACCAATACCATCATGAAAACCCTGGATGATAAAGT
TCGTCTGGCCAAAAACATCATTGAATTTTGTGGTAGCCTGTAAGGTTGATGGCTATACAGAAATAAAAAATGATCAATGG
CATGCGGTACGGCAAACGCTTGCTGAACCACTCACGAAGAATGCACTATCCACTACGCATGCGCCGCCTTTGGGCTTTTG
TTTCTCCCTTGTCATCTTCGAATTGGCGCGGCGACTGGCAGACAATCGCTGCGCTATTTCCCTATTTGATGGTTTATAAA
GGGCGCGTCGCCTTCGCACTTGCCTGTCTGGTTGCCGCAAAACTCGCTAATCTAGGGGTTCCGGTGGTGATGAAAGCGAT
TGTCGACCAATTATCAGAAATTGACCGGTTGGCAACGCAGGAATCGTCATTTTCCTTGTTCCACGGCGTGAGTTTTCTTA
TTGTCGCTTATGCGGTCGTGCGTTTATCAAGCTCGCTTTTTGCTGAGCTGCGTGAAATTATCTTTGGAAAAGTGGCTTAC
AACGCTGCGCGACAAGTGGCACTCGCAGTATTCCGGCATCTGCACACGTTGTCGTTGTGTTTCACCTTGAGCGGCAAAC
AGGTGGCCTATCGCGCGATATTGAGCGAGGCACGCGTGGCGTCAAGACGCTTGTTTCATATTCGCTCTATAATATCCTGC
CAATATGTGTTGAAGTCATATTGGTACTCATTTTTTTTGCTATTCGTTACGACATCTATTATACGGTCGTCACGCTATCC
GTGCTTGCGGCTTACATCACGTTTACCGTGATGGTGACAGAATGGCGTACGCGTTTAAGGCAGGAGATGAATAAGCTTGA
TTCACGCTCCAATACGCTAATGGTTGATTCGCTTATTAATTACGAGACAGTCAAGAACTTTGGTAACGAGCAACACGAAG
TACAGCGCTATGATGAAAGCATGATGCTTTATCATGATGCGGCAGTTCGATCGCAGAAATCGCTTTCGTTCATGAATCTT
GGCCAGCAGTCGATTATTGCAATCTGCATGATTGCCGTTTTATGGAGGGCGACGCAGCAGGTTGTGGATAAGCAATTGAC
ATTGGGCGACTTTGTGTTAATCAATACATTTATGTTGCAGATATATATTCCGCTGAGTTTTTTGGGGAATATGTATCGAA
CTTTGAAGCAAAGTCTGACCGATATGGATCAGATGTTTTCTTTGTTAAGGCTCAGACGTGAAGTTGACGATATTCAAGGT
TCATCTCCGCTTGCTGTAAGAAGCGCTGAAGTACGCTTTGAGCATGTCAGTTTTTCATATGAACCGCAACGGCAAATTTT
GCGGGATGTCACCTTTACGATTGCAGCGGGTACGACGACCGCAATTGTCGGACATAGCGGCTCAGGTAAGTCGACACTCG
CACGGCTGTTGCTTCGCTTTTATGATGTAGAGCATGGCGCTGGTCGTATTTTGATTGATGGCCAAGACATTCGGGCTGTG
ACGCAAGACTCGCTACGCGCGGCCGTTGGCATTGTGCCGCAAGATACGGTGCTTTTCCGTGACACGATCTATTACAATAT
TGCGTACGGACGGTTGTCTGCGTCGCCAGAAGAAGTGATAGCGGCTGCGCGCGCCGCTCATATTCATGCTTTTATTGAAA
GCTTGCCCGCTGGCTATTCCACTATAGTGGGTGAGCGTGGTTTGAAATTATCTGGCGGTGAAAAACAGCGCATCGCAATT
GCCCGTACCTTGCTAAAAAAGCCGCCTATTCTCATTTTTGATGAAGCGACTTCAGCACTCGATTCACGTGCCGAGCGTGC

GATTCAGCGCGAGTTGAAACAGCTTGCGCGTCATCGTACAACACTAATCATTGCGCATCGTCTGTCGACCATTACGCATG
CGCAACAGATTCTTGTTATGGATCAAGGTCGTATCGTTGAGTGTGGCACCCACAGAACATTGCTGAATGCAGGTGGCCTA
TATGCGCAAATGTGGGCATTGCAACACAAGCAGCCTGAATAAGTTTAAACCAGCAAGAAGCATGACTGTATCCGCCATCG
GTGGCATTCCCTTAATACAAACAGGGGATGATCTTGGCCAGATTATTAAGGAGGCGATCCACAAAAACGGTATCGCTCTA
GAAAATGGCGACGTGTTAGTGCTTGCGCAAAAGATCGTTTCAAAGTCCGAAGGACGATGGGCTGCACTATCTTCTGTCAC
GCCCGGCAAACAAGCTATCGATTTGGCGCAGAAAGTGGATAAAGATCCACGGCTCGTTGAACTGATCCTATCGGAGTCGG
CAGAAATCGTTGCACACAGGCAAGACGGTGTACTGATCACCGCTCACCGTCTCGGATGCGTGATGGCGAATGCCGGTATC
GATCATTCCAATGTTGGTGATGAAGACAGTGTGCTTCTGCTTCCGAAAGACCCGGATCACAGTGCACGCGAATTAAAAAA
ACAGTTTCATCGCTTGTGCGGCGTCGATGTCCACATCATCATCAACGATAGTTTTGGCAGAGTGTGGCGCCATGGCACGG
CAGGATGCGCAATCGGTGTAGCGGGCTTTTCCCCGCTTAAAAATTATATTGGGAAGCCGGATCTGTTTGGGCAGCTCTTA
CGAACAACCCAAGTGGCAGTGGCCGATGAACTCGCCGCAGCAGCGTCTTTCTTAATGGGACAAGCCGATGAAGGATCGCC
TGTTGTGCTGATCCGCGGCGCGAATTTGCCCCCCGCTGATGGAACGGTCCAGCAGCTTATTCGCCCCAAAGAGGAAGATT
TATTTCGTGAAACCTCCTCCACACTGGCGGTGGAAATGGCCTAAACTAGTGGATCCGAATTCGAGCTCGGCGCGCCTGCA
GGTCGACAAGCTTGCGGCCGCATAATGCTTAAGTCGAACAGAAAGTAATCGTATTGTACACGGCCGCATAATCGAAATTA
ATACGACTCACTATAGGGGAATTGTGAGCGGATAACAATTCCCCATCTTAGTATATTAGTTAAGTATAAGAAGGAGATAT
ACATATGGCAGATCTCAATTGGATATCGGCCGGCCACGCGATCGCTGACGTCGGTACCCTCGAGTCTGGTAAAGAAACCG
CTGCTGCGAAATTTGAACGCCAGCACATGGACTCGTCTACTAGCGCAGCTTAATTAACCTAGGCTGCTGCCACCGCTGAG
CAATAACTAGCATAACCCCTTGGGGCCTCTAAACGGGTCTTGAGGGGTTTTTTGCTGAAAGGAGGAACTATATCCGGATT
GGCGAATGGGACGCGCCCTGTAGCGGCGCATTAAGCGCGGCGGGTGTGGTGGTTACGCGCAGCGTGACCGCTACACTTGC
CAGCGCCCTAGCGCCCGCTCCTTTCGCTTTCTTCCCTTCCTTTCTCGCCACGTTCGCCGGCTTTCCCCGTCAAGCTCTAA
ATCGGGGGCTCCCTTTAGGGTTCCGATTTAGTGCTTTACGGCACCTCGACCCCAAAAAACTTGATTAGGGTGATGGTTCA
CGTAGTGGGCCATCGCCCTGATAGACGGTTTTTCGCCCTTTGACGTTGGAGTCCACGTTCTTTAATAGTGGACTCTTGTT
CCAAACTGGAACAACACTCAACCCTATCTCGGTCTATTCTTTTGATTTATAAGGGATTTTGCCGATTTCGGCCTATTGGT
TAAAAAATGAGCTGATTTAACAAAAATTTAACGCGAATTTTAACAAAATATTAACGTTTACAATTTCTGGCGGCACGATG
GCATGAGATTATCAAAAAGGATCTTCACCTAGATCCTTTTAAATTAAAAATGAAGTTTTAAATCAATCTAAAGTATATAT
GAGTAAACTTGGTCTGACAGTTACCAATGCTTAATCAGTGAGGCACCTATCTCAGCGATCTGTCTATTTCGTTCATCCAT
AGTTGCCTGACTCCCCGTCGTGTAGATAACTACGATACGGGAGGGCTTACCATCTGGCCCCAGTGCTGCAATGATACCGC
GAGACCCACGCTCACCGGCTCCAGATTTATCAGCAATAAACCAGCCAGCCGGAAGGGCCGAGCGCAGAAGTGGTCCTGCA
ACTTTATCCGCCTCCATCCAGTCTATTAATTGTTGCCGGGAAGCTAGAGTAAGTAGTTCGCCAGTTAATAGTTTGCGCAA
CGTTGTTGCCATTGCTACAGGCATCGTGGTGTCACGCTCGTCGTTTGGTATGGCTTCATTCAGCTCCGGTTCCCAACGAT
CAAGGCGAGTTACATGATCCCCCATGTTGTGCAAAAAAGCGGTTAGCTCCTTCGGTCCTCCGATCGTTGTCAGAAGTAAG
TTGGCCGCAGTGTTATCACTCATGGTTATGGCAGCACTGCATAATTCTCTTACTGTCATGCCATCCGTAAGATGCTTTTC
TGTGACTGGTGAGTACTCAACCAAGTCATTCTGAGAATAGTGTATGCGGCGACCGAGTTGCTCTTGCCCGGCGTCAATAC
GGGATAATACCGCGCCACATAGCAGAACTTTAAAAGTGCTCATCATTGGAAAACGTTCTTCGGGGCGAAAACTCTCAAGG
ATCTTACCGCTGTTGAGATCCAGTTCGATGTAACCCACTCGTGCACCCAACTGATCTTCAGCATCTTTTACTTTCACCAG
CGTTTCTGGGTGAGCAAAAACAGGAAGGCAAAATGCCGCAAAAAAGGGAATAAGGGCGACACGGAAATGTTGAATACTCA
TACTCTTCCTTTTTCAATCATGATTGAAGCATTTATCAGGGTTATTGTCTCATGAGCGGATACATATTTGAATGTATTTA
GAAAAATAAACAAATAGGTCATGACCAAAATCCCTTAACGTGAGTTTTCGTTCCACTGAGCGTCAGACCCCGTAGAAAAG
ATCAAAGGATCTTCTTGAGATCCTTTTTTTCTGCGCGTAATCTGCTGCTTGCAAACAAAAAAACCACCGCTACCAGCGGT
GGTTTGTTTGCCGGATCAAGAGCTACCAACTCTTTTTCCGAAGGTAACTGGCTTCAGCAGAGCGCAGATACCAAATACTG
TCCTTCTAGTGTAGCCGTAGTTAGGCCACCACTTCAAGAACTCTGTAGCACCGCCTACATACCTCGCTCTGCTAATCCTG
TTACCAGTGGCTGCTGCCAGTGGCGATAAGTCGTGTCTTACCGGGTTGGACTCAAGACGATAGTTACCGGATAAGGCGCA
GCGGTCGGGCTGAACGGGGGGTTCGTGCACACAGCCCAGCTTGGAGCGAACGACCTACACCGAACTGAGATACCTACAGC
GTGAGCTATGAGAAAGCGCCACGCTTCCCGAAGGGAGAAAGGCGGACAGGTATCCGGTAAGCGGCAGGGTCGGAACAGGA
GAGCGCACGAGGGAGCTTCCAGGGGGAAACGCCTGGTATCTTTATAGTCCTGTCGGGTTTCGCCACCTCTGACTTGAGCG
TCGATTTTTGTGATGCTCGTCAGGGGGGCGGAGCCTATGGAAAAACGCCAGCAACGCGGCCTTTTTACGGTTCCTGGCCT
TTTGCTGGCCTTTTGCTCACATGTTCTTTCCTGCGTTATCCCCTGATTCTGTGGATAACCGTATTACCGCCTTTGAGTGA
GCTGATACCGCTCGCCGCAGCCGAACGACCGAGCGCAGCGAGTCAGTGAGCGAGGAAGCGGAAGAGCGCCTGATGCGGTA
TTTTCTCCTTACGCATCTGTGCGGTATTTCACACCGCATATATGGTGCACTCTCAGTACAATCTGCTCTGATGCCGCATA
GTTAAGCCAGTATACACTCCGCTATCGCTACGTGACTGGGTCATGGCTGCGCCCCGACACCCGCCAACACCCGCTGACGC
GCCCTGACGGGCTTGTCTGCTCCCGGCATCCGCTTACAGACAAGCTGTGACCGTCTCCGGGAGCTGCATGTGTCAGAGGT
TTTCACCGTCATCACCGAAACGCGCGAGGCAGCTGCGGTAAAGCTCATCAGCGTGGTCGTGAAGCGATTCACAGATGTCT
GCCTGTTCATCCGCGTCCAGCTCGTTGAGTTTCTCCAGAAGCGTTAATGTCTGGCTTCTGATAAAGCGGGCCATGTTAAG
GGCGGTTTTTTCCTGTTTGGTCACTGATGCCTCCGTGTAAGGGGGATTTCTGTTCATGGGGGTAATGATACCGATGAAAC
GAGAGAGGATGCTCACGATACGGGTTACTGATGATGAACATGCCCGGTTACTGGAACGTTGTGAGGGTAAACAACTGGCG
GTATGGATGCGGCGGGACCAGAGAAAAATCACTCAGGGTCAATGCCAGCGCTTCGTTAATACAGATGTAGGTGTTCCACA
GGGTAGCCAGCAGCATCCTGCGATGCAGATCCGGAACATAATGGTGCAGGGCGCTGACTTCCGCGTTTCCAGACTTTACG
AAACACGGAAACCGAAGACCATTCATGTTGTTGCTCAGGTCGCAGACGTTTTGCAGCAGCAGTCGCTTCACGTTCGCTCG
CGTATCGGTGATTCATTCTGCTAACCAGTAAGGCAACCCCGCCAGCCTAGCCGGGTCCTCAACGACAGGAGCACGATCAT
GCTAGTCATGCCCCGCGCCCACCGGAAGGAGCTGACTGGGTTGAAGGCTCTCAAGGGCATCGGTCGAGATCCCGGTGCCT
AATGAGTGAGCTAACTTACATTAATTGCGTTGCGCTCACTGCCCGCTTTCCAGTCGGGAAACCTGTCGTGCCAGCTGCAT
TAATGAATCGGCCAACGCGCGGGGAGAGGCGGTTTGCGTATTGGGCGCCAGGGTGGTTTTTCTTTTCACCAGTGAGACGG
GCAACAGCTGATTGCCCTTCACCGCCTGGCCCTGAGAGAGTTGCAGCAAGCGGTCCACGCTGGTTTGCCCCAGCAGGCGA
AAATCCTGTTTGATGGTGGTTAACGGCGGGATATAACATGAGCTGTCTTCGGTATCGTCGTATCCCACTACCGAGATGTC

CGCACCAACGCGCAGCCCGGACTCGGTAATGGCGCGCATTGCGCCCAGCGCCATCTGATCGTTGGCAACCAGCATCGCAG
TGGGAACGATGCCCTCATTCAGCATTTGCATGGTTTGTTGAAAACCGGACATGGCACTCCAGTCGCCTTCCCGTTCCGCT
ATCGGCTGAATTTGATTGCGAGTGAGATATTTATGCCAGCCAGCCAGACGCAGACGCGCCGAGACAGAACTTAATGGGCC
CGCTAACAGCGCGATTTGCTGGTGACCCAATGCGACCAGATGCTCCACGCCCAGTCGCGTACCGTCTTCATGGGAGAAAA
TAATACTGTTGATGGGTGTCTGGTCAGAGACATCAAGAAATAACGCCGGAACATTAGTGCAGGCAGCTTCCACAGCAATG
GCATCCTGGTCATCCAGCGGATAGTTAATGATCAGCCCACTGACGCGTTGCGCGAGAAGATTGTGCACCGCCGCTTTACA
GGCTTCGACGCCGCTTCGTTCTACCATCGACACCACCACGCTGGCACCCAGTTGATCGGCGCGAGATTTAATCGCCGCGA
CAATTTGCGACGGCGCGTGCAGGGCCAGACTGGAGGTGGCAACGCCAATCAGCAACGACTGTTTGCCCGCCAGTTGTTGT
GCCACGCGGTTGGGAATGTAATTCAGCTCCGCCATCGCCGCTTCCACTTTTTCCCGCGTTTTCGCAGAAACGTGGCTGGC
CTGGTTCACCACGCGGGAAACGGTCTGATAAGAGACACCGGCATACTCTGCGACATCGTATAACGTTACTGGTTTCACAT
TCACCACCCTGAATTGACTCTCTTCCGGGCGCTATCATGCCATACCGCGAAAGGTTTTGCGCCATTCGATGGTGTCCGGG
ATCTCGACGCTCTCCCTTATGCGACTCCTGCATTAGGAAGCAGCCCAGTAGTAGGTTGAGGCCGTTGAGCACCGCCGCCG
CAAGGAATGGTGCATGCAAGGAGATGGCGCCCAACAGTCCCCCGGCCACGGGGCCTGCCACCATACCCACGCCGAAACAA
GCGCTCATGAGCCCGAAGTGGCGAGCCCGATCTTCCCCATCGGTGATGTCGGCGATATAGGCGCCAGCAACCGCACCTGT
GGCGCCGGTGATGCCGGCCACGATGCGTCCGGCGTAGAGGATCGAGATCGATCTCGATCCCGCGAAATTAATACGACTCA
CTATA

>pDB061

CCGACACCATCGAATGGTGCAAAACCTTTCGCGGTATGGCATGATAGCGCCCGGAAGAGAGTCAATTCAGGGTGGTGAAT
GTGAAACCAGTAACGTTATACGATGTCGCAGAGTATGCCGGTGTCTCTTATCAGACCGTTTCCCGCGTGGTGAACCAGGC
CAGCCACGTTTCTGCGAAAACGCGGGAAAAAGTGGAAGCGGCGATGGCGGAGCTGAATTACATTCCCAACCGCGTGGCAC
AACAACTGGCGGGCAAACAGTCGTTGCTGATTGGCGTTGCCACCTCCAGTCTGGCCCTGCACGCGCCGTCGCAAATTGTC
GCGGCGATTAAATCTCGCGCCGATCAACTGGGTGCCAGCGTGGTGGTGTCGATGGTAGAACGAAGCGGCGTCGAAGCCTG
TAAAGCGGCGGTGCACAATCTTCTCGCGCAACGCGTCAGTGGGCTGATCATTAACTATCCGCTGGATGACCAGGATGCCA
TTGCTGTGGAAGCTGCCTGCACTAATGTTCCGGCGTTATTTCTTGATGTCTCTGACCAGACACCCATCAACAGTATTATT
TTCTCCCATGAAGACGGTACGCGACTGGGCGTGGAGCATCTGGTCGCATTGGGTCACCAGCAAATCGCGCTGTTAGCGGG
CCCATTAAGTTCTGTCTCGGCGCGTCTGCGTCTGGCTGGCTGGCATAAATATCTCACTCGCAATCAAATTCAGCCGATAG
CGGAACGGGAAGGCGACTGGAGTGCCATGTCCGGTTTTCAACAAACCATGCAAATGCTGAATGAGGGCATCGTTCCCACT
GCGATGCTGGTTGCCAACGATCAGATGGCGCTGGGCGCAATGCGCGCCATTACCGAGTCCGGGCTGCGCGTTGGTGCGGA
TATCTCGGTAGTGGGATACGACGATACCGAAGACAGCTCATGTTATATCCCGCCGTTAACCACCATCAAACAGGATTTTC
GCCTGCTGGGGCAAACCAGCGTGGACCGCTTGCTGCAACTCTCTCAGGGCCAGGCGGTGAAGGGCAATCAGCTGTTGCCC
GTCTCACTGGTGAAAAGAAAAACCACCCTGGCGCCCAATACGCAAACCGCCTCTCCCCGCGCGTTGGCCGATTCATTAAT
GCAGCTGGCACGACAGGTTTCCCGACTGGAAAGCGGGCAGTGAGCGCAACGCAATTAATGTAAGTTAGCTCACTCATTAG
GCACAATTCTCATGTTTGACAGCTTATCATCGACTGCACGGTGCACCAATGCTTCTGGCGTCAGGCAGCCATCGGAAGCT
GTGGTATGGCTGTGCAGGTCGTAAATCACTGCATAATTCGTGTCGCTCAAGGCGCACTCCCGTTCTGGATAATGTTTTTT
GCGCCGACATCATAACGGTTCTGGCAAATATTCTGAAATGAGCTGTTGACAATTAATCATCGGCTCGTATAATGTGTGGA
ATTGTGAGCGGATAACAATTTCACACAGGAAACAGCCAGTCCGTTTAGGTGTTTTCACGAGCACTTCACCAACAAGGACC
ATAGCATATGAAAATCGAAGAAGGTAAACTGGTAATCTGGATTAACGGCGATAAAGGCTATAACGGTCTCGCTGAAGTCG
GTAAGAAATTCGAGAAAGATACCGGAATTAAAGTCACCGTTGAGCATCCGGATAAACTGGAAGAGAAATTCCCACAGGTT
GCGGCAACTGGCGATGGCCCTGACATTATCTTCTGGGCACACGACCGCTTTGGTGGCTACGCTCAATCTGGCCTGTTGGC
TGAAATCACCCCGGACAAAGCGTTCCAGGACAAGCTGTATCCGTTTACCTGGGATGCCGTACGTTACAACGGCAAGCTGA
TTGCTTACCCGATCGCTGTTGAAGCGTTATCGCTGATTTATAACAAAGATCTGCTGCCGAACCCGCCAAAAACCTGGGAA
GAGATCCCGGCGCTGGATAAAGAACTGAAAGCGAAAGGTAAGAGCGCGCTGATGTTCAACCTGCAAGAACCGTACTTCAC
CTGGCCGCTGATTGCTGCTGACGGGGGTTATGCGTTCAAGTATGAAAACGGCAAGTACGACATTAAAGACGTGGGCGTGG
ATAACGCTGGCGCGAAAGCGGGTCTGACCTTCCTGGTTGACCTGATTAAAAACAAACACATGAATGCAGACACCGATTAC
TCCATCGCAGAAGCTGCCTTTAATAAAGGCGAAACAGCGATGACCATCAACGGCCCGTGGGCATGGTCCAACATCGACAC
CAGCAAAGTGAATTATGGTGTAACGGTACTGCCGACCTTCAAGGGTCAACCATCCAAACCGTTCGTTGGCGTGCTGAGCG
CAGGTATTAACGCCGCCAGTCCGAACAAAGAGCTGGCAAAAGAGTTCCTCGAAAACTATCTGCTGACTGATGAAGGTCTG
GAAGCGGTTAATAAAGACAAACCGCTGGGTGCCGTAGCGCTGAAGTCTTACGAGGAAGAGTTGGCGAAAGATCCACGTAT
TGCCGCCACTATGGAAAACGCCCAGAAAGGTGAAATCATGCCGAACATCCCGCAGATGTCCGCTTTCTGGTATGCCGTGC
GTACTGCGGTGATCAACGCCGCCAGCGGTCGTCAGACTGTCGATGAAGCCCTGAAAGACGCGCAGACTAATTCGAGCTCG
AACAACAACAACAATAACAATAACAACAACCTCGGGATCGAGGGAAGGATTTCAGAAAATCTTTATTTTCAAGGTCATCA
TCATCATCATCATAGCAGCGGCATGGCAGATACCAGCCGTCCGCTGAATGCAAAACAGCTGGAACGTCTGAATGCCAAAA
GCACCGGTACACTGATTAAATGGATGAGCCGTTTTCAGACCTTTCTGTTTAAAACCACCAATGGTAAACTGGGCAACAAA
TTTCTGCGTGGCACCGAAGTTGGTATTCTGACCACCATTGGTCGTAAAAGCGGTGAACCGCGTGATACACCGCTGCTGTT
TCTGCAAGAAGGTCGTCGTATTGTTCTGGTTGCAAGCCAAGGTGGTCGTGCAACCAATCCGATGTGGTATCTGAATCTGA
AAGCAAATCCGAAAGTTACCTTTCAGACCCGTAGCGAAAAACTGGCACTGGTTGCCCGTGAAGCAACCGATGCAGAACGT
GATGAATATTGGCCGAAACTGGATGCAATGTATCCGGATTTTGCAAACTATCGTAGCTATACCGATCGCAAAATTCCGAT
TGTTATTTGCGATCCGGCATAAGTCGACCTGCAGGCAAGCTTGGCACTGGCCGTCGTTTTACAACGTCGTGACTGGGAAA
ACCCTGGCGTTACCCAACTTAATCGCCTTGCAGCACATCCCCCTTTCGCCAGCTGGCGTAATAGCGAAGAGGCCCGCACC
GATCGCCCTTCCCAACAGTTGCGCAGCCTGAATGGCGAATGGCAGCTTGGCTGTTTTGGCGGATGAGATAAGATTTTCAG
CCTGATACAGATTAAATCAGAACGCAGAAGCGGTCTGATAAAACAGAATTTGCCTGGCGGCAGTAGCGCGGTGGTCCCAC
CTGACCCCATGCCGAACTCAGAAGTGAAACGCCGTAGCGCCGATGGTAGTGTGGGGTCTCCCCATGCGAGAGTAGGGAAC
TGCCAGGCATCAAATAAAACGAAAGGCTCAGTCGAAAGACTGGGCCTTTCGTTTTATCTGTTGTTTGTCGGTGAACGCTC
TCCTGAGTAGGACAAATCCGCCGGGAGCGGATTTGAACGTTGCGAAGCAACGGCCCGGAGGGTGGCGGGCAGGACGCCCG

CCATAAACTGCCAGGCATCAAATTAAGCAGAAGGCCATCCTGACGGATGGCCTTTTTGCGTTTCTACAAACTCTTTTGTT
TATTTTTCTAAATACATTCAAATATGTATCCGCTCATGAGACAATAACCCTGATAAATGCTTCAATAATATTGAAAAAGG
AAGAGTATGAGTATTCAACATTTCCGTGTCGCCCTTATTCCCTTTTTTGCGGCATTTTGCCTTCCTGTTTTTGCTCACCC
AGAAACGCTGGTGAAAGTAAAAGATGCTGAAGATCAGTTGGGTGCACGAGTGGGTTACATCGAACTGGATCTCAACAGCG
GTAAGATCCTTGAGAGTTTTCGCCCCGAAGAACGTTTTCCAATGATGAGCACTTTTAAAGTTCTGCTATGTGGCGCGGTA
TTATCCCGTGTTGACGCCGGGCAAGAGCAACTCGGTCGCCGCATACACTATTCTCAGAATGACTTGGTTGAGTACTCACC
AGTCACAGAAAAGCATCTTACGGATGGCATGACAGTAAGAGAATTATGCAGTGCTGCCATAACCATGAGTGATAACACTG
CGGCCAACTTACTTCTGACAACGATCGGAGGACCGAAGGAGCTAACCGCTTTTTTGCACAACATGGGGGATCATGTAACT
CGCCTTGATCGTTGGGAACCGGAGCTGAATGAAGCCATACCAAACGACGAGCGTGACACCACGATGCCTGTAGCAATGGC
AACAACGTTGCGCAAACTATTAACTGGCGAACTACTTACTCTAGCTTCCCGGCAACAATTAATAGACTGGATGGAGGCGG
ATAAAGTTGCAGGACCACTTCTGCGCTCGGCCCTTCCGGCTGGCTGGTTTATTGCTGATAAATCTGGAGCCGGTGAGCGT
GGGTCTCGCGGTATCATTGCAGCACTGGGGCCAGATGGTAAGCCCTCCCGTATCGTAGTTATCTACACGACGGGGAGTCA
GGCAACTATGGATGAACGAAATAGACAGATCGCTGAGATAGGTGCCTCACTGATTAAGCATTGGTAACTGTCAGACCAAG
TTTACTCATATATACTTTAGATTGATTTACCCCGGTTGATAATCAGAAAAGCCCCAAAAACAGGAAGATTGTATAAGCAA
ATATTTAAATTGTAAACGTTAATATTTTGTTAAAATTCGCGTTAAATTTTTGTTAAATCAGCTCATTTTTTAACCAATAG
GCCGAAATCGGCAAAATCCCTTATAAATCAAAAGAATAGACCGAGATAGGGTTGAGTGTTGTTCCAGTTTGGAACAAGAG
TCCACTATTAAAGAACGTGGACTCCAACGTCAAAGGGCGAAAAACCGTCTATCAGGGCGATGGCCCACTACGTGAACCAT
CACCCAAATCAAGTTTTTTGGGGTCGAGGTGCCGTAAAGCACTAAATCGGAACCCTAAAGGGAGCCCCCGATTTAGAGCT
TGACGGGGAAAGCCGGCGAACGTGGCGAGAAAGGAAGGGAAGAAAGCGAAAGGAGCGGGCGCTAGGGCGCTGGCAAGTGT
AGCGGTCACGCTGCGCGTAACCACCACACCCGCCGCGCTTAATGCGCCGCTACAGGGCGCGTAAAAGGATCTAGGTGAAG
ATCCTTTTTGATAATCTCATGACCAAAATCCCTTAACGTGAGTTTTCGTTCCACTGAGCGTCAGACCCCGTAGAAAAGAT
CAAAGGATCTTCTTGAGATCCTTTTTTTCTGCGCGTAATCTGCTGCTTGCAAACAAAAAAACCACCGCTACCAGCGGTGG
TTTGTTTGCCGGATCAAGAGCTACCAACTCTTTTTCCGAAGGTAACTGGCTTCAGCAGAGCGCAGATACCAAATACTGTC
CTTCTAGTGTAGCCGTAGTTAGGCCACCACTTCAAGAACTCTGTAGCACCGCCTACATACCTCGCTCTGCTAATCCTGTT
ACCAGTGGCTGCTGCCAGTGGCGATAAGTCGTGTCTTACCGGGTTGGACTCAAGACGATAGTTACCGGATAAGGCGCAGC
GGTCGGGCTGAACGGGGGGTTCGTGCACACAGCCCAGCTTGGAGCGAACGACCTACACCGAACTGAGATACCTACAGCGT
GAGCTATGAGAAAGCGCCACGCTTCCCGAAGGGAGAAAGGCGGACAGGTATCCGGTAAGCGGCAGGGTCGGAACAGGAGA
GCGCACGAGGGAGCTTCCAGGGGGAAACGCCTGGTATCTTTATAGTCCTGTCGGGTTTCGCCACCTCTGACTTGAGCGTC
GATTTTTGTGATGCTCGTCAGGGGGGCGGAGCCTATGGAAAAACGCCAGCAACGCGGCCTTTTTACGGTTCCTGGCCTTT
TGCTGGCCTTTTGCTCACATGTTCTTTCCTGCGTTATCCCCTGATTCTGTGGATAACCGTATTACCGCCTTTGAGTGAGC
TGATACCGCTCGCCGCAGCCGAACGACCGAGCGCAGCGAGTCAGTGAGCGAGGAAGCGGAAGAGCGCCTGATGCGGTATT
TTCTCCTTACGCATCTGTGCGGTATTTCACACCGCATATATGGTGCACTCTCAGTACAATCTGCTCTGATGCCGCATAGT
TAAGCCAGTATACACTCCGCTATCGCTACGTGACTGGGTCATGGCTGCGCCCCGACACCCGCCAACACCCGCTGACGCGC
CCTGACGGGCTTGTCTGCTCCCGGCATCCGCTTACAGACAAGCTGTGACCGTCTCCGGGAGCTGCATGTGTCAGAGGTTT
TCACCGTCATCACCGAAACGCGCGAGGCAGCTGCGGTAAAGCTCATCAGCGTGGTCGTGCAGCGATTCACAGATGTCTGC
CTGTTCATCCGCGTCCAGCTCGTTGAGTTTCTCCAGAAGCGTTAATGTCTGGCTTCTGATAAAGCGGGCCATGTTAAGGG
CGGTTTTTTCCTGTTTGGTCACTGATGCCTCCGTGTAAGGGGGATTTCTGTTCATGGGGGTAATGATACCGATGAAACGA
GAGAGGATGCTCACGATACGGGTTACTGATGATGAACATGCCCGGTTACTGGAACGTTGTGAGGGTAAACAACTGGCGGT
ATGGATGCGGCGGGACCAGAGAAAAATCACTCAGGGTCAATGCCAGCGCTTCGTTAATACAGATGTAGGTGTTCCACAGG
GTAGCCAGCAGCATCCTGCGATGCAGATCCGGAACATAATGGTGCAGGGCGCTGACTTCCGCGTTTCCAGACTTTACGAA
ACACGGAAACCGAAGACCATTCATGTTGTTGCTCAGGTCGCAGACGTTTTGCAGCAGCAGTCGCTTCACGTTCGCTCGCG
TATCGGTGATTCATTCTGCTAACCAGTAAGGCAACCCCGCCAGCCTAGCCGGGTCCTCAACGACAGGAGCACGATCATGC
GCACCCGTGGCCAGGACCCAACGCTGCCCGAAATT
>pDB065
GGGGAATTGTGAGCGGATAACAATTCCCCTGTAGAAATAATTTTGTTTAACTTTAATAAGGAGATATACCATGCGTGTGG
CTCTGCTGGGCGGTACGGGCAACCTGGGCAAAGGTCTGGCACTGCGTCTGGCAACCCTGGGTCATGAAATCGTGGTCGGC
TCACGTCGCGAAGAAAAAGCGGAAGCCAAAGCGGCCGAATATCGTCGCATTGCAGGCGATGCTTCGATCACCGGTATGAA
AAACGAAGACGCAGCTGAAGCGTGCGATATTGCCGTGCTGACCATCCCGTGGGAACATGCAATTGACACGGCTCGTGATC
TGAAAAATATTCTGCGCGAAAAAATCGTTGTTAGTCCGCTGGTGCCGGTTTCCCGTGGTGCCAAAGGTTTTACCTACAGC
TCTGAACGCTCAGCGGCCGAAATTGTTGCCGAAGTCCTGGAAAGCGAAAAAGTCGTGTCTGCCCTGCACACGATCCCGGC
AGCTCGTTTTGCAAACCTGGATGAAAAATTCGACTGGGATGTCCCGGTGTGTGGCGATGACGATGAAAGCAAAAAAGTTG
TCATGTCACTGATTTCGGAAATTGATGGTCTGCGTCCGCTGGATGCCGGTCCGCTGAGTAATTCCCGCCTGGTTGAATCT
CTGACGCCGCTGATTCTGAACATTATGCGTTTTAACGGTATGGGCGAACTGGGTATCAAATTTCTGTGAGGATCCGAATT
CGAGCTCGGCGCGCCTGCAGGTCGACAAGCTTGCGGCCGCATAATGCTTAAGTCGAACAGAAAGTAATCGTATTGTACAC
GGCCGCATAATCGAAATTAATACGACTCACTATAGGGGAATTGTGAGCGGATAACAATTCCCCATCTTAGTATATTAGTT
AAGTATAAGAAGGAGATATACATATGGCAGATCTCAATTGGATATCGGCCGGCCACGCGATCGCTGACGTCGGTACCCTC
GAGTCTGGTAAAGAAACCGCTGCTGCGAAATTTGAACGCCAGCACATGGACTCGTCTACTAGCGCAGCTTAATTAACCTA
GGCTGCTGCCACCGCTGAGCAATAACTAGCATAACCCCTTGGGGCCTCTAAACGGGTCTTGAGGGGTTTTTTGCTGAAAC
CTCAGGCATTTGAGAAGCACACGGTCACACTGCTTCCGGTAGTCAATAAACCGGTAAACCAGCAATAGACATAAGCGGCT
ATTTAACGACCCTGCCCTGAACCGACGACCGGGTCATCGTGGCCGGATCTTGCGGCCCCTCGGCTTGAACGAATTGTTAG
ACATTATTTGCCGACTACCTTGGTGATCTCGCCTTTCACGTAGTGGACAAATTCTTCCAACTGATCTGCGCGCGAGGCCA
AGCGATCTTCTTCTTGTCCAAGATAAGCCTGTCTAGCTTCAAGTATGACGGGCTGATACTGGGCCGGCAGGCGCTCCATT
GCCCAGTCGGCAGCGACATCCTTCGGCGCGATTTTGCCGGTTACTGCGCTGTACCAAATGCGGGACAACGTAAGCACTAC
ATTTCGCTCATCGCCAGCCCAGTCGGGCGGCGAGTTCCATAGCGTTAAGGTTTCATTTAGCGCCTCAAATAGATCCTGTT

CAGGAACCGGATCAAAGAGTTCCTCCGCCGCTGGACCTACCAAGGCAACGCTATGTTCTCTTGCTTTTGTCAGCAAGATA
GCCAGATCAATGTCGATCGTGGCTGGCTCGAAGATACCTGCAAGAATGTCATTGCGCTGCCATTCTCCAAATTGCAGTTC
GCGCTTAGCTGGATAACGCCACGGAATGATGTCGTCGTGCACAACAATGGTGACTTCTACAGCGCGGAGAATCTCGCTCT
CTCCAGGGGAAGCCGAAGTTTCCAAAAGGTCGTTGATCAAAGCTCGCCGCGTTGTTTCATCAAGCCTTACGGTCACCGTA
ACCAGCAAATCAATATCACTGTGTGGCTTCAGGCCGCCATCCACTGCGGAGCCGTACAAATGTACGGCCAGCAACGTCGG
TTCGAGATGGCGCTCGATGACGCCAACTACCTCTGATAGTTGAGTCGATACTTCGGCGATCACCGCTTCCCTCATACTCT
TCCTTTTTCAATATTATTGAAGCATTTATCAGGGTTATTGTCTCATGAGCGGATACATATTTGAATGTATTTAGAAAAAT
AAACAAATAGCTAGCTCACTCGGTCGCTACGCTCCGGGCGTGAGACTGCGGCGGGCGCTGCGGACACATACAAAGTTACC
CACAGATTCCGTGGATAAGCAGGGGACTAACATGTGAGGCAAAACAGCAGGGCCGCGCCGGTGGCGTTTTTCCATAGGCT
CCGCCCTCCTGCCAGAGTTCACATAAACAGACGCTTTTCCGGTGCATCTGTGGGAGCCGTGAGGCTCAACCATGAATCTG
ACAGTACGGGCGAAACCCGACAGGACTTAAAGATCCCCACCGTTTCCGGCGGGTCGCTCCCTCTTGCGCTCTCCTGTTCC
GACCCTGCCGTTTACCGGATACCTGTTCCGCCTTTCTCCCTTACGGGAAGTGTGGCGCTTTCTCATAGCTCACACACTGG
TATCTCGGCTCGGTGTAGGTCGTTCGCTCCAAGCTGGGCTGTAAGCAAGAACTCCCCGTTCAGCCCGACTGCTGCGCCTT
ATCCGGTAACTGTTCACTTGAGTCCAACCCGGAAAAGCACGGTAAAACGCCACTGGCAGCAGCCATTGGTAACTGGGAGT
TCGCAGAGGATTTGTTTAGCTAAACACGCGGTTGCTCTTGAAGTGTGCGCCAAAGTCCGGCTACACTGGAAGGACAGATT
TGGTTGCTGTGCTCTGCGAAAGCCAGTTACCACGGTTAAGCAGTTCCCCAACTGACTTAACCTTCGATCAAACCACCTCC
CCAGGTGGTTTTTTCGTTTACAGGGCAAAAGATTACGCGCAGAAAAAAAGGATCTCAAGAAGATCCTTTGATCTTTTCTA
CTGAACCGCTCTAGATTTCAGTGCAATTTATCTCTTCAAATGTAGCACCTGAAGTCAGCCCCATACGATATAAGTTGTAA
TTCTCATGTTAGTCATGCCCCGCGCCCACCGGAAGGAGCTGACTGGGTTGAAGGCTCTCAAGGGCATCGGTCGAGATCCC
GGTGCCTAATGAGTGAGCTAACTTACATTAATTGCGTTGCGCTCACTGCCCGCTTTCCAGTCGGGAAACCTGTCGTGCCA
GCTGCATTAATGAATCGGCCAACGCGCGGGGAGAGGCGGTTTGCGTATTGGGCGCCAGGGTGGTTTTTCTTTTCACCAGT
GAGACGGGCAACAGCTGATTGCCCTTCACCGCCTGGCCCTGAGAGAGTTGCAGCAAGCGGTCCACGCTGGTTTGCCCCAG
CAGGCGAAAATCCTGTTTGATGGTGGTTAACGGCGGGATATAACATGAGCTGTCTTCGGTATCGTCGTATCCCACTACCG
AGATGTCCGCACCAACGCGCAGCCCGGACTCGGTAATGGCGCGCATTGCGCCCAGCGCCATCTGATCGTTGGCAACCAGC
ATCGCAGTGGGAACGATGCCCTCATTCAGCATTTGCATGGTTTGTTGAAAACCGGACATGGCACTCCAGTCGCCTTCCCG
TTCCGCTATCGGCTGAATTTGATTGCGAGTGAGATATTTATGCCAGCCAGCCAGACGCAGACGCGCCGAGACAGAACTTA
ATGGGCCCGCTAACAGCGCGATTTGCTGGTGACCCAATGCGACCAGATGCTCCACGCCCAGTCGCGTACCGTCTTCATGG
GAGAAAATAATACTGTTGATGGGTGTCTGGTCAGAGACATCAAGAAATAACGCCGGAACATTAGTGCAGGCAGCTTCCAC
AGCAATGGCATCCTGGTCATCCAGCGGATAGTTAATGATCAGCCCACTGACGCGTTGCGCGAGAAGATTGTGCACCGCCG
CTTTACAGGCTTCGACGCCGCTTCGTTCTACCATCGACACCACCACGCTGGCACCCAGTTGATCGGCGCGAGATTTAATC
GCCGCGACAATTTGCGACGGCGCGTGCAGGGCCAGACTGGAGGTGGCAACGCCAATCAGCAACGACTGTTTGCCCGCCAG
TTGTTGTGCCACGCGGTTGGGAATGTAATTCAGCTCCGCCATCGCCGCTTCCACTTTTTCCCGCGTTTTCGCAGAAACGT
GGCTGGCCTGGTTCACCACGCGGGAAACGGTCTGATAAGAGACACCGGCATACTCTGCGACATCGTATAACGTTACTGGT
TTCACATTCACCACCCTGAATTGACTCTCTTCCGGGCGCTATCATGCCATACCGCGAAAGGTTTTGCGCCATTCGATGGT
GTCCGGGATCTCGACGCTCTCCCTTATGCGACTCCTGCATTAGGAAATTAATACGACTCACTATA
>pDB070
GGGGAATTGTGAGCGGATAACAATTCCCCTCTAGACCCGGGGGTGATCCGATGCAACTACAATGCCATGCTGGAAGCAGC
GAGTACGTGACACATGTCGGCAAAGCAGCGATGCCAACTCGTTACGGAAATTTTGTTGCCCATGCATTTCGTTCGACAAG
CGATAACAATGAGTATCTCGCGCTGGTGATGGGTGACGTTTGCCAACGTGAATCAGTACTGACGCGACTGCACTCCGAGT
GCCTCACCGGTGATGTACTAGGATCACTGCGCTGCGATTGTGGTGAACAATTGGACGCCGCATTACGCCATATCGCATCT
GAAGGCGTTGGCGCGCTATTGTATTTGCGAGGCCATGAAGGACGCGGCATTGGCTTGTGTAACAAAATTCTTGCATACGG
ACTCCAAGAACAAGGACTTGACACCGTCGATGCCAATCGCGACCTGGGTTTGCCGGATGACGCGCGCGAGTATGACTGCG
CGGCGAGCATCGTGCGCCAACTCGGCATCCTGTCGGTACGGCTGATGAGCAACAATCCGGATAAGTTTGAAGCGCTGCAG
CGCCATGGCATTCCAGTATGTGAGCGCGTTGACCTCGCAATTGCATTACGCGAGGAAAATGAGCGTTATATCTGGACTAA
GCGCAATCGTTTCGGGCATTATTTTGACGAGAGTGAGCTTCAGTGCCCGTTGTCATTGTAGAGGCCTATATGAGATGCAA
TGGTATGAACTGCGGCATCAAAATGAAAGTGATTATTCCGGTTAGCCCGATCAATAGCCTGAAAACCCGTCTGAGCGAAT
TTCTGAGCGGTGAAGAACGTAAAAATCTGCTGCTGAATATGCTGAAGGATATCATTAAAGCACTGGATGGTCTGGATATT
GTTATTGTTAGCCGTGATGAAGAGATCCTGGATTTTGCCAAAAATGAACTGAAAGCCGAAACCATCAAAGAGAAATACAA
AGGCCTGAACAACGCAATCAAACAGGCGTTTGAAGAAATCGAAGATAAAGAGGTGATTATCATTCCGGCAGATATTCCGC
TGATCAAAAAAAAGCACATCGAGGACATTCTGAAGCTGAGCAAAAACTATGATCTGATTATTGCACCGAGCCGTGGTGGT
GGCACCAATCTGCTGTATCTGAAAAGCAAAGATCTGATCGAGATCAAATATGAGGGCTTCAGCTTTCTGAAACATCTGGA
AGAAGCCAAAAAACGCAATCTGCGCTATTATATCTATGATAGCTTCCTGATTAGCGTGGATATCAATACACCGGAAGATC
TGGGTGAAATCTTTATTCATGGCAACGATACCTACACCAAAAACTACCTGAAAAGCCTGGGTATTGATGTGGAACCGAAA
CATTCAAGCGCAGGTCGTTTTGTTGTTAAACGTCGTTAATGCCGACCTTGAGGAATTCGAGTCTTCCATCAAAGACGCCA
AGAGGAGGGTGCTATGCCAAATCTAGTAAGCAATCCTATTTACAAAATGATCCAAGAGGTCGAACGTTCCGCTAAGGGCA
GTCAGCCAAGGCTAGATGACAAGGCTTTGGCGTATGAGCTTGAAGCGGTGCAAGATTGGCGTGAGTTGGCGGGTCTGGCA
TCGGCATGGCGCGATATCGGTTGGGGCAACGTCATCACCTATTCGCGCAAGGTTTTTATTCCGCTCACCCATCTTTGTCG
AGATGTTTGCCACTACTGCACCTTTGCCAAAGCGCCACGGGTGATTGGCCAAGCATTTCTTACCGTGGACCAAGCGCTGG
ACATTGCGCGGGCGGGTGCTGCAGTTGGCTGTCGTGAGGCTTTGTTTACGCTCGGAGATCGGCCCGAAGCGCGTTATGCC
GCTGCACGCGATGCGCTGCAGCAACTGGGTCATGCTACGACAGCCGACTATGTCGCTGAGGTCGCGCAACGCGTACGAGC
TGAAACCGGGCTGCTGCCCCACTTTAATATGGGCGTGCTAAGCGCGGCAGAATACCAGATGCTGCGGCCCCATGCCCCTT
CCTTCGGGTTGATGCTAGAGACGGCTTCTGAACGTCTTTCCGAGCGTGGTGGGCCGCATTACGGCTCTCCCGATAAGCAC
CCGGCAGCGCGGCTTGAAACGCTTCGCCTTGCCGGCGAAGCGCATATACCGATCACCTCAGGCATATTGATTGGTATCGG
GGAGACACGCCGCGAGCGGCTTGAATCGCTCTTTGCGCTGCGGGATTTGCATGATCGCTATGGTCATGTGCAGGAGGTCA

TTATTCAGAACTTTCGCGCGAAACCCGGCACCAAAATGGTTAACGCGCCTGAGCCGTCAATGGACGAATTGTGTTGGACA
ACAGCGGTCGCCCGGTTGGTCCTGGGTTCCGCAATGAGCATCCAGGTGCCGCCTAATCTTTTCGACGGTGACTTGACCGA
TTTGATTCGCGCGGGAATCAATGATTGGGGCGGCGTATCGCCGGTCACGCCTGATCACGTTAATCCAGAAGCGCCGTGGC
CGCACCTTGACCGCTTAAGTGAGGATACAGACAGGGGCGGGAAGACGTTGACCGAGCGGATAACCGTCTATCCTGCCTAT
ATCGAGGAACGTGATCGTTGGATCGACCCGGGCTTGCATGCTGATGTGCTACGCCATTCAGATTGCAGTGGATTGGCAAC
AAGCGATATCTGGAAAGCCGGCTCCACGACGATAGGGGATATCGCTTGCCTGGCAGTCACTCCCACGGCTTCCCAGGTTG
ATCGTGTTGGTTCGCAGACGATCGATATCGTCGAGAAATGCGTGGCGGGTGAGCGCCTCGAAGAAGCCGAGCTTGTACAT
TTATTTAACGCACGCGGCCATGATTTTACGCATGTGACGCACACGGCGGACCGTTTGCGCAGGCAGACAAAGGGCGACAC
GATCACTTACGTTGTCAATCGGAACATCAACTACACCAACGTCTGCCAATACCATTGCAATTTTTGTGCTTTTTCACGAG
GACCGATACAAGAAGACTTGCGCGAAAAGCCCTATATCGTTGACCTCAACGAAATCCGTCGCCGCGTCAAAGAAGCGTGG
GATCGAGGGGCGACGGAGGTCTGCTTGCAAGGGGGCATTCATCCCGACTATACGGGACGTACTTATCTTGACATCTGCGA
AGCGGCTAAAACAGAATGTCCAGACATGCACATTCACGCCTTTTCGCCCCTTGAGGTGCTGCATGGTGCCACGACCCTGG
GGACGTCAATCAGCGCTTTTTTGTCTGATTTGAAAAGCGCTGGATTAAGTACGTTGCCTGGGACCGCAGCCGAGATCCTG
GACGACGAAGTGCGGCGACAAATCTGTCCGGATAAGTTGAATACGCAAGAATGGTTGACCGTCGTGCGATCGGCACATGA
GTTGGGATTGCGGACCACTTGCACGATCATGTTCGGTCATGTCGAATCGTATAAGCATTGGGCGCGACATATCCTCAGGC
TTGCCGAGCTGCAACGTGATACAGGCGGGCTGACTGAGTTTGTTCCATTGCCTTTTGTCCATGAGGAAGCGCCATTATTT
AAAAAGAAAGGTGCGCGTCAGGGGCCGACTTTACGAGAAGCCATTTTGATGCATGCCGTCGGGCGTCTGGCGTTTCATGG
CTTGATTGACAATATTCAAACGTCTTGGGTGAAAATGGGTGAGCACGGCGCACAGCTTTGCTTGCAGGCGGGGGCTAACG
ATTTGGGCGGTACCCTAATGAATGAATCGATTAGCCGCGCAGCCGGTGCAGCCCACGGACAGGAAATGCCGCCTGCCGCC
ATGGAAGCCCTGGCCGCCAGACTCGGGCGCAATGCGATGCAGCGTACCCCGCTTTATCGTAACGTCAGTCCTGAGCGGCA
GCAAGCTTCGCATGCTGCTCCTGCGCTGGTGCCTGTGCGATTCACTAAGGCCGGACGCTTGGTGCGTGGGCACGGCCATC
AGGTCTTGAGTGAAGCGGAACAATGATTGGACCTGCATAGGTGGATCATAGGTGGATTCATGGTGTGGCTTGCATGTTGG
CAGCCTCAAATCGATGTTGTGATGGCTCTCACGCAGCGCTTCCCATCATGGATTCACCAAGGTGCTTAATTACCCTATTA
AATTAGACAGACGCTATTCGGTTTGACACGATCGGAAGGGTGATCGGTTTGGGTCAGAGAAATGCTTGATCTTTACGCAG
GAAGATAGTGATTACCGTTCTGAGCGGTGGTACAGGCACCCCGAAACTGCTGCAGGGTCTGAAACGTGTTGTTAATAATG
AAGAACTGGCCGTGATTGTGAATACCGGTGAAGATACCTGGATTGGTGATCTGTATCTGAGTCCGGATGTTGATACCGTG
CTGTATACCCTGGCAGATCTGATTAACGAAGAAACCTGGTATGGTGTGAAAGAGGATACCTTTTATACCCACGAACAGCT
GAAAAATCTGGGCTTTGATGAAGTTCTGCGCATTGGTGATAAAGATCGTGCACTGAAAATGCACAAAACCTATTATCTGA
AACGCGGTCATAAACTGAGCGAAGTTGTTGATATGGAAAAAGTTGCCCTGGGCATTAAAGCAAAAGTTATTCCGATGACC
GATGATCGTGTGGAAACCAAAATTCTGGCAAAAGTTGATGGTAAAGTGGACCTGCTGAAATTCCATGATTTTTGGGTTAA
ACGCAAAGGTGATGTTGAAGTGCTGGATGTGATTTATGAAAACAGCCTGTATGCAAAACCGTGCGAAAAAGCAGTTGAAG
CCATTAAAAACAGCGATCTGGTTATTATTGGTCCGAGCAATCCGATTACCAGCATCGGTCCGATTCTGAGCCTGAATGGT
ATTAAAGAACTGCTGAAAGACAAAAAAGTTGTTGTGGTTAGCCCGATTGTTGGTAATAGCGCAGTTAGCGGTCCGGCAGG
TAAACTGATGAAAGCCAAAGGTTATGATGTTAGCGTGAAAGGCATCTACGAGTTCTATAAAGATATTGTGGATGTGCTGG
TGATCGACAACGTGGATAAAGAAATTGCAAAAGAAATTCCGTGCGAAGTGCTGATTACCAATACCATCATGAAAACCCTG
GATGATAAAGTTCGTCTGGCCAAAAACATCATTGAATTTTGTGGTAGCCTGTAAGGTTGATGGCTATACAGAAATAAAAA
ATGATCAATGGCATGCGGTACGGCAAACGCTTGCTGAACCACTCACGAAGAATGCACTATCCACTACGCATGCGCCGCCT
TTGGGCTTTTGTTTCTCCCTTGTCATCTTCGAATTGGCGCGGCGACTGGCAGACAATCGCTGCGCTATTTCCCTATTTGA
TGGTTTATAAAGGGCGCGTCGCCTTCGCACTTGCCTGTCTGGTTGCCGCAAAACTCGCTAATCTAGGGGTTCCGGTGGTG
ATGAAAGCGATTGTCGACCAATTATCAGAAATTGACCGGTTGGCAACGCAGGAATCGTCATTTTCCTTGTTCCACGGCGT
GAGTTTTCTTATTGTCGCTTATGCGGTCGTGCGTTTATCAAGCTCGCTTTTTGCTGAGCTGCGTGAAATTATCTTTGGAA
AAGTGGCTTACAACGCTGCGCGACAAGTGGCACTCGCAGTATTCCGGCATCTGCACACGTTGTCGTTGTGTTTTCACCTT
GAGCGGCAAACAGGTGGCCTATCGCGCGATATTGAGCGAGGCACGCGTGGCGTCAAGACGCTTGTTTCATATTCGCTCTA
TAATATCCTGCCAATATGTGTTGAAGTCATATTGGTACTCATTTTTTTTGCTATTCGTTACGACATCTATTATACGGTCG
TCACGCTATCCGTGCTTGCGGCTTACATCACGTTTACCGTGATGGTGACAGAATGGCGTACGCGTTTAAGGCAGGAGATG
AATAAGCTTGATTCACGCTCCAATACGCTAATGGTTGATTCGCTTATTAATTACGAGACAGTCAAGAACTTTGGTAACGA
GCAACACGAAGTACAGCGCTATGATGAAAGCATGATGCTTTATCATGATGCGGCAGTTCGATCGCAGAAATCGCTTTCGT
TCATGAATCTTGGCCAGCAGTCGATTATTGCAATCTGCATGATTGCCGTTTTATGGAGGGCGACGCAGCAGGTTGTGGAT
AAGCAATTGACATTGGGCGACTTTGTGTTAATCAATACATTTATGTTGCAGATATATATTCCGCTGAGTTTTTTGGGGAA
TATGTATCGAACTTTGAAGCAAAGTCTGACCGATATGGATCAGATGTTTTCTTTGTTAAGGCTCAGACGTGAAGTTGACG
ATATTCAAGGTTCATCTCCGCTTGCTGTAAGAAGCGCTGAAGTACGCTTTGAGCATGTCAGTTTTTCATATGAACCGCAA
CGGCAAATTTTGCGGGATGTCACCTTTACGATTGCAGCGGGTACGACGACCGCAATTGTCGGACATAGCGGCTCAGGTAA
GTCGACACTCGCACGGCTGTTGCTTCGCTTTTATGATGTAGAGCATGGCGCTGGTCGTATTTTGATTGATGGCCAAGACA
TTCGGGCTGTGACGCAAGACTCGCTACGCGCGGCCGTTGGCATTGTGCCGCAAGATACGGTGCTTTTCCGTGACACGATC
TATTACAATATTGCGTACGGACGGTTGTCTGCGTCGCCAGAAGAAGTGATAGCGGCTGCGCGCGCCGCTCATATTCATGC
TTTTATTGAAAGCTTGCCCGCTGGCTATTCCACTATAGTGGGTGAGCGTGGTTTGAAATTATCTGGCGGTGAAAAACAGC
GCATCGCAATTGCCCGTACCTTGCTAAAAAAGCCGCCTATTCTCATTTTTGATGAAGCGACTTCAGCACTCGATTCACGT
GCCGAGCGTGCGATTCAGCGCGAGTTGAAACAGCTTGCGCGTCATCGTACAACACTAATCATTGCGCATCGTCTGTCGAC
CATTACGCATGCGCAACAGATTCTTGTTATGGATCAAGGTCGTATCGTTGAGTGTGGCACCCACAGAACATTGCTGAATG
CAGGTGGCCTATATGCGCAAATGTGGGCATTGCAACACAAGCAGCCTGAATAAGTTTAAACCAGCAAGAAGCATGACTGT
ATCCGCCATCGGTGGCATTCCCTTAATACAAACAGGGGATGATCTTGGCCAGATTATTAAGGAGGCGATCCACAAAAACG
GTATCGCTCTAGAAAATGGCGACGTGTTAGTGCTTGCGCAAAAGATCGTTTCAAAGTCCGAAGGACGATGGGCTGCACTA
TCTTCTGTCACGCCCGGCAAACAAGCTATCGATTTGGCGCAGAAAGTGGATAAAGATCCACGGCTCGTTGAACTGATCCT
ATCGGAGTCGGCAGAAATCGTTGCACACAGGCAAGACGGTGTACTGATCACCGCTCACCGTCTCGGATGCGTGATGGCGA

ATGCCGGTATCGATCATTCCAATGTTGGTGATGAAGACAGTGTGCTTCTGCTTCCGAAAGACCCGGATCACAGTGCACGC
GAATTAAAAAAACAGTTTCATCGCTTGTGCGGCGTCGATGTCCACATCATCATCAACGATAGTTTTGGCAGAGTGTGGCG
CCATGGCACGGCAGGATGCGCAATCGGTGTAGCGGGCTTTTCCCCGCTTAAAAATTATATTGGGAAGCCGGATCTGTTTG
GGCAGCTCTTACGAACAACCCAAGTGGCAGTGGCCGATGAACTCGCCGCAGCAGCGTCTTTCTTAATGGGACAAGCCGAT
GAAGGATCGCCTGTTGTGCTGATCCGCGGCGCGAATTTGCCCCCCGCTGATGGAACGGTCCAGCAGCTTATTCGCCCCAA
AGAGGAAGATTTATTTCGTGAAACCTCCTCCACACTGGCGGTGGAAATGGCCTAAACTAGTGGATCCGAATTCGAGCTCG
GCGCGCCTGCAGGTCGACAAGCTTGCGGCCGCATAATGCTTAAGTCGAACAGAAAGTAATCGTATTGTACACGGCCGCAT
AATCGAAATTAATACGACTCACTATAGGGGAATTGTGAGCGGATAACAATTCCCCATCTTAGTATATTAGTTAAGTATAA
GAAGGAGATATACATATGGCAGATCTCAATTGGATATCGGCCGGCCACGCGATCGCTGACGTCGGTACCCTCGAGTCTGG
TAAAGAAACCGCTGCTGCGAAATTTGAACGCCAGCACATGGACTCGTCTACTAGCGCAGCTTAATTAACCTAGGCTGCTG
CCACCGCTGAGCAATAACTAGCATAACCCCTTGGGGCCTCTAAACGGGTCTTGAGGGGTTTTTTGCTGAAAGGAGGAACT
ATATCCGGATTGGCGAATGGGACGCGCCCTGTAGCGGCGCATTAAGCGCGGCGGGTGTGGTGGTTACGCGCAGCGTGACC
GCTACACTTGCCAGCGCCCTAGCGCCCGCTCCTTTCGCTTTCTTCCCTTCCTTTCTCGCCACGTTCGCCGGCTTTCCCCG
TCAAGCTCTAAATCGGGGGCTCCCTTTAGGGTTCCGATTTAGTGCTTTACGGCACCTCGACCCCAAAAAACTTGATTAGG
GTGATGGTTCACGTAGTGGGCCATCGCCCTGATAGACGGTTTTTCGCCCTTTGACGTTGGAGTCCACGTTCTTTAATAGT
GGACTCTTGTTCCAAACTGGAACAACACTCAACCCTATCTCGGTCTATTCTTTTGATTTATAAGGGATTTTGCCGATTTC
GGCCTATTGGTTAAAAAATGAGCTGATTTAACAAAAATTTAACGCGAATTTTAACAAAATATTAACGTTTACAATTTCTG
GCGGCACGATGGCATGAGATTATCAAAAAGGATCTTCACCTAGATCCTTTTAAATTAAAAATGAAGTTTTAAATCAATCT
AAAGTATATATGAGTAAACTTGGTCTGACAGTTACCAATGCTTAATCAGTGAGGCACCTATCTCAGCGATCTGTCTATTT
CGTTCATCCATAGTTGCCTGACTCCCCGTCGTGTAGATAACTACGATACGGGAGGGCTTACCATCTGGCCCCAGTGCTGC
AATGATACCGCGAGACCCACGCTCACCGGCTCCAGATTTATCAGCAATAAACCAGCCAGCCGGAAGGGCCGAGCGCAGAA
GTGGTCCTGCAACTTTATCCGCCTCCATCCAGTCTATTAATTGTTGCCGGGAAGCTAGAGTAAGTAGTTCGCCAGTTAAT
AGTTTGCGCAACGTTGTTGCCATTGCTACAGGCATCGTGGTGTCACGCTCGTCGTTTGGTATGGCTTCATTCAGCTCCGG
TTCCCAACGATCAAGGCGAGTTACATGATCCCCCATGTTGTGCAAAAAAGCGGTTAGCTCCTTCGGTCCTCCGATCGTTG
TCAGAAGTAAGTTGGCCGCAGTGTTATCACTCATGGTTATGGCAGCACTGCATAATTCTCTTACTGTCATGCCATCCGTA
AGATGCTTTTCTGTGACTGGTGAGTACTCAACCAAGTCATTCTGAGAATAGTGTATGCGGCGACCGAGTTGCTCTTGCCC
GGCGTCAATACGGGATAATACCGCGCCACATAGCAGAACTTTAAAAGTGCTCATCATTGGAAAACGTTCTTCGGGGCGAA
AACTCTCAAGGATCTTACCGCTGTTGAGATCCAGTTCGATGTAACCCACTCGTGCACCCAACTGATCTTCAGCATCTTTT
ACTTTCACCAGCGTTTCTGGGTGAGCAAAAACAGGAAGGCAAAATGCCGCAAAAAAGGGAATAAGGGCGACACGGAAATG
TTGAATACTCATACTCTTCCTTTTTCAATCATGATTGAAGCATTTATCAGGGTTATTGTCTCATGAGCGGATACATATTT
GAATGTATTTAGAAAAATAAACAAATAGGTCATGACCAAAATCCCTTAACGTGAGTTTTCGTTCCACTGAGCGTCAGACC
CCGTAGAAAAGATCAAAGGATCTTCTTGAGATCCTTTTTTTCTGCGCGTAATCTGCTGCTTGCAAACAAAAAAACCACCG
CTACCAGCGGTGGTTTGTTTGCCGGATCAAGAGCTACCAACTCTTTTTCCGAAGGTAACTGGCTTCAGCAGAGCGCAGAT
ACCAAATACTGTCCTTCTAGTGTAGCCGTAGTTAGGCCACCACTTCAAGAACTCTGTAGCACCGCCTACATACCTCGCTC
TGCTAATCCTGTTACCAGTGGCTGCTGCCAGTGGCGATAAGTCGTGTCTTACCGGGTTGGACTCAAGACGATAGTTACCG
GATAAGGCGCAGCGGTCGGGCTGAACGGGGGGTTCGTGCACACAGCCCAGCTTGGAGCGAACGACCTACACCGAACTGAG
ATACCTACAGCGTGAGCTATGAGAAAGCGCCACGCTTCCCGAAGGGAGAAAGGCGGACAGGTATCCGGTAAGCGGCAGGG
TCGGAACAGGAGAGCGCACGAGGGAGCTTCCAGGGGGAAACGCCTGGTATCTTTATAGTCCTGTCGGGTTTCGCCACCTC
TGACTTGAGCGTCGATTTTTGTGATGCTCGTCAGGGGGGCGGAGCCTATGGAAAAACGCCAGCAACGCGGCCTTTTTACG
GTTCCTGGCCTTTTGCTGGCCTTTTGCTCACATGTTCTTTCCTGCGTTATCCCCTGATTCTGTGGATAACCGTATTACCG
CCTTTGAGTGAGCTGATACCGCTCGCCGCAGCCGAACGACCGAGCGCAGCGAGTCAGTGAGCGAGGAAGCGGAAGAGCGC
CTGATGCGGTATTTTCTCCTTACGCATCTGTGCGGTATTTCACACCGCATATATGGTGCACTCTCAGTACAATCTGCTCT
GATGCCGCATAGTTAAGCCAGTATACACTCCGCTATCGCTACGTGACTGGGTCATGGCTGCGCCCCGACACCCGCCAACA
CCCGCTGACGCGCCCTGACGGGCTTGTCTGCTCCCGGCATCCGCTTACAGACAAGCTGTGACCGTCTCCGGGAGCTGCAT
GTGTCAGAGGTTTTCACCGTCATCACCGAAACGCGCGAGGCAGCTGCGGTAAAGCTCATCAGCGTGGTCGTGAAGCGATT
CACAGATGTCTGCCTGTTCATCCGCGTCCAGCTCGTTGAGTTTCTCCAGAAGCGTTAATGTCTGGCTTCTGATAAAGCGG
GCCATGTTAAGGGCGGTTTTTTCCTGTTTGGTCACTGATGCCTCCGTGTAAGGGGGATTTCTGTTCATGGGGGTAATGAT
ACCGATGAAACGAGAGAGGATGCTCACGATACGGGTTACTGATGATGAACATGCCCGGTTACTGGAACGTTGTGAGGGTA
AACAACTGGCGGTATGGATGCGGCGGGACCAGAGAAAAATCACTCAGGGTCAATGCCAGCGCTTCGTTAATACAGATGTA
GGTGTTCCACAGGGTAGCCAGCAGCATCCTGCGATGCAGATCCGGAACATAATGGTGCAGGGCGCTGACTTCCGCGTTTC
CAGACTTTACGAAACACGGAAACCGAAGACCATTCATGTTGTTGCTCAGGTCGCAGACGTTTTGCAGCAGCAGTCGCTTC
ACGTTCGCTCGCGTATCGGTGATTCATTCTGCTAACCAGTAAGGCAACCCCGCCAGCCTAGCCGGGTCCTCAACGACAGG
AGCACGATCATGCTAGTCATGCCCCGCGCCCACCGGAAGGAGCTGACTGGGTTGAAGGCTCTCAAGGGCATCGGTCGAGA
TCCCGGTGCCTAATGAGTGAGCTAACTTACATTAATTGCGTTGCGCTCACTGCCCGCTTTCCAGTCGGGAAACCTGTCGT
GCCAGCTGCATTAATGAATCGGCCAACGCGCGGGGAGAGGCGGTTTGCGTATTGGGCGCCAGGGTGGTTTTTCTTTTCAC
CAGTGAGACGGGCAACAGCTGATTGCCCTTCACCGCCTGGCCCTGAGAGAGTTGCAGCAAGCGGTCCACGCTGGTTTGCC
CCAGCAGGCGAAAATCCTGTTTGATGGTGGTTAACGGCGGGATATAACATGAGCTGTCTTCGGTATCGTCGTATCCCACT
ACCGAGATGTCCGCACCAACGCGCAGCCCGGACTCGGTAATGGCGCGCATTGCGCCCAGCGCCATCTGATCGTTGGCAAC
CAGCATCGCAGTGGGAACGATGCCCTCATTCAGCATTTGCATGGTTTGTTGAAAACCGGACATGGCACTCCAGTCGCCTT
CCCGTTCCGCTATCGGCTGAATTTGATTGCGAGTGAGATATTTATGCCAGCCAGCCAGACGCAGACGCGCCGAGACAGAA
CTTAATGGGCCCGCTAACAGCGCGATTTGCTGGTGACCCAATGCGACCAGATGCTCCACGCCCAGTCGCGTACCGTCTTC
ATGGGAGAAAATAATACTGTTGATGGGTGTCTGGTCAGAGACATCAAGAAATAACGCCGGAACATTAGTGCAGGCAGCTT
CCACAGCAATGGCATCCTGGTCATCCAGCGGATAGTTAATGATCAGCCCACTGACGCGTTGCGCGAGAAGATTGTGCACC
GCCGCTTTACAGGCTTCGACGCCGCTTCGTTCTACCATCGACACCACCACGCTGGCACCCAGTTGATCGGCGCGAGATTT

AATCGCCGCGACAATTTGCGACGGCGCGTGCAGGGCCAGACTGGAGGTGGCAACGCCAATCAGCAACGACTGTTTGCCCG
CCAGTTGTTGTGCCACGCGGTTGGGAATGTAATTCAGCTCCGCCATCGCCGCTTCCACTTTTTCCCGCGTTTTCGCAGAA
ACGTGGCTGGCCTGGTTCACCACGCGGGAAACGGTCTGATAAGAGACACCGGCATACTCTGCGACATCGTATAACGTTAC
TGGTTTCACATTCACCACCCTGAATTGACTCTCTTCCGGGCGCTATCATGCCATACCGCGAAAGGTTTTGCGCCATTCGA
TGGTGTCCGGGATCTCGACGCTCTCCCTTATGCGACTCCTGCATTAGGAAGCAGCCCAGTAGTAGGTTGAGGCCGTTGAG
CACCGCCGCCGCAAGGAATGGTGCATGCAAGGAGATGGCGCCCAACAGTCCCCCGGCCACGGGGCCTGCCACCATACCCA
CGCCGAAACAAGCGCTCATGAGCCCGAAGTGGCGAGCCCGATCTTCCCCATCGGTGATGTCGGCGATATAGGCGCCAGCA
ACCGCACCTGTGGCGCCGGTGATGCCGGCCACGATGCGTCCGGCGTAGAGGATCGAGATCGATCTCGATCCCGCGAAATT
AATACGACTCACTATA

>pDB071

TAATACGACTCACTATAGGGGAATTGTGAGCGGATAACAATTCCCCTGTAGAAATAATTTTGTTTAACTTTAATAAGGAG
ATATACCATGCGTGTGGCTCTGCTGGGCGGTACGGGCAACCTGGGCAAAGGTCTGGCACTGCGTCTGGCAACCCTGGGTC
ATGAAATCGTGGTCGGCTCACGTCGCGAAGAAAAAGCGGAAGCCAAAGCGGCCGAATATCGTCGCATTGCAGGCGATGCT
TCGATCACCGGTATGAAAAACGAAGACGCAGCTGAAGCGTGCGATATTGCCGTGCTGACCATCCCGTGGGAACATGCAAT
TGACACGGCTCGTGATCTGAAAAATATTCTGCGCGAAAAAATCGTTGTTAGTCCGCTGGTGCCGGTTTCCCGTGGTGCCA
AAGGTTTTACCTACAGCTCTGAACGCTCAGCGGCCGAAATTGTTGCCGAAGTCCTGGAAAGCGAAAAAGTCGTGTCTGCC
CTGCACACGATCCCGGCAGCTCGTTTTGCAAACCTGGATGAAAAATTCGACTGGGATGTCCCGGTGTGTGGCGATGACGA
TGAAAGCAAAAAAGTTGTCATGTCACTGATTTCGGAAATTGATGGTCTGCGTCCGCTGGATGCCGGTCCGCTGAGTAATT
CCCGCCTGGTTGAATCTCTGACGCCGCTGATTCTGAACATTATGCGTTTTAACGGTATGGGCGAACTGGGTATCAAATTT
CTGTGAGGATCCGAATTCGAGCTCGGCGCGCCTGCAGGTCGACAAGCTTGCGGCCGCATAATGCTTAAGTCGAACAGAAA
GTAATCGTATTGTACACGGCCGCATAATCGAAATTAATACGACTCACTATAGGGGAATTGTGAGCGGATAACAATTCCCC
ATCTTAGTATATTAGGAGATGCAATGGTATGTCTCCTGTTTCTGTCGCCCAGCGCAATACGGGCATCTGGGCGGTCGTGC
CGTTGAAAGCGCCTGAGTGTGCCAAAACCCGGTTGTCTGGCGTTTTAAGCCATGCTGCGCGCCAAGCGTTGTTTTTTTCC
ATGGCCAGTCATGTCATTGGTACGTTACGCGCATCGCCTCGGATTGCTTCCCTATTGGTGGTGACGCCGTCGGAAAGTAC
GGCGGAAATGGCCCGGGCAGCCGGTGCTGAGATCTTGTGGGGACCACCGGACGAAGGCATGGCGAATGCCTGTTCGCGAG
CGATGGCTCATATTGCGGCAGCAGGCGGAGAGCGTGTGATGTTTGTCCCCGGTGATTTGCCCCTGTTGGATGGGGCGGCC
ATCGACATGTTGAGCCGTGCACCGGTCGATGCGATTGGCATGGCGCCAAACCGGGACGGTCATGGCACGAATGGGCTGAT
TTGCCGACCTGGCGCTATTCCGTTGTTTTTCAGCGGGCCAAGCTTTTCCGCTCATCAAAACGCCGCTCGGTGCGCCGGAA
TTGATGTTTGGATTGTCCGTTCAAGAGAGTGGGCGTTGGATGTAGACTTGCCTGCCGACCTTGAGGAATTCGAGTCTTCC
ATCAAAGACGCCAAGAGGAGGGTGCTATGCCAAATCTAGTAAGCAATCCTATTTACAAAATGATCCAAGAGGTCGAACGT
TCCGCTAAGGGCAGTCAGCCAAGGCTAGATGACAAGGCTTTGGCGTATGAGCTTGAAGCGGTGCAAGATTGGCGTGAGTT
GGCGGGTCTGGCATCGGCATGGCGCGATATCGGTTGGGGCAACGTCATCACCTATTCGCGCAAGGTTTTATTCCGCTCA
CCCATCTTTGTCGAGATGTTTGCCACTACTGCACCTTTGCCAAAGCGCCACGGGTGATTGGCCAAGCATTTCTTACCGTG
GACCAAGCGCTGGACATTGCGCGGGCGGGTGCTGCAGTTGGCTGTCGTGAGGCTTTGTTTACGCTCGGAGATCGGCCCGA
AGCGCGTTATGCCGCTGCACGCGATGCGCTGCAGCAACTGGGTCATGCTACGACAGCCGACTATGTCGCTGAGGTCGCGC
AACGCGTACGAGCTGAAACCGGGCTGCTGCCCCACTTTAATATGGGCGTGCTAAGCGCGGCAGAATACCAGATGCTGCGG
CCCCATGCCCCTTCCTTCGGGTTGATGCTAGAGACGGCTTCTGAACGTCTTTCCGAGCGTGGTGGGCCGCATTACGGCTC
TCCCGATAAGCACCCGGCAGCGCGGCTTGAAACGCTTCGCCTTGCCGGCGAAGCGCATATACCGATCACCTCAGGCATAT
TGATTGGTATCGGGGAGACACGCCGCGAGCGGCTTGAATCGCTCTTTGCGCTGCGGGATTTGCATGATCGCTATGGTCAT
GTGCAGGAGGTCATTATTCAGAACTTTCGCGCGAAACCCGGCACCAAAATGGTTAACGCGCCTGAGCCGTCAATGGACGA
ATTGTGTTGGACAACAGCGGTCGCCCGGTTGGTCCTGGGTTCCGCAATGAGCATCCAGGTGCCGCCTAATCTTTTCGACG
GTGACTTGACCGATTTGATTCGCGCGGGAATCAATGATTGGGGCGGCGTATCGCCGGTCACGCCTGATCACGTTAATCCA
GAAGCGCCGTGGCCGCACCTTGACCGCTTAAGTGAGGATACAGACAGGGGCGGGAAGACGTTGACCGAGCGGATAACCGT
CTATCCTGCCTATATCGAGGAACGTGATCGTTGGATCGACCCGGGCTTGCATGCTGATGTGCTACGCCATTCAGATTGCA
GTGGATTGGCAACAAGCGATATCTGGAAAGCCGGCTCCACGACGATAGGGGATATCGCTTGCCTGGCAGTCACTCCCACG
GCTTCCCAGGTTGATCGTGTTGGTTCGCAGACGATCGATATCGTCGAGAAATGCGTGGCGGGTGAGCGCCTCGAAGAAGC
CGAGCTTGTACATTTATTTAACGCACGCGGCCATGATTTTACGCATGTGACGCACACGGCGGACCGTTTGCGCAGGCAGA
CAAAGGGCGACACGATCACTTACGTTGTCAATCGGAACATCAACTACACCAACGTCTGCCAATACCATTGCAATTTTTGT
GCTTTTTCACGAGGACCGATACAAGAAGACTTGCGCGAAAAGCCCTATATCGTTGACCTCAACGAAATCCGTCGCCGCGT
CAAAGAAGCGTGGGATCGAGGGGCGACGGAGGTCTGCTTGCAAGGGGGCATTCATCCCGACTATACGGGACGTACTTATC
TTGACATCTGCGAAGCGGCTAAAACAGAATGTCCAGACATGCACATTCACGCCTTTTCGCCCCTTGAGGTGCTGCATGGT
GCCACGACCCTGGGGACGTCAATCAGCGCTTTTTTGTCTGATTTGAAAAGCGCTGGATTAAGTACGTTGCCTGGGACCGC
AGCCGAGATCCTGGACGACGAAGTGCGGCGACAAATCTGTCCGGATAAGTTGAATACGCAAGAATGGTTGACCGTCGTGC
GATCGGCACATGAGTTGGGATTGCGGACCACTTGCACGATCATGTTCGGTCATGTCGAATCGTATAAGCATTGGGCGCGA
CATATCCTCAGGCTTGCCGAGCTGCAACGTGATACAGGCGGGCTGACTGAGTTTGTTCCATTGCCTTTTGTCCATGAGGA
AGCGCCATTATTTAAAAAGAAAGGTGCGCGTCAGGGGCCGACTTTACGAGAAGCCATTTTGATGCATGCCGTCGGGCGTC
TGGCGTTTCATGGCTTGATTGACAATATTCAAACGTCTTGGGTGAAAATGGGTGAGCACGGCGCACAGCTTTGCTTGCAG
GCGGGGGCTAACGATTTGGGCGGTACCCTAATGAATGAATCGATTAGCCGCGCAGCCGGTGCAGCCCACGGACAGGAAAT
GCCGCCTGCCGCCATGGAAGCCCTGGCCGCCAGACTCGGGCGCAATGCGATGCAGCGTACCCCGCTTTATCGTAACGTCA
GTCCTGAGCGGCAGCAAGCTTCGCATGCTGCTCCTGCGCTGGTGCCTGTGCGATTCACTAAGGCCGGACGCTTGGTGCGT
GGGCACGGCCATCAGGTCTTGAGTGAAGCGGAACAATGATTGGACCTGCATAGGTGGATCATAGGTGGATTCATGGTGTG
GCTTGCATGTTGGCAGCCTCAAATCGATGTTGTGATGGCTCTCACGCAGCGCTTCCCATCATGGATTCACCAAGGTGCTT
AATTACCCTATTAAATTAGACAGACGCTATTCGGTTTGACACGATCGGAAGGGTGATCGGTTTGGGTCAGAGAAATGCTT
GATCTTTACGCAGGAAGATAATGGCAAAGTATGTTGCGTTGTGCGGCGGGGTGGGCGGCGCTAAGCTGGCCTATGGTCTA

GCGCAGGTGCTTTCTGCCGACGAGTTGACGATCGTTGTGAACACGGGCGACGATTTTGAGCATCTTGGCTTGCTCATCTG
TCCAGACCTGGACACGGTTGTCTATACGCTGGCCGATGTGGCAGATGCAAAAAAAGGATGGGGGCGGGCGAACGAGAGCT
GGTCTTCGCAAGAAGCGCTCGCTCGTTTGGGCGGACCGGTTTGGTTCCAGCTGGGTGACAAGGACCTTGCGCTGCATCTT
TATCGGCGCAGCTTGCTTGACGGTGGTGCCAGCTTGTGTGATGTAACCGAGACCATTGCGAAAGCGTTAGGTGTCAAGCA
TTCAATCGTCCCAATGTCTGATGATCCGGTCAGAACCATCGTTGAGACAGACGAAGGCGATTTGCCTTTCCAGACGTATT
TCGTTAAACGGCGATGTGAGCCACGCGTGTCGGGGTTCCGCTTTGAAGGGGCTTCGCTAGCACGTTTATCGTCGCCATTT
GAAGCAGCGTTGAGTACGCCTGACCTGGCTGGCGTAATCTTATGCCCTTCGAATCCTTTTGTCAGCATTGGACCGATTCT
GGCATTACCGGGCGTGCGTGACAGGCTGCATGCGTGTAATGTGCCTGTTTTAGCTGTTGCGCCGCTTGTCGGTGGTGAAG
CAGTCAAAGGCCCGTTGACTAAGATGATGCATGAACTGGGCATGAGTGTTTCGGTTGGCGAGATTGCTTCGCTTTATGCT
GATTTCCTTGATCTGCTGGTGATTGATCCGTTGGATGAATGCGATCATGATCTTTTGCGTAAGGATCGTGTGGCAGTCAA
TAAAGTCAAGACGCTCATGACAACACCTGATGAGCGTATCGCGCTTGCGCGACACGTGCTGGCTTGCGTTGAACATCACC
GAAACAATCGACAGACAGCTCACGTTGTGCAGCACAGCACCGCATAACGAGTCTGGTAAAGAAACCGCTGCTGCGAAATT
TGAACGCCAGCACATGGACTCGTCTACTAGCGCAGCTTAATTAACCTAGGCTGCTGCCACCGCTGAGCAATAACTAGCAT
AACCCCTTGGGGCCTCTAAACGGGTCTTGAGGGGTTTTTTGCTGAAACCTCAGGCATTTGAGAAGCACACGGTCACACTG
CTTCCGGTAGTCAATAAACCGGTAAACCAGCAATAGACATAAGCGGCTATTTAACGACCCTGCCCTGAACCGACGACCGG
GTCATCGTGGCCGGATCTTGCGGCCCCTCGGCTTGAACGAATTGTTAGACATTATTTGCCGACTACCTTGGTGATCTCGC
CTTTCACGTAGTGGACAAATTCTTCCAACTGATCTGCGCGCGAGGCCAAGCGATCTTCTTCTTGTCCAAGATAAGCCTGT
CTAGCTTCAAGTATGACGGGCTGATACTGGGCCGGCAGGCGCTCCATTGCCCAGTCGGCAGCGACATCCTTCGGCGCGAT
TTTGCCGGTTACTGCGCTGTACCAAATGCGGGACAACGTAAGCACTACATTTCGCTCATCGCCAGCCCAGTCGGGCGGCG
AGTTCCATAGCGTTAAGGTTTCATTTAGCGCCTCAAATAGATCCTGTTCAGGAACCGGATCAAAGAGTTCCTCCGCCGCT
GGACCTACCAAGGCAACGCTATGTTCTCTTGCTTTTGTCAGCAAGATAGCCAGATCAATGTCGATCGTGGCTGGCTCGAA
GATACCTGCAAGAATGTCATTGCGCTGCCATTCTCCAAATTGCAGTTCGCGCTTAGCTGGATAACGCCACGGAATGATGT
CGTCGTGCACAACAATGGTGACTTCTACAGCGCGGAGAATCTCGCTCTCTCCAGGGGAAGCCGAAGTTTCCAAAAGGTCG
TTGATCAAAGCTCGCCGCGTTGTTTCATCAAGCCTTACGGTCACCGTAACCAGCAAATCAATATCACTGTGTGGCTTCAG
GCCGCCATCCACTGCGGAGCCGTACAAATGTACGGCCAGCAACGTCGGTTCGAGATGGCGCTCGATGACGCCAACTACCT
CTGATAGTTGAGTCGATACTTCGGCGATCACCGCTTCCCTCATACTCTTCCTTTTTCAATATTATTGAAGCATTTATCAG
GGTTATTGTCTCATGAGCGGATACATATTTGAATGTATTTAGAAAAATAAACAAATAGCTAGCTCACTCGGTCGCTACGC
TCCGGGCGTGAGACTGCGGCGGGCGCTGCGGACACATACAAAGTTACCCACAGATTCCGTGGATAAGCAGGGGACTAACA
TGTGAGGCAAAACAGCAGGGCCGCGCCGGTGGCGTTTTTCCATAGGCTCCGCCCTCCTGCCAGAGTTCACATAAACAGAC
GCTTTTCCGGTGCATCTGTGGGAGCCGTGAGGCTCAACCATGAATCTGACAGTACGGGCGAAACCCGACAGGACTTAAAG
ATCCCCACCGTTTCCGGCGGGTCGCTCCCTCTTGCGCTCTCCTGTTCCGACCCTGCCGTTTACCGGATACCTGTTCCGCC
TTTCTCCCTTACGGGAAGTGTGGCGCTTTCTCATAGCTCACACACTGGTATCTCGGCTCGGTGTAGGTCGTTCGCTCCAA
GCTGGGCTGTAAGCAAGAACTCCCCGTTCAGCCCGACTGCTGCGCCTTATCCGGTAACTGTTCACTTGAGTCCAACCCGG
AAAAGCACGGTAAAACGCCACTGGCAGCAGCCATTGGTAACTGGGAGTTCGCAGAGGATTTGTTTAGCTAAACACGCGGT
TGCTCTTGAAGTGTGCGCCAAAGTCCGGCTACACTGGAAGGACAGATTTGGTTGCTGTGCTCTGCGAAAGCCAGTTACCA
CGGTTAAGCAGTTCCCCAACTGACTTAACCTTCGATCAAACCACCTCCCCAGGTGGTTTTTTCGTTTACAGGGCAAAAGA
TTACGCGCAGAAAAAAAGGATCTCAAGAAGATCCTTTGATCTTTTCTACTGAACCGCTCTAGATTTCAGTGCAATTTATC
TCTTCAAATGTAGCACCTGAAGTCAGCCCCATACGATATAAGTTGTAATTCTCATGTTAGTCATGCCCCGCGCCCACCGG
AAGGAGCTGACTGGGTTGAAGGCTCTCAAGGGCATCGGTCGAGATCCCGGTGCCTAATGAGTGAGCTAACTTACATTAAT
TGCGTTGCGCTCACTGCCCGCTTTCCAGTCGGGAAACCTGTCGTGCCAGCTGCATTAATGAATCGGCCAACGCGCGGGGA
GAGGCGGTTTGCGTATTGGGCGCCAGGGTGGTTTTTCTTTTCACCAGTGAGACGGGCAACAGCTGATTGCCCTTCACCGC
CTGGCCCTGAGAGAGTTGCAGCAAGCGGTCCACGCTGGTTTGCCCCAGCAGGCGAAAATCCTGTTTGATGGTGGTTAACG
GCGGGATATAACATGAGCTGTCTTCGGTATCGTCGTATCCCACTACCGAGATGTCCGCACCAACGCGCAGCCCGGACTCG
GTAATGGCGCGCATTGCGCCCAGCGCCATCTGATCGTTGGCAACCAGCATCGCAGTGGGAACGATGCCCTCATTCAGCAT
TTGCATGGTTTGTTGAAAACCGGACATGGCACTCCAGTCGCCTTCCCGTTCCGCTATCGGCTGAATTTGATTGCGAGTGA
GATATTTATGCCAGCCAGCCAGACGCAGACGCGCCGAGACAGAACTTAATGGGCCCGCTAACAGCGCGATTTGCTGGTGA
CCCAATGCGACCAGATGCTCCACGCCCAGTCGCGTACCGTCTTCATGGGAGAAAATAATACTGTTGATGGGTGTCTGGTC
AGAGACATCAAGAAATAACGCCGGAACATTAGTGCAGGCAGCTTCCACAGCAATGGCATCCTGGTCATCCAGCGGATAGT
TAATGATCAGCCCACTGACGCGTTGCGCGAGAAGATTGTGCACCGCCGCTTTACAGGCTTCGACGCCGCTTCGTTCTACC
ATCGACACCACCACGCTGGCACCCAGTTGATCGGCGCGAGATTTAATCGCCGCGACAATTTGCGACGGCGCGTGCAGGGC
CAGACTGGAGGTGGCAACGCCAATCAGCAACGACTGTTTGCCCGCCAGTTGTTGTGCCACGCGGTTGGGAATGTAATTCA
GCTCCGCCATCGCCGCTTCCACTTTTTCCCGCGTTTTCGCAGAAACGTGGCTGGCCTGGTTCACCACGCGGGAAACGGTC
TGATAAGAGACACCGGCATACTCTGCGACATCGTATAACGTTACTGGTTTCACATTCACCACCCTGAATTGACTCTCTTC
CGGGCGCTATCATGCCATACCGCGAAAGGTTTTGCGCCATTCGATGGTGTCCGGGATCTCGACGCTCTCCCTTATGCGAC
TCCTGCATTAGGAAAT
>pDL008
GCGCCAGCAACCGCACCTGTGGCGCCGGTGATGCCGGCCACGATGCGTCCGGCGTAGAGGATCGAGATCTCGATCCCGCG
AAATTAATACGACTCACTATAGGGGAATTGTGAGCGGATAACAATTCCCCTCTAGAAATAATTTTGTTTAACTTTAAGAA
GGAGATATACCATGCGTGTGGCTCTGCTGGGCGGTACGGGCAACCTGGGCAAAGGTCTGGCACTGCGTCTGGCAACCCTG
GGTCATGAAATCGTGGTCGGCTCACGTCGCGAAGAAAAAGCGGAAGCCAAAGCGGCCGAATATCGTCGCATTGCAGGCGA
TGCTTCGATCACCGGTATGAAAAACGAAGACGCAGCTGAAGCGTGCGATATTGCCGTGCTGACCATCCCGTGGGAACATG
CAATTGACACGGCTCGTGATCTGAAAAATATTCTGCGCGAAAAAATCGTTGTTAGTCCGCTGGTGCCGGTTTCCCGTGGT
GCCAAAGGTTTTACCTACAGCTCTGAACGCTCAGCGGCCGAAATTGTTGCCGAAGTCCTGGAAAGCGAAAAAGTCGTGTC
TGCCCTGCACACGATCCCGGCAGCTCGTTTTGCAAACCTGGATGAAAAATTCGACTGGGATGTCCCGGTGTGTGGCGATG

ACGATGAAAGCAAAAAAGTTGTCATGTCACTGATTTCGGAAATTGATGGTCTGCGTCCGCTGGATGCCGGTCCGCTGAGT
AATTCCCGCCTGGTTGAATCTCTGACGCCGCTGATTCTGAACATTATGCGTTTTAACGGTATGGGCGAACTGGGTATCAA
ATTTCTGGCGGCCGCACTCGAGCACCACCACCACCACCACTGAGATCCGGCTGCTAACAAAGCCCGAAAGGAAGCTGAGT
TGGCTGCTGCCACCGCTGAGCAATAACTAGCATAACCCCTTGGGGCCTCTAAACGGGTCTTGAGGGGTTTTTTGCTGAAA
GGAGGAACTATATCCGGATTGGCGAATGGGACGCGCCCTGTAGCGGCGCATTAAGCGCGGCGGGTGTGGTGGTTACGCGC
AGCGTGACCGCTACACTTGCCAGCGCCCTAGCGCCCGCTCCTTTCGCTTTCTTCCCTTCCTTTCTCGCCACGTTCGCCGG
CTTTCCCCGTCAAGCTCTAAATCGGGGGCTCCCTTTAGGGTTCCGATTTAGTGCTTTACGGCACCTCGACCCCAAAAAAC
TTGATTAGGGTGATGGTTCACGTAGTGGGCCATCGCCCTGATAGACGGTTTTTCGCCCTTTGACGTTGGAGTCCACGTTC
TTTAATAGTGGACTCTTGTTCCAAACTGGAACAACACTCAACCCTATCTCGGTCTATTCTTTTGATTTATAAGGGATTTT
GCCGATTTCGGCCTATTGGTTAAAAAATGAGCTGATTTAACAAAAATTTAACGCGAATTTTAACAAAATATTAACGTTTA
CAATTTCAGGTGGCACTTTTCGGGGAAATGTGCGCGGAACCCCTATTTGTTTATTTTTCTAAATACATTCAAATATGTAT
CCGCTCATGAATTAATTCTTAGAAAAACTCATCGAGCATCAAATGAAACTGCAATTTATTCATATCAGGATTATCAATAC
CATATTTTTGAAAAAGCCGTTTCTGTAATGAAGGAGAAAACTCACCGAGGCAGTTCCATAGGATGGCAAGATCCTGGTAT
CGGTCTGCGATTCCGACTCGTCCAACATCAATACAACCTATTAATTTCCCCTCGTCAAAAATAAGGTTATCAAGTGAGAA
ATCACCATGAGTGACGACTGAATCCGGTGAGAATGGCAAAAGTTTATGCATTTCTTTCCAGACTTGTTCAACAGGCCAGC
CATTACGCTCGTCATCAAAATCACTCGCATCAACCAAACCGTTATTCATTCGTGATTGCGCCTGAGCGAGACGAAATACG
CGATCGCTGTTAAAAGGACAATTACAAACAGGAATCGAATGCAACCGGCGCAGGAACACTGCCAGCGCATCAACAATATT
TTCACCTGAATCAGGATATTCTTCTAATACCTGGAATGCTGTTTTCCCGGGGATCGCAGTGGTGAGTAACCATGCATCAT
CAGGAGTACGGATAAAATGCTTGATGGTCGGAAGAGGCATAAATTCCGTCAGCCAGTTTAGTCTGACCATCTCATCTGTA
ACATCATTGGCAACGCTACCTTTGCCATGTTTCAGAAACAACTCTGGCGCATCGGGCTTCCCATACAATCGATAGATTGT
CGCACCTGATTGCCCGACATTATCGCGAGCCCATTTATACCCATATAAATCAGCATCCATGTTGGAATTTAATCGCGGCC
TAGAGCAAGACGTTTCCCGTTGAATATGGCTCATAACACCCCTTGTATTACTGTTTATGTAAGCAGACAGTTTTATTGTT
CATGACCAAAATCCCTTAACGTGAGTTTTCGTTCCACTGAGCGTCAGACCCCGTAGAAAAGATCAAAGGATCTTCTTGAG
ATCCTTTTTTTCTGCGCGTAATCTGCTGCTTGCAAACAAAAAAACCACCGCTACCAGCGGTGGTTTGTTTGCCGGATCAA
GAGCTACCAACTCTTTTTCCGAAGGTAACTGGCTTCAGCAGAGCGCAGATACCAAATACTGTCCTTCTAGTGTAGCCGTA
GTTAGGCCACCACTTCAAGAACTCTGTAGCACCGCCTACATACCTCGCTCTGCTAATCCTGTTACCAGTGGCTGCTGCCA
GTGGCGATAAGTCGTGTCTTACCGGGTTGGACTCAAGACGATAGTTACCGGATAAGGCGCAGCGGTCGGGCTGAACGGGG
GGTTCGTGCACACAGCCCAGCTTGGAGCGAACGACCTACACCGAACTGAGATACCTACAGCGTGAGCTATGAGAAAGCGC
CACGCTTCCCGAAGGGAGAAAGGCGGACAGGTATCCGGTAAGCGGCAGGGTCGGAACAGGAGAGCGCACGAGGGAGCTTC
CAGGGGGAAACGCCTGGTATCTTTATAGTCCTGTCGGGTTTCGCCACCTCTGACTTGAGCGTCGATTTTTGTGATGCTCG
TCAGGGGGGCGGAGCCTATGGAAAAACGCCAGCAACGCGGCCTTTTTACGGTTCCTGGCCTTTTGCTGGCCTTTTGCTCA
CATGTTCTTTCCTGCGTTATCCCCTGATTCTGTGGATAACCGTATTACCGCCTTTGAGTGAGCTGATACCGCTCGCCGCA
GCCGAACGACCGAGCGCAGCGAGTCAGTGAGCGAGGAAGCGGAAGAGCGCCTGATGCGGTATTTTCTCCTTACGCATCTG
TGCGGTATTTCACACCGCATATATGGTGCACTCTCAGTACAATCTGCTCTGATGCCGCATAGTTAAGCCAGTATACACTC
CGCTATCGCTACGTGACTGGGTCATGGCTGCGCCCCGACACCCGCCAACACCCGCTGACGCGCCCTGACGGGCTTGTCTG
CTCCCGGCATCCGCTTACAGACAAGCTGTGACCGTCTCCGGGAGCTGCATGTGTCAGAGGTTTTCACCGTCATCACCGAA
ACGCGCGAGGCAGCTGCGGTAAAGCTCATCAGCGTGGTCGTGAAGCGATTCACAGATGTCTGCCTGTTCATCCGCGTCCA
GCTCGTTGAGTTTCTCCAGAAGCGTTAATGTCTGGCTTCTGATAAAGCGGGCCATGTTAAGGGCGGTTTTTTCCTGTTTG
GTCACTGATGCCTCCGTGTAAGGGGGATTTCTGTTCATGGGGGTAATGATACCGATGAAACGAGAGAGGATGCTCACGAT
ACGGGTTACTGATGATGAACATGCCCGGTTACTGGAACGTTGTGAGGGTAAACAACTGGCGGTATGGATGCGGCGGGACC
AGAGAAAAATCACTCAGGGTCAATGCCAGCGCTTCGTTAATACAGATGTAGGTGTTCCACAGGGTAGCCAGCAGCATCCT
GCGATGCAGATCCGGAACATAATGGTGCAGGGCGCTGACTTCCGCGTTTCCAGACTTTACGAAACACGGAAACCGAAGAC
CATTCATGTTGTTGCTCAGGTCGCAGACGTTTTGCAGCAGCAGTCGCTTCACGTTCGCTCGCGTATCGGTGATTCATTCT
GCTAACCAGTAAGGCAACCCCGCCAGCCTAGCCGGGTCCTCAACGACAGGAGCACGATCATGCGCACCCGTGGGGCCGCC
ATGCCGGCGATAATGGCCTGCTTCTCGCCGAAACGTTTGGTGGCGGGACCAGTGACGAAGGCTTGAGCGAGGGCGTGCAA
GATTCCGAATACCGCAAGCGACAGGCCGATCATCGTCGCGCTCCAGCGAAAGCGGTCCTCGCCGAAAATGACCCAGAGCG
CTGCCGGCACCTGTCCTACGAGTTGCATGATAAAGAAGACAGTCATAAGTGCGGCGACGATAGTCATGCCCCGCGCCCAC
CGGAAGGAGCTGACTGGGTTGAAGGCTCTCAAGGGCATCGGTCGAGATCCCGGTGCCTAATGAGTGAGCTAACTTACATT
AATTGCGTTGCGCTCACTGCCCGCTTTCCAGTCGGGAAACCTGTCGTGCCAGCTGCATTAATGAATCGGCCAACGCGCGG
GGAGAGGCGGTTTGCGTATTGGGCGCCAGGGTGGTTTTTCTTTTCACCAGTGAGACGGGCAACAGCTGATTGCCCTTCAC
CGCCTGGCCCTGAGAGAGTTGCAGCAAGCGGTCCACGCTGGTTTGCCCCAGCAGGCGAAAATCCTGTTTGATGGTGGTTA
ACGGCGGGATATAACATGAGCTGTCTTCGGTATCGTCGTATCCCACTACCGAGATATCCGCACCAACGCGCAGCCCGGAC
TCGGTAATGGCGCGCATTGCGCCCAGCGCCATCTGATCGTTGGCAACCAGCATCGCAGTGGGAACGATGCCCTCATTCAG
CATTTGCATGGTTTGTTGAAAACCGGACATGGCACTCCAGTCGCCTTCCCGTTCCGCTATCGGCTGAATTTGATTGCGAG
TGAGATATTTATGCCAGCCAGCCAGACGCAGACGCGCCGAGACAGAACTTAATGGGCCCGCTAACAGCGCGATTTGCTGG
TGACCCAATGCGACCAGATGCTCCACGCCCAGTCGCGTACCGTCTTCATGGGAGAAAATAATACTGTTGATGGGTGTCTG
GTCAGAGACATCAAGAAATAACGCCGGAACATTAGTGCAGGCAGCTTCCACAGCAATGGCATCCTGGTCATCCAGCGGAT
AGTTAATGATCAGCCCACTGACGCGTTGCGCGAGAAGATTGTGCACCGCCGCTTTACAGGCTTCGACGCCGCTTCGTTCT
ACCATCGACACCACCACGCTGGCACCCAGTTGATCGGCGCGAGATTTAATCGCCGCGACAATTTGCGACGGCGCGTGCAG
GGCCAGACTGGAGGTGGCAACGCCAATCAGCAACGACTGTTTGCCCGCCAGTTGTTGTGCCACGCGGTTGGGAATGTAAT
TCAGCTCCGCCATCGCCGCTTCCACTTTTTCCCGCGTTTTCGCAGAAACGTGGCTGGCCTGGTTCACCACGCGGGAAACG
GTCTGATAAGAGACACCGGCATACTCTGCGACATCGTATAACGTTACTGGTTTCACATTCACCACCCTGAATTGACTCTC
TTCCGGGCGCTATCATGCCATACCGCGAAAGGTTTTGCGCCATTCGATGGTGTCCGGGATCTCGACGCTCTCCCTTATGC
GACTCCTGCATTAGGAAGCAGCCCAGTAGTAGGTTGAGGCCGTTGAGCACCGCCGCCGCAAGGAATGGTGCATGCAAGGA

GATGGCGCCCAACAGTCCCCCGGCCACGGGGCCTGCCACCATACCCACGCCGAAACAAGCGCTCATGAGCCCGAAGTGGC
GAGCCCGATCTTCCCCATCGGTGATGTCGGCGATATAG
>pMH01
TGGCGAATGGGACGCGCCCTGTAGCGGCGCATTAAGCGCGGCGGGTGTGGTGGTTACGCGCAGCGTGACCGCTACACTTG
CCAGCGCCCTAGCGCCCGCTCCTTTCGCTTTCTTCCCTTCCTTTCTCGCCACGTTCGCCGGCTTTCCCCGTCAAGCTCTA
AATCGGGGGCTCCCTTTAGGGTTCCGATTTAGTGCTTTACGGCACCTCGACCCCAAAAAACTTGATTAGGGTGATGGTTC
ACGTAGTGGGCCATCGCCCTGATAGACGGTTTTTCGCCCTTTGACGTTGGAGTCCACGTTCTTTAATAGTGGACTCTTGT
TCCAAACTGGAACAACACTCAACCCTATCTCGGTCTATTCTTTTGATTTATAAGGGATTTTGCCGATTTCGGCCTATTGG
TTAAAAAATGAGCTGATTTAACAAAAATTTAACGCGAATTTTAACAAAATATTAACGTTTACAATTTCAGGTGGCACTTT
TCGGGGAAATGTGCGCGGAACCCCTATTTGTTTATTTTTCTAAATACATTCAAATATGTATCCGCTCATGAATTAATTCT
TAGAAAAACTCATCGAGCATCAAATGAAACTGCAATTTATTCATATCAGGATTATCAATACCATATTTTTGAAAAAGCCG
TTTCTGTAATGAAGGAGAAAACTCACCGAGGCAGTTCCATAGGATGGCAAGATCCTGGTATCGGTCTGCGATTCCGACTC
GTCCAACATCAATACAACCTATTAATTTCCCCTCGTCAAAAATAAGGTTATCAAGTGAGAAATCACCATGAGTGACGACT
GAATCCGGTGAGAATGGCAAAAGTTTATGCATTTCTTTCCAGACTTGTTCAACAGGCCAGCCATTACGCTCGTCATCAAA
ATCACTCGCATCAACCAAACCGTTATTCATTCGTGATTGCGCCTGAGCGAGACGAAATACGCGATCGCTGTTAAAAGGAC
AATTACAAACAGGAATCGAATGCAACCGGCGCAGGAACACTGCCAGCGCATCAACAATATTTTCACCTGAATCAGGATAT
TCTTCTAATACCTGGAATGCTGTTTTCCCGGGGATCGCAGTGGTGAGTAACCATGCATCATCAGGAGTACGGATAAAATG
CTTGATGGTCGGAAGAGGCATAAATTCCGTCAGCCAGTTTAGTCTGACCATCTCATCTGTAACATCATTGGCAACGCTAC
CTTTGCCATGTTTCAGAAACAACTCTGGCGCATCGGGCTTCCCATACAATCGATAGATTGTCGCACCTGATTGCCCGACA
TTATCGCGAGCCCATTTATACCCATATAAATCAGCATCCATGTTGGAATTTAATCGCGGCCTAGAGCAAGACGTTTCCCG
TTGAATATGGCTCATAACACCCCTTGTATTACTGTTTATGTAAGCAGACAGTTTTATTGTTCATGACCAAAATCCCTTAA
CGTGAGTTTTCGTTCCACTGAGCGTCAGACCCCGTAGAAAAGATCAAAGGATCTTCTTGAGATCCTTTTTTTCTGCGCGT
AATCTGCTGCTTGCAAACAAAAAAACCACCGCTACCAGCGGTGGTTTGTTTGCCGGATCAAGAGCTACCAACTCTTTTTC
CGAAGGTAACTGGCTTCAGCAGAGCGCAGATACCAAATACTGTCCTTCTAGTGTAGCCGTAGTTAGGCCACCACTTCAAG
AACTCTGTAGCACCGCCTACATACCTCGCTCTGCTAATCCTGTTACCAGTGGCTGCTGCCAGTGGCGATAAGTCGTGTCT
TACCGGGTTGGACTCAAGACGATAGTTACCGGATAAGGCGCAGCGGTCGGGCTGAACGGGGGGTTCGTGCACACAGCCCA
GCTTGGAGCGAACGACCTACACCGAACTGAGATACCTACAGCGTGAGCTATGAGAAAGCGCCACGCTTCCCGAAGGGAGA
AAGGCGGACAGGTATCCGGTAAGCGGCAGGGTCGGAACAGGAGAGCGCACGAGGGAGCTTCCAGGGGGAAACGCCTGGTA
TCTTTATAGTCCTGTCGGGTTTCGCCACCTCTGACTTGAGCGTCGATTTTTGTGATGCTCGTCAGGGGGGCGGAGCCTAT
GGAAAAACGCCAGCAACGCGGCCTTTTTACGGTTCCTGGCCTTTTGCTGGCCTTTTGCTCACATGTTCTTTCCTGCGTTA
TCCCCTGATTCTGTGGATAACCGTATTACCGCCTTTGAGTGAGCTGATACCGCTCGCCGCAGCCGAACGACCGAGCGCAG
CGAGTCAGTGAGCGAGGAAGCGGAAGAGCGCCTGATGCGGTATTTTCTCCTTACGCATCTGTGCGGTATTTCACACCGCA
TATATGGTGCACTCTCAGTACAATCTGCTCTGATGCCGCATAGTTAAGCCAGTATACACTCCGCTATCGCTACGTGACTG
GGTCATGGCTGCGCCCCGACACCCGCCAACACCCGCTGACGCGCCCTGACGGGCTTGTCTGCTCCCGGCATCCGCTTACA
GACAAGCTGTGACCGTCTCCGGGAGCTGCATGTGTCAGAGGTTTTCACCGTCATCACCGAAACGCGCGAGGCAGCTGCGG
TAAAGCTCATCAGCGTGGTCGTGAAGCGATTCACAGATGTCTGCCTGTTCATCCGCGTCCAGCTCGTTGAGTTTCTCCAG
AAGCGTTAATGTCTGGCTTCTGATAAAGCGGGCCATGTTAAGGGCGGTTTTTTCCTGTTTGGTCACTGATGCCTCCGTGT
AAGGGGGATTTCTGTTCATGGGGGTAATGATACCGATGAAACGAGAGAGGATGCTCACGATACGGGTTACTGATGATGAA
CATGCCCGGTTACTGGAACGTTGTGAGGGTAAACAACTGGCGGTATGGATGCGGCGGGACCAGAGAAAAATCACTCAGGG
TCAATGCCAGCGCTTCGTTAATACAGATGTAGGTGTTCCACAGGGTAGCCAGCAGCATCCTGCGATGCAGATCCGGAACA
TAATGGTGCAGGGCGCTGACTTCCGCGTTTCCAGACTTTACGAAACACGGAAACCGAAGACCATTCATGTTGTTGCTCAG
GTCGCAGACGTTTTGCAGCAGCAGTCGCTTCACGTTCGCTCGCGTATCGGTGATTCATTCTGCTAACCAGTAAGGCAACC
CCGCCAGCCTAGCCGGGTCCTCAACGACAGGAGCACGATCATGCGCACCCGTGGGGCCGCCATGCCGGCGATAATGGCCT
GCTTCTCGCCGAAACGTTTGGTGGCGGGACCAGTGACGAAGGCTTGAGCGAGGGCGTGCAAGATTCCGAATACCGCAAGC
GACAGGCCGATCATCGTCGCGCTCCAGCGAAAGCGGTCCTCGCCGAAAATGACCCAGAGCGCTGCCGGCACCTGTCCTAC
GAGTTGCATGATAAAGAAGACAGTCATAAGTGCGGCGACGATAGTCATGCCCCGCGCCCACCGGAAGGAGCTGACTGGGT
TGAAGGCTCTCAAGGGCATCGGTCGAGATCCCGGTGCCTAATGAGTGAGCTAACTTACATTAATTGCGTTGCGCTCACTG
CCCGCTTTCCAGTCGGGAAACCTGTCGTGCCAGCTGCATTAATGAATCGGCCAACGCGCGGGGAGAGGCGGTTTGCGTAT
TGGGCGCCAGGGTGGTTTTTCTTTTCACCAGTGAGACGGGCAACAGCTGATTGCCCTTCACCGCCTGGCCCTGAGAGAGT
TGCAGCAAGCGGTCCACGCTGGTTTGCCCCAGCAGGCGAAAATCCTGTTTGATGGTGGTTAACGGCGGGATATAACATGA
GCTGTCTTCGGTATCGTCGTATCCCACTACCGAGATATCCGCACCAACGCGCAGCCCGGACTCGGTAATGGCGCGCATTG
CGCCCAGCGCCATCTGATCGTTGGCAACCAGCATCGCAGTGGGAACGATGCCCTCATTCAGCATTTGCATGGTTTGTTGA
AAACCGGACATGGCACTCCAGTCGCCTTCCCGTTCCGCTATCGGCTGAATTTGATTGCGAGTGAGATATTTATGCCAGCC
AGCCAGACGCAGACGCGCCGAGACAGAACTTAATGGGCCCGCTAACAGCGCGATTTGCTGGTGACCCAATGCGACCAGAT
GCTCCACGCCCAGTCGCGTACCGTCTTCATGGGAGAAAATAATACTGTTGATGGGTGTCTGGTCAGAGACATCAAGAAAT
AACGCCGGAACATTAGTGCAGGCAGCTTCCACAGCAATGGCATCCTGGTCATCCAGCGGATAGTTAATGATCAGCCCACT
GACGCGTTGCGCGAGAAGATTGTGCACCGCCGCTTTACAGGCTTCGACGCCGCTTCGTTCTACCATCGACACCACCACGC
TGGCACCCAGTTGATCGGCGCGAGATTTAATCGCCGCGACAATTTGCGACGGCGCGTGCAGGGCCAGACTGGAGGTGGCA
ACGCCAATCAGCAACGACTGTTTGCCCGCCAGTTGTTGTGCCACGCGGTTGGGAATGTAATTCAGCTCCGCCATCGCCGC
TTCCACTTTTTCCCGCGTTTTCGCAGAAACGTGGCTGGCCTGGTTCACCACGCGGGAAACGGTCTGATAAGAGACACCGG
CATACTCTGCGACATCGTATAACGTTACTGGTTTCACATTCACCACCCTGAATTGACTCTCTTCCGGGCGCTATCATGCC
ATACCGCGAAAGGTTTTGCGCCATTCGATGGTGTCCGGGATCTCGACGCTCTCCCTTATGCGACTCCTGCATTAGGAAGC
AGCCCAGTAGTAGGTTGAGGCCGTTGAGCACCGCCGCCGCAAGGAATGGTGCATGCAAGGAGATGGCGCCCAACAGTCCC
CCGGCCACGGGGCCTGCCACCATACCCACGCCGAAACAAGCGCTCATGAGCCCGAAGTGGCGAGCCCGATCTTCCCCATC

GGTGATGTCGGCGATATAGGCGCCAGCAACCGCACCTGTGGCGCCGGTGATGCCGGCCACGATGCGTCCGGCGTAGAGGA
TCGAGATCTCGATCCCGCGAAATTAATACGACTCACTATAGGGGAATTGTGAGCGGATAACAATTCCCCTCTAGAAATAA
TTTTGTTTAACTTTAAGAAGGAGATATACCATGGGCAGCAGCCATCATCATCATCATCACAGCAGCGGCCTGGTGCCGCG
CGGCAGCCATATGGCTAGCATGACTGGTGGACAGCAAATGGGTCGCGGATCCATGATCAAGGAAAAGCGCAAAGTTGAAG
TGATTGGTCTGGAACTGCCGATTTTTAAAGGCGGCGAACAGATTAATCTGAGTGAACTGATTGCCCAGTATCCGATTGAA
GATGGTGACATTATTGTTATTGCAGAAACCCTGATTAGCAAACTGGAAGGTGGTGTTATTGATCGTGATAAAATTATTCC
GAGCAAAGAAGCCATTGAACTGGCCAAAAAGACTGGTAAAGATCCGAAAGTTGTTCAGGTTATTCTGGATGAAGCCAAAG
AAATTGTGAAAGTTGGCAAAAATTTCATCATCACCGAAACCAAACATGGCTTTGTTTGCGCCAATAGCGGTGTGGATGAA
AGCAATATCTATAAAGGTATTAAGATCCTGCCGAAAAATCCGGATGAAAGTGCCGAAAAAATTCGCAAAGAAATTGAAAA
ACTGACCGGCAAACGCGTTGGCGTTATTATTAGTGATAGCGTGGGTCGCCCGTTTCGCAAAGGCGCAGTTGGTATTGCCA
TTGGCGTTAGCGGCATTCTGGCACTGTGGGATCGCAAAGGTGAAAAAGATCTGTTTGGTCGTGAACTGAAAACCACCGAA
GTGGCCATTGCCGATGAACTGGCCAGCATGGCCAATGTGGTTATGGGCGAAGCCGATGAAGGCATTCCGGTTGTTATTAT
TCGTGGCGCAAATGTGCCGTTTGGCAATGGCAAAGGCCGTGATCTGATTCGTCCGAAAGAAGAAGATGTGTTTCGCAATT
AAAAGCTTGCGGCCGCACTCGAGCACCACCACCACCACCACTGAGATCCGGCTGCTAACAAAGCCCGAAAGGAAGCTGAG
TTGGCTGCTGCCACCGCTGAGCAATAACTAGCATAACCCCTTGGGGCCTCTAAACGGGTCTTGAGGGGTTTTTTGCTGAA
AGGAGGAACTATATCCGGAT

>pMH02
GGGGAATTGTGAGCGGATAACAATTCCCCTGTAGAAATAATTTTGTTTAACTTTAATAAGGAGATATACCATGTCTCCTG
TTTCTGTCGCCCAGCGCAATACGGGCATCTGGGCGGTCGTGCCGTTGAAAGCGCCTGAGTGTGCCAAAACCCGGTTGTCT
GGCGTTTTAAGCCATGCTGCGCGCCAAGCGTTGTTTTTTTCCATGGCCAGTCATGTCATTGGTACGTTACGCGCATCGCC
TCGGATTGCTTCCCTATTGGTGGTGACGCCGTCGGAAAGTACGGCGGAAATGGCCCGGGCAGCCGGTGCTGAGATCTTGT
GGGGACCACCGGACGAAGGCATGGCGAATGCCTGTTCGCGAGCGATGGCTCATATTGCGGCAGCAGGCGGAGAGCGTGTG
ATGTTTGTCCCCGGTGATTTGCCCCTGTTGGATGGGGCGGCCATCGACATGTTGAGCCGTGCACCGGTCGATGCGATTGG
CATGGCGCCAAACCGGGACGGTCATGGCACGAATGGGCTGATTTGCCGACCTGGCGCTATTCCGTTGTTTTTCAGCGGGC
CAAGCTTTTCCGCTCATCAAAACGCCGCTCGGTGCGCCGGAATTGATGTTTGGATTGTCCGTTCAAGAGAGTGGGCGTTG
GATGTAGACTTGCCTGCCGACCTTGAGGAATTCGAGTCTTCCATCAAAGACGCCAAGAGGAGGGTGCTATGCCAAATCTA
GTAAGCAATCCTATTTACAAAATGATCCAAGAGGTCGAACGTTCCGCTAAGGGCAGTCAGCCAAGGCTAGATGACAAGGC
TTTGGCGTATGAGCTTGAAGCGGTGCAAGATTGGCGTGAGTTGGCGGGTCTGGCATCGGCATGGCGCGATATCGGTTGGG
GCAACGTCATCACCTATTCGCGCAAGGTTTTTATTCCGCTCACCCATCTTTGTCGAGATGTTTGCCACTACTGCACCTTT
GCCAAAGCGCCACGGGTGATTGGCCAAGCATTTCTTACCGTGGACCAAGCGCTGGACATTGCGCGGGCGGGTGCTGCAGT
TGGCTGTCGTGAGGCTTTGTTTACGCTCGGAGATCGGCCCGAAGCGCGTTATGCCGCTGCACGCGATGCGCTGCAGCAAC
TGGGTCATGCTACGACAGCCGACTATGTCGCTGAGGTCGCGCAACGCGTACGAGCTGAAACCGGGCTGCTGCCCCACTTT
AATATGGGCGTGCTAAGCGCGGCAGAATACCAGATGCTGCGGCCCCATGCCCCTTCCTTCGGGTTGATGCTAGAGACGGC
TTCTGAACGTCTTTCCGAGCGTGGTGGGCCGCATTACGGCTCTCCCGATAAGCACCCGGCAGCGCGGCTTGAAACGCTTC
GCCTTGCCGGCGAAGCGCATATACCGATCACCTCAGGCATATTGATTGGTATCGGGGAGACACGCCGCGAGCGGCTTGAA
TCGCTCTTTGCGCTGCGGGATTTGCATGATCGCTATGGTCATGTGCAGGAGGTCATTATTCAGAACTTTCGCGCGAAACC
CGGCACCAAAATGGTTAACGCGCCTGAGCCGTCAATGGACGAATTGTGTTGGACAACAGCGGTCGCCCGGTTGGTCCTGG
GTTCCGCAATGAGCATCCAGGTGCCGCCTAATCTTTTCGACGGTGACTTGACCGATTTGATTCGCGCGGGAATCAATGAT
TGGGGCGGCGTATCGCCGGTCACGCCTGATCACGTTAATCCAGAAGCGCCGTGGCCGCACCTTGACCGCTTAAGTGAGGA
TACAGACAGGGGCGGGAAGACGTTGACCGAGCGGATAACCGTCTATCCTGCCTATATCGAGGAACGTGATCGTTGGATCG
ACCCGGGCTTGCATGCTGATGTGCTACGCCATTCAGATTGCAGTGGATTGGCAACAAGCGATATCTGGAAAGCCGGCTCC
ACGACGATAGGGGATATCGCTTGCCTGGCAGTCACTCCCACGGCTTCCCAGGTTGATCGTGTTGGTTCGCAGACGATCGA
TATCGTCGAGAAATGCGTGGCGGGTGAGCGCCTCGAAGAAGCCGAGCTTGTACATTTATTTAACGCACGCGGCCATGATT
TTACGCATGTGACGCACACGGCGGACCGTTTGCGCAGGCAGACAAAGGGCGACACGATCACTTACGTTGTCAATCGGAAC
ATCAACTACACCAACGTCTGCCAATACCATTGCAATTTTTGTGCTTTTTCACGAGGACCGATACAAGAAGACTTGCGCGA
AAAGCCCTATATCGTTGACCTCAACGAAATCCGTCGCCCGCGTCAAAGAAGCGTGGGATCGAGGGGCGACGGAGGTCTGCT
TGCAAGGGGGCATTCATCCCGACTATACGGGACGTACTTATCTTGACATCTGCGAAGCGGCTAAAACAGAATGTCCAGAC
ATGCACATTCACGCCTTTTCGCCCCTTGAGGTGCTGCATGGTGCCACGACCCTGGGGACGTCAATCAGCGCTTTTTTGTC
TGATTTGAAAAGCGCTGGATTAAGTACGTTGCCTGGGACCGCAGCCGAGATCCTGGACGACGAAGTGCGGCGACAAATCT
GTCCGGATAAGTTGAATACGCAAGAATGGTTGACCGTCGTGCGATCGGCACATGAGTTGGGATTGCGGACCACTTGCACG
ATCATGTTCGGTCATGTCGAATCGTATAAGCATTGGGCGCGACATATCCTCAGGCTTGCCGAGCTGCAACGTGATACAGG
CGGGCTGACTGAGTTTGTTCCATTGCCTTTTGTCCATGAGGAAGCGCCATTATTTAAAAAGAAAGGTGCGCGTCAGGGGC
CGACTTTACGAGAAGCCATTTTGATGCATGCCGTCGGGCGTCTGGCGTTTCATGGCTTGATTGACAATATTCAAACGTCT
TGGGTGAAAATGGGTGAGCACGGCGCACAGCTTTGCTTGCAGGCGGGGGCTAACGATTTGGGCGGTACCCTAATGAATGA
ATCGATTAGCCGCGCAGCCGGTGCAGCCCACGGACAGGAAATGCCGCCTGCCGCCATGGAAGCCCTGGCCGCCAGACTCG
GGCGCAATGCGATGCAGCGTACCCCGCTTTATCGTAACGTCAGTCCTGAGCGGCAGCAAGCTTCGCATGCTGCTCCTGCG
CTGGTGCCTGTGCGATTCACTAAGGCCGGACGCTTGGTGCGTGGGCACGGCCATCAGGTCTTGAGTGAAGCGGAACAATG
ATTGGACCTGCATAGGTGGATCATAGGTGGATTCATGGTGTGGCTTGCATGTTGGCAGCCTCAAATCGATGTTGTGATGG
CTCTCACGCAGCGCTTCCCATCATGGATTCACCAAGGTGCTTAATTACCCTATTAAATTAGACAGACGCTATTCGGTTTG
ACACGATCGGAAGGGTGATCGGTTTGGGTCAGAGAAATGCTTGATCTTTACGCAGGAAGATAATGGCAAAGTATGTTGCG
TTGTGCGGCGGGGTGGGCGGCGCTAAGCTGGCCTATGGTCTAGCGCAGGTGCTTTCTGCCGACGAGTTGACGATCGTTGT
GAACACGGGCGACGATTTTGAGCATCTTGGCTTGCTCATCTGTCCAGACCTGGACACGGTTGTCTATACGCTGGCCGATG
TGGCAGATGCAAAAAAAGGATGGGGGCGGGCGAACGAGAGCTGGTCTTCGCAAGAAGCGCTCGCTCGTTTGGGCGGACCG

GTTTGGTTCCAGCTGGGTGACAAGGACCTTGCGCTGCATCTTTATCGGCGCAGCTTGCTTGACGGTGGTGCCAGCTTGTG
TGATGTAACCGAGACCATTGCGAAAGCGTTAGGTGTCAAGCATTCAATCGTCCCAATGTCTGATGATCCGGTCAGAACCA
TCGTTGAGACAGACGAAGGCGATTTGCCTTTCCAGACGTATTTCGTTAAACGGCGATGTGAGCCACGCGTGTCGGGGTTC
CGCTTTGAAGGGGCTTCGCTAGCACGTTTATCGTCGCCATTTGAAGCAGCGTTGAGTACGCCTGACCTGGCTGGCGTAAT
CTTATGCCCTTCGAATCCTTTTGTCAGCATTGGACCGATTCTGGCATTACCGGGCGTGCGTGACAGGCTGCATGCGTGTA
ATGTGCCTGTTTTAGCTGTTGCGCCGCTTGTCGGTGGTGAAGCAGTCAAAGGCCCGTTGACTAAGATGATGCATGAACTG
GGCATGAGTGTTTCGGTTGGCGAGATTGCTTCGCTTTATGCTGATTTCCTTGATCTGCTGGTGATTGATCCGTTGGATGA
ATGCGATCATGATCTTTTGCGTAAGGATCGTGTGGCAGTCAATAAAGTCAAGACGCTCATGACAACACCTGATGAGCGTA
TCGCGCTTGCGCGACACGTGCTGGCTTGCGTTGAACATCACCGAAACAATCGACAGACAGCTCACGTTGTGCAGCACAGC
ACCGCATAAGGATCCGAATTCGAGCTCGGCGCGCCTGCAGGTCGACAAGCTTGCGGCCGCATAATGCTTAAGTCGAACAG
AAAGTAATCGTATTGTACACGGCCGCATAATCGAAATTAATACGACTCACTATAGGGGAATTGTGAGCGGATAACAATTC
CCCATCTTAGTATATTAGTTAAGTATAAGAAGGAGATATACATATGGCAGATCTCAATTGGATATCGGCCGGCCACGCGA
TCGCTGACGTCGGTACCCTCGAGTCTGGTAAAGAAACCGCTGCTGCGAAATTTGAACGCCAGCACATGGACTCGTCTACT
AGCGCAGCTTAATTAACCTAGGCTGCTGCCACCGCTGAGCAATAACTAGCATAACCCCTTGGGGCCTCTAAACGGGTCTT
GAGGGGTTTTTTGCTGAAACCTCAGGCATTTGAGAAGCACACGGTCACACTGCTTCCGGTAGTCAATAAACCGGTAAACC
AGCAATAGACATAAGCGGCTATTTAACGACCCTGCCCTGAACCGACGACCGGGTCATCGTGGCCGGATCTTGCGGCCCCT
CGGCTTGAACGAATTGTTAGACATTATTTGCCGACTACCTTGGTGATCTCGCCTTTCACGTAGTGGACAAATTCTTCCAA
CTGATCTGCGCGCGAGGCCAAGCGATCTTCTTCTTGTCCAAGATAAGCCTGTCTAGCTTCAAGTATGACGGGCTGATACT
GGGCCGGCAGGCGCTCCATTGCCCAGTCGGCAGCGACATCCTTCGGCGCGATTTTGCCGGTTACTGCGCTGTACCAAATG
CGGGACAACGTAAGCACTACATTTCGCTCATCGCCAGCCCAGTCGGGCGGCGAGTTCCATAGCGTTAAGGTTTCATTTAG
CGCCTCAAATAGATCCTGTTCAGGAACCGGATCAAAGAGTTCCTCCGCCGCTGGACCTACCAAGGCAACGCTATGTTCTC
TTGCTTTTGTCAGCAAGATAGCCAGATCAATGTCGATCGTGGCTGGCTCGAAGATACCTGCAAGAATGTCATTGCGCTGC
CATTCTCCAAATTGCAGTTCGCGCTTAGCTGGATAACGCCACGGAATGATGTCGTCGTGCACAACAATGGTGACTTCTAC
AGCGCGGAGAATCTCGCTCTCTCCAGGGGAAGCCGAAGTTTCCAAAAGGTCGTTGATCAAAGCTCGCCGCGTTGTTTCAT
CAAGCCTTACGGTCACCGTAACCAGCAAATCAATATCACTGTGTGGCTTCAGGCCGCCATCCACTGCGGAGCCGTACAAA
TGTACGGCCAGCAACGTCGGTTCGAGATGGCGCTCGATGACGCCAACTACCTCTGATAGTTGAGTCGATACTTCGGCGAT
CACCGCTTCCCTCATACTCTTCCTTTTTCAATATTATTGAAGCATTTATCAGGGTTATTGTCTCATGAGCGGATACATAT
TTGAATGTATTTAGAAAAATAAACAAATAGCTAGCTCACTCGGTCGCTACGCTCCGGGCGTGAGACTGCGGCGGGCGCTG
CGGACACATACAAAGTTACCCACAGATTCCGTGGATAAGCAGGGGACTAACATGTGAGGCAAAACAGCAGGGCCGCGCCG
GTGGCGTTTTTCCATAGGCTCCGCCCTCCTGCCAGAGTTCACATAAACAGACGCTTTTCCGGTGCATCTGTGGGAGCCGT
GAGGCTCAACCATGAATCTGACAGTACGGGCGAAACCCGACAGGACTTAAAGATCCCCACCGTTTCCGGCGGGTCGCTCC
CTCTTGCGCTCTCCTGTTCCGACCCTGCCGTTTACCGGATACCTGTTCCGCCTTTCTCCCTTACGGGAAGTGTGGCGCTT
TCTCATAGCTCACACACTGGTATCTCGGCTCGGTGTAGGTCGTTCGCTCCAAGCTGGGCTGTAAGCAAGAACTCCCCGTT
CAGCCCGACTGCTGCGCCTTATCCGGTAACTGTTCACTTGAGTCCAACCCGGAAAAGCACGGTAAAACGCCACTGGCAGC
AGCCATTGGTAACTGGGAGTTCGCAGAGGATTTGTTTAGCTAAACACGCGGTTGCTCTTGAAGTGTGCGCCAAAGTCCGG
CTACACTGGAAGGACAGATTTGGTTGCTGTGCTCTGCGAAAGCCAGTTACCACGGTTAAGCAGTTCCCCAACTGACTTAA
CCTTCGATCAAACCACCTCCCCAGGTGGTTTTTTCGTTTACAGGGCAAAAGATTACGCGCAGAAAAAAAGGATCTCAAGA
AGATCCTTTGATCTTTTCTACTGAACCGCTCTAGATTTCAGTGCAATTTATCTCTTCAAATGTAGCACCTGAAGTCAGCC
CCATACGATATAAGTTGTAATTCTCATGTTAGTCATGCCCCGCGCCCACCGGAAGGAGCTGACTGGGTTGAAGGCTCTCA
AGGGCATCGGTCGAGATCCCGGTGCCTAATGAGTGAGCTAACTTACATTAATTGCGTTGCGCTCACTGCCCGCTTTCCAG
TCGGGAAACCTGTCGTGCCAGCTGCATTAATGAATCGGCCAACGCGCGGGGAGAGGCGGTTTGCGTATTGGGCGCCAGGG
TGGTTTTTCTTTTCACCAGTGAGACGGGCAACAGCTGATTGCCCTTCACCGCCTGGCCCTGAGAGAGTTGCAGCAAGCGG
TCCACGCTGGTTTGCCCCAGCAGGCGAAAATCCTGTTTGATGGTGGTTAACGGCGGGATATAACATGAGCTGTCTTCGGT
ATCGTCGTATCCCACTACCGAGATGTCCGCACCAACGCGCAGCCCGGACTCGGTAATGGCGCGCATTGCGCCCAGCGCCA
TCTGATCGTTGGCAACCAGCATCGCAGTGGGAACGATGCCCTCATTCAGCATTTGCATGGTTTGTTGAAAACCGGACATG
GCACTCCAGTCGCCTTCCCGTTCCGCTATCGGCTGAATTTGATTGCGAGTGAGATATTTATGCCAGCCAGCCAGACGCAG
ACGCGCCGAGACAGAACTTAATGGGCCCGCTAACAGCGCGATTTGCTGGTGACCCAATGCGACCAGATGCTCCACGCCCA
GTCGCGTACCGTCTTCATGGGAGAAAATAATACTGTTGATGGGTGTCTGGTCAGAGACATCAAGAAATAACGCCGGAACA
TTAGTGCAGGCAGCTTCCACAGCAATGGCATCCTGGTCATCCAGCGGATAGTTAATGATCAGCCCACTGACGCGTTGCGC
GAGAAGATTGTGCACCGCCGCTTTACAGGCTTCGACGCCGCTTCGTTCTACCATCGACACCACCACGCTGGCACCCAGTT
GATCGGCGCGAGATTTAATCGCCGCGACAATTTGCGACGGCGCGTGCAGGGCCAGACTGGAGGTGGCAACGCCAATCAGC
AACGACTGTTTGCCCGCCAGTTGTTGTGCCACGCGGTTGGGAATGTAATTCAGCTCCGCCATCGCCGCTTCCACTTTTTC
CCGCGTTTTCGCAGAAACGTGGCTGGCCTGGTTCACCACGCGGGAAACGGTCTGATAAGAGACACCGGCATACTCTGCGA
CATCGTATAACGTTACTGGTTTCACATTCACCACCCTGAATTGACTCTCTTCCGGGCGCTATCATGCCATACCGCGAAAG
GTTTTGCGCCATTCGATGGTGTCCGGGATCTCGACGCTCTCCCTTATGCGACTCCTGCATTAGGAAATTAATACGACTCA
CTATA
>pFS01
GCGCCAGCAACCGCACCTGTGGCGCCGGTGATGCCGGCCACGATGCGTCCGGCGTAGAGGATCGAGATCTCGATCCCGCG
AAATTAATACGACTCACTATAGGGGAATTGTGAGCGGATAACAATTCCCCTCTAGAAATAATTTTGTTTAACTTTAAGAA
GGAGATATACCATGGGCAGCAGCCATCATCATCATCATCACAGCAGCGGCCTGGTGCCGCGCGGCAGCCATATGGCTAGC
ATGACTGGTGGACAGCAAATGGGTCGCGGATCCATGACTGTATCCGCCATCGGTGGCATTCCCTTAATACAAACAGGGGA
TGATCTTGGCCAGATTATTAAGGAGGCGATCCACAAAAACGGTATCGCTCTAGAAAATGGCGACGTGTTAGTGCTTGCGC
AAAAGATCGTTTCAAAGTCCGAAGGACGATGGGCTGCACTATCTTCTGTCACGCCCGGCAAACAAGCTATCGATTTGGCG
CAGAAAGTGGATAAAGATCCACGGCTCGTTGAACTGATCCTATCGGAGTCGGCAGAAATCGTTGCACACAGGCAAGACGG

TGTACTGATCACCGCTCACCGTCTCGGATGCGTGATGGCGAATGCCGGTATCGATCATTCCAATGTTGGTGATGAAGACA
GTGTGCTTCTGCTTCCGAAAGACCCGGATCACAGTGCACGCGAATTAAAAAAACAGTTTCATCGCTTGTGCGGCGTCGAT
GTCCACATCATCATCAACGATAGTTTTGGCAGAGTGTGGCGCCATGGCACGGCAGGATGCGCAATCGGTGTAGCGGGCTT
TTCCCCGCTTAAAAATTATATTGGGAAGCCGGATCTGTTTGGGCAGCTCTTACGAACAACCCAAGTGGCAGTGGCCGATG
AACTCGCCGCAGCAGCGTCTTTCTTAATGGGACAAGCCGATGAAGGATCGCCTGTTGTGCTGATCCGCGGCGCGAATTTG
CCCCCCGCTGATGGAACGGTCCAGCAGCTTATTCGCCCCAAAGAGGAAGATTTATTTCGTGAAACCTCCTCCACACTGGC
GGTGGAAATGGCCTAAGAATTCGAGCTCCGTCGACAAGCTTGCGGCCGCACTCGAGCACCACCACCACCACCACTGAGAT
CCGGCTGCTAACAAAGCCCGAAAGGAAGCTGAGTTGGCTGCTGCCACCGCTGAGCAATAACTAGCATAACCCCTTGGGGC
CTCTAAACGGGTCTTGAGGGGTTTTTTGCTGAAAGGAGGAACTATATCCGGATTGGCGAATGGGACGCGCCCTGTAGCGG
CGCATTAAGCGCGGCGGGTGTGGTGGTTACGCGCAGCGTGACCGCTACACTTGCCAGCGCCCTAGCGCCCGCTCCTTTCG
CTTTCTTCCCTTCCTTTCTCGCCACGTTCGCCGGCTTTCCCCGTCAAGCTCTAAATCGGGGGCTCCCTTTAGGGTTCCGA
TTTAGTGCTTTACGGCACCTCGACCCCAAAAAACTTGATTAGGGTGATGGTTCACGTAGTGGGCCATCGCCCTGATAGAC
GGTTTTTCGCCCTTTGACGTTGGAGTCCACGTTCTTTAATAGTGGACTCTTGTTCCAAACTGGAACAACACTCAACCCTA
TCTCGGTCTATTCTTTTGATTTATAAGGGATTTTGCCGATTTCGGCCTATTGGTTAAAAAATGAGCTGATTTAACAAAAA
TTTAACGCGAATTTTAACAAAATATTAACGTTTACAATTTCAGGTGGCACTTTTCGGGGAAATGTGCGCGGAACCCCTAT
TTGTTTATTTTTCTAAATACATTCAAATATGTATCCGCTCATGAATTAATTCTTAGAAAAACTCATCGAGCATCAAATGA
AACTGCAATTTATTCATATCAGGATTATCAATACCATATTTTTGAAAAAGCCGTTTCTGTAATGAAGGAGAAAACTCACC
GAGGCAGTTCCATAGGATGGCAAGATCCTGGTATCGGTCTGCGATTCCGACTCGTCCAACATCAATACAACCTATTAATT
TCCCCTCGTCAAAAATAAGGTTATCAAGTGAGAAATCACCATGAGTGACGACTGAATCCGGTGAGAATGGCAAAAGTTTA
TGCATTTCTTTCCAGACTTGTTCAACAGGCCAGCCATTACGCTCGTCATCAAAATCACTCGCATCAACCAAACCGTTATT
CATTCGTGATTGCGCCTGAGCGAGACGAAATACGCGATCGCTGTTAAAAGGACAATTACAAACAGGAATCGAATGCAACC
GGCGCAGGAACACTGCCAGCGCATCAACAATATTTTCACCTGAATCAGGATATTCTTCTAATACCTGGAATGCTGTTTTC
CCGGGGATCGCAGTGGTGAGTAACCATGCATCATCAGGAGTACGGATAAAATGCTTGATGGTCGGAAGAGGCATAAATTC
CGTCAGCCAGTTTAGTCTGACCATCTCATCTGTAACATCATTGGCAACGCTACCTTTGCCATGTTTCAGAAACAACTCTG
GCGCATCGGGCTTCCCATACAATCGATAGATTGTCGCACCTGATTGCCCGACATTATCGCGAGCCCATTTATACCCATAT
AAATCAGCATCCATGTTGGAATTTAATCGCGGCCTAGAGCAAGACGTTTCCCGTTGAATATGGCTCATAACACCCCTTGT
ATTACTGTTTATGTAAGCAGACAGTTTTATTGTTCATGACCAAAATCCCTTAACGTGAGTTTTCGTTCCACTGAGCGTCA
GACCCCGTAGAAAAGATCAAAGGATCTTCTTGAGATCCTTTTTTTCTGCGCGTAATCTGCTGCTTGCAAACAAAAAAACC
ACCGCTACCAGCGGTGGTTTGTTTGCCGGATCAAGAGCTACCAACTCTTTTTCCGAAGGTAACTGGCTTCAGCAGAGCGC
AGATACCAAATACTGTCCTTCTAGTGTAGCCGTAGTTAGGCCACCACTTCAAGAACTCTGTAGCACCGCCTACATACCTC
GCTCTGCTAATCCTGTTACCAGTGGCTGCTGCCAGTGGCGATAAGTCGTGTCTTACCGGGTTGGACTCAAGACGATAGTT
ACCGGATAAGGCGCAGCGGTCGGGCTGAACGGGGGGTTCGTGCACACAGCCCAGCTTGGAGCGAACGACCTACACCGAAC
TGAGATACCTACAGCGTGAGCTATGAGAAAGCGCCACGCTTCCCGAAGGGAGAAAGGCGGACAGGTATCCGGTAAGCGGC
AGGGTCGGAACAGGAGAGCGCACGAGGGAGCTTCCAGGGGGAAACGCCTGGTATCTTTATAGTCCTGTCGGGTTTCGCCA
CCTCTGACTTGAGCGTCGATTTTTGTGATGCTCGTCAGGGGGGCGGAGCCTATGGAAAAACGCCAGCAACGCGGCCTTTT
TACGGTTCCTGGCCTTTTGCTGGCCTTTTGCTCACATGTTCTTTCCTGCGTTATCCCCTGATTCTGTGGATAACCGTATT
ACCGCCTTTGAGTGAGCTGATACCGCTCGCCGCAGCCGAACGACCGAGCGCAGCGAGTCAGTGAGCGAGGAAGCGGAAGA
GCGCCTGATGCGGTATTTTCTCCTTACGCATCTGTGCGGTATTTCACACCGCATATATGGTGCACTCTCAGTACAATCTG
CTCTGATGCCGCATAGTTAAGCCAGTATACACTCCGCTATCGCTACGTGACTGGGTCATGGCTGCGCCCCGACACCCGCC
AACACCCGCTGACGCGCCCTGACGGGCTTGTCTGCTCCCGGCATCCGCTTACAGACAAGCTGTGACCGTCTCCGGGAGCT
GCATGTGTCAGAGGTTTTCACCGTCATCACCGAAACGCGCGAGGCAGCTGCGGTAAAGCTCATCAGCGTGGTCGTGAAGC
GATTCACAGATGTCTGCCTGTTCATCCGCGTCCAGCTCGTTGAGTTTCTCCAGAAGCGTTAATGTCTGGCTTCTGATAAA
GCGGGCCATGTTAAGGGCGGTTTTTTCCTGTTTGGTCACTGATGCCTCCGTGTAAGGGGGATTTCTGTTCATGGGGGTAA
TGATACCGATGAAACGAGAGAGGATGCTCACGATACGGGTTACTGATGATGAACATGCCCGGTTACTGGAACGTTGTGAG
GGTAAACAACTGGCGGTATGGATGCGGCGGGACCAGAGAAAAATCACTCAGGGTCAATGCCAGCGCTTCGTTAATACAGA
TGTAGGTGTTCCACAGGGTAGCCAGCAGCATCCTGCGATGCAGATCCGGAACATAATGGTGCAGGGCGCTGACTTCCGCG
TTTCCAGACTTTACGAAACACGGAAACCGAAGACCATTCATGTTGTTGCTCAGGTCGCAGACGTTTTGCAGCAGCAGTCG
CTTCACGTTCGCTCGCGTATCGGTGATTCATTCTGCTAACCAGTAAGGCAACCCCGCCAGCCTAGCCGGGTCCTCAACGA
CAGGAGCACGATCATGCGCACCCGTGGGGCCGCCATGCCGGCGATAATGGCCTGCTTCTCGCCGAAACGTTTGGTGGCGG
GACCAGTGACGAAGGCTTGAGCGAGGGCGTGCAAGATTCCGAATACCGCAAGCGACAGGCCGATCATCGTCGCGCTCCAG
CGAAAGCGGTCCTCGCCGAAAATGACCCAGAGCGCTGCCGGCACCTGTCCTACGAGTTGCATGATAAAGAAGACAGTCAT
AAGTGCGGCGACGATAGTCATGCCCCGCGCCCACCGGAAGGAGCTGACTGGGTTGAAGGCTCTCAAGGGCATCGGTCGAG
ATCCCGGTGCCTAATGAGTGAGCTAACTTACATTAATTGCGTTGCGCTCACTGCCCGCTTTCCAGTCGGGAAACCTGTCG
TGCCAGCTGCATTAATGAATCGGCCAACGCGCGGGGAGAGGCGGTTTGCGTATTGGGCGCCAGGGTGGTTTTTCTTTTCA
CCAGTGAGACGGGCAACAGCTGATTGCCCTTCACCGCCTGGCCCTGAGAGAGTTGCAGCAAGCGGTCCACGCTGGTTTGC
CCCAGCAGGCGAAAATCCTGTTTGATGGTGGTTAACGGCGGGATATAACATGAGCTGTCTTCGGTATCGTCGTATCCCAC
TACCGAGATATCCGCACCAACGCGCAGCCCGGACTCGGTAATGGCGCGCATTGCGCCCAGCGCCATCTGATCGTTGGCAA
CCAGCATCGCAGTGGGAACGATGCCCTCATTCAGCATTTGCATGGTTTGTTGAAAACCGGACATGGCACTCCAGTCGCCT
TCCCGTTCCGCTATCGGCTGAATTTGATTGCGAGTGAGATATTTATGCCAGCCAGCCAGACGCAGACGCGCCGAGACAGA
ACTTAATGGGCCCGCTAACAGCGCGATTTGCTGGTGACCCAATGCGACCAGATGCTCCACGCCCAGTCGCGTACCGTCTT
CATGGGAGAAAATAATACTGTTGATGGGTGTCTGGTCAGAGACATCAAGAAATAACGCCGGAACATTAGTGCAGGCAGCT
TCCACAGCAATGGCATCCTGGTCATCCAGCGGATAGTTAATGATCAGCCCACTGACGCGTTGCGCGAGAAGATTGTGCAC
CGCCGCTTTACAGGCTTCGACGCCGCTTCGTTCTACCATCGACACCACCACGCTGGCACCCAGTTGATCGGCGCGAGATT
TAATCGCCGCGACAATTTGCGACGGCGCGTGCAGGGCCAGACTGGAGGTGGCAACGCCAATCAGCAACGACTGTTTGCCC

GCCAGTTGTTGTGCCACGCGGTTGGGAATGTAATTCAGCTCCGCCATCGCCGCTTCCACTTTTTCCCGCGTTTTCGCAGA
AACGTGGCTGGCCTGGTTCACCACGCGGGAAACGGTCTGATAAGAGACACCGGCATACTCTGCGACATCGTATAACGTTA
CTGGTTTCACATTCACCACCCTGAATTGACTCTCTTCCGGGCGCTATCATGCCATACCGCGAAAGGTTTTGCGCCATTCG
ATGGTGTCCGGGATCTCGACGCTCTCCCTTATGCGACTCCTGCATTAGGAAGCAGCCCAGTAGTAGGTTGAGGCCGTTGA
GCACCGCCGCCGCAAGGAATGGTGCATGCAAGGAGATGGCGCCCAACAGTCCCCCGGCCACGGGGCCTGCCACCATACCC
ACGCCGAAACAAGCGCTCATGAGCCCGAAGTGGCGAGCCCGATCTTCCCCATCGGTGATGTCGGCGATATAG
>pFS03
GGGGAATTGTGAGCGGATAACAATTCCCCTGTAGAAATAATTTTGTTTAACTTTAATAAGGAGATATACCATGGGCAGCA
GCCATCACCATCATCACCACAGCCAGGATCCGAATTCGAGCTCGGCGCGCCTGCAGGTCGACAAGCTTGCGATGTCTCCT
GTTTCTGTCGCCCAGCGCAATACGGGCATCTGGGCGGTCGTGCCGTTGAAAGCGCCTGAGTGTGCCAAAACCCGGTTGTC
TGGCGTTTTAAGCCATGCTGCGCGCCAAGCGTTGTTTTTTTCCATGGCCAGTCATGTCATTGGTACGTTACGCGCATCGC
CTCGGATTGCTTCCCTATTGGTGGTGACGCCGTCGGAAAGTACGGCGGAAATGGCCCGGGCAGCCGGTGCTGAGATCTTG
TGGGGACCACCGGACGAAGGCATGGCGAATGCCTGTTCGCGAGCGATGGCTCATATTGCGGCAGCAGGCGGAGAGCGTGT
GATGTTTGTCCCCGGTGATTTGCCCCTGTTGGATGGGGCGGCCATCGACATGTTGAGCCGTGCACCGGTCGATGCGATTG
GCATGGCGCCAAACCGGGACGGTCATGGCACGAATGGGCTGATTTGCCGACCTGGCGCTATTCCGTTGTTTTCAGCGGG
CCAAGCTTTTCCGCTCATCAAAACGCCGCTCGGTGCGCCGGAATTGATGTTTGGATTGTCCGTTCAAGAGAGTGGGCGTT
GGATGTAGACTTGCCTGCCGACCTTGAGGAATTCGAGTCTTCCATCAAAGACGCCAAGAGGAGGGTGCTATGCCAAATCT
AGGCCGCATAATGCTTAAGTCGAACAGAAAGTAATCGTATTGTACACGGCCGCATAATCGAAATTAATACGACTCACTAT
AGGGGAATTGTGAGCGGATAACAATTCCCCATCTTAGTATATTAGTTAAGTATAAGAAGGAGATATACATATGGCAGATC
TCAATTGGATATCGGCCGGCCACGCGATCGCTGACGTCGGTACCCTCGAGTCTGGTAAAGAAACCGCTGCTGCGAAATTT
GAACGCCAGCACATGGACTCGTCTACTAGCGCAGCTTAATTAACCTAGGCTGCTGCCACCGCTGAGCAATAACTAGCATA
ACCCCTTGGGGCCTCTAAACGGGTCTTGAGGGGTTTTTTGCTGAAACCTCAGGCATTTGAGAAGCACACGGTCACACTGC
TTCCGGTAGTCAATAAACCGGTAAACCAGCAATAGACATAAGCGGCTATTTAACGACCCTGCCCTGAACCGACGACCGGG
TCGAATTTGCTTTCGAATTTCTGCCATTCATCCGCTTATTATCACTTATTCAGGCGTAGCACCAGGCGTTTAAGGGCACC
AATAACTGCCTTAAAAAAATTACGCCCCGCCCTGCCACTCATCGCAGTACTGTTGTAATTCATTAAGCATTCTGCCGACA
TGGAAGCCATCACAGACGGCATGATGAACCTGAATCGCCAGCGGCATCAGCACCTTGTCGCCTTGCGTATAATATTTGCC
CATAGTGAAAACGGGGGCGAAGAAGTTGTCCATATTGGCCACGTTTAAATCAAAACTGGTGAAACTCACCCAGGGATTGG
CTGAGACGAAAAACATATTCTCAATAAACCCTTTAGGGAAATAGGCCAGGTTTTCACCGTAACACGCCACATCTTGCGAA
TATATGTGTAGAAACTGCCGGAAATCGTCGTGGTATTCACTCCAGAGCGATGAAAACGTTTCAGTTTGCTCATGGAAAAC
GGTGTAACAAGGGTGAACACTATCCCATATCACCAGCTCACCGTCTTTCATTGCCATACGGAACTCCGGATGAGCATTCA
TCAGGCGGGCAAGAATGTGAATAAAGGCCGGATAAAACTTGTGCTTATTTTTCTTTACGGTCTTTAAAAAGGCCGTAATA
TCCAGCTGAACGGTCTGGTTATAGGTACATTGAGCAACTGACTGAAATGCCTCAAAATGTTCTTTACGATGCCATTGGGA
TATATCAACGGTGGTATATCCAGTGATTTTTTTCTCCATTTTAGCTTCCTTAGCTCCTGAAAATCTCGATAACTCAAAAA
ATACGCCCGGTAGTGATCTTATTTCATTATGGTGAAAGTTGGAACCTCTTACGTGCCGATCAACGTCTCATTTTCGCCAA
AAGTTGGCCCAGGGCTTCCCGGTATCAACAGGGACACCAGGATTTATTTATTCTGCGAAGTGATCTTCCGTCACAGGTAT
TTATTCGGCGCAAAGTGCGTCGGGTGATGCTGCCAACTTACTGATTTAGTGTATGATGGTGTTTTTGAGGTGCTCCAGTG
GCTTCTGTTTCTATCAGCTGTCCCTCCTGTTCAGCTACTGACGGGGTGGTGCGTAACGGCAAAAGCACCGCCGGACATCA
GCGCTAGCGGAGTGTATACTGGCTTACTATGTTGGCACTGATGAGGGTGTCAGTGAAGTGCTTCATGTGGCAGGAGAAAA
AAGGCTGCACCGGTGCGTCAGCAGAATATGTGATACAGGATATATTCCGCTTCCTCGCTCACTGACTCGCTACGCTCGGT
CGTTCGACTGCGGCGAGCGGAAATGGCTTACGAACGGGGCGGAGATTTCCTGGAAGATGCCAGGAAGATACTTAACAGGG
AAGTGAGAGGGCCGCGGCAAAGCCGTTTTTCCATAGGCTCCGCCCCCCTGACAAGCATCACGAAATCTGACGCTCAAATC
AGTGGTGGCGAAACCCGACAGGACTATAAAGATACCAGGCGTTTCCCCTGGCGGCTCCCTCGTGCGCTCTCCTGTTCCTG
CCTTTCGGTTTACCGGTGTCATTCCGCTGTTATGGCCGCGTTTGTCTCATTCCACGCCTGACACTCAGTTCCGGGTAGGC
AGTTCGCTCCAAGCTGGACTGTATGCACGAACCCCCCGTTCAGTCCGACCGCTGCGCCTTATCCGGTAACTATCGTCTTG
AGTCCAACCCGGAAAGACATGCAAAAGCACCACTGGCAGCAGCCACTGGTAATTGATTTAGAGGAGTTAGTCTTGAAGTC
ATGCGCCGGTTAAGGCTAAACTGAAAGGACAAGTTTTGGTGACTGCGCTCCTCCAAGCCAGTTACCTCGGTTCAAAGAGT
TGGTAGCTCAGAGAACCTTCGAAAAACCGCCCTGCAAGGCGGTTTTTTCGTTTTCAGAGCAAGAGATTACGCGCAGACCA
AAACGATCTCAAGAAGATCATCTTATTAATCAGATAAAATATTTCTAGATTTCAGTGCAATTTATCTCTTCAAATGTAGC
ACCTGAAGTCAGCCCCATACGATATAAGTTGTAATTCTCATGTTAGTCATGCCCCGCGCCCACCGGAAGGAGCTGACTGG
GTTGAAGGCTCTCAAGGGCATCGGTCGAGATCCCGGTGCCTAATGAGTGAGCTAACTTACATTAATTGCGTTGCGCTCAC
TGCCCGCTTTCCAGTCGGGAAACCTGTCGTGCCAGCTGCATTAATGAATCGGCCAACGCGCGGGGAGAGGCGGTTTGCGT
ATTGGGCGCCAGGGTGGTTTTTCTTTTCACCAGTGAGACGGGCAACAGCTGATTGCCCTTCACCGCCTGGCCCTGAGAGA
GTTGCAGCAAGCGGTCCACGCTGGTTTGCCCCAGCAGGCGAAAATCCTGTTTGATGGTGGTTAACGGCGGGATATAACAT
GAGCTGTCTTCGGTATCGTCGTATCCCACTACCGAGATGTCCGCACCAACGCGCAGCCCGGACTCGGTAATGGCGCGCAT
TGCGCCCAGCGCCATCTGATCGTTGGCAACCAGCATCGCAGTGGGAACGATGCCCTCATTCAGCATTTGCATGGTTTGTT
GAAAACCGGACATGGCACTCCAGTCGCCTTCCCGTTCCGCTATCGGCTGAATTTGATTGCGAGTGAGATATTTATGCCAG
CCAGCCAGACGCAGACGCGCCGAGACAGAACTTAATGGGCCCGCTAACAGCGCGATTTGCTGGTGACCCAATGCGACCAG
ATGCTCCACGCCCAGTCGCGTACCGTCTTCATGGGAGAAAATAATACTGTTGATGGGTGTCTGGTCAGAGACATCAAGAA
ATAACGCCGGAACATTAGTGCAGGCAGCTTCCACAGCAATGGCATCCTGGTCATCCAGCGGATAGTTAATGATCAGCCCA
CTGACGCGTTGCGCGAGAAGATTGTGCACCGCCGCTTTACAGGCTTCGACGCCGCTTCGTTCTACCATCGACACCACCAC
GCTGGCACCCAGTTGATCGGCGCGAGATTTAATCGCCGCGACAATTTGCGACGGCGCGTGCAGGGCCAGACTGGAGGTGG
CAACGCCAATCAGCAACGACTGTTTGCCCGCCAGTTGTTGTGCCACGCGGTTGGGAATGTAATTCAGCTCCGCCATCGCC
GCTTCCACTTTTTCCCGCGTTTTCGCAGAAACGTGGCTGGCCTGGTTCACCACGCGGGAAACGGTCTGATAAGAGACACC
GGCATACTCTGCGACATCGTATAACGTTACTGGTTTCACATTCACCACCCTGAATTGACTCTCTTCCGGGCGCTATCATG

CCATACCGCGAAAGGTTTTGCGCCATTCGATGGTGTCCGGGATCTCGACGCTCTCCCTTATGCGACTCCTGCATTAGGAA
ATTAATACGACTCACTATA
>pFS04
GCGCCAGCAACCGCACCTGTGGCGCCGGTGATGCCGGCCACGATGCGTCCGGCGTAGAGGATCGAGATCTCGATCCCGCG
AAATTAATACGACTCACTATAGGGGAATTGTGAGCGGATAACAATTCCCCTCTAGAAATAATTTTGTTTAACTTTAAGAA
GGAGATATACCATGGGCAGCAGCCATCATCATCATCATCACAGCAGCGGCCTGGTGCCGCGCGGCAGCCATATGGCTAGC
ATGACTGGTGGACAGCAAATGGGTCGCGGATCCGTGATTACCGTTCTGAGCGGTGGTACAGGCACCCCGAAACTGCTGCA
GGGTCTGAAACGTGTTGTTAATAATGAAGAACTGGCCGTGATTGTGAATACCGGTGAAGATACCTGGATTGGTGATCTGT
ATCTGAGTCCGGATGTTGATACCGTGCTGTATACCCTGGCAGATCTGATTAACGAAGAAACCTGGTATGGTGTGAAAGAG
GATACCTTTTATACCCACGAACAGCTGAAAAATCTGGGCTTTGATGAAGTTCTGCGCATTGGTGATAAAGATCGTGCACT
GAAAATGCACAAAACCTATTATCTGAAACGCGGTCATAAACTGAGCGAAGTTGTTGATATGGAAAAAGTTGCCCTGGGCA
TTAAAGCAAAAGTTATTCCGATGACCGATGATCGTGTGGAAACCAAAATTCTGGCAAAAGTTGATGGTAAAGTGGACCTG
CTGAAATTCCATGATTTTTGGGTTAAACGCAAAGGTGATGTTGAAGTGCTGGATGTGATTTATGAAAACAGCCTGTATGC
AAAACCGTGCGAAAAAGCAGTTGAAGCCATTAAAAACAGCGATCTGGTTATTATTGGTCCGAGCAATCCGATTACCAGCA
TCGGTCCGATTCTGAGCCTGAATGGTATTAAAGAACTGCTGAAAGACAAAAAAGTTGTTGTGGTTAGCCCGATTGTTGGT
AATAGCGCAGTTAGCGGTCCGGCAGGTAAACTGATGAAAGCCAAAGGTTATGATGTTAGCGTGAAAGGCATCTACGAGTT
CTATAAAGATATTGTGGATGTGCTGGTGATCGACAACGTGGATAAAGAAATTGCAAAAGAAATTCCGTGCGAAGTGCTGA
TTACCAATACCATCATGAAAACCCTGGATGATAAAGTTCGTCTGGCCAAAAACATCATTGAATTTTGTGGTAGCCTGTAA
GAATTCGAGCTCCGTCGACAAGCTTGCGGCCGCACTCGAGCACCACCACCACCACCACTGAGATCCGGCTGCTAACAAAG
CCCGAAAGGAAGCTGAGTTGGCTGCTGCCACCGCTGAGCAATAACTAGCATAACCCCTTGGGGCCTCTAAACGGGTCTTG
AGGGGTTTTTTGCTGAAAGGAGGAACTATATCCGGATTGGCGAATGGGACGCGCCCTGTAGCGGCGCATTAAGCGCGGCG
GGTGTGGTGGTTACGCGCAGCGTGACCGCTACACTTGCCAGCGCCCTAGCGCCCGCTCCTTTCGCTTTCTTCCCTTCCTT
TCTCGCCACGTTCGCCGGCTTTCCCCGTCAAGCTCTAAATCGGGGGCTCCCTTTAGGGTTCCGATTTAGTGCTTTACGGC
ACCTCGACCCCAAAAAACTTGATTAGGGTGATGGTTCACGTAGTGGGCCATCGCCCTGATAGACGGTTTTTCGCCCTTTG
ACGTTGGAGTCCACGTTCTTTAATAGTGGACTCTTGTTCCAAACTGGAACAACACTCAACCCTATCTCGGTCTATTCTTT
TGATTTATAAGGGATTTTGCCGATTTCGGCCTATTGGTTAAAAAATGAGCTGATTTAACAAAAATTTAACGCGAATTTTA
ACAAAATATTAACGTTTACAATTTCAGGTGGCACTTTTCGGGGAAATGTGCGCGGAACCCCTATTTGTTTATTTTTCTAA
ATACATTCAAATATGTATCCGCTCATGAATTAATTCTTAGAAAAACTCATCGAGCATCAAATGAAACTGCAATTTATTCA
TATCAGGATTATCAATACCATATTTTTGAAAAAGCCGTTTCTGTAATGAAGGAGAAAACTCACCGAGGCAGTTCCATAGG
ATGGCAAGATCCTGGTATCGGTCTGCGATTCCGACTCGTCCAACATCAATACAACCTATTAATTTCCCCTCGTCAAAAAT
AAGGTTATCAAGTGAGAAATCACCATGAGTGACGACTGAATCCGGTGAGAATGGCAAAAGTTTATGCATTTCTTTCCAGA
CTTGTTCAACAGGCCAGCCATTACGCTCGTCATCAAAATCACTCGCATCAACCAAACCGTTATTCATTCGTGATTGCGCC
TGAGCGAGACGAAATACGCGATCGCTGTTAAAAGGACAATTACAAACAGGAATCGAATGCAACCGGCGCAGGAACACTGC
CAGCGCATCAACAATATTTTCACCTGAATCAGGATATTCTTCTAATACCTGGAATGCTGTTTTCCCGGGGATCGCAGTGG
TGAGTAACCATGCATCATCAGGAGTACGGATAAAATGCTTGATGGTCGGAAGAGGCATAAATTCCGTCAGCCAGTTTAGT
CTGACCATCTCATCTGTAACATCATTGGCAACGCTACCTTTGCCATGTTTCAGAAACAACTCTGGCGCATCGGGCTTCCC
ATACAATCGATAGATTGTCGCACCTGATTGCCCGACATTATCGCGAGCCCATTTATACCCATATAAATCAGCATCCATGT
TGGAATTTAATCGCGGCCTAGAGCAAGACGTTTCCCGTTGAATATGGCTCATAACACCCCTTGTATTACTGTTTATGTAA
GCAGACAGTTTTATTGTTCATGACCAAAATCCCTTAACGTGAGTTTTCGTTCCACTGAGCGTCAGACCCCGTAGAAAAGA
TCAAAGGATCTTCTTGAGATCCTTTTTTTCTGCGCGTAATCTGCTGCTTGCAAACAAAAAAACCACCGCTACCAGCGGTG
GTTTGTTTGCCGGATCAAGAGCTACCAACTCTTTTTCCGAAGGTAACTGGCTTCAGCAGAGCGCAGATACCAAATACTGT
CCTTCTAGTGTAGCCGTAGTTAGGCCACCACTTCAAGAACTCTGTAGCACCGCCTACATACCTCGCTCTGCTAATCCTGT
TACCAGTGGCTGCTGCCAGTGGCGATAAGTCGTGTCTTACCGGGTTGGACTCAAGACGATAGTTACCGGATAAGGCGCAG
CGGTCGGGCTGAACGGGGGGTTCGTGCACACAGCCCAGCTTGGAGCGAACGACCTACACCGAACTGAGATACCTACAGCG
TGAGCTATGAGAAAGCGCCACGCTTCCCGAAGGGAGAAAGGCGGACAGGTATCCGGTAAGCGGCAGGGTCGGAACAGGAG
AGCGCACGAGGGAGCTTCCAGGGGGAAACGCCTGGTATCTTTATAGTCCTGTCGGGTTTCGCCACCTCTGACTTGAGCGT
CGATTTTTGTGATGCTCGTCAGGGGGGCGGAGCCTATGGAAAAACGCCAGCAACGCGGCCTTTTTACGGTTCCTGGCCTT
TTGCTGGCCTTTTGCTCACATGTTCTTTCCTGCGTTATCCCCTGATTCTGTGGATAACCGTATTACCGCCTTTGAGTGAG
CTGATACCGCTCGCCGCAGCCGAACGACCGAGCGCAGCGAGTCAGTGAGCGAGGAAGCGGAAGAGCGCCTGATGCGGTAT
TTTCTCCTTACGCATCTGTGCGGTATTTCACACCGCATATATGGTGCACTCTCAGTACAATCTGCTCTGATGCCGCATAG
TTAAGCCAGTATACACTCCGCTATCGCTACGTGACTGGGTCATGGCTGCGCCCCGACACCCGCCAACACCCGCTGACGCG
CCCTGACGGGCTTGTCTGCTCCCGGCATCCGCTTACAGACAAGCTGTGACCGTCTCCGGGAGCTGCATGTGTCAGAGGTT
TTCACCGTCATCACCGAAACGCGCGAGGCAGCTGCGGTAAAGCTCATCAGCGTGGTCGTGAAGCGATTCACAGATGTCTG
CCTGTTCATCCGCGTCCAGCTCGTTGAGTTTCTCCAGAAGCGTTAATGTCTGGCTTCTGATAAAGCGGGCCATGTTAAGG
GCGGTTTTTTCCTGTTTGGTCACTGATGCCTCCGTGTAAGGGGGATTTCTGTTCATGGGGGTAATGATACCGATGAAACG
AGAGAGGATGCTCACGATACGGGTTACTGATGATGAACATGCCCGGTTACTGGAACGTTGTGAGGGTAAACAACTGGCGG
TATGGATGCGGCGGGACCAGAGAAAAATCACTCAGGGTCAATGCCAGCGCTTCGTTAATACAGATGTAGGTGTTCCACAG
GGTAGCCAGCAGCATCCTGCGATGCAGATCCGGAACATAATGGTGCAGGGCGCTGACTTCCGCGTTTCCAGACTTTACGA
AACACGGAAACCGAAGACCATTCATGTTGTTGCTCAGGTCGCAGACGTTTTGCAGCAGCAGTCGCTTCACGTTCGCTCGC
GTATCGGTGATTCATTCTGCTAACCAGTAAGGCAACCCCGCCAGCCTAGCCGGGTCCTCAACGACAGGAGCACGATCATG
CGCACCCGTGGGGCCGCCATGCCGGCGATAATGGCCTGCTTCTCGCCGAAACGTTTGGTGGCGGGACCAGTGACGAAGGC
TTGAGCGAGGGCGTGCAAGATTCCGAATACCGCAAGCGACAGGCCGATCATCGTCGCGCTCCAGCGAAAGCGGTCCTCGC
CGAAAATGACCCAGAGCGCTGCCGGCACCTGTCCTACGAGTTGCATGATAAAGAAGACAGTCATAAGTGCGGCGACGATA
GTCATGCCCCGCGCCCACCGGAAGGAGCTGACTGGGTTGAAGGCTCTCAAGGGCATCGGTCGAGATCCCGGTGCCTAATG

AGTGAGCTAACTTACATTAATTGCGTTGCGCTCACTGCCCGCTTTCCAGTCGGGAAACCTGTCGTGCCAGCTGCATTAAT
GAATCGGCCAACGCGCGGGGAGAGGCGGTTTGCGTATTGGGCGCCAGGGTGGTTTTTCTTTTCACCAGTGAGACGGGCAA
CAGCTGATTGCCCTTCACCGCCTGGCCCTGAGAGAGTTGCAGCAAGCGGTCCACGCTGGTTTGCCCCAGCAGGCGAAAAT
CCTGTTTGATGGTGGTTAACGGCGGGATATAACATGAGCTGTCTTCGGTATCGTCGTATCCCACTACCGAGATATCCGCA
CCAACGCGCAGCCCGGACTCGGTAATGGCGCGCATTGCGCCCAGCGCCATCTGATCGTTGGCAACCAGCATCGCAGTGGG
AACGATGCCCTCATTCAGCATTTGCATGGTTTGTTGAAAACCGGACATGGCACTCCAGTCGCCTTCCCGTTCCGCTATCG
GCTGAATTTGATTGCGAGTGAGATATTTATGCCAGCCAGCCAGACGCAGACGCGCCGAGACAGAACTTAATGGGCCCGCT
AACAGCGCGATTTGCTGGTGACCCAATGCGACCAGATGCTCCACGCCCAGTCGCGTACCGTCTTCATGGGAGAAAATAAT
ACTGTTGATGGGTGTCTGGTCAGAGACATCAAGAAATAACGCCGGAACATTAGTGCAGGCAGCTTCCACAGCAATGGCAT
CCTGGTCATCCAGCGGATAGTTAATGATCAGCCCACTGACGCGTTGCGCGAGAAGATTGTGCACCGCCGCTTTACAGGCT
TCGACGCCGCTTCGTTCTACCATCGACACCACCACGCTGGCACCCAGTTGATCGGCGCGAGATTTAATCGCCGCGACAAT
TTGCGACGGCGCGTGCAGGGCCAGACTGGAGGTGGCAACGCCAATCAGCAACGACTGTTTGCCCGCCAGTTGTTGTGCCA
CGCGGTTGGGAATGTAATTCAGCTCCGCCATCGCCGCTTCCACTTTTTCCCGCGTTTTCGCAGAAACGTGGCTGGCCTGG
TTCACCACGCGGGAAACGGTCTGATAAGAGACACCGGCATACTCTGCGACATCGTATAACGTTACTGGTTTCACATTCAC
CACCCTGAATTGACTCTCTTCCGGGCGCTATCATGCCATACCGCGAAAGGTTTTGCGCCATTCGATGGTGTCCGGGATCT
CGACGCTCTCCCTTATGCGACTCCTGCATTAGGAAGCAGCCCAGTAGTAGGTTGAGGCCGTTGAGCACCGCCGCCGCAAG
GAATGGTGCATGCAAGGAGATGGCGCCCAACAGTCCCCCGGCCACGGGGCCTGCCACCATACCCACGCCGAAACAAGCGC
TCATGAGCCCGAAGTGGCGAGCCCGATCTTCCCCATCGGTGATGTCGGCGATATAG
>pFS06
GCGCCAGCAACCGCACCTGTGGCGCCGGTGATGCCGGCCACGATGCGTCCGGCGTAGAGGATCGAGATCTCGATCCCGCG
AAATTAATACGACTCACTATAGGGGAATTGTGAGCGGATAACAATTCCCCTCTAGAAATAATTTTGTTTAACTTTAAGAA
GGAGATATACCATGGGCAGCAGCCATCATCATCATCATCACAGCAGCGGCCTGGTGCCGCGCGGCAGCCATATGGCTAGC
ATGACTGGTGGACAGCAAATGGGTCGCGGATCCATGAGCGCAGCAGCAAATGCAGAACATGGTGCAGCAGATCGTGTTGA
AATTCTGCCGGTTCCGGGTCTGCCGGAATTTCGTCCGGGTGATGATCTGGTTGGTAGCCTGGCCGAAGCAGCACCGTGGC
TGCGTGATGGTGATGTGCTGGTTGTTACCAGCAAAGTTGTTAGCAAATGTGAAGGTCGTATTGTTGCAGCACCGAGCGAT
CCGGAAGAACGTGATACCCTGCGTCGTAAACTGATTGATGATGAAGCAGTTCGTGTTCTGGCACGTAAAGGTCGTACCCT
GATTACCGAAAATGCAATTGGTCTGGTTCAGGCAGCAGCCGGTGTTGATGGTAGCAATGTTGGTAGCACCGAACTGGCAC
TGCTGCCGGTTGATCCTGATCGTAGCGCAGCAACCCTGCGTGAAGGTCTGCGTGAACGCCTGGGTGTTACCGTTGGTGTT
GTTATTACCGATACCATGGGTCGTGCATGGCGTACCGGTCAGACCGATTTTGCCATTGGTGCAAGCGGTCTGACAGTTCT
GCAGGGTTATGCAGGTAGCCGTGATCGTCATGGTAATGAACTGGTTGTGACCGAAGTTGCAGTTGCAGATGAAATTGCAG
CAGCAGCGGATCTGGTTAAAGGTAAACTGACCGCAATTCCGGTTGCCGTTGTTCGTGGTCTGCGCCTGCCGGATGATGGT
AGTACCGCACATCGTCTGGTTCGTGCCGGTGAAGATGACCTGTTTTGGCTGGGCACCGCAGAAGCAATTGAACTGGGTCG
TCGTCAGGCACAGCTGCTGCGTCGTTCAGTTCGTCGTTTTAGCGCAGAACCGGTGCCGCATGATGCCATTGAAGCAGCAG
TTGGTGAAGCCCTGACCGCACCGGCACCGCATCATACCCGTCCGGTTCGTTTTGTTTGGGTTCAAGATAGCGAAACCCGT
ACACGTCTGCTGGATCGTATGAAAGAACAGTGGCGTGCAGATCTGACCGCAGATGGTCTGGATGCAGATGCAGTTGATCG
TCGTGTTGCACGTGGTCAGATTCTGTATGATGCACCGGAACTGGTTATTCCGTTTCTGGTTCCGGATGGTGCACATAGCT
ATCCTGATGATGCACGTACCGCAGCAGAACATACCATGTTTACCGTTGCAGTGGGTGCAGCCGTTCAGGGTCTGCTGGTT
GCACTGGCCGTTCGTGATATTGGTAGCTGTTGGATTGGTTCAACCATTTTTGCCGCAGATCTGGTTCGCGCAGAACTGGA
ACTGCCTGATGATTGGGAACCGCTGGGTGCAATTGCGATTGGTTATCCGGAACAGACACCGCAGCCGCTGGGTCCGCGTG
ATCCGGTTCCGACCGATGAACTGCTGGTTCGTAAATAAGAATTCGAGCTCCGTCGACAAGCTTGCGGCCGCACTCGAGCA
CCACCACCACCACCACTGAGATCCGGCTGCTAACAAAGCCCGAAAGGAAGCTGAGTTGGCTGCTGCCACCGCTGAGCAAT
AACTAGCATAACCCCTTGGGGCCTCTAAACGGGTCTTGAGGGGTTTTTTGCTGAAAGGAGGAACTATATCCGGATTGGCG
AATGGGACGCGCCCTGTAGCGGCGCATTAAGCGCGGCGGGTGTGGTGGTTACGCGCAGCGTGACCGCTACACTTGCCAGC
GCCCTAGCGCCCGCTCCTTTCGCTTTCTTCCCTTCCTTTCTCGCCACGTTCGCCGGCTTTCCCCGTCAAGCTCTAAATCG
GGGGCTCCCTTTAGGGTTCCGATTTAGTGCTTTACGGCACCTCGACCCCAAAAAACTTGATTAGGGTGATGGTTCACGTA
GTGGGCCATCGCCCTGATAGACGGTTTTTCGCCCTTTGACGTTGGAGTCCACGTTCTTTAATAGTGGACTCTTGTTCCAA
ACTGGAACAACACTCAACCCTATCTCGGTCTATTCTTTTGATTTATAAGGGATTTTGCCGATTTCGGCCTATTGGTTAAA
AAATGAGCTGATTTAACAAAAATTTAACGCGAATTTTAACAAAATATTAACGTTTACAATTTCAGGTGGCACTTTTCGGG
GAAATGTGCGCGGAACCCCTATTTGTTTATTTTTCTAAATACATTCAAATATGTATCCGCTCATGAATTAATTCTTAGAA
AAACTCATCGAGCATCAAATGAAACTGCAATTTATTCATATCAGGATTATCAATACCATATTTTTGAAAAAGCCGTTTCT
GTAATGAAGGAGAAAACTCACCGAGGCAGTTCCATAGGATGGCAAGATCCTGGTATCGGTCTGCGATTCCGACTCGTCCA
ACATCAATACAACCTATTAATTTCCCCTCGTCAAAAATAAGGTTATCAAGTGAGAAATCACCATGAGTGACGACTGAATC
CGGTGAGAATGGCAAAAGTTTATGCATTTCTTTCCAGACTTGTTCAACAGGCCAGCCATTACGCTCGTCATCAAAATCAC
TCGCATCAACCAAACCGTTATTCATTCGTGATTGCGCCTGAGCGAGACGAAATACGCGATCGCTGTTAAAAGGACAATTA
CAAACAGGAATCGAATGCAACCGGCGCAGGAACACTGCCAGCGCATCAACAATATTTTCACCTGAATCAGGATATTCTTC
TAATACCTGGAATGCTGTTTTCCCGGGGATCGCAGTGGTGAGTAACCATGCATCATCAGGAGTACGGATAAAATGCTTGA
TGGTCGGAAGAGGCATAAATTCCGTCAGCCAGTTTAGTCTGACCATCTCATCTGTAACATCATTGGCAACGCTACCTTTG
CCATGTTTCAGAAACAACTCTGGCGCATCGGGCTTCCCATACAATCGATAGATTGTCGCACCTGATTGCCCGACATTATC
GCGAGCCCATTTATACCCATATAAATCAGCATCCATGTTGGAATTTAATCGCGGCCTAGAGCAAGACGTTTCCCGTTGAA
TATGGCTCATAACACCCCTTGTATTACTGTTTATGTAAGCAGACAGTTTTATTGTTCATGACCAAAATCCCTTAACGTGA
GTTTTCGTTCCACTGAGCGTCAGACCCCGTAGAAAAGATCAAAGGATCTTCTTGAGATCCTTTTTTTCTGCGCGTAATCT
GCTGCTTGCAAACAAAAAAACCACCGCTACCAGCGGTGGTTTGTTTGCCGGATCAAGAGCTACCAACTCTTTTTCCGAAG
GTAACTGGCTTCAGCAGAGCGCAGATACCAAATACTGTCCTTCTAGTGTAGCCGTAGTTAGGCCACCACTTCAAGAACTC
TGTAGCACCGCCTACATACCTCGCTCTGCTAATCCTGTTACCAGTGGCTGCTGCCAGTGGCGATAAGTCGTGTCTTACCG

GGTTGGACTCAAGACGATAGTTACCGGATAAGGCGCAGCGGTCGGGCTGAACGGGGGGTTCGTGCACACAGCCCAGCTTG
GAGCGAACGACCTACACCGAACTGAGATACCTACAGCGTGAGCTATGAGAAAGCGCCACGCTTCCCGAAGGGAGAAAGGC
GGACAGGTATCCGGTAAGCGGCAGGGTCGGAACAGGAGAGCGCACGAGGGAGCTTCCAGGGGGAAACGCCTGGTATCTTT
ATAGTCCTGTCGGGTTTCGCCACCTCTGACTTGAGCGTCGATTTTTGTGATGCTCGTCAGGGGGGCGGAGCCTATGGAAA
AACGCCAGCAACGCGGCCTTTTTACGGTTCCTGGCCTTTTGCTGGCCTTTTGCTCACATGTTCTTTCCTGCGTTATCCCC
TGATTCTGTGGATAACCGTATTACCGCCTTTGAGTGAGCTGATACCGCTCGCCGCAGCCGAACGACCGAGCGCAGCGAGT
CAGTGAGCGAGGAAGCGGAAGAGCGCCTGATGCGGTATTTTCTCCTTACGCATCTGTGCGGTATTTCACACCGCATATAT
GGTGCACTCTCAGTACAATCTGCTCTGATGCCGCATAGTTAAGCCAGTATACACTCCGCTATCGCTACGTGACTGGGTCA
TGGCTGCGCCCCGACACCCGCCAACACCCGCTGACGCGCCCTGACGGGCTTGTCTGCTCCCGGCATCCGCTTACAGACAA
GCTGTGACCGTCTCCGGGAGCTGCATGTGTCAGAGGTTTTCACCGTCATCACCGAAACGCGCGAGGCAGCTGCGGTAAAG
CTCATCAGCGTGGTCGTGAAGCGATTCACAGATGTCTGCCTGTTCATCCGCGTCCAGCTCGTTGAGTTTCTCCAGAAGCG
TTAATGTCTGGCTTCTGATAAAGCGGGCCATGTTAAGGGCGGTTTTTTCCTGTTTGGTCACTGATGCCTCCGTGTAAGGG
GGATTTCTGTTCATGGGGGTAATGATACCGATGAAACGAGAGAGGATGCTCACGATACGGGTTACTGATGATGAACATGC
CCGGTTACTGGAACGTTGTGAGGGTAAACAACTGGCGGTATGGATGCGGCGGGACCAGAGAAAAATCACTCAGGGTCAAT
GCCAGCGCTTCGTTAATACAGATGTAGGTGTTCCACAGGGTAGCCAGCAGCATCCTGCGATGCAGATCCGGAACATAATG
GTGCAGGGCGCTGACTTCCGCGTTTCCAGACTTTACGAAACACGGAAACCGAAGACCATTCATGTTGTTGCTCAGGTCGC
AGACGTTTTGCAGCAGCAGTCGCTTCACGTTCGCTCGCGTATCGGTGATTCATTCTGCTAACCAGTAAGGCAACCCCGCC
AGCCTAGCCGGGTCCTCAACGACAGGAGCACGATCATGCGCACCCGTGGGGCCGCCATGCCGGCGATAATGGCCTGCTTC
TCGCCGAAACGTTTGGTGGCGGGACCAGTGACGAAGGCTTGAGCGAGGGCGTGCAAGATTCCGAATACCGCAAGCGACAG
GCCGATCATCGTCGCGCTCCAGCGAAAGCGGTCCTCGCCGAAAATGACCCAGAGCGCTGCCGGCACCTGTCCTACGAGTT
GCATGATAAAGAAGACAGTCATAAGTGCGGCGACGATAGTCATGCCCCGCGCCCACCGGAAGGAGCTGACTGGGTTGAAG
GCTCTCAAGGGCATCGGTCGAGATCCCGGTGCCTAATGAGTGAGCTAACTTACATTAATTGCGTTGCGCTCACTGCCCGC
TTTCCAGTCGGGAAACCTGTCGTGCCAGCTGCATTAATGAATCGGCCAACGCGCGGGGAGAGGCGGTTTGCGTATTGGGC
GCCAGGGTGGTTTTTCTTTTCACCAGTGAGACGGGCAACAGCTGATTGCCCTTCACCGCCTGGCCCTGAGAGAGTTGCAG
CAAGCGGTCCACGCTGGTTTGCCCCAGCAGGCGAAAATCCTGTTTGATGGTGGTTAACGGCGGGATATAACATGAGCTGT
CTTCGGTATCGTCGTATCCCACTACCGAGATATCCGCACCAACGCGCAGCCCGGACTCGGTAATGGCGCGCATTGCGCCC
AGCGCCATCTGATCGTTGGCAACCAGCATCGCAGTGGGAACGATGCCCTCATTCAGCATTTGCATGGTTTGTTGAAAACC
GGACATGGCACTCCAGTCGCCTTCCCGTTCCGCTATCGGCTGAATTTGATTGCGAGTGAGATATTTATGCCAGCCAGCCA
GACGCAGACGCGCCGAGACAGAACTTAATGGGCCCGCTAACAGCGCGATTTGCTGGTGACCCAATGCGACCAGATGCTCC
ACGCCCAGTCGCGTACCGTCTTCATGGGAGAAAATAATACTGTTGATGGGTGTCTGGTCAGAGACATCAAGAAATAACGC
CGGAACATTAGTGCAGGCAGCTTCCACAGCAATGGCATCCTGGTCATCCAGCGGATAGTTAATGATCAGCCCACTGACGC
GTTGCGCGAGAAGATTGTGCACCGCCGCTTTACAGGCTTCGACGCCGCTTCGTTCTACCATCGACACCACCACGCTGGCA
CCCAGTTGATCGGCGCGAGATTTAATCGCCGCGACAATTTGCGACGGCGCGTGCAGGGCCAGACTGGAGGTGGCAACGCC
AATCAGCAACGACTGTTTGCCCGCCAGTTGTTGTGCCACGCGGTTGGGAATGTAATTCAGCTCCGCCATCGCCGCTTCCA
CTTTTTCCCGCGTTTTCGCAGAAACGTGGCTGGCCTGGTTCACCACGCGGGAAACGGTCTGATAAGAGACACCGGCATAC
TCTGCGACATCGTATAACGTTACTGGTTTCACATTCACCACCCTGAATTGACTCTCTTCCGGGCGCTATCATGCCATACC
GCGAAAGGTTTTGCGCCATTCGATGGTGTCCGGGATCTCGACGCTCTCCCTTATGCGACTCCTGCATTAGGAAGCAGCCC
AGTAGTAGGTTGAGGCCGTTGAGCACCGCCGCCGCAAGGAATGGTGCATGCAAGGAGATGGCGCCCAACAGTCCCCCGGC
CACGGGGCCTGCCACCATACCCACGCCGAAACAAGCGCTCATGAGCCCGAAGTGGCGAGCCCGATCTTCCCCATCGGTGA
TGTCGGCGATATAG
